# An experimental census of retrons for DNA production and genome editing

**DOI:** 10.1101/2024.01.25.577267

**Authors:** Asim G. Khan, Matías Rojas-Montero, Alejandro González-Delgado, Santiago C. Lopez, Rebecca F. Fang, Seth L. Shipman

## Abstract

Retrons are bacterial immune systems that use reverse transcribed DNA as a detector of phage infection. They are also increasingly deployed as a component of biotechnology. For genome editing, for instance, retrons are modified so that the reverse transcribed DNA (RT-DNA) encodes an editing donor. Retrons are commonly found in bacterial genomes; thousands of unique retrons have now been predicted bioinformatically. However, only a small number have been characterized experimentally. Here, we add substantially to the corpus of experimentally studied retrons. We synthesized >100 previously untested retrons to identify the natural sequence of RT-DNA they produce, quantify their RT-DNA production, and test the relative efficacy of editing using retron-derived donors to edit bacterial genomes, phage genomes, and human genomes. We add 62 new empirically determined, natural RT-DNAs, which are not predictable from the retron sequence alone. We report a large diversity in RT-DNA production and editing rates across retrons, finding that top performing editors outperform those used in previous studies, and are drawn from a subset of the retron phylogeny.

## INTRODUCTION

The signature of a retron – copious single-stranded DNA of a uniform length produced spontaneously by bacteria – was first observed in 1984^1^. Five years later, it was shown that this DNA was the product of an endogenous reverse transcriptase (RT), the first RT discovered in prokaryotes^2^. The template for the reverse transcribed DNA (RT-DNA) of a retron is a short, highly structured RNA, which is produced from the same operon as the retron RT. The RT recognizes this structured, noncoding RNA (ncRNA) and partially reverse transcribes it into RT-DNA, polymerizing from a critical 2’ hydroxyl of the ncRNA^3,4^. The result is a transcriptional/reverse-transcriptional cascade that amplifies a single locus in the bacterial genome up to hundreds of copies of RT-DNA per cell^5^. This phenomenon of RT-DNA production was observed from 16 natural retrons over the next thirty years, without any conclusive evidence for a cellular function^6^.

More recently, a cellular role for retrons has become clear. Retrons confer phage resistance to bacteria^7–10^. Full mechanistic details are still emerging, but the general model is that retron RT-DNA is a sensor of phage infection. Either directly or indirectly, the phage modifies or degrades the RT-DNA, which releases an accessory retron protein that acts as a cellular toxin to remove the infected cell and spare the bacterial population^8,9,11^. Individual retrons have been shown to confer specific resistance to particular phages^8,12^. This specificity is due, at least in part, to the unique sequence of RT-DNA produced by the retron^8^. More than a thousand retrons have now been identified in genomic and metagenomic databases by homology to known retron RTs^10^. However, the sequence of the ncRNA has remained difficult to identify bioinformatically and the exact RT-DNA produced by each retron is, thus far, impossible to predict.

Retrons, like many other bacterial immune systems, have also proven valuable as a component of biotechnology. Specifically, the faithful production of abundant single-stranded DNA in cells by retrons has been used to produce templates for genome engineering, transcriptional receipts for molecular recorders, and transcription factor decoys^13–17^. Yet, to date, these technologies have been built using only a small number of the early-discovered retrons.

It is critical that we survey the diversity of retrons, both to inform investigations of bacterial immunity and to improve retron-based technologies. Of the thousand plus retrons identified^10^, only eighteen have actually been shown to produce RT-DNA^6,8,9^, only one has been used for bacterial recombineering^13,16,18^, and only eight have been tested for human genome editing^18–20^. Here, we address the lack of experimental testing across the diversity of predicted retrons by undertaking a census of RT-DNA production and editing in multiple cellular contexts. We provide validation of noncoding RNA sequence and RT-DNA production for retrons across the phylogeny, and report sequences of the unpredictable RT-DNA that are critical to inform new studies of function and technology. We identify the first subsets of retrons that appear to lack RT-DNA production, which will also inform future work in the field. We also find that particular subtypes of retrons produce higher amounts of RT-DNA and better editing performance.

## RESULTS

In order to expand the corpus of experimentally validated retrons, we synthesized 163 never-before-tested retron RTs and ncRNAs distributed across various experiment types to enable quantification of RT-DNA production, as well as editing in bacterial, phage, and human genomes. These retrons were drawn from a set of 1,928 recently published retrons^10^ that were bioinformatically predicted based on protein homology to known retron RTs. While the RT component was previously annotated, the ncRNA component of retrons is difficult to predict across diverse retrons. We identified and annotated the cognate ncRNAs of each RT by simulating RNA folding in regions within 500bp of the RT and comparing the overall secondary structure to the consensus ncRNA structures. This consists of conserved characteristics such as an inverted repeat (the a1/a2 region of a retron ncRNA) >8bp, a priming guanine near the beginning of the ncRNA, and multiple hairpins in close proximity to a retron RT. Retrons have been recently shown to engage in phage defense that requires one or more additional accessory proteins, multiple of which have been shown to be cellular toxins^8,9,12^. The accessory proteins and phage defense mechanisms are extremely diverse among retrons^10^. For this work, we chose to focus on a census of the core reverse transcription machinery and use in biotechnology and thus excluded accessory genes from our analysis. However, it is important to note that in some cases the retron RT is fused to accessory domains and, in these instances, we did not truncate the protein.

### RT-DNA production by a diverse set of retrons

For testing RT-DNA production, we synthesized 98 new retron RTs and ncRNAs, and one known standard, retron-Eco1 (ec86). These retrons were chosen to give a broad representation of the RT phylogeny (**Fig 1a**), including retrons from each RT clade (**Fig 1b**). Retron ncRNAs are frequently found immediately upstream of the retron RT. However, in this natural arrangement, putative ribosomal binding sites can be contained within the ncRNA, which makes standardization impossible. Therefore, we chose to invert the architecture and create operons driven by a T7/lac promoter, followed by a strong RBS, the RT coding region, and finally the ncRNA (**Fig 1c**). In this arrangement, no additional surrounding sequence other than the inverted repeat a1/a2 regions are present with the ncRNA, so in cases where an RT-DNA is produced, we can be sure that we have rigorously identified the ncRNA.

**Figure 1.**
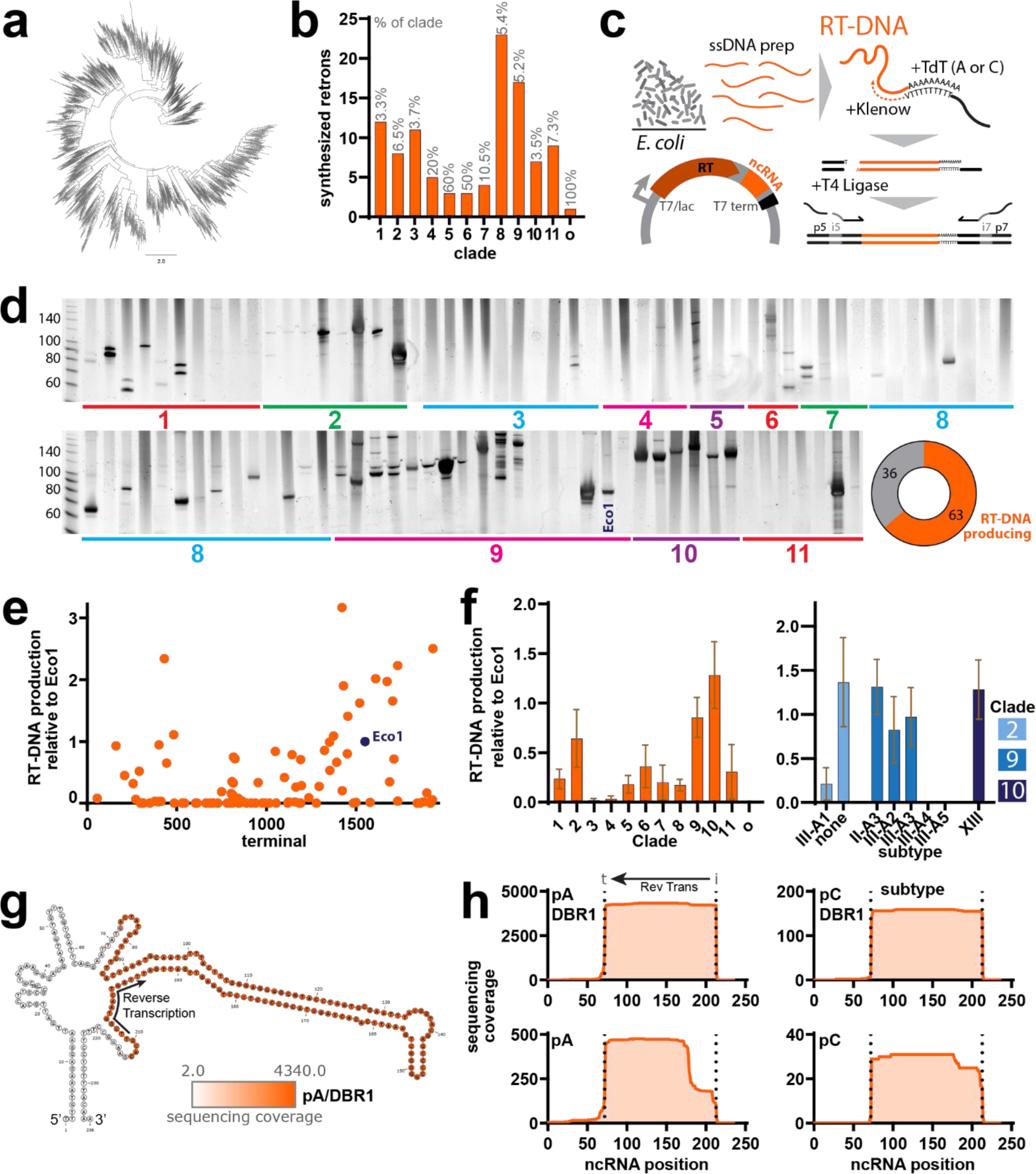
RT-DNA production by a diverse set of retrons. **a.** Phylogenic analysis of retron RTs^10^. **b.** Number of retrons synthesized by clade (percentage of clade synthesized indicated in gray above each bar). Bar o indicates an orphan, clade-less retron. **c.** Schematic of RT-DNA quantification experiment. Bottom left shows the architecture of retrons for expression, right shows strategy for unbiased sequencing of RT-DNA. **d.** PAGE analysis of all retrons tested (composite of multiple gels, uncropped gels in **Supplemental** Figure 6) and proportion of retrons with detectable RT-DNA (bottom right donut). **e.** Quantification of RT-DNA production by density, relative to retron-Eco1 (marked in blue) arranged by order (terminal number) in the RT phylogeny shown in 1a. **f.** RT-DNA production by clade (one-way ANOVA, effect of clade *P*=0.0003). Bar o indicates an orphan, clade-less retron. Plot to the right splits production of the top clades by retron subtype. Bars show mean ±SEM. **g.** RT-DNA sequencing coverage plotted onto folded ncRNA for retron Mestre-1673. Orange indicates the bases that are reverse transcribed. **h.** Plots of RT-DNA sequencing coverage by ncRNA position for retron Mestre-1673. Conditions include pA or pC tailing of the RT-DNA for sequencing prep and inclusion or exclusion of DBR1 prior to sequencing prep. Dashed lines indicate the initiation and termination points, defined from the pA/DBR1 condition (initiation: coverage increases to ≥15% of maximum; termination: coverage decreases to ≤50% of maximum). Additional statistical details in **Supplemental Table 2**.

We transformed each of these 99 retrons (98 new plus one reference standard) into BL21-AI *E. coli* and induced expression of the synthetic operon for five hours. We prepared non-genomic nucleotides by midiprep and visualized these nucleotides on a TBE/Urea gel to quantify the presence of a characteristic RT-DNA band. Next, we prepared any resulting RT-DNA for sequencing. Because the retron RT-DNA sequences were unknown to us, we adapted an unbiased sequencing approach that we have previously used to examine Eco1 variant libraries^12^. Briefly, we added a single polynucleotide handle to the 5’ end of RT-DNA using TdT, used a complementary polynucleotide anchored primer and Klenow to synthesize a strand complementary to the RT-DNA, and then ligated adapters at the 3’ end of the now double-stranded RT-DNA (**Fig 1c**). The resulting molecules were indexed and sequenced on an Illumina machine.

We found that 62 of the 98 newly tested retrons produced RT-DNA that were detectable on a TBE/Urea gel, which varied in both length and abundance (**Fig 1d**). We quantified the intensity of these bands relative to a standard, retron-Eco1. When these relative production values are plotted by phylogenetic terminal, we observed that retrons in certain regions of the phylogeny produce more RT-DNA than others (**Fig 1e**). When examined by RT clade, we see greatest production from clades 2, 9, and 10; almost no production from clades 3 and 4; and intermediate production from the rest (**Fig 1f**). This can be further broken down by retron subtype – which is based on features of the accessory proteins rather than the RT itself – within the high producing clades. Here, we see that within clades 2 and 9, the highest producing retrons are confined to certain subtypes. This subtype dependence is also evident in some lower producing clades (**Supplementary** Fig 1).

We next used the sequencing data to determine the sequence of RT-DNA from retrons that produced it. We aligned RT-DNA sequences determined from the blind prep of single-stranded DNA to each retron ncRNA. An example of sequencing coverage is shown for Mestre-1673, plotting coverage onto the secondary structure of the ncRNA (**Fig 1g**). Each RT-DNA was prepped in parallel using four similar approaches, varying two parameters. First, we varied the nucleotide that we used to extend the sequence (poly adenine, pA, or poly cytosine, pC) for unbiased sequencing as TdT polymerase has nucleotide preferences that could affect the efficiency of the prep based on the terminating nucleotide of the RT-DNA. Second, we varied whether we pretreated the nucleotides with a debranching enzyme (DBR1) prior to sequencing prep. The characteristic 2’-5’ RNA to DNA linkage created when retron RTs polymerize from a structured RNA inhibits our sequencing preparation. For some retrons, this branched molecule is long lasting and DBR1 is required for efficient preparation. However, other retrons are debranched *in vivo* by nucleases that cleave in the RT-DNA. In these cases, DBR1 is not required for efficient sequencing preparation. By including both conditions, we can estimate whether a given retron remains branched or is debranched in vivo.

Figure 1h shows each of these four conditions for Mestre-1673, with sequencing coverage plotted over linear ncRNA position. In the pA/DBR1 condition (also shown in Figure 1g), we see even coverage of the RT-DNA region of the ncRNA. We define the initiation point (dashed line i) as the point where coverage increases to ≥15% of maximum coverage and termination (dashed line t) as the point where coverage decreases to ≤50% of maximum coverage. These initiation and termination points are propagated to the other conditions. In this case the pC/DBR1 condition closely matches the pA, but with fewer reads, which is typical of all RT-DNA sequencing. The conditions below, which lack DBR1 addition during sequencing prep, have substantially fewer reads. This is consistent with a retron that remains 2’-5’ branched *in vivo*. These coverage maps also illustrate a pattern that occurs frequently among retrons. In the absence of DBR1, the 5’ end of the RT-DNA begins further from the branched initiation point. This likely represents a fraction of RT-DNA that was debranched *in vivo* by cleavage of the RT-DNA at a position 3’ to the branch point or partial nuclease degradation from the free 5’ end of a debranched fraction.

### Characteristics of RT-DNA production across retrons

We generated RT-DNA coverage maps for 54 retrons. Sequencing coverage (RT-DNA) plotted onto retron ncRNA secondary structures is shown for a subset of retrons in Figure 2a. Coverage and sequencing data for all retrons is provided in accompanying supplemental data (**Supplemental Data 1**). When quantified across retrons, the initiation point is typically quite stringent, with RT-DNA sequencing coverage rising from 15% of maximum to 85% of maximum within 3 nucleotides for >65% of retrons tested (Fig 2b). In contrast, termination tended to occur over a wider window, suggesting more variability in the termination point or some degradation of the 3’ end after termination (Fig 2c). The branching nucleotide of the ncRNA from which the RT-DNA is built has been found to be most frequently a guanine often preceded by TTA, which we also see in our set of functional retrons (**Supplemental** Fig 2). We find that the first reverse transcribed base is also most frequently a guanine and that the termination point of RT-DNA typically precedes an AT rich region of the ncRNA (Fig 2d). The length of the ncRNA varies across retrons, and is correlated with the length of the RT-DNA (Fig 2e). We can use the ratio of DBR1+ sequencing reads to DBR1-sequencing reads to infer which retrons remain branched *in vivo*. Retrons with similar numbers of reads regardless of the presence or absence of DBR1 in the sequencing prep are likely unbranched already *in vivo*, whereas retrons with many more reads in the DBR1+ condition likely remain branched *in vivo* (Fig 2f). Finally, this RT-DNA sequencing data can be harnessed to estimate the relative fidelity of the retrons by quantifying the errors per base in the RT-DNA with respect to the retron’s ncRNA reference. Critically, these are not absolute error rates of the RT as this includes errors introduced by sequencing prep or Illumina sequencing. We find that most retrons are of a similar fidelity, with just a few retrons making ∼5x greater errors (Fig 2g).

**Figure 2.**
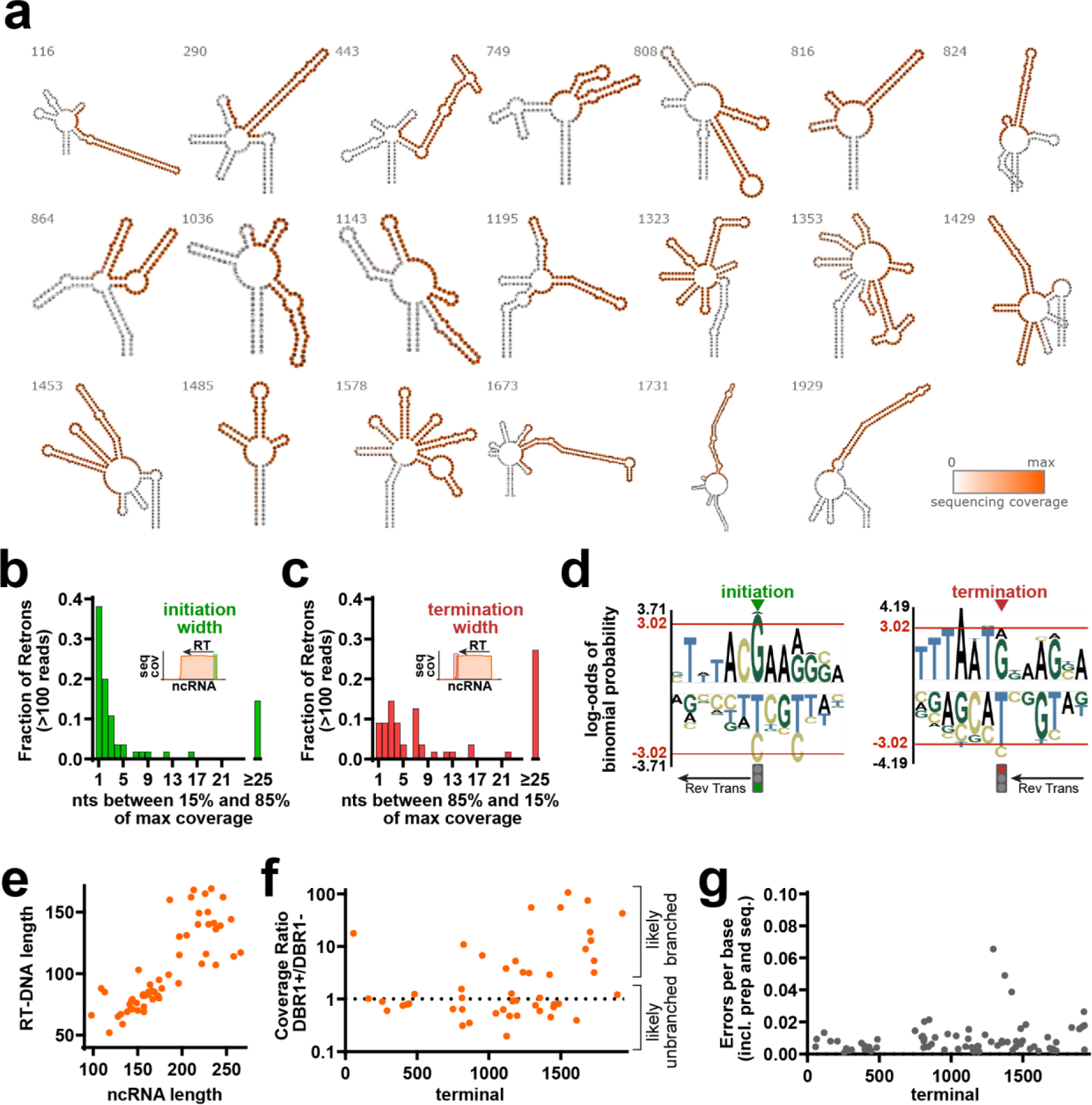
Characteristics of RT-DNA production across retrons. **a.** Examples of RT-DNA sequencing coverage plotted on folded ncRNA for a subset of sequenced retrons (additional data in **Supplemental Data 1**). **b.** Strictness of initiation. For all retrons with at least 100 sequencing reads, number of nucleotides at the RT initiation point between 15% an 85% of maximum coverage. **c.** Strictness of termination. For all retrons with at least 100 sequencing reads, number of nucleotides at the RT termination point from 85% to 15% of maximum coverage. **d.** Probability logos showing overrepresented (positive) and underrepresented (negative) nucleotides at the initiation (left) and termination (right) points. Red line indicates *P*<0.05. **e.** RT-DNA length versus ncRNA length (Pearson Correlation, r^2^=0.6586, *P*<0.0001). **f.** Ratio of the sequencing coverage in the +DBR1 condition versus the –DBR1 condition for each retron. Retrons with a high ratio are more likely to exist in a 2’-5’ branched form with their msr, which requires DBR1 for efficient sequencing, whereas retrons with ratios around or under 1 are equally sequenced in the absence or presence of DBR1 indication that they were not 2’-5’ branched. **g.** Errors per base versus the ncRNA reference for retrons with at least 1,000 sequenced bases, calculated using Levenshtein Distance. Errors include fidelity of the RT, but also errors introduced by the sequencing prep and sequencing. Additional statistical details in **Supplemental Table 2**.

### Surveying retrons for bacterial and phage recombineering

The ability of retrons to continuously produce a ssDNA containing a precise target mutation has been recently exploited for recombineering purposes in both bacteria and phages^13,16,18,21–23^. Retron-based recombineering relies on the reverse transcription of a modified RT-DNA that contains an editing donor (Fig 3a). The host single-stranded binding protein (SSB) will bind the resulting RT-Donor to promote its interaction with a single-stranded annealing protein (SSAP, e.g. CspRecT), which leads to installation of the edit in the lagging strand during chromosome replication^24^. However, only retron-Eco1 has been extensively characterized and optimized for use in prokaryotic genome engineering.

**Figure 3.**
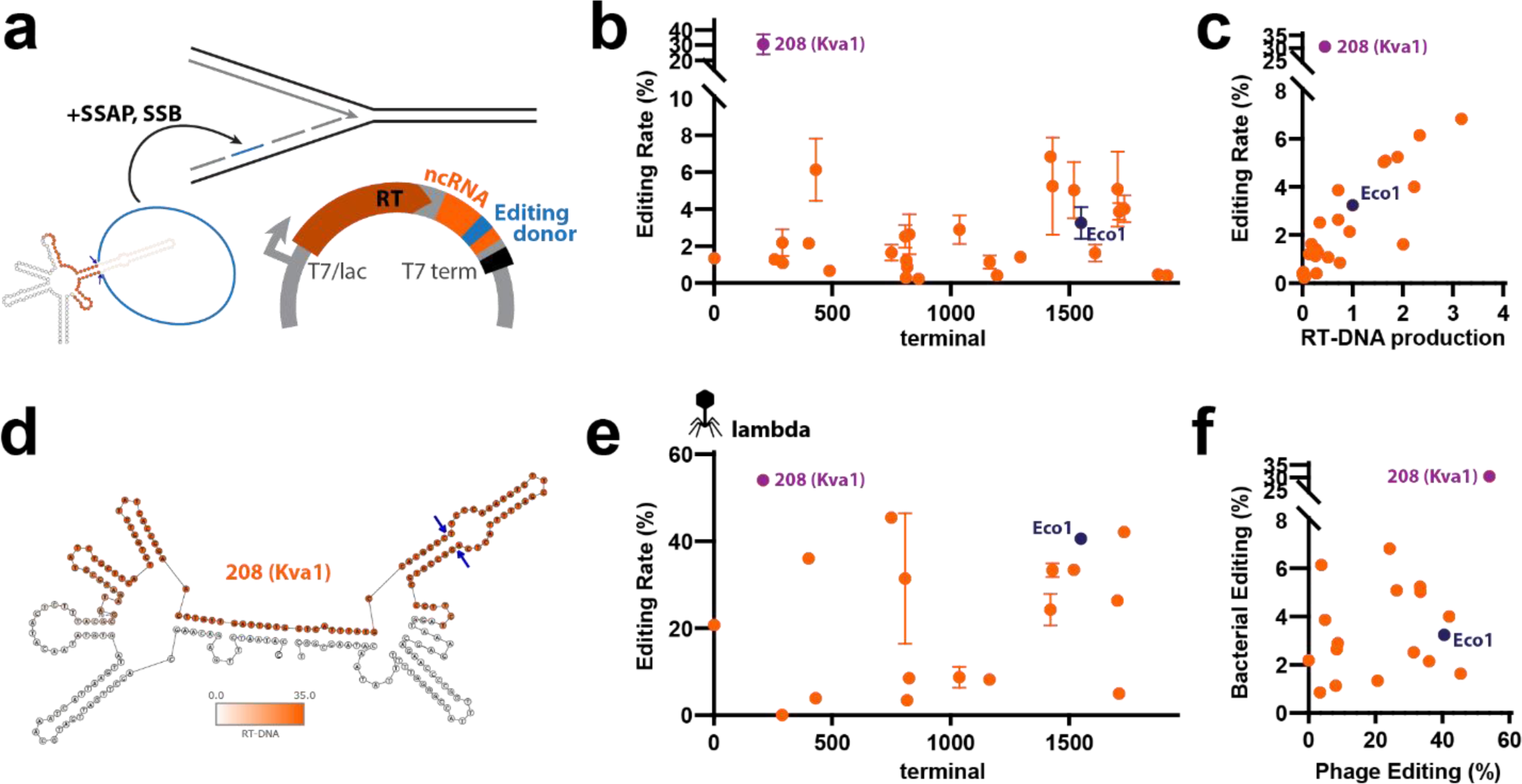
Bacterial editing across retrons. **a.** Schematic of modifications to the retron architecture for encoding a recombineering donor. **b.** Precise editing rate across retrons for bacterial genome recombineering. Points show mean ±SEM. **c.** RT-DNA production versus editing rate (r^2^=0.7477, *P*<0.0001). **d.** Predicted secondary structure of Kva1 ncRNA with sequencing data in orange. **e.** Precise editing rate across retrons for phage genome recombineering. Points show mean ±SEM. **f.** Bacterial versus phage editing rate (r^2^=0.2184, *P*=0.0505). Additional statistical details in **Supplemental Table 2**.

To survey the ability of retrons to support recombineering, a batch of 29 candidates were selected from our retron library, prioritizing higher RT-DNA producers across the phylogenic tree, and including retron-Eco1 as a reference. The ncRNAs of these retrons were modified to contain a 90 bp template donor to make a precise single nucleotide mutation in *rpoB* gene. The template location was placed within the RT-DNA region as determined by our sequencing data and in a centrally located hairpin stem. After analyzing recombineering efficiencies, we found that 8 retrons yield higher editing rates than retron-Eco1 (Fig 3b). We find that RT-DNA production from the wildtype ncRNA and editing using a modified DNA are strongly correlated, consistent with previous work showing that the abundance of recombineering donor is a limiting reagent in editing (Fig 3c). However, the most effective retron for bacterial editing deviates from this correlation. Strikingly, a retron from *Klebsiella variicola*, referred from now on as retron-Kva1 (Mestre-208, clade 1), shows a 10-fold increase editing efficiency compared to Eco1 despite producing only a moderate amount of RT-DNA. Thus, retron-Kva1 represents a particular case that will require further characterization to understand whether another specific feature, such as its predicted non-canonical ncRNA structure, could explain its high recombineering abilities (Fig 3d).

18 out of the 29 retron candidates were also assessed for their ability to support phage genome recombineering. These retrons were drawn from a subset of high RT-DNA producers across clades, modified to edit phage lambda *xis* gene (stop codon TGA>TAA) using a 70 bp RT-DNA donor. In this case, 4 retrons, including retron-Kva1, show higher recombineering rates than wild-type retron-Eco1 (Fig 3f). We found no strong correlation between RT-DNA production and lambda recombineering rates for this set of retrons (**Supplemental** Fig 3); the relative editing of bacterial and phage genomes by individual retrons is only weakly correlated (Fig 3f). This could indicate that the biological mechanism of bacterial and phage recombineering differ and that phage biology, RT-DNA structure or additional factors could impact editing rates. However, care should be taken with this interpretation, given that the set of retrons tested for phage editing was selected based on RT-DNA production and genome editing in bacteria.

### Human precise editing by a diverse set of retrons

To extend the retron census beyond function in bacteria, we tested a diverse set of retrons for their ability to precisely edit the genomes of cultured human cells. Analogous to bacterial and phage editing, a modified RT-DNA encodes an editing donor. In eukaryotic editing, however, this donor is used to precisely repair a genomic site after a targeted double strand break by a Cas9 nuclease. For clarity, we term this combination of a retron with Cas9 for precise editing as an editron.

We synthesized a set of 136 never-before-tested editrons for human editing and four editrons that have been previously tested as internal references. For human editing of the EMX1 locus, a human codon-optimized RT was driven by a constitutive CAG promoter, while a modified ncRNA fused to a CRISPR sgRNA was driven by a Pol III promoter (Fig 4a). We modified each ncRNA to encode an editing donor in the RT-DNA, which is designed to insert a retron-specific 10 base barcode. The donor also contains a recoding of the PAM nucleotides to protect the ncRNA/sgRNA/RT plasmid and prevent further cutting of the genome after precise editing. Based on previous findings that the length of the retron ncRNA’s a1/a2 region can affect editing rates^18^, we increased this region of the ncRNA up to 16 bases for any editrons where the endogenous a1/a2 was shorter than 16 bases. We used an H1 promoter to express the ncRNA/sgRNA for most editrons, but used a U6 for a small subset where the H1 promoter was incompatible with synthesis of the retron components. In a parallel experiment, we found that no difference between H1 and U6 promoters for retron-Eco1 (**Supplementary** Fig 4a).

**Figure 4.**
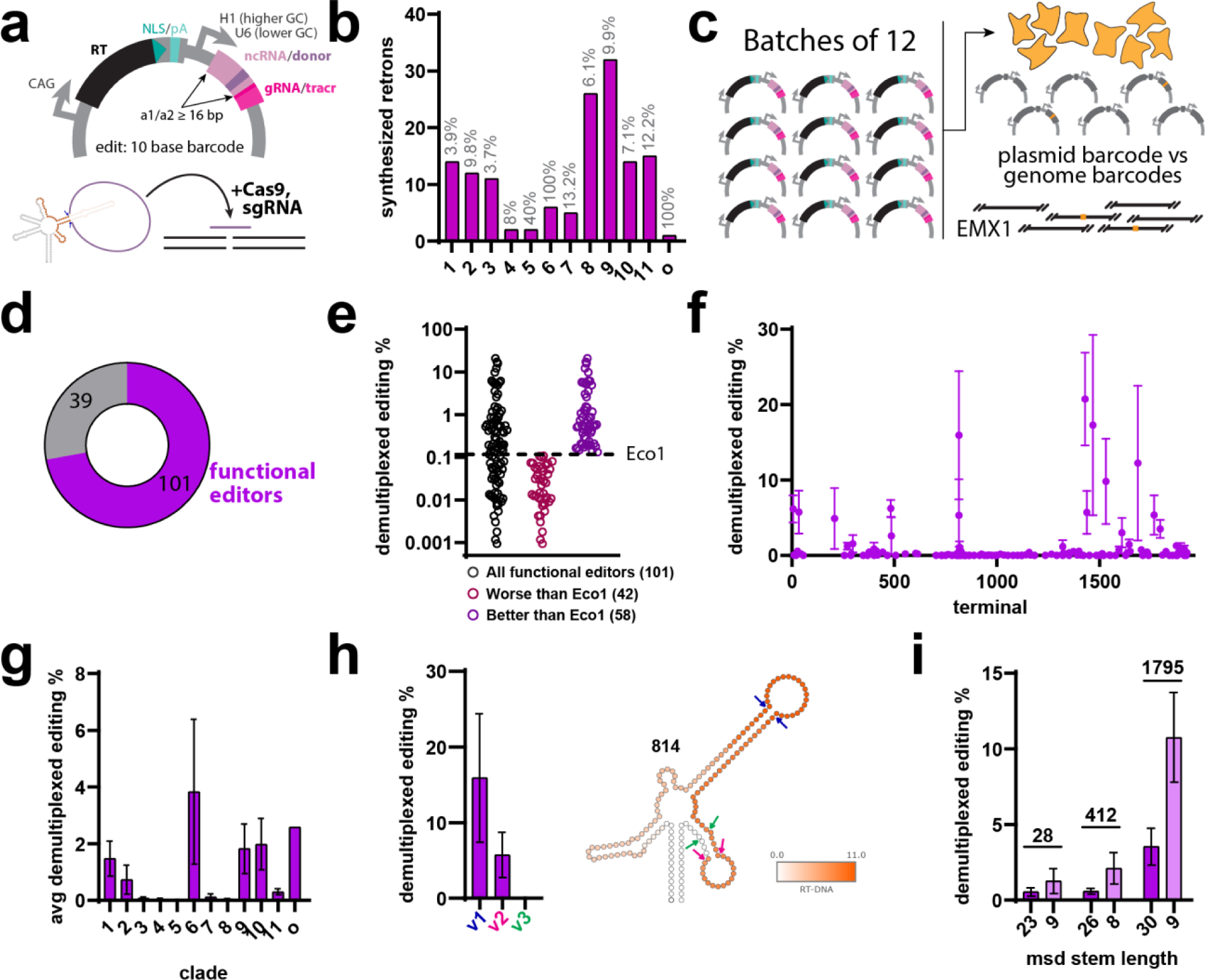
Human precise editing by a diverse set of retrons. **a.** Schematic of synthesized retron architecture for editing human genomes. **b.** Number of retrons synthesized by clade (percentage of clade synthesized indicated in gray above each bar). Bar o indicates an orphan, clade-less retron. **c.** Schematic of experimental workflow to test retrons in batches of 12. **d.** Proportion of retrons that enable precise editing at any efficiency. **e.** Demultiplexed editing percentage of functional editors, showing a range of efficiencies both above and below the prior reference retron-Eco1. **f.** Demultiplexed editing by retron. Points are mean ±SEM. **g.** Demultiplexed editing by clade (one-way ANOVA, effect of clade, *P*=0.2197). Bars are mean ±SEM. Bar o indicates an orphan, clade-less retron. **h.** Demultiplexed editing for retron Mestre-814 with the editing donor placed in different stem positions of the msd (one-way ANOVA, effect of donor location, *P=*0.3774). **i.** Demultiplexed editing for retrons tested with long and short versions of the msd stem (two-way ANOVA, effect of stem length, *P*=0.0017). Additional statistical details in **Supplemental Table 2**.

As in the bacterial work, we chose the human editrons to sample the phylogenic diversity of retrons, representing each RT clade. Of the 99 total retrons quantified for RT-DNA production and 140 total retrons quantified for human editing, 72 were tested in both conditions. We quantified editing in multiplexed pools of twelve editrons at a time to facilitate testing, using the unique 10 base pair insertion to demultiplex the edits and extrapolate the relative editing rate for each individual member of a pool. Specifically, we quantified the total precise editing rate for a given pool, then assigned a subset of that editing to each member of the pool based on the relative number of barcodes for that retron in the cell genomes and the relative number of plasmids for that retron in the particular pool (Fig 4c). Each editron appeared in at least two separate pools to mitigate effects of specific cross-reactivity within a pool.

The editron plasmids were transfected into HEK293T cells containing Cas9 pre-integrated into their genomes. Cells were collected three days after transfection. We generated two amplicons per sample for targeted Illumina sequencing: one which amplified the genome (to quantify editing) and one which amplified the plasmid (to account for differences in the relative abundance of retrons plasmids in the pool). We tested retrons in cells with either constitutively expressing or doxycycline inducible versions of the Cas9 and found similar results in both (**Supplemental** Fig 4b), and therefore present a merged dataset.

We found that the majority of retrons (101/140) were able to support precise editing to some degree (Fig 4d). Of the functional editrons, we find that 58 of the 101 support higher rates of precise editing than the previous standard retron, retron-Eco1 (Fig 4e). As in the case of RT-DNA production, we find that the functional retrons are not randomly distributed across the RT phylogeny, but rather occur in hotspots (Fig 4f). In fact, the best editrons were drawn from the same clades (1, 2, 6, 9, 10) as the highest RT-DNA producing retrons (Fig 4g). For one retron where we empirically determined the RT-DNA, we tested the placement of the donor at various positions either inside or outside the reverse transcribed region and found, unsurprisingly, that a donor in the core of the RT-DNA had a tendency to support higher levels of editing, consistent with the use of the RT-DNA as the editing donor and not the plasmid itself (Fig 4h). Additionally, we tested the effect of donor placement within the RT-DNA stem for three retrons and found that shortening the stem length that flanks the donor yielded higher rates of editing (Fig 4i).

## DISCUSSION

Retrons are notable for their innate immune function against phages and their applications in biotechnology, particularly for genome engineering. Despite their diversity, prior studies have only explored a limited range of retrons. Herethis study, we conduct a comprehensive survey of these bacterial systems, focusing on RT-DNA production, bacterial and phage recombineering, and precise human genome editing. This work complements and advances earlier work on retron systems, including an expansion of validated retron RTs, ncRNAs and RT-DNAs. Some of the validated ncRNAs are derived from retron subtypes with no previously predicted ncRNAs (e.g. clade 8, type XI).

The data regarding RT-DNA position within the ncRNA, which can currently only be determined empirically, are particularly valuable. As the number of empirically determined RT-DNA sequences increases, it may become possible to predict future sequences without experimentation. We observed significant variability in RT-DNA production among different retrons, with retrons from certain clades consistently producing more RT-DNA. Another interesting finding is the lack of RT-DNA production in clades 3 and 4. Perhaps the ncRNA from these retrons is noncanonical or positioned far from the retron operon. Alternatively, these retrons may require additional host factors for RT-DNA production.

Our study also sheds light on reverse transcription initiation and termination within retrons. We confirm the presence of a preceding TTA consensus in functional retrons and identify a prevalent AT-rich region at the site of termination area. It is also important to note that some of retrons were visualized as double bands on the gel, and different branched/debranched conformations in sequencing. It has been described that retrons undergo varying maturation processes, which may explain these results^2,25–27^.

This research not only provides biological insights but also expands the genome editing toolkit for both prokaryotic and eukaryotic cells. As in previous work^18^, we found a strong correlation between RT-DNA production and bacterial editing rates. When editing lambda phage genomes, this correlation was lost. Speculatively, this could be due to anti-retron systems in the lambda genome. Recent work has suggested the presence of an anti-retron protein in phage T5 that reduces the amount of RT-DNA inside the cell^28^. We also show that retron-Kva1 significantly outperforms the previous standard retron-Eco1 for both bacterial and phage recombineering, becoming an ideal candidate for further characterization and optimization.

We also characterized the ability of phylogenetically diverse retrons to edit mammalian cells, an application which has been limited by a small number of validated retrons. Our work identifies 58 retrons that edit mammalian cells at greater efficiencies than the previous biotechnological standard of retron-Eco1. Interestingly, the top retrons for bacterial RT-DNA production, bacterial recombineering, and human editors are all derived from an overlapping set of RT clades. However, bacterial RT-DNA production is not correlated with human editing at the level of individual retrons (**Supplemental** Fig 5). This difference could reflect RT expression differences in human versus bacterial cells, as well as reverse transcription efficiency. We also demonstrate a new method to test editing of retrons in multiplexed pools which harnesses the unique RT-DNA produced by each retron. This enables a more efficient and trackable screening platform for genome engineering.

Overall, our study is the most extensive characterization of retrons to date, validating a larger set of retrons and identifying retrons with better RT-DNA production, bacterial and phage recombineering, and human editing.

## METHODS

Biological replicates were taken from distinct samples, not the same sample measured repeatedly.

### Plasmids

For bacterial expression, retron ncRNA-RT pairs were synthesized into high-copy pET21 backbones with T7/lac inducible promoters (Twist). Codons were optimized only if necessary for synthesis. Constructs for recombineering were cloned from these plasmids to add an rpoB editing donor (S512P) or lambda xis editing donor (stop codon TGA to TAA). pORTMAGE-Ec1 was generated previously^29^.

All human vectors are derivatives of pSCL.273, itself a derivative of pCAGGS^30^. pCAGGS was modified by replacing the MCS and rb_glob_polyA sequence with an IDT gblock containing inverted BbsI restriction sites and a SpCas9 tracrRNA, using Gibson Assembly. The resulting plasmid, pSCL.273, contains an SV40 ori for plasmid maintenance in HEK293T cells. The strong CAG promoter is followed by the BbsI sites and SpCas9 tracrRNA. BbsI-mediated digestion of pSCL.273 yields a backbone for single or library cloning of plasmids with inserts that contain (retron RT - pol III promoter - modified retron ncRNA), by Gibson Assembly or Golden Gate cloning.

Retrons for human editing were synthesized into pSCL.273 (Twist). The RT is driven by the CAG promoter. The ncRNA/gRNA cassette was placed under a H1/U6 promoter, with the choice of promoter not impacting editing rates (Supplementary Fig 4a). The donor location loop was determined by viewing RNAFold structures. Typically, the donor was placed at the first unpaired bases of the predicted RT-DNA stem. The reverse transcribed repair template was slightly asymmetric (49 bp of genome site homology upstream of the Cas9 cut site; 71 bp of genome site homology downstream of the cut site) and was complementary to the target strand; in practice, this means that after reverse transcription, the repair template RT-DNA is complementary to the non-target strand, as recommended in previous studies^31^. The repair template for human editing carried three distinct mutations to the EMX1 locus: the first inserts a 10-bp sequence at the Cas9 cut site, with a unique sequence generated for each retron plasmid. The second changes a G>A after the 10-bp insert for the gRNA. The third recodes the Cas9 PAM (NGG → NTT). The gRNA is 20 bp and targets EMX1.

Retron accession information as well as all plasmids and ncRNA sequences are listed in Supplemental Table 1.

### Bacterial Strains and Growth Conditions

The E. coli strains used in this study were DH5α (New England Biolabs) for cloning, bSLS.114^18^ for RT-DNA production, bMS.346^15^ for bacterial and phage retron recombineering assays. bSLS.114 was constructed from BL21-AI cells using lambda-red replacement to remove the retron-Eco1 locus. bMS.346 was generated from *E. coli* MG1655 by inactivating the *exoI* and *recJ* genes with early stop codons. Bacterial cultures were grown in LB (supplemented with 0.1 mM MnCl2 and 5 mM MgCl2 (MMB) for phage assays), shaking at 37 °C with appropriate inducers and antibiotics. Inducers and antibiotics were used at the following working concentrations: 1mM m-toluic acid (Sigma-Aldrich), 1 mM IPTG (GoldBio), 2 mg/ml L-arabinose (GoldBio), 35 µg/ml kanamycin (GoldBio), and 100 µg/ml carbenicillin (GoldBio).

### RT-DNA expression and gel analysis

RT-DNA expression and analysis was performed as previously described^12^. Briefly, retron plasmids were transformed into bSLS.114 for expression. A starter culture from a single clone was grown overnight in 3 ml LB plus antibiotic. After 16h, the culture was diluted (1:100) into 25 ml LB. This culture was allowed to reach OD ∼0.5 (approximately 2 hours) and then induced with 1M IPTG and 200 ug/ml l-arabinose. After 5 hours, OD600 was measured and bacteria were harvested for RT-DNA analysis.

RT-DNA was recovered using a Qiagen Plasmid Plus Midi kit, eluted into a volume of 150 ul. Volume RT-DNA prep was adjusted based on bacterial OD, measured at the point of collection, to normalize the input prior to loading into Novex TBE-Urea gels (15% Invitrogen). The gels were run (45 minutes at 200 V) in pre-heated (>75°C) TBE running buffer. Gels were stained with SYBR Gold (ThermoFisher) and then imaged on a Gel Doc Imager (BioRad). To quantify the amount of RT-DNA production relative to retron-Eco1, a retron-Eco1 expressing strain was included every batch of culture grown, and the resulting prep of was always run on the same gel as the experimental retron for quantification. The density of the strongest band of each retron was quantified with ImageJ software.

### Multiplexed RT-DNA Sequencing

Following Qiagen midipreps, mixes of up to 11 different retrons (including Eco1) were subjected to a DBR1 or sham-DBR1 (water) treatment in the following reaction: 79 ul of RT-DNA prep split evenly depending on the number of retrons in the mix, 10 ul of DBR1 (50 ng/uL), 1 ul of RNaseH (NEB), 10 ul rCutSmart Buffer (NEB). The mixes of retrons were organized according to their relative production to retron-Eco1, with the highest producing retrons grouped together and lowest producing retrons grouped together. This minimized the skew in sequencing among the retrons in a pool.

These reactions were cleaned up with ssDNA/RNA clean & concentrator kit (Zymo Research) and RT-DNA was prepped for sequencing by taking the resulting material and extending the 3′ end in parallel with two types of nucleotides: dCTP or dATP, using terminal deoxynucleotidyl transferase (TdT) (NEB). This reaction was carried out in 1× TdT buffer, with 60 units of TdT and 125 M dATP for 60 s, or 125 M dCTP for 4 minutes at room temperature with the aim of adding ∼25 adenosines adenosines or ∼6 cytosines before inactivating the TdT at 70°C for 5 min. Next, a primer a polynucleotide, anchored primer was used to create a complementary strand to the TdT extended products using 15 units of Klenow Fragment (3′→5′ exo-) (NEB) in 1× NEB2, 1 mM dNTP and 50 nM of primer containing an Illumina adapter sequence, nine thymines (for the pA extended version) or six guanines (for the pC extended version), and a non-thymine (V) or non-guanine (H) anchor. This product was cleaned up using Qiagen PCR cleanup kit and eluted in 10 ul water. Finally, Illumina adapters were ligated on at the 3′ end of the complementary strand using 1× TA Ligase Master Mix (NEB). All products were indexed and sequenced on an Illumina MiSeq instrument. Previous data from Palka et al. 2022^12^ was included in the sequencing analysis for retron-Sen2 (Mestre-116), retron-Eco6 (Mestre-488), retron-Eco2 (Mestre-1036), and retron-Eco4 (Mestre-64).

### Bacterial Recombineering and Analysis

To edit bacterial genomes, the different retron cassettes encoded in a pET-21 (+) plasmid and the CspRecT and mutLE32K in the plasmid pORTMAGE-Ec1 were transformed into the bMS.346 strain by electroporation. Experiments were conducted in 500uL cultures in a deep 96-well plate. 3 individual colonies of every retron tested were grown in LB with kanamycin and carbenicillin for 16 h at 37°C. A 1:1000 dilution of these cultures was grown into LB with 1 mM IPTG, 0.2% arabinose and 1 mM m-toluic acid for 16 h with shaking at 37°C. A volume of 25 μl of culture was collected, mixed with 25 μl of water and incubated at 95°C for 10 min. A volume of 1 μl of this boiled culture was used as a template in 25-μl reactions with primers flanking the edit site, which additionally contained adapters for Illumina sequencing preparation. These amplicons were indexed and sequenced on an Illumina MiSeq instrument and processed with custom Python software to quantify the percentage of precisely edited genomes.

### Lambda Propagation and Plaque Assays

Lambda was propagated from an ATCC stock (#97538) into a 2mL culture of *E. coli* bMS.346 strain at 37°C at OD600 0.25 in MMB medium until culture collapse. The culture was then centrifuged and the supernatant was filtered to remove bacterial remnants. Lysate titer was determined using the full plate plaque assay method as described by Kropinski et al. 2009^32^.

### Lambda Recombineering and analysis

The diverse retron cassettes with modified ncRNAs to contain a lambda donor to edit *xis* gene, was co-expressed with CspRecT and mutL E32K from the plasmid pORTMAGE-Ec1. Experiments were conducted in 500uL cultures in a deep 96-well plate. 3 individual cultures were grown for every retron tested for 16h at 37°C. A 1:100 dilution of each culture was induced for 2 h at 37°C. The OD600 of each culture was measured to approximate cell density and cultures were diluted to OD600 0.25. A volume of pre-titered lambda was added to the culture to reach a multiplicity of infection (MOI) of 0.1. The infected culture was grown overnight for 16 h, before being centrifuged for 10 min at 4000 rpm to remove the cells. For amplicon-based sequencing, a volume of 25 μl of the medium containing the phage was collected, mixed with 25 μl of water and incubated at 95°C for 10 min. A volume of 1 μl of this boiled culture was used as a template in 25-μl reactions with primers flanking the edit site, which additionally contained adapters for Illumina sequencing preparation. Sequencing and editing rates were analyzed as described previously for bacterial recombineering.

### Human Editing Pool Design

Editrons were pooled in groups of 12. All editrons appear in at least two different pools (**Supplementary Table 1**). Individual plasmid cultures were started from glycerol stocks in 500 uL of LB Media +Carb for ∼6 hours of growth at 37 degrees. Then, OD600 was measured using SpectraMax platereader to determine how much of each culture to add to a mixed culture for equal distribution. Mixed cultures were grown overnight at 37 degrees and Midiprepped according to manufacturer’s protocol (Qiagen).

### Human Cell Culture

For pooled experiments, two Cas9-expressing HEK293T cell lines were used. The first expresses Cas9 from a piggybac integrated, TRE3G driven, doxycycline-inducible (1 µg/ml) cassette, which we have previously described^18^. The second expresses Cas9 constitutively from a CBh promoter in the AAVS1 Safe Harbor locus (GeneCopoeia #SL502). All HEK cells were cultured in DMEM +GlutaMax supplement (Thermofisher #10566016).

T25 cultures were transiently transfected with 12.76 ug of pooled plasmid per T25 using Lipofectamine 3000. Cultures were passaged for an additional 48 h. In the inducible Cas9 line, doxycycline was refreshed at passaging. Three days after transfection, cells were collected for sequencing analysis. To prepare samples for sequencing, cell pellets were collected, and gDNA was extracted using a QIAamp DNA mini kit according to the manufacturer’s instructions. DNA was eluted in 200 µl of ultra-pure, nuclease-free water.

### Human Sample Preparation and Analysis

For pooled samples, 0.5 µl of the gDNA was used as template in 25-µl PCR reactions with primer pairs to amplify the locus of interest and a PCR reaction with primer pairs to amplify the plasmid region, both of which also contained adapters for Illumina sequencing preparation. Lastly, the amplicons were indexed and sequenced on an Illumina MiSeq instrument and processed with custom Python software to quantify the percentage of on-target precise genomic edits normalized to the representation of each plasmid in the pool.

## Data Availability

All data supporting the findings of this study are available within the article and its supplementary information, or will be made available from the authors upon request. Sequencing data associated with this study is be available on NCBI SRA (PRJNA1047666).

## Code Availability

Custom code to process or analyze data from this study will be made available on GitHub prior to peer-reviewed publication.

## Acknowledgements

Work was supported by funding from the National Science Foundation (MCB 2137692), the National Institute of Biomedical Imaging and Bioengineering (R21EB031393), the National Institute of General Medical Sciences (1DP2GM140917), and research support from Retronix Bio. S.L.S. is a Chan Zuckerberg Biohub – San Francisco Investigator and acknowledges additional funding support from the L.K. Whittier Foundation and the Pew Biomedical Scholars Program. A.G.-D. was supported by the California Institute of Regenerative Medicine (CIRM) scholar program. S.C.L. was supported by a Berkeley Fellowship for Graduate Study. R.F.F. was supported by a UCSF Discovery Fellowship. We thank Karen Zhang and David Wen for comments on the manuscript.

## Author Contributions

S.C.L, A.G.-D, and S.L.S. conceived the study. M.R.-M. and A.G.-D. performed the experiments in bacteria and phage. A.G.K performed experiments in human cells. S.L.S., A.G.K., M.R.-M., A.G.-D, S.C.L. and R.F.F. analyzed the data. A.G.K., M.R.-M. A.G.-D, and S.L.S. wrote the manuscript with inputs from all authors.

## Supplementary Figures

**Supplementary Figure 1.**
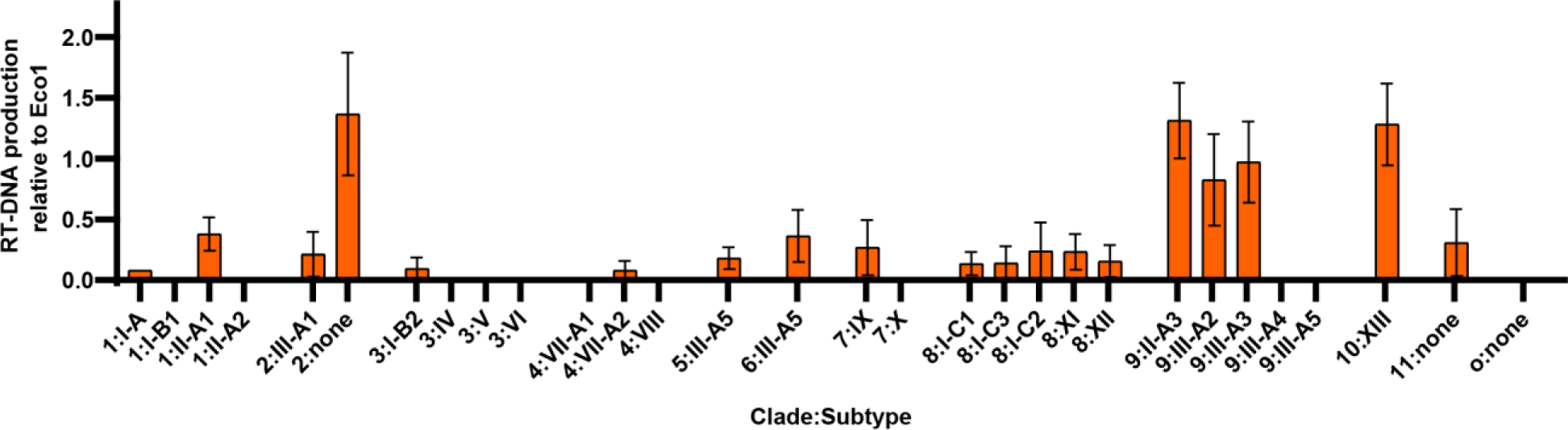
Related to Figure 1. RT-DNA production relative to Eco1 by clade, separated by subtype. Bars show mean ±SEM.

**Supplementary Figure 2.**
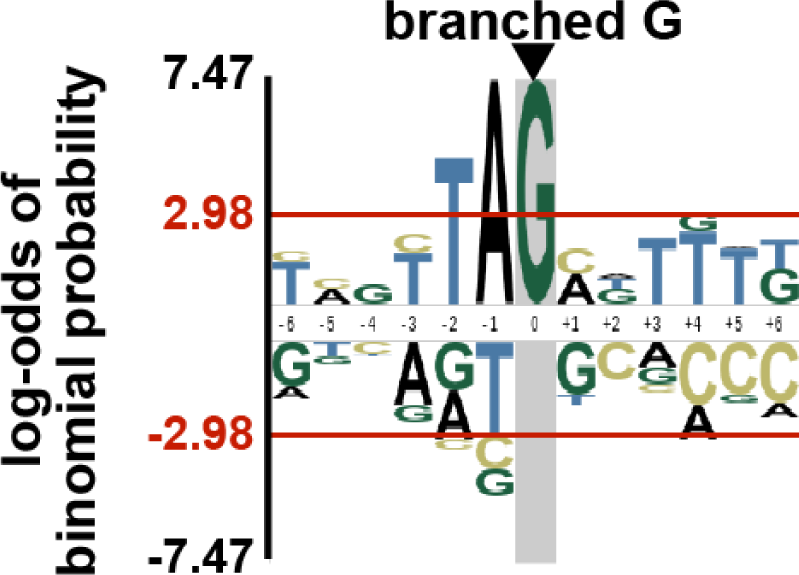
Related to Figure 2. Probability logo of ncRNA nucleotides adjacent to the branching guanosine for retrons that produce RT-DNA.

**Supplementary Figure 3.**
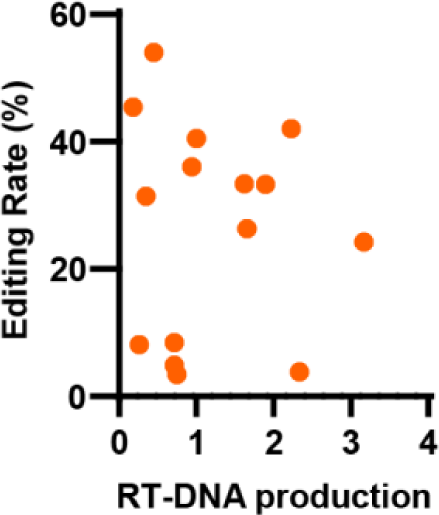
Related to Figure 3. RT-DNA production versus with phage lambda recombineering rates (Pearson Correlation (r^2^=0.001926, *P*=0.8766).

**Supplementary Figure 4.**
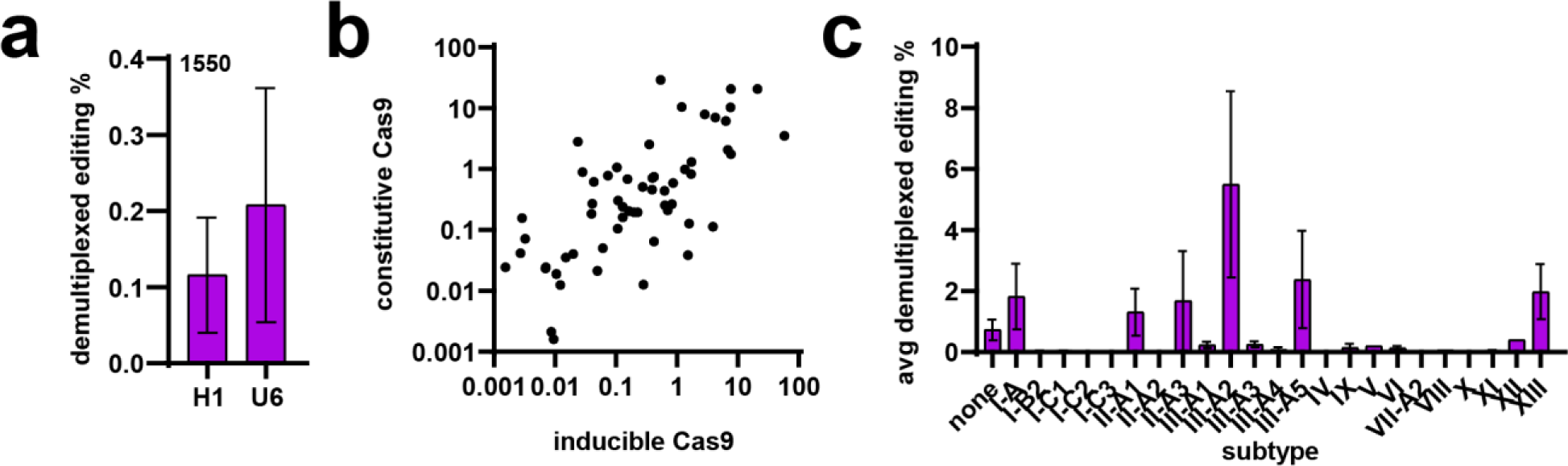
Related to Figure 4. **a.** Demultiplexed editing percentage by retron-Eco1 (Mestre-1550) comparing the use of an H1 and U6 promoter to drive ncRNA/sgRNA (unpaired T-test, *P*=0.6256). Bars are mean ±SEM. **b.** Comparison of demultiplexed editing rates of editrons in the constitutive and inducible Cas9 cell lines (Pearson Correlation, r^2^=0.4196, *P*<0.0001). **c.** Average demultiplexed editing rates across retron subtypes. Bars are mean ±SEM.

**Supplementary Figure 5.**
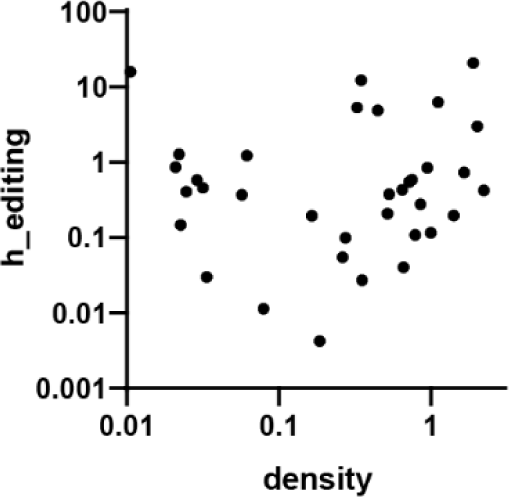
Related to Figure 4. Bacterial RT-DNA production compared with human demultiplexed editing.

