## Supplementary Figure 6 for "An experimental census of retrons for DNA production and genome editing"

Supplementary Figure 6: Figure 1d, uncropped gels

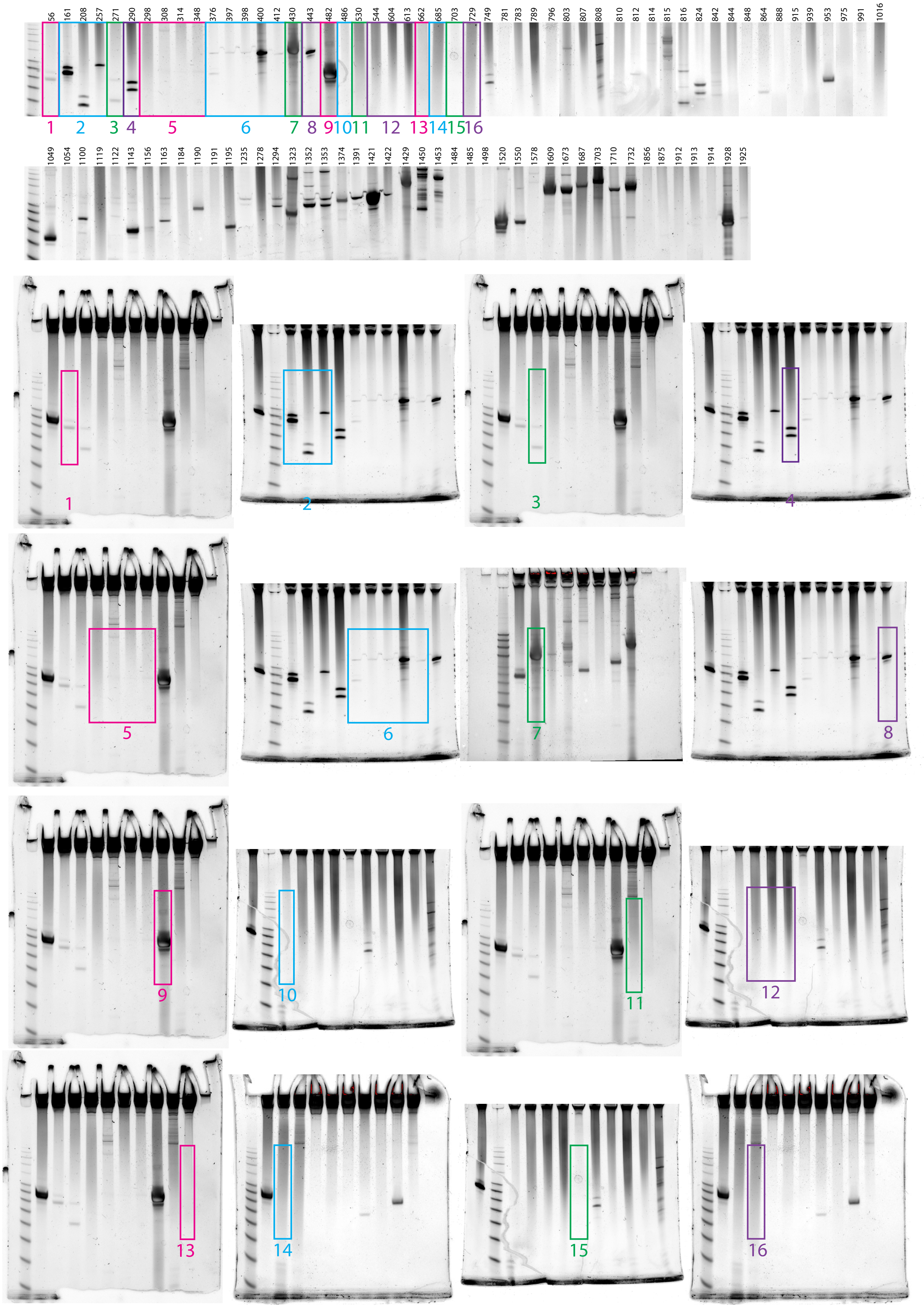

Supplementary Figure 6: Figure 1d, uncropped gels (continued)

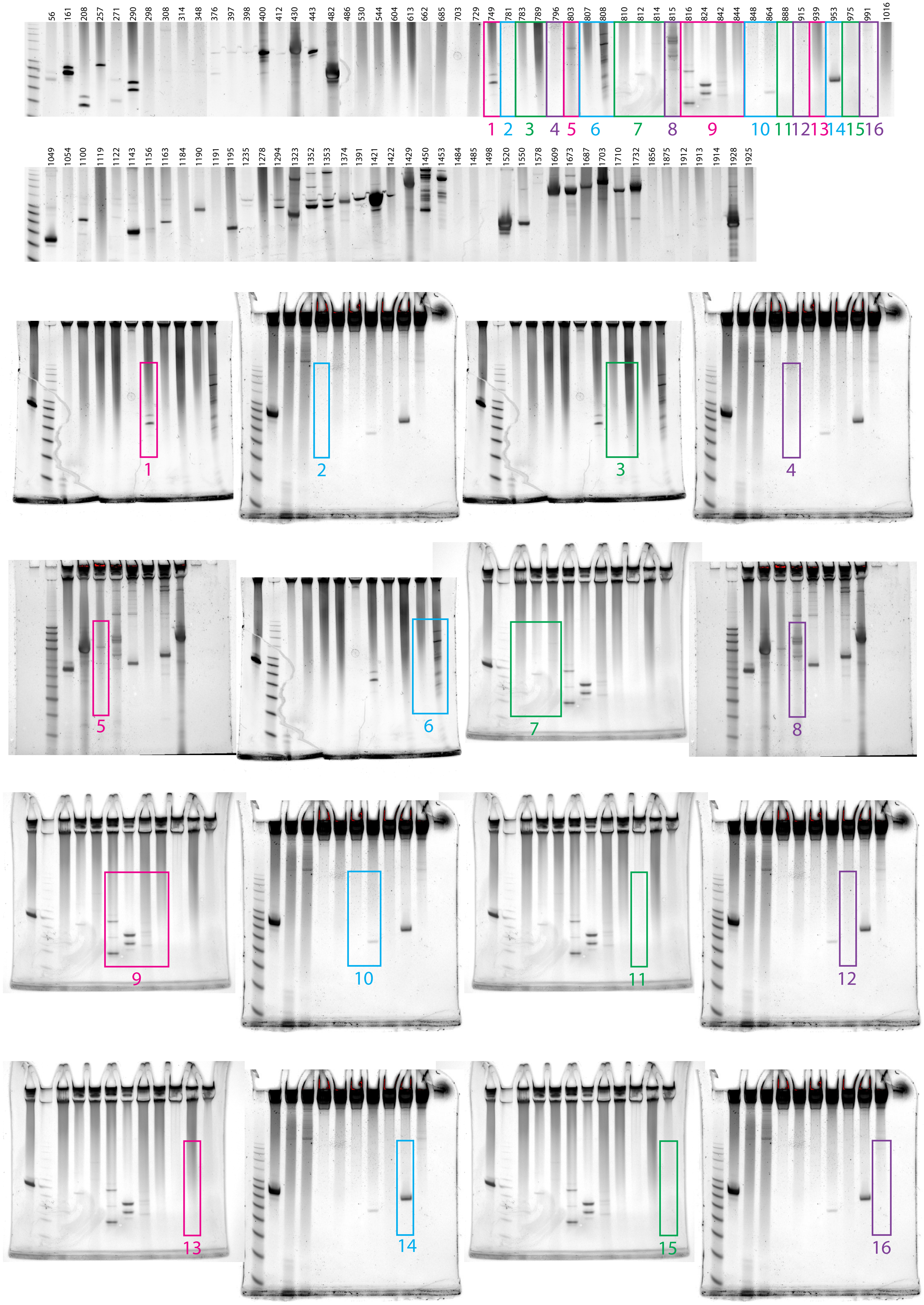

Supplementary Figure 6: Figure 1d, uncropped gels (continued)

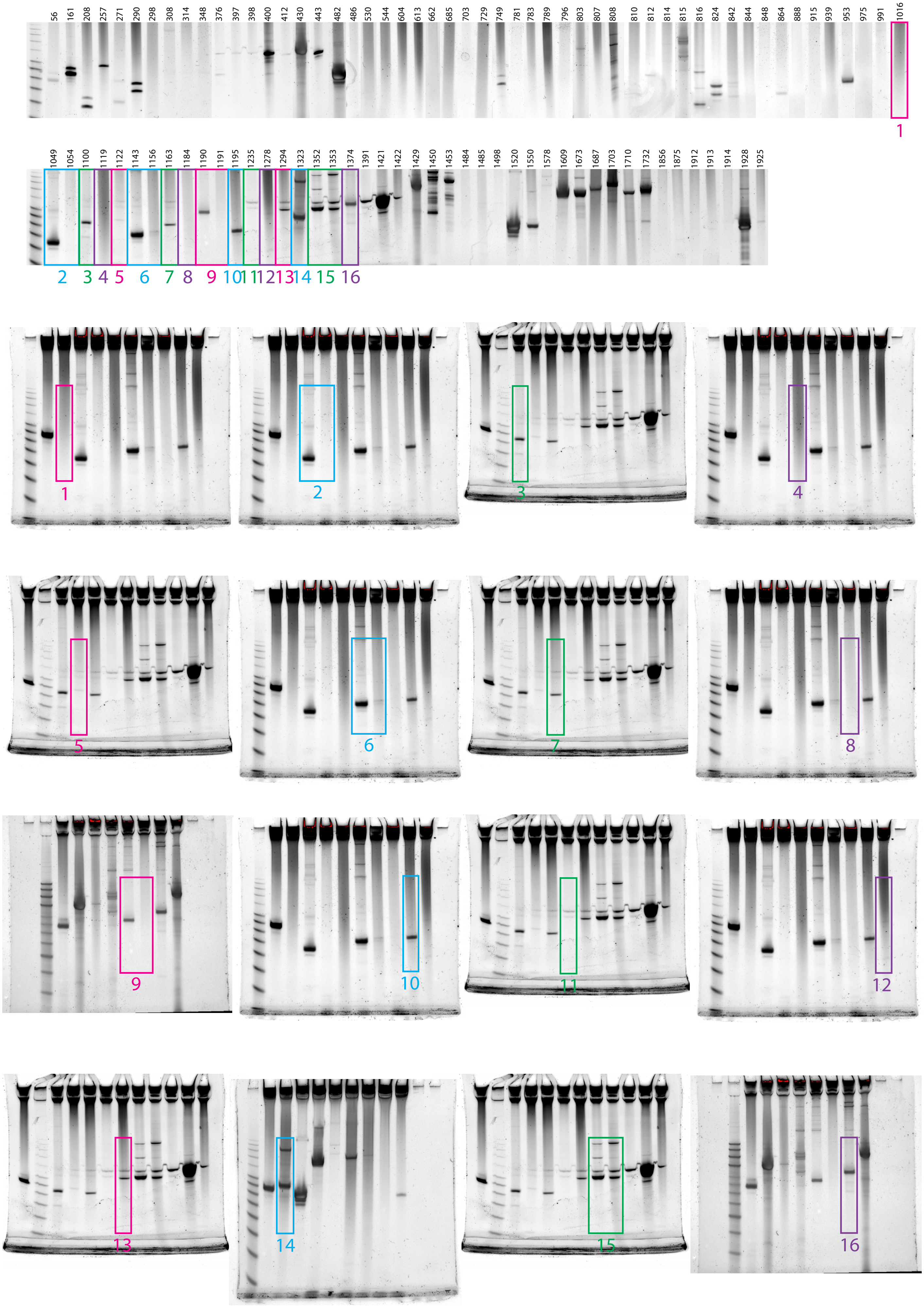

Supplementary Figure 6: Figure 1d, uncropped gels (continued)

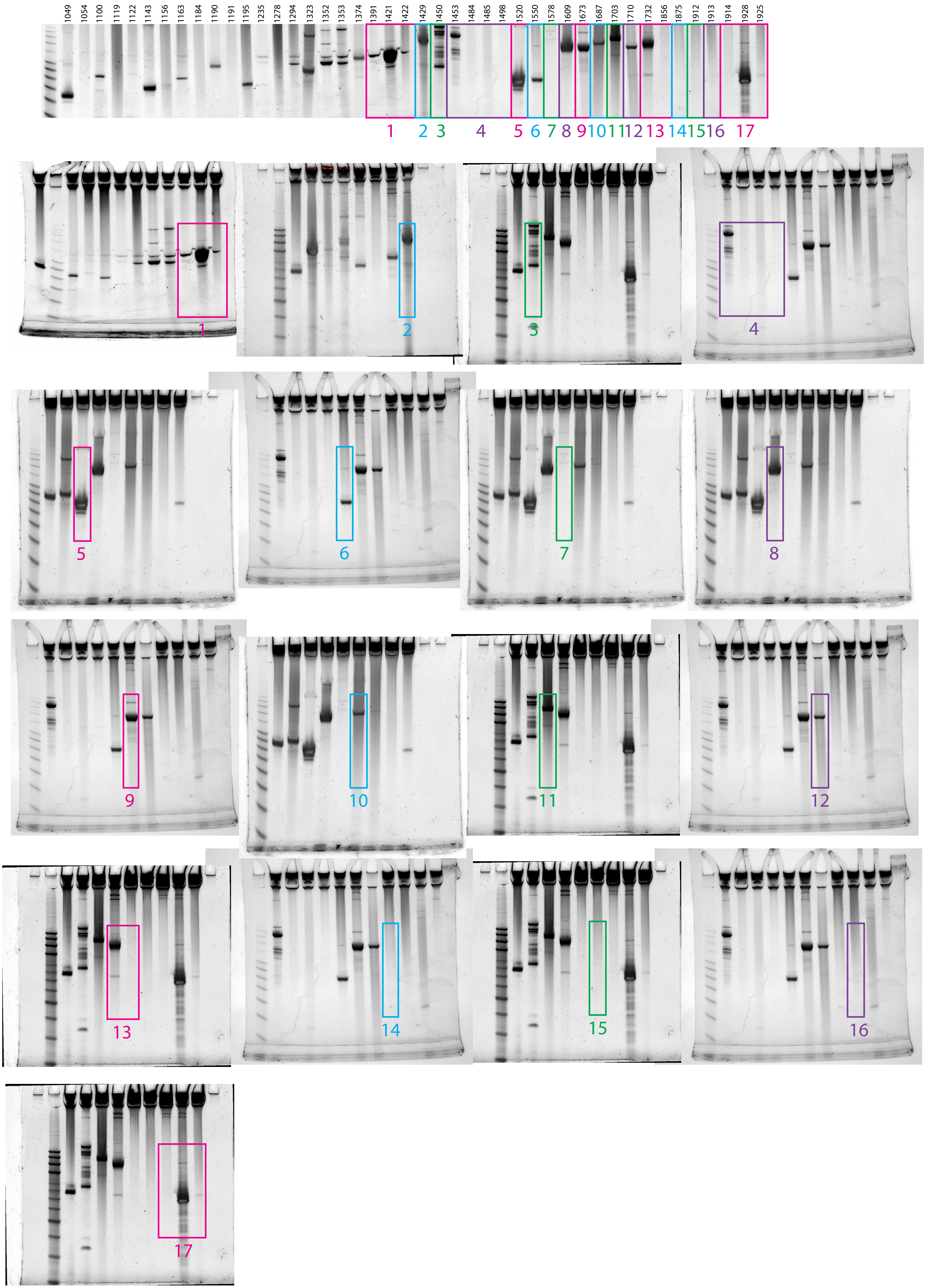
