## Supplemental Data 1 for "An experimental census of retrons for DNA production and genome editing"

### Retron Terminal 6: Proteobacteria

**Bacterial Vector for RTDNA: None**

**Bacterial Vector for Editing: None**

**Phage Vector for Editing: None**

**Human Vector for Editing: pSLS.948**

**RTDNA Production (relative to Eco1): undetermined**

**Bacterial Editing: undetermined**

**Phage Editing: undetermined**

**Human Editing (demultiplexed): 6.161448 percent precise (53.07429516x Eco1)**

**ncRNA:**

**TCAACTCTTTAGCGTGGACGTATTACGTCTAGTCGGGTGATTAGCCAGACTCTAACTTATTGAACG**
**CATTTGGGGTTGCGAAAGTGTCGCACCCCTCACTCGTTCCTCCCTCGGGTTGCGACACTTTCGCTT**
**CCTCCAGTAAAGAGTTGA**

**no RTDNA sequencing data**

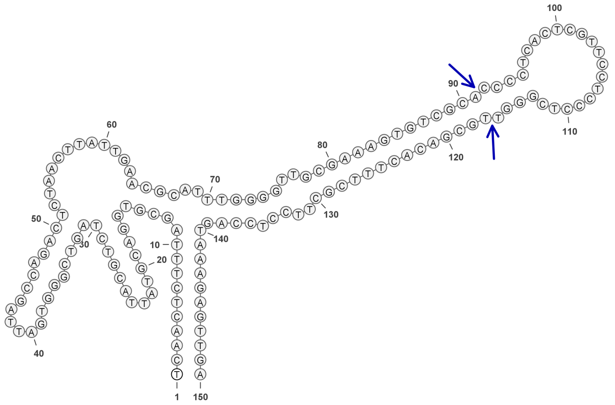

### Retron Terminal 10: Proteobacteria

**Bacterial Vector for RTDNA: None**

**Bacterial Vector for Editing: None**

**Phage Vector for Editing: None**

**Human Vector for Editing: pSLS.928**

**RTDNA Production (relative to Eco1): undetermined**

**Bacterial Editing: undetermined**

**Phage Editing: undetermined**

**Human Editing (demultiplexed): 0.074212 percent precise (0.639257134x Eco1)**

**ncRNA:**

**GTGCGATAAGCTTAATAAACTCTTTAGCGTAGGACATATTATGTCTTGTCGGGTGATTAGCCAGAC**
**ACTAATTTATTGCTCACATTTAGGGTTGCGAAAGTATCGCAACCTGATCTCGCGAGCTCCTTCAGG**
**TTGCGATACTTTCGCTTCTTCTAGTAAAGAGTAAAATTCGGGAGTACATCCGTACTCCCGTTTTCTT**
**ACAGGTGAAATTATGCAGATGATTAAACAATTAGCTGATTCGTTACAACAAGATGAATCAGATATA**
**AAAAAATTTTTGCAAAATGGACCCCGTAAATATAAGGTTTATCGCAT**

**no RTDNA sequencing data**

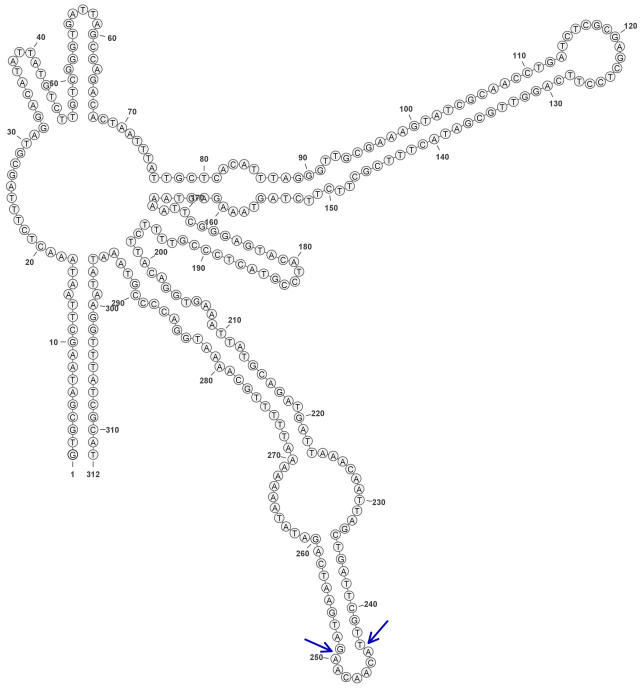


**10**

### Retron Terminal 21: Proteobacteria

**Bacterial Vector for RTDNA: None**

**Bacterial Vector for Editing: None**

**Phage Vector for Editing: None**

**Human Vector for Editing: pSLS.949 (U6)**

**RTDNA Production (relative to Eco1): undetermined**

**Bacterial Editing: undetermined**

**Phage Editing: undetermined**

**Human Editing (demultiplexed): 0 percent precise (0x Eco1)**

**ncRNA:**

**AGCGTCAGTTGCAAGATTTTATTATATTTCCTAAACATAAATGCGCAATGGCGTATCGCAAAGGAT**
**TGAGTATAAAAAATAATGCATTGTCTCATGTGAATAATTCGTATCTTCTTAAGTTAGACTTAGAGAA**
**TTTTTTTAATAGTATTACACCGGATCTTTTTTGGAGTGTATGGTCAAGAACTTGGTCTTTACCATCAA**
**AAGAAGACCAATTTTTTTTAGAGAAATTGATTTTTTGGCGTT**

**no RTDNA sequencing data**

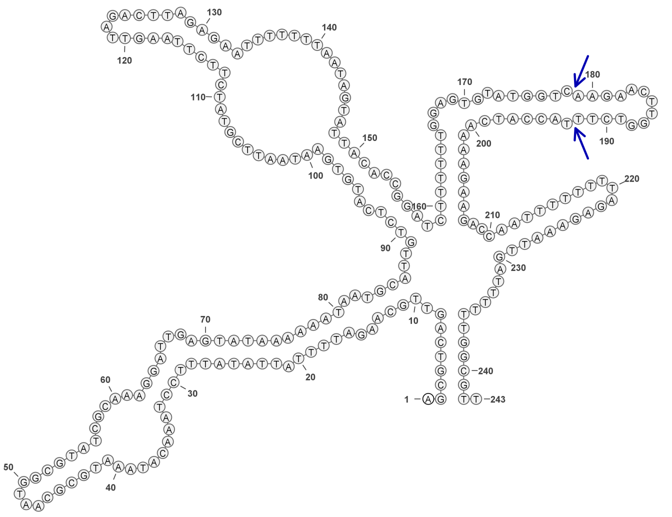

### Retron Terminal 28: Proteobacteria

**Bacterial Vector for RTDNA: None**

**Bacterial Vector for Editing: None**

**Phage Vector for Editing: None**

**Human Vector for Editing: pSLS.843**

**RTDNA Production (relative to Eco1): undetermined**

**Bacterial Editing: undetermined**

**Phage Editing: undetermined**

**Human Editing (demultiplexed): 0.538586 percent precise (4.639343274x Eco1)**

**ncRNA:**

**TAGTTTGTCTTTTAGCGAATGAGGCATTTATGCCTAGTCGGGTGTTTAGCCAGACTCTAACTTATTG**
**AACGAGATTGTATGCGATATCTGCGCATACAAAGCTCGTTCCTCACTTTGTATGCGCAGATATCGC**
**ATTAACTAGTAAAAGACAACCTA**

**no RTDNA sequencing data**

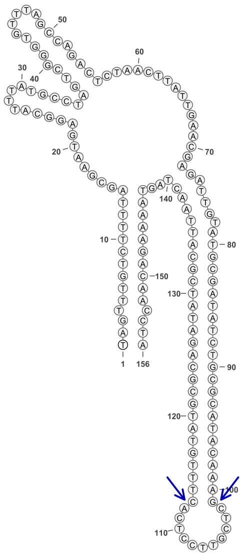

28

### Retron Terminal 34: Proteobacteria

**Bacterial Vector for RTDNA: None**

**Bacterial Vector for Editing: None**

**Phage Vector for Editing: None**

**Human Vector for Editing: pSLS.950**

**RTDNA Production (relative to Eco1): undetermined**

**Bacterial Editing: undetermined**

**Phage Editing: undetermined**

**Human Editing (demultiplexed): 5.750998 percent precise (49.5387067x Eco1)**

**ncRNA:**

**CTCTCTAGCTTAGGACTCTATGTCCAGTCGGGTGTTTAGCCAGACTCTAATTTATTGAACGGTTTGG**
**GGTTGCGAAAGACTCGCATCCTAAACTCGTTCCTCATTTAGGATGCGAGTCTTTCGCTTCCTCTAG**
**TAGAGAG**

**no RTDNA sequencing data**

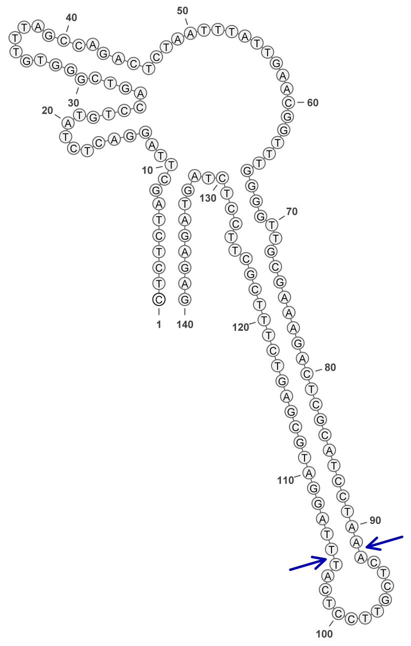


34

### Retron Terminal 44: Proteobacteria

**Bacterial Vector for RTDNA: None**

**Bacterial Vector for Editing: None**

**Phage Vector for Editing: None**

**Human Vector for Editing: pSLS.943**

**RTDNA Production (relative to Eco1): undetermined**

**Bacterial Editing: undetermined**

**Phage Editing: undetermined**

**Human Editing (demultiplexed): 0.208382 percent precise (1.794988414x Eco1)**

**ncRNA:**

**TATGACTCATTAAAATCAAAACATATTTTTAGTTTGATGGGAGATGGTGGTGATTGGACAAAAGAA**
**AAAATCAAAGAGCATCAAACATATATACTGGCTGTATTAACAAAACACTATGATGACGAAAAATAA**
**GTGAATAAGAATTAGTTTGAAGGTAATATGCACTTGATATATAATTAAAAAGCTCTTTAGCTTAAGA**
**CGAATTTCGTCTAGTCGGGTGTTTAGCCAGACTCTAATTTATTGAACGACATTATGGTTGCGATATG**
**TTCGCAACCAGATCTCGTACCTCAATCAGGTTGCGAACATATCGCTTCCTCTAGTAGAGAGCAAAG**
**CCCAAAAGCGATGCTTTTGGGCTTTTTTAATGGGTATA**

**no RTDNA sequencing data**

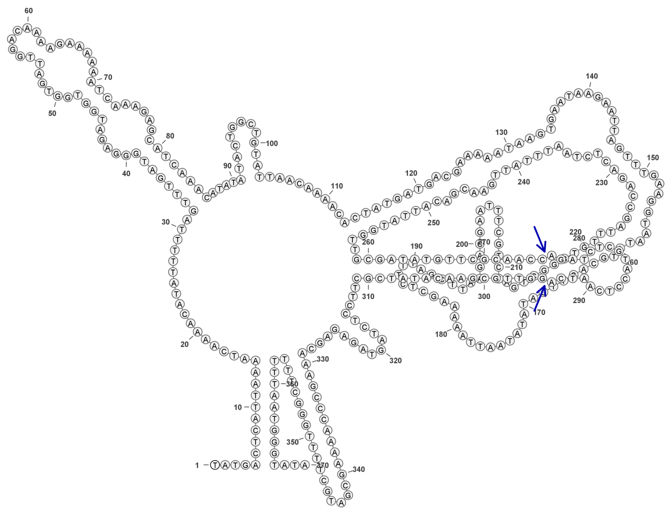

**44**

### Retron Terminal 56: Proteobacteria

**Bacterial Vector for RTDNA: pSLS.654**

**Bacterial Vector for Editing: None**

**Phage Vector for Editing: None**

**Human Vector for Editing: pSLS.899**

**RTDNA Production (relative to Eco1): 0.0794138 by PAGE**

**Bacterial Editing: undetermined**

**Phage Editing: undetermined**

**Human Editing (demultiplexed): 0.011369 percent precise (0.097931795x Eco1)**

**ncRNA:**

**GCTCTTTAGTTATGGGTTCTATGGGCCTAGTCGGGTGTTAAGCCAGACTCAAATTTATTGAGAGCA
TTGAGCGTTGCGAAAGGTAGATACCTTTCGCTCAATGTTCGCACATATTCGCAAAAGGTATCTACC
TTTCGCTTCGTATGAGTAGAGAGT**

**RTDNA:**

**CGAAGCGAAAGGTAGATACCTTTTGCGAATATGTGCGAACATTGAGCGAAAGGTATCTACCTTTC
GCAACGCTCAATGCTCTC**

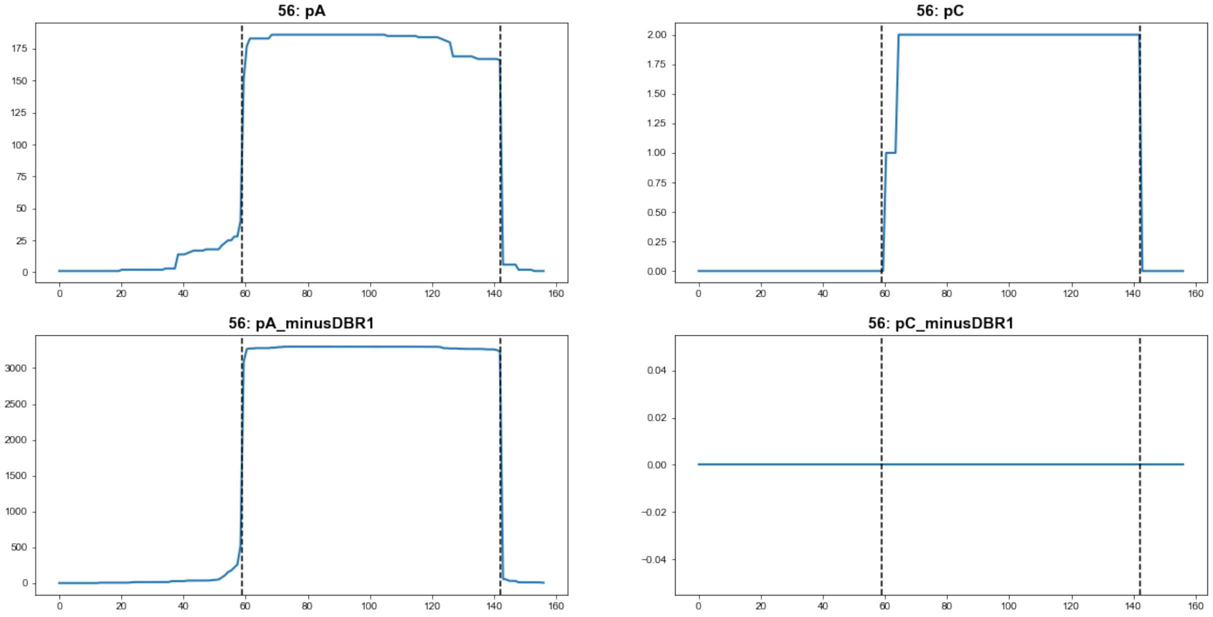


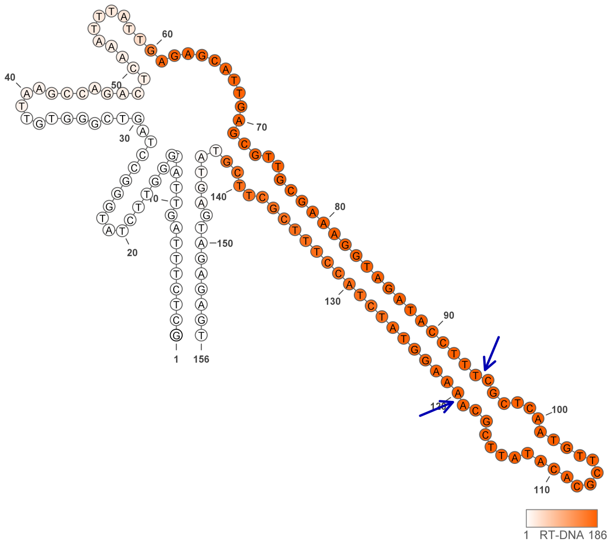

### Retron Terminal 64: Proteobacteria

**Bacterial Vector for RTDNA: pSLS.490**

**Bacterial Vector for Editing: None**

**Phage Vector for Editing: None**

**Human Vector for Editing: None**

**RTDNA Production (relative to Eco1): undetermined**

**Bacterial Editing: undetermined**

**Phage Editing: undetermined**

**Human Editing (demultiplexed): undetermined**

**ncRNA:**

**GCTCTTTAGCGTTTTATGGATTTACCACCTGATTGGTCAAATCTAGTTGGGCGTTGCGCCAAACTCT AATTTATTGATTACATTTACAGTTGCGGAACAAACTTTTTGAGCCGCAATCCAGTAGGTTGCGGAT CAAAAAGTTTGTTCCGCGGCTTCAAGTAAAGAGC**

**RTDNA:**

**AGCCGCGGAACAAACTTTTTGATCCGCAACCTACTGGATTGCGGCTCAAAAAGTTTGTTCCGCAA CTGTAAATGTAATC**

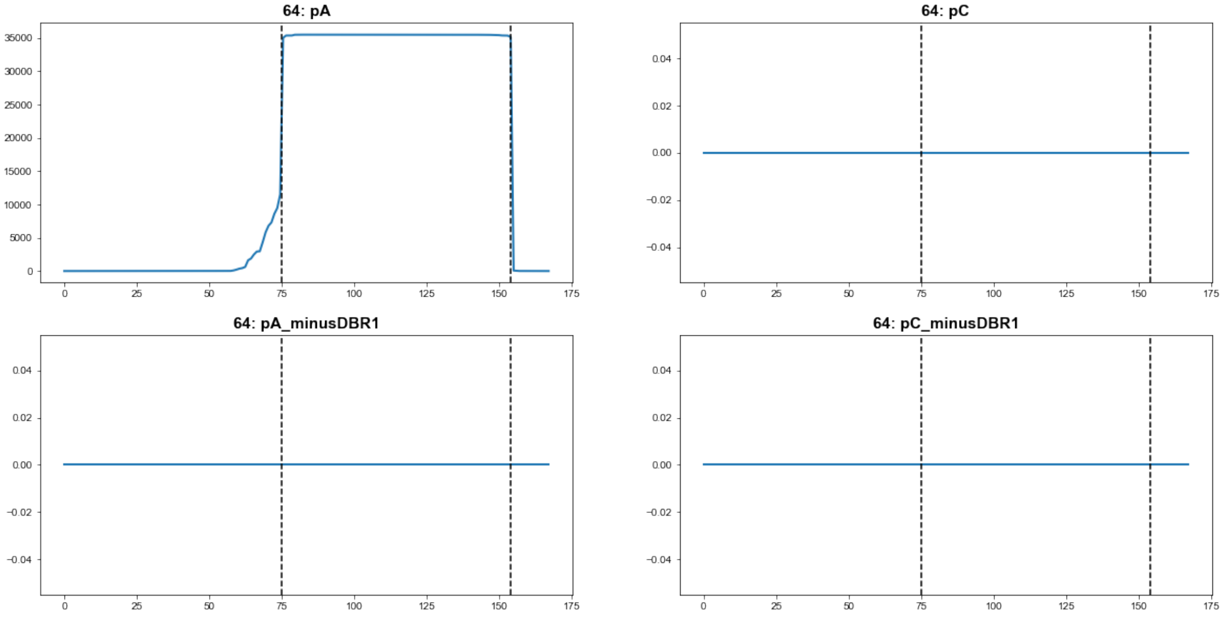


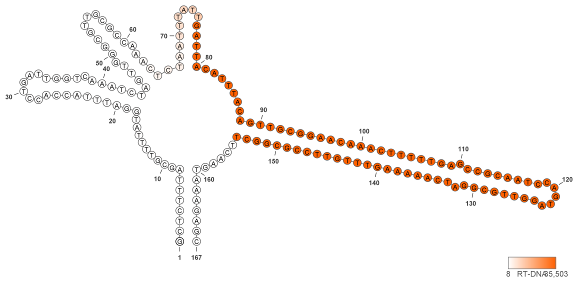

**64**

### Retron Terminal 116: Proteobacteria

**Bacterial Vector for RTDNA: pSLS.488**

**Bacterial Vector for Editing: None**

**Phage Vector for Editing: None**

**Human Vector for Editing: None**

**RTDNA Production (relative to Eco1): undetermined**

**Bacterial Editing: undetermined**

**Phage Editing: undetermined**

**Human Editing (demultiplexed): undetermined**

**ncRNA:**

**ACTCTTTAGCGTTAGGCTTTGATTTATAGCCTTGTCGAGCGTTTCGCCAGACACTAACTTATTGAGT
ACTTTTAGGGTTGCGCTAGAAAGTTTTCTACCGATCCTAGAAGTCTCTAGGATCGGTAGAAAACTT
TCTAGCGCCTCCTCTAGTAAAGAGT**

**RTDNA:**

**GGAGGCGCTAGAAAGTTTTCTACCGATCCTAGAGACTTCTAGGATCGGTAGAAAACTTTCTAGCG
CAACCCTAAAAGTACTC**

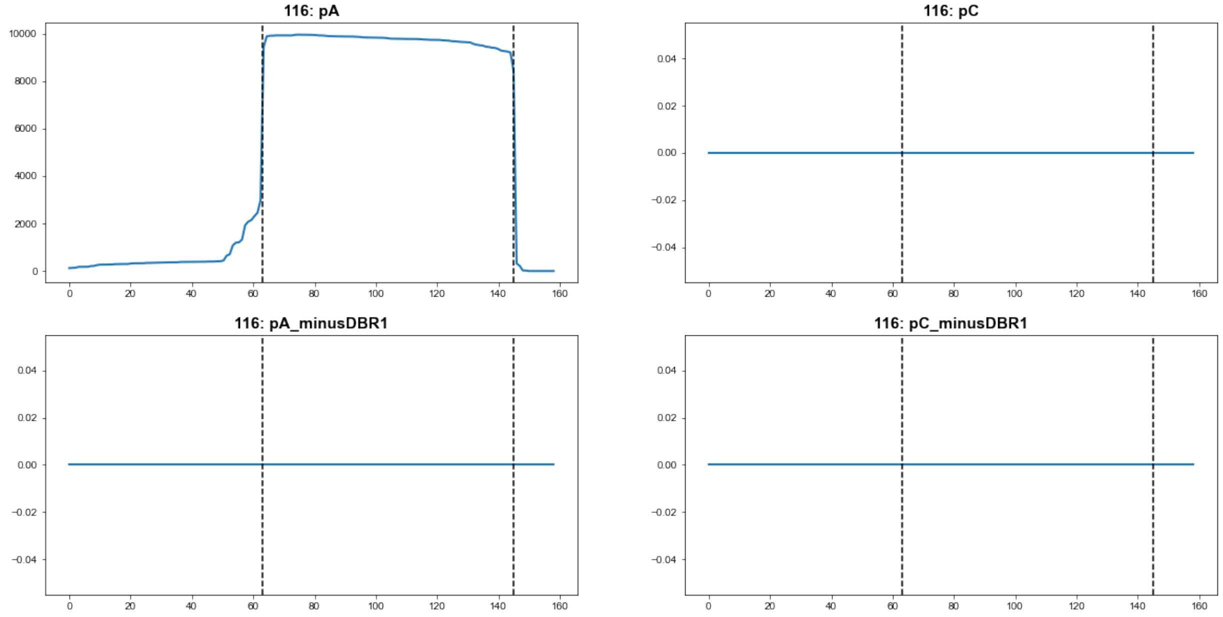


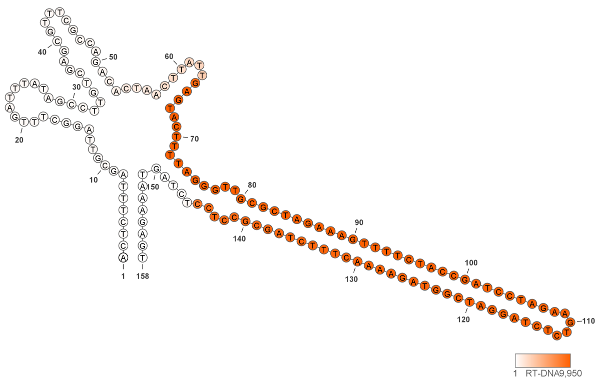


**116**

### Retron Terminal 161: Proteobacteria

**Bacterial Vector for RTDNA: pSLS.598**

**Bacterial Vector for Editing: None**

**Phage Vector for Editing: None**

**Human Vector for Editing: pSLS.833**

**RTDNA Production (relative to Eco1): 0.930615785 by PAGE**

**Bacterial Editing: undetermined**

**Phage Editing: undetermined**

**Human Editing (demultiplexed): undetermined**

**ncRNA:**

**GGGCTCAATTACGCGCGGTAGCAGACATTTTTTTGCTATCAATTGTCGGGCGTTTCGCCAGACTGA GCTGTAGTGGACTACAACAACGGCACGAAAGCGGCATGCGCCGAATCCGTGTCCAGTGGGTCCC GATTCGGCGCATGCCGCTTTCGTGCCTATCTTCAGGTAATTGACCT**

**RTDNA:**

**GATAGGCACGAAAGCGGCATGCGCCGAATCGGGACCCACTGGACACGGATTCGGCGCATGCCG CTTTCGTGCCGTTGTTGTAGTCC**

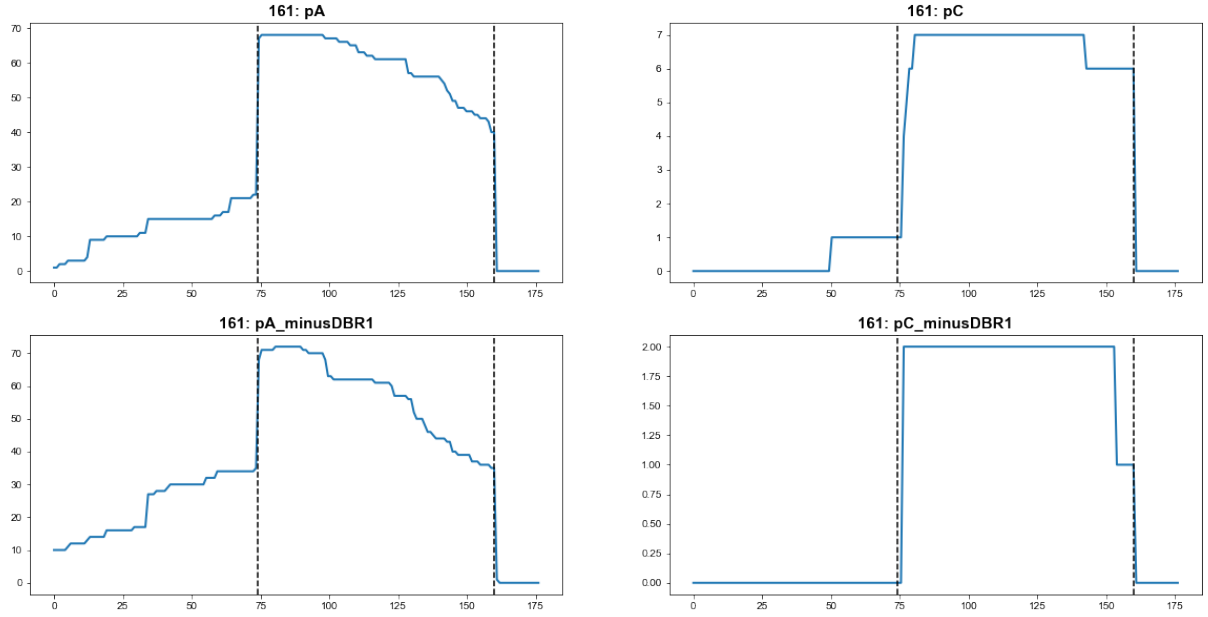


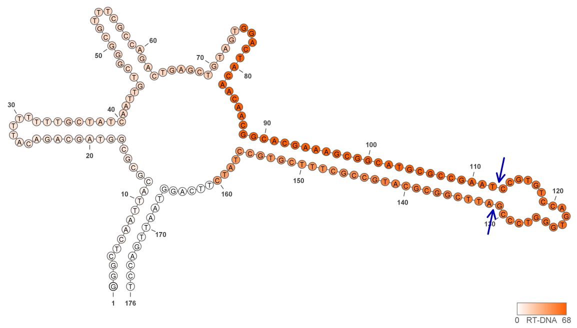


**161**

### Retron Terminal 208: Proteobacteria

**Bacterial Vector for RTDNA: pSLS.588**

**Bacterial Vector for Editing: pAGD289**

**Phage Vector for Editing: pAGD380**

**Human Vector for Editing: pSLS.823**

**RTDNA Production (relative to Eco1): 0.447202949 by PAGE**

**Bacterial Editing: 30.5988386 percent precise (9.415646145x Eco1)**

**Phage Editing: 54.05041585 percent precise (1.333135789x Eco1)**

**Human Editing (demultiplexed): 4.910143 percent precise (42.29563877x Eco1)**

**ncRNA:**

**CCATAATCTTGAGACAAGCAGCTTGATTGATGCAAATCATTAAGTTAGTGTATAACATACTCTTTAC**
**GCGATAGAGGCGTATCTGCTTCATCTGGTGTATCACCGGAACTTGTTGATTGTGGTCATTTATGCC**
**ACGGCCTTCCCAAAAATCTTCGATTTTTACTCAGGCCGTGCCCTTCTAGGTAAAGAGCAGAACCCC**
**GGTTTACCGGGGTTTTTTATAACATAAGCGGCT**

**RTDNA:**

**AGAAGGGCACGGCCTGAGTAAAAATCGAAGATTTTTGGGAAGGCCGTGGCATAAATGACCACAA**
**TCAACAAGTTCCGGTGATACACCAGATGAAGCAGATACGCCTCTATC**

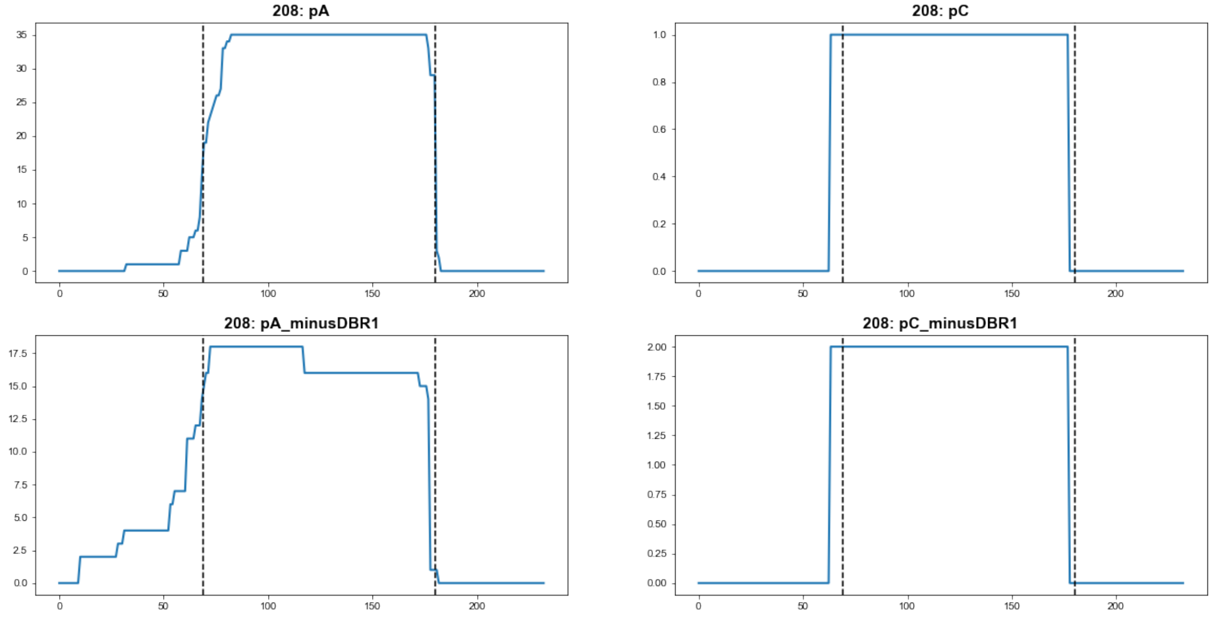


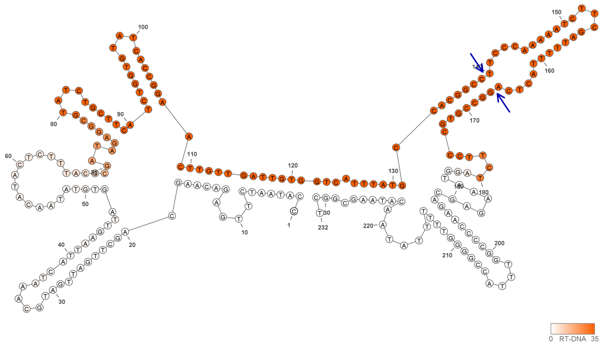


**208**

### Retron Terminal 257: Proteobacteria

**Bacterial Vector for RTDNA: pSLS.589**

**Bacterial Vector for Editing: pAGD290**

**Phage Vector for Editing: None**

**Human Vector for Editing: pSLS.824**

**RTDNA Production (relative to Eco1): 0.317920642 by PAGE**

**Bacterial Editing: 1.27042207 percent precise (0.390924794x Eco1)**

**Phage Editing: undetermined**

**Human Editing (demultiplexed): 0 percent precise (0x Eco1)**

**ncRNA:**

**TATCCTTAGTTTGAAGAAAGTGCCTCGGTATTTTCTTTTGTGGTTGGTGTATAGCCGACCCGTTCGT**
**ATTGTTCGGAATTGTAGTACGTAACTACGTAAATTCCGAACCGCTCGGAACTCGCTGAGCGTAATT**
**TACGTAGTTACGTACAGTCGCAGGCTCCTCTAGTAAGGATA**

**RTDNA:**

**TAGAGGAGCCTGCGACTGTACGTAACTACGTAAATTACGCTCAGCGAGTTCCGAGCGGTTCGGAA**
**TTTACGTAGTTACGTACTACAATTCCGAAC**

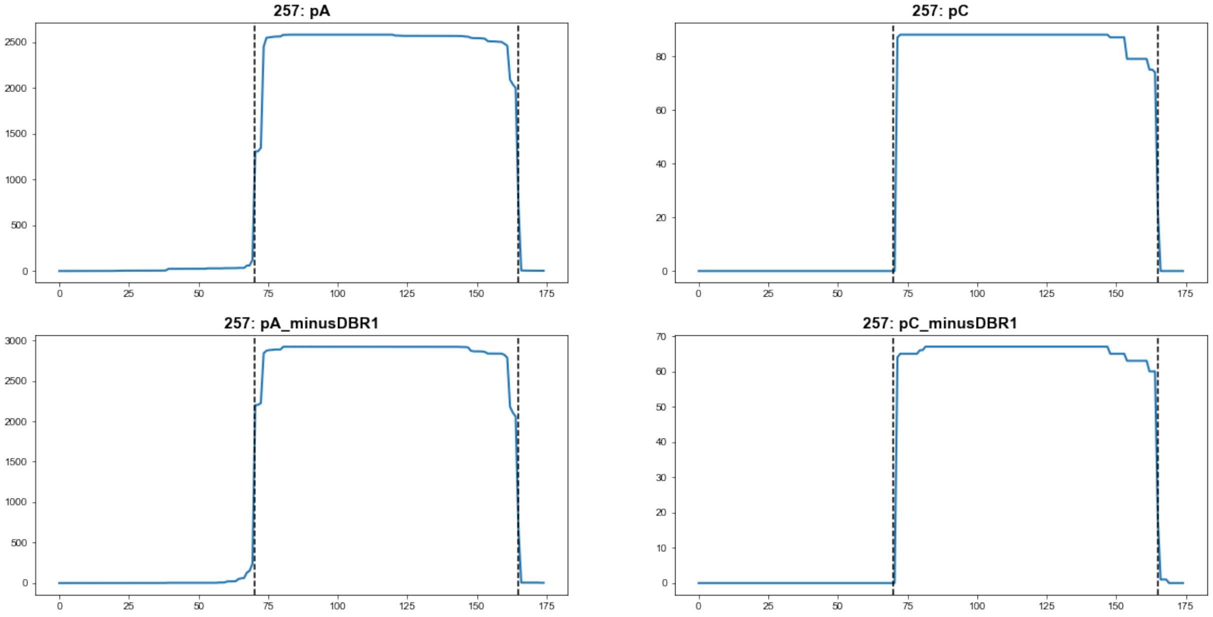


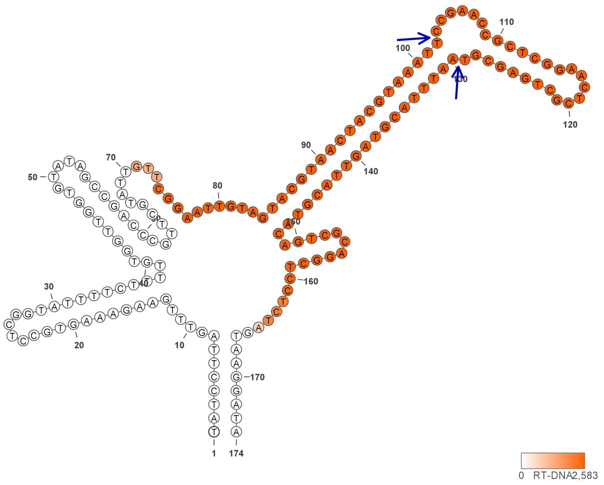


**257**

### Retron Terminal 271: Acidobacteria

**Bacterial Vector for RTDNA: pSLS.655**

**Bacterial Vector for Editing: None**

**Phage Vector for Editing: None**

**Human Vector for Editing: pSLS.900**

**RTDNA Production (relative to Eco1): 0.061622281 by PAGE**

**Bacterial Editing: undetermined**

**Phage Editing: undetermined**

**Human Editing (demultiplexed): 1.231894 percent precise (10.61145136x Eco1)**

**ncRNA:**

**GCCCTTTACGTGGATTGCCGAGATCGTTCATTCGGTGTGTCTGGCGTTTCGCTGGACCGAAATTGA TTGGTAAGATTTACCGGCGCGCAGGAACAGTTTCAGCCTACGCGCTACGCGCTCCGAAGAAACTG TTCCTGCGCGCCTATCTCAAAGGTAAGGGGC**

**RTDNA:**

**TTGAGATAGGCGCGCAGGAACAGTTTCTTCGGAGCGCGTAGCGCGTAGGCTGAAACTGTTCCTGC GCGCCGGTAAATCTTACC**

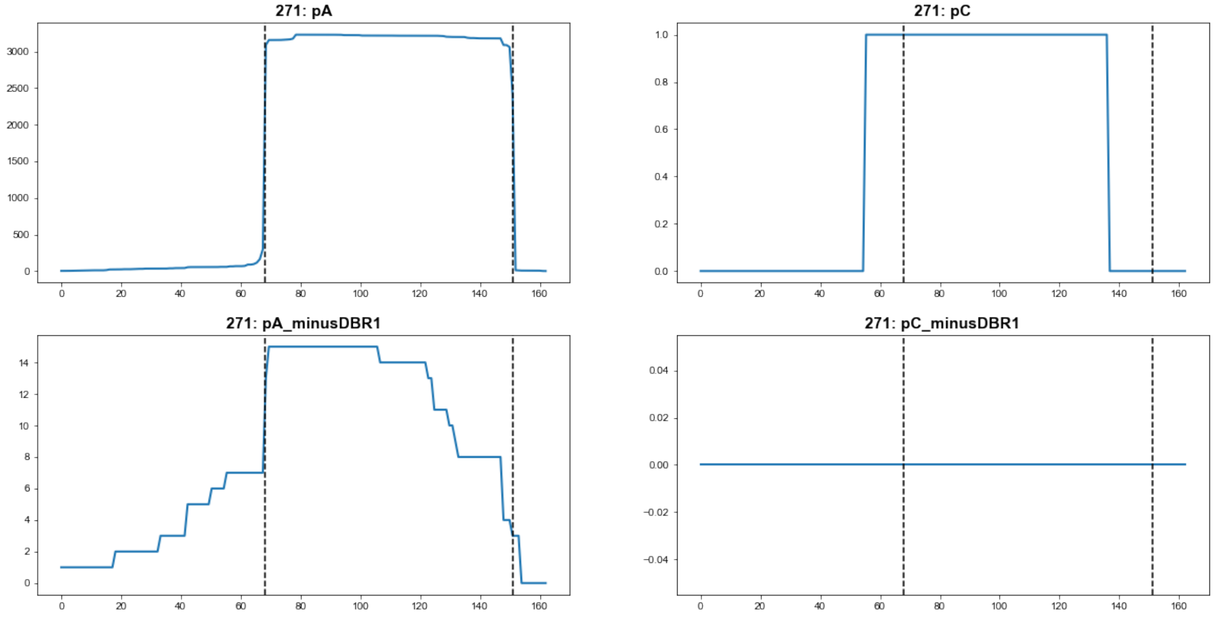


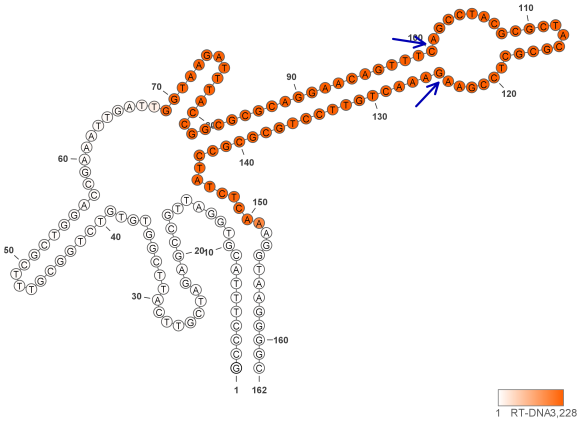


**271**

### Retron Terminal 288: Proteobacteria

**Bacterial Vector for RTDNA: pSLS.577**

**Bacterial Vector for Editing: None**

**Phage Vector for Editing: None**

**Human Vector for Editing: pSLS.820 (H1)**

**RTDNA Production (relative to Eco1): undetermined**

**Bacterial Editing: undetermined**

**Phage Editing: undetermined**

**Human Editing (demultiplexed): 0.009126 percent precise (0.078610745x Eco1)**

**ncRNA:**

**AAAGCCAGACCTAGCAATTTATGGGTTAATAGCCCATCGGGCCATGAGTCATGGTTCCGCCTAGT**
**ATTTTAGCAATGCCCGTCGTTCAGTTCGCTGAGCGGCGGCTGGGGGCCACCGCTCAGCGAACTGA**
**ACGACGTGCTCAAGTAGGTTTGGTTTT**

**no RTDNA sequencing data**

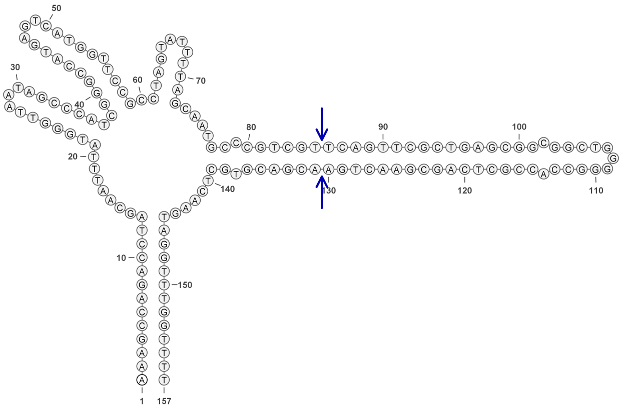

**288**

### Retron Terminal 290: Proteobacteria

**Bacterial Vector for RTDNA: pSLS.578**

**Bacterial Vector for Editing: pAGD282**

**Phage Vector for Editing: None**

**Human Vector for Editing: pSLS.811 (H1)**

**RTDNA Production (relative to Eco1): 0.517779705 by PAGE**

**Bacterial Editing: 1.07932745 percent precise (0.332122584x Eco1)**

**Phage Editing: undetermined**

**Human Editing (demultiplexed): 0.207333 percent precise (1.785952399x Eco1)**

**ncRNA:**

**AGAGCCAAACCTAGCATTTTATGGGTTAATAGCCCATCGGGCCATGAGTCATGGTTTCGCCTAGTA**
**TTTTAGCTATGCCCGTCGATCAGTTCGCTGAGCGGCGGCTGGGGGCCACCGATCAGCGAACTGAT**
**CGACGTGCTCAAGTAGGTTTGGCTCT**

**RTDNA:**

**GCACGTCGATCAGTTCGCTGATCGGTGGCCCCCAGCCGCCGCTCAGCGAACTGATCGACGGGCA**
**TAGCT**

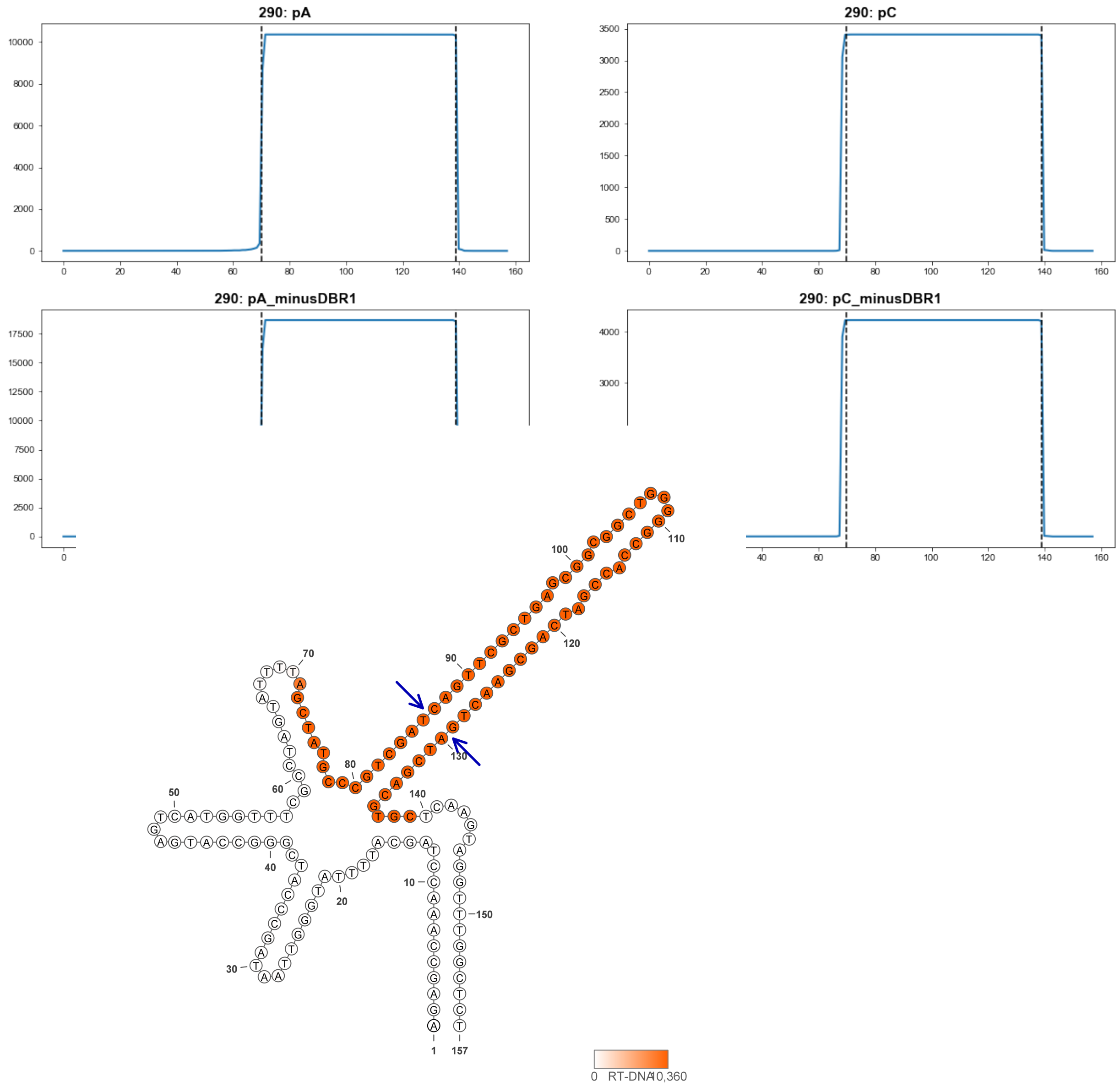


**290**

### Retron Terminal 298: Firmicutes

**Bacterial Vector for RTDNA: pSLS.656**

**Bacterial Vector for Editing: None**

**Phage Vector for Editing: None**

**Human Vector for Editing: pSLS.901**

**RTDNA Production (relative to Eco1): 0 by PAGE**

**Bacterial Editing: undetermined**

**Phage Editing: undetermined**

**Human Editing (demultiplexed): 1.539784 percent precise (13.26359494x Eco1)**

**ncRNA:**

**AAAATAAAGCTAAGAGATTCTAGTATTAATCTAAAGGCCTATGGCCTTTGGTTCTCTGAGTTGAGTT**
**CAGAGAATGAAAGATTGCTGACATTCGTGGCGAGTTAAAGGACTCGAACCGATGCCAGCATTGGT**
**TCGAGTCCTTTAACTCGCTTCCGGTTTAATGTAGAGTCTCTTTTTATTTT**

**RTDNA: None**

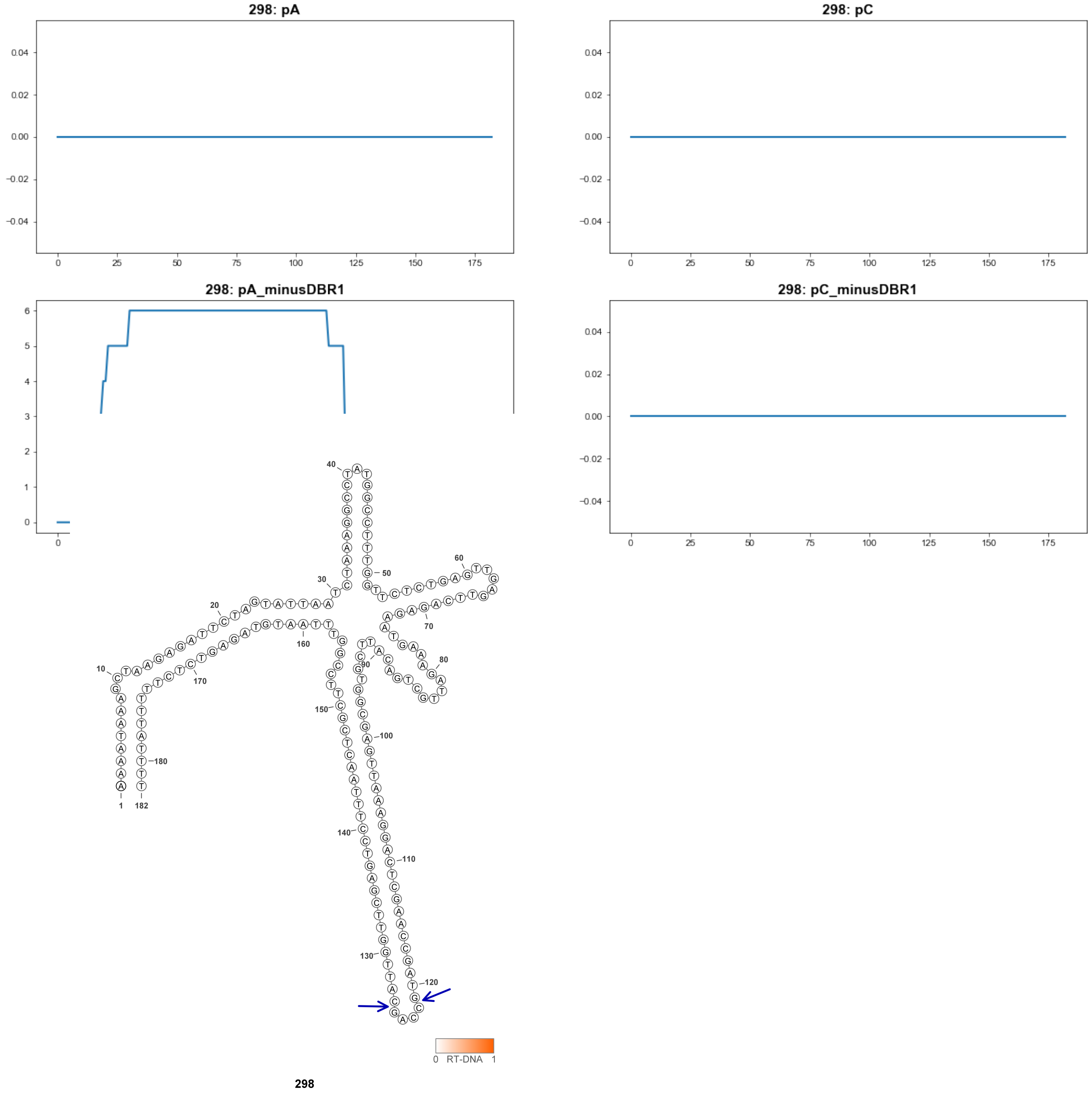

### Retron Terminal 308: Proteobacteria

**Bacterial Vector for RTDNA: pSLS.657**

**Bacterial Vector for Editing: None**

**Phage Vector for Editing: None**

**Human Vector for Editing: None**

**RTDNA Production (relative to Eco1): 0 by PAGE**

**Bacterial Editing: undetermined**

**Phage Editing: undetermined**

**Human Editing (demultiplexed): undetermined**

**ncRNA:**

**TGCATGCATGTCAGCGACTAGGAGCATCTCCAGTCATCAGCCTGACCTCTAGTCGCCTTTATCTGG TTTTACTGTTGTGTGGCTCGCTAGCGAAATCCTCTAGCGAGCCACACAAATAGTAAAAAATCCAGT TTTCTGCTTTTGCATCTGCCGCGTCA**

**RTDNA:**

**TTTTTTACTATTTGTGTGGCTCGCTAGAGGATTTCGCTAGCGAGCCACACAACAGTAAAACCAGAT AAAGGCGACTAGAGGTCAGGCTGATGACTGGAGATGCTCCTAGTCGCTGAC**

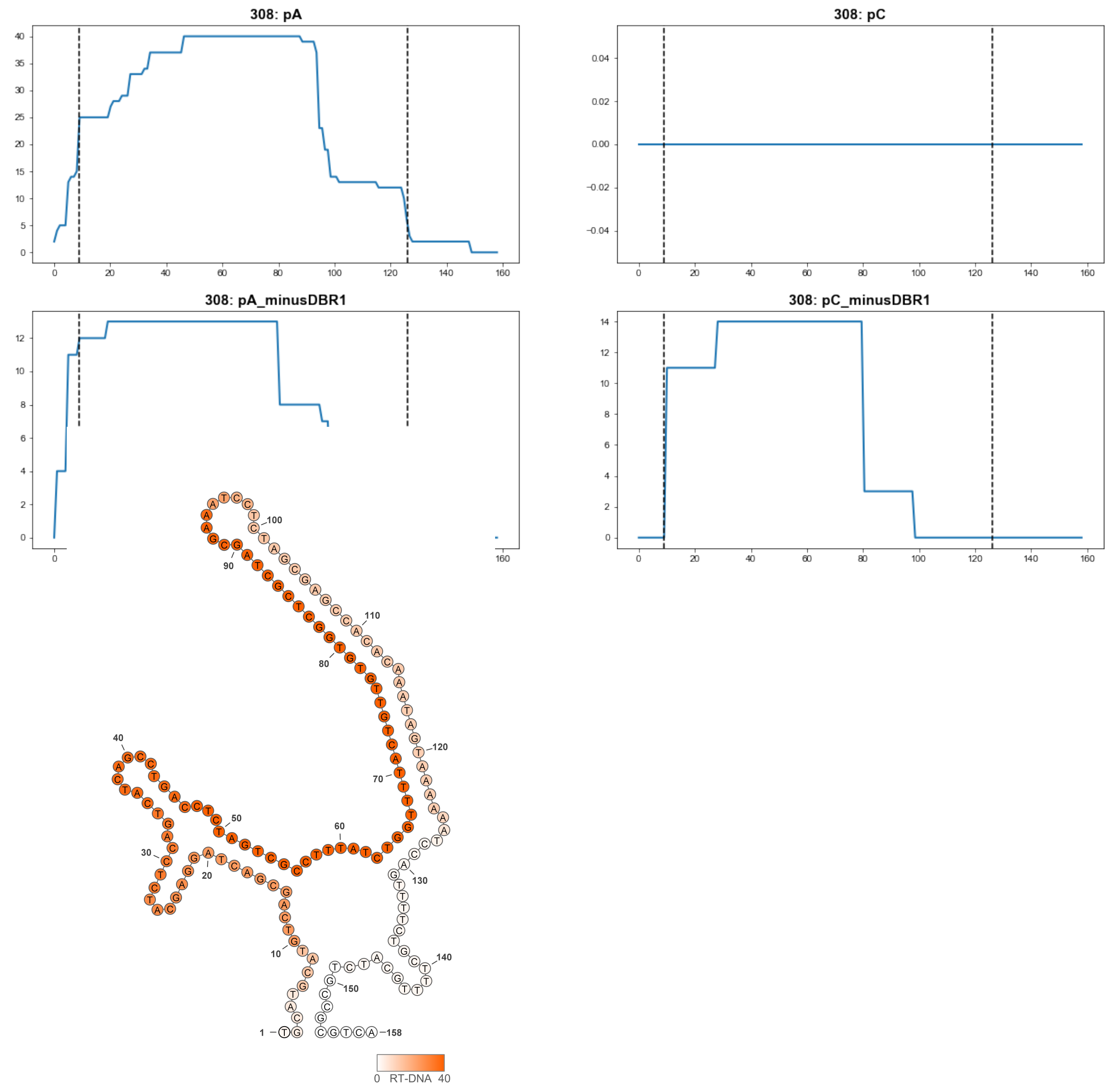


308

### Retron Terminal 314: Proteobacteria

**Bacterial Vector for RTDNA: pSLS.658**

**Bacterial Vector for Editing: None**

**Phage Vector for Editing: None**

**Human Vector for Editing: pSLS.951**

**RTDNA Production (relative to Eco1): 0 by PAGE**

**Bacterial Editing: undetermined**

**Phage Editing: undetermined**

**Human Editing (demultiplexed): undetermined**

**ncRNA:**

**CGGCTCTGCATCTACTTTACGATTGGGCGGTGCGCCTCCTCCAGGCCGCCGCCCAAATTGTAACC**
**GCTGTCAAAAGCGACCTTCTCGCTCCCCCACTCTGCCACCCGTTCCGTCATCTTTTATGTGACGAT**
**GGCGTTGCATCGGCACTACGCACATGCCGACAGCACAATCGCACTCGGTCGGTCGTTTCGCGTGT**
**CGCGAGACGGGTCCGTTGACCTTACCTCACTCGTCTCCATGACTAAGAAGCCACTCGCAGAGCTG**

**RTDNA: None**

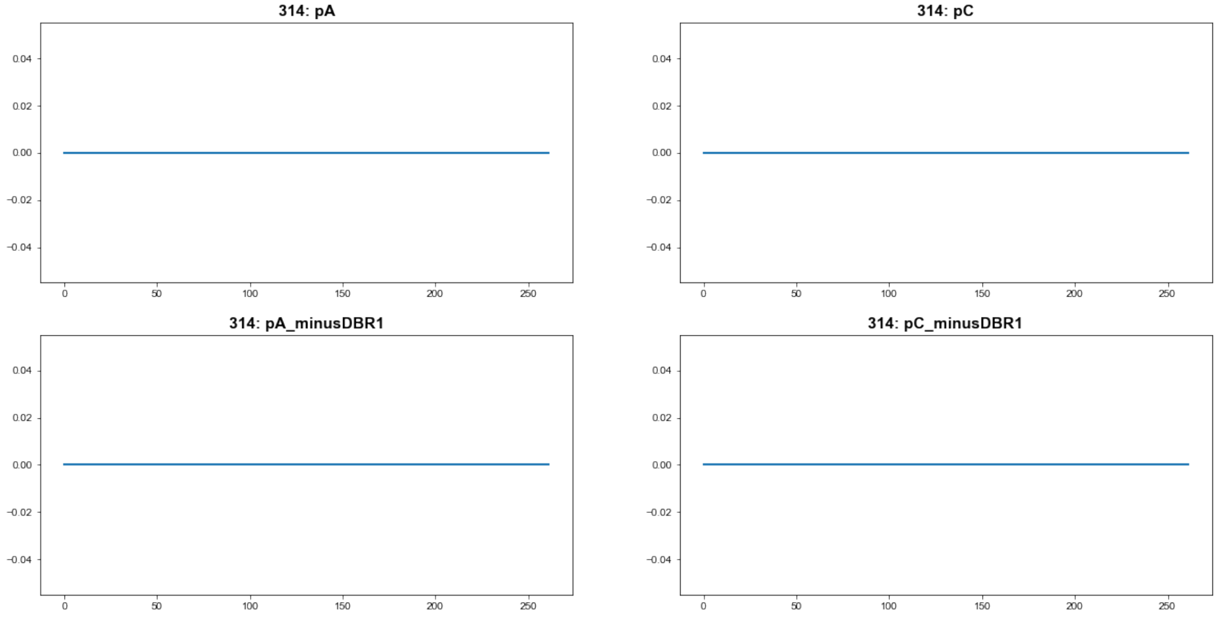


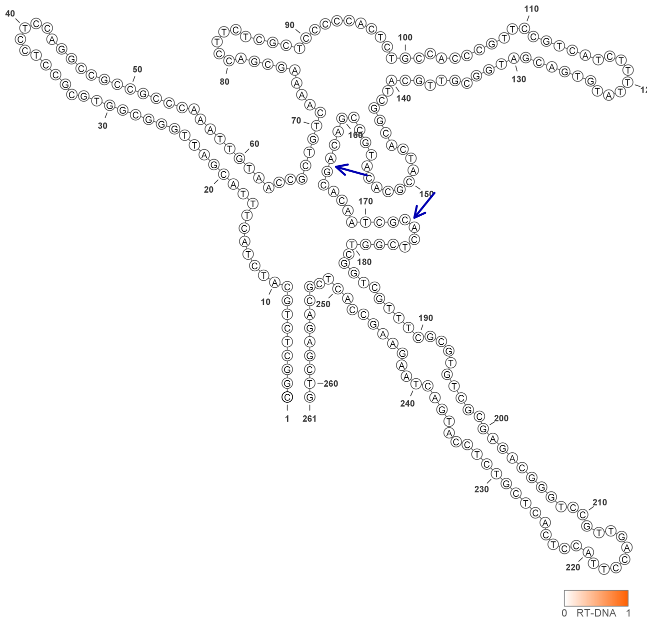


314

### Retron Terminal 348: Proteobacteria

**Bacterial Vector for RTDNA: pSLS.660**

**Bacterial Vector for Editing: None**

**Phage Vector for Editing: None**

**Human Vector for Editing: pSLS.902**

**RTDNA Production (relative to Eco1): 0 by PAGE**

**Bacterial Editing: undetermined**

**Phage Editing: undetermined**

**Human Editing (demultiplexed): 0 percent precise (0x Eco1)**

**ncRNA:**

**GCTGGCCTCAACTGAAACAGAGAAAATGAGCGGCGATTCGCCGTTCTTAACGTTAAGGCACATTT**
**TAAAAGAGGTTCCGCACCCAAGCCTGCCTTGGGCGACCGGGACCCTGCCCAGCCGCCCAAGGCA**
**GGCTTGGGTGCGGATCTATGAGCAATTAAAGATTGGTTCGACGTTCGGTGATGAGGCCAGC**

**RTDNA: None**

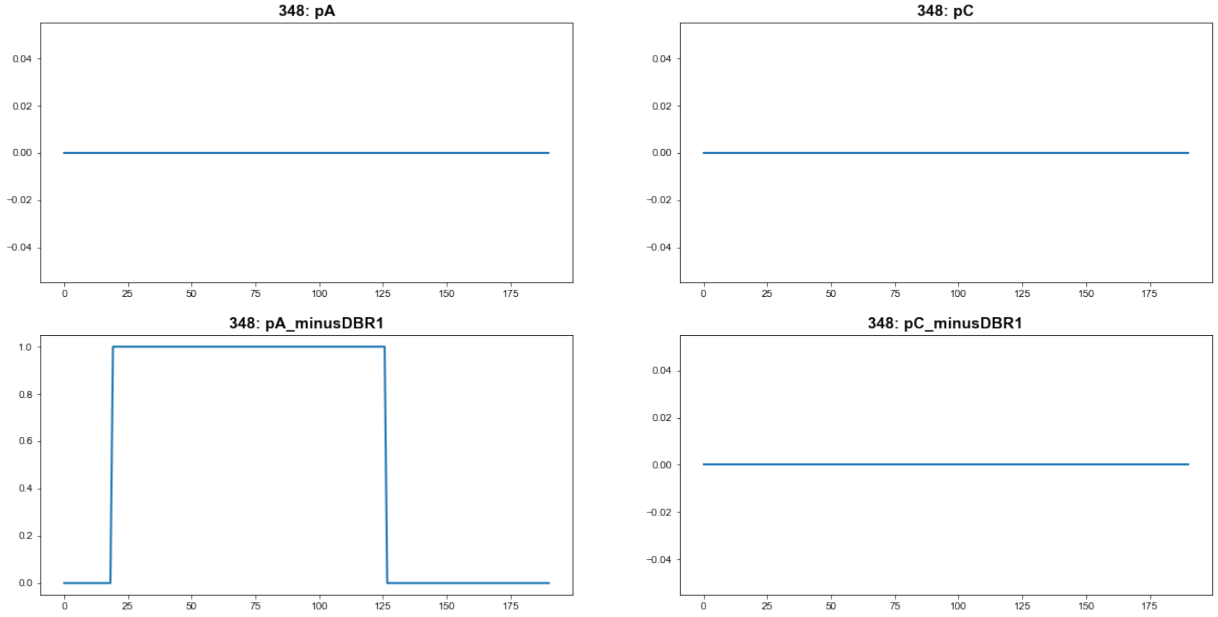


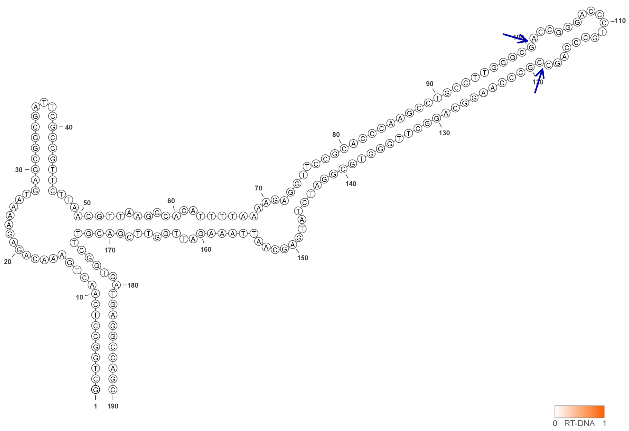


**348**

### Retron Terminal 370: Bacteroidetes

**Bacterial Vector for RTDNA: None**

**Bacterial Vector for Editing: None**

**Phage Vector for Editing: None**

**Human Vector for Editing: pSLS.952**

**RTDNA Production (relative to Eco1): undetermined**

**Bacterial Editing: undetermined**

**Phage Editing: undetermined**

**Human Editing (demultiplexed): 0 percent precise (0x Eco1)**

**ncRNA:**

**TAGCGTCGGAGCCGTGAGTCACTGTTCTGTAATTGAATGTTCTTTCAAGCACTGGTTTAAAAGGCT**
**CTACGCAGAGCTCCGAACCTTTTAAACCCGTATTCCTAATGGGAGTGAGTGACTCTAAAAGTCCTC**
**ACTCCATTAATGATTCAGACGCTA**

**no RTDNA sequencing data**

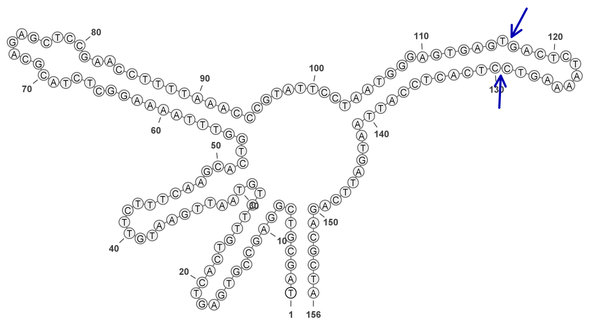

**370**

### Retron Terminal 376: Bacteroidetes

**Bacterial Vector for RTDNA: pSLS.599**

**Bacterial Vector for Editing: None**

**Phage Vector for Editing: None**

**Human Vector for Editing: pSLS.834 (U6)**

**RTDNA Production (relative to Eco1): 0.031602342 by PAGE**

**Bacterial Editing: undetermined**

**Phage Editing: undetermined**

**Human Editing (demultiplexed): 0.459221 percent precise (3.955698547x Eco1)**

**ncRNA:**

**TAAATTATTTTTTTAAATTAATATAATGGGAAGTGAGTCACTAACCCTTATTTGGGAACTAAATTTCA AGCTCAATGAGCTTGTAACTAAGCAAAATCTACGATATTTGCTTAGTTAAAGCTCATTTTTCCGTTT GGGAGCTATGCTCCCAAACATAAAGGAAATAAATAATCATGAAAGATTTTGAATTATATAAAAAGC AATTTA**

**RTDNA:**

**GCTTTTTATATAATTCAAAATCTTTCATGATTATTTATTTCCTTTATGTTTGGGAGCATAGCTCCCAA ACGGAAAAATGAGCTTTAACTAAGCAAATATCGTAGATTTTGCTTAGTTACAAGCTCATTGAGCTT G**

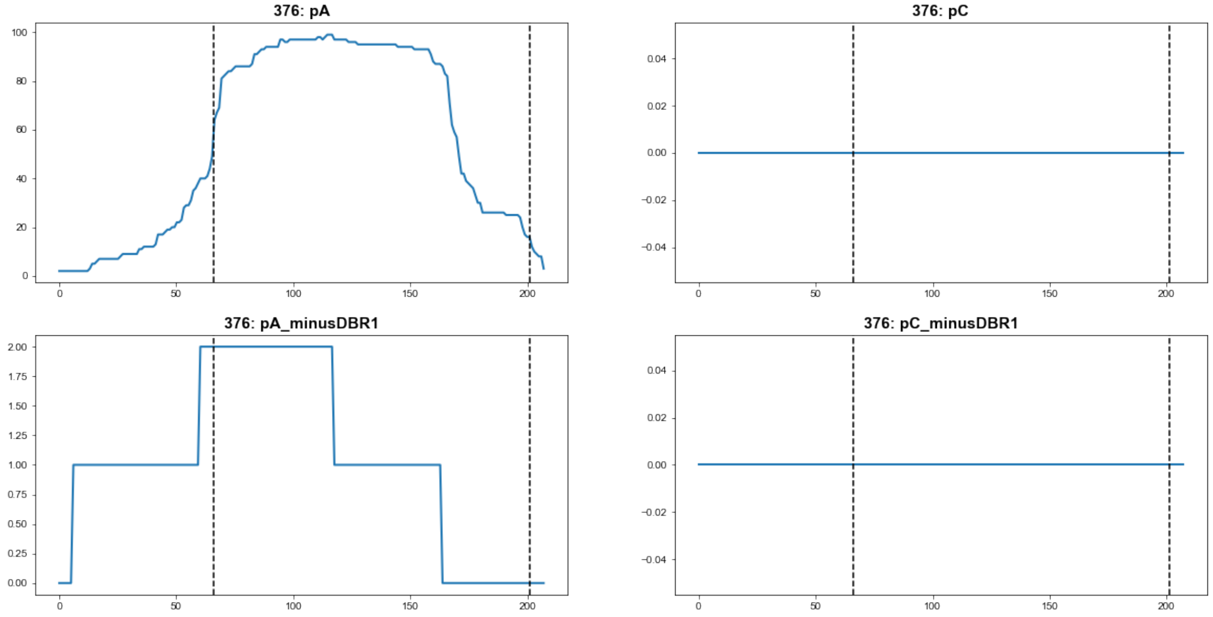


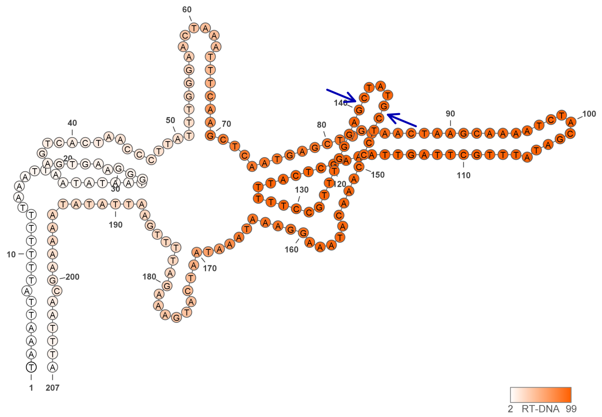


376

### Retron Terminal 397: Bacteroidetes

**Bacterial Vector for RTDNA: pSLS.614**

**Bacterial Vector for Editing: None**

**Phage Vector for Editing: None**

**Human Vector for Editing: None**

**RTDNA Production (relative to Eco1): 0.026777971 by PAGE**

**Bacterial Editing: undetermined**

**Phage Editing: undetermined**

**Human Editing (demultiplexed): undetermined**

**ncRNA:**

**TTTTGTTCCTTCACTTGATGAGTTGCCAGATTAAATAATTATATTTAGATTTGTTTTGGTATCTGATTT CGGATACCATTATAGGGAATTTGACTTCCGAGCAGTTCCGATTCAAAAGATCCTCCAGAGTCGTCC CTTTTGAATCGGCCTTCCAAAATATGGGACCAATTTATTTGGTCCCTTTGAGATTATTAAAATATGG ATTTTATTAGTTATAAGCATAAATTTATATTAGAAGCACAAAA**

**RTDNA:**

**TATAACTAATAAAATCCATATTTTAATAATCTCAAAGGGACCAAATAAATTGGTCCCATATTTTGGA AGGCCGATTCAAAAGGGACGACTCTGGAGGATCTTTTGAATCGGAACTGCTCGGAAGTCAAATTC CCTATAATGGTATCCGAAATCAGATACCAAAACAAATCT**

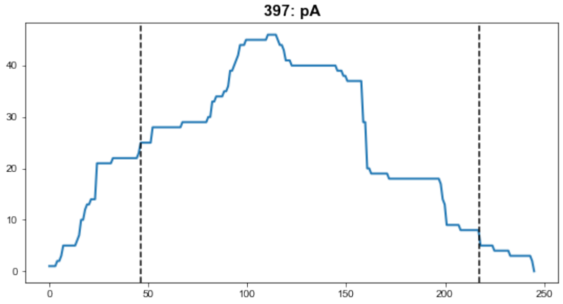


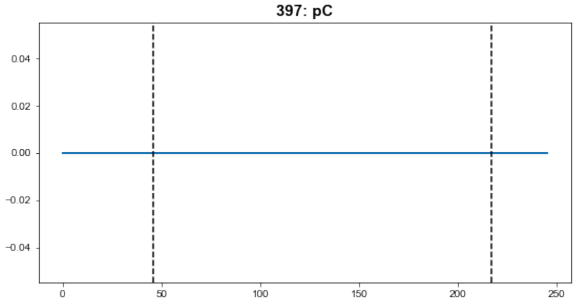


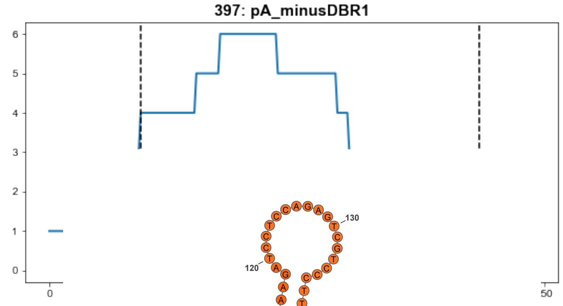


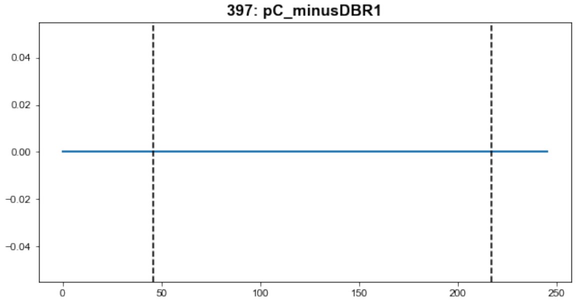


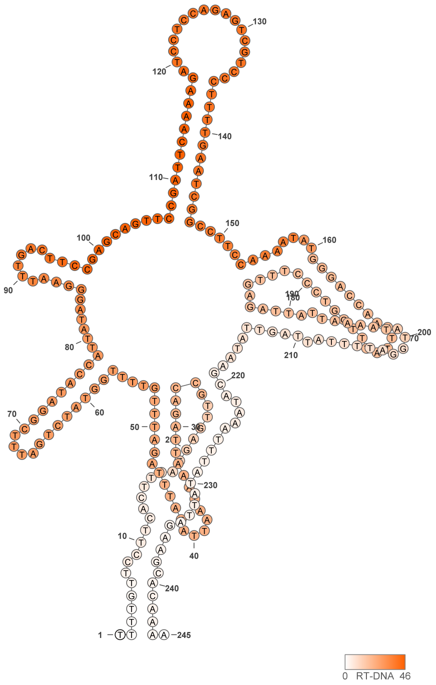


397

### Retron Terminal 398: Bacteroidetes

**Bacterial Vector for RTDNA: pSLS.615**

**Bacterial Vector for Editing: None**

**Phage Vector for Editing: None**

**Human Vector for Editing: pSLS.836 (U6)**

**RTDNA Production (relative to Eco1): 0.02254987 by PAGE**

**Bacterial Editing: undetermined**

**Phage Editing: undetermined**

**Human Editing (demultiplexed): 0.147896 percent precise (1.273966113x Eco1)**

**ncRNA:**

**GGATTTCTCCAAAAATTGATGAAATAATTTCTCCAATCCAAGAAAATTATATATATTTGCCGAATAA
TACAAAAAGTTTAAAATAACATATCAATTTGTCCTAGGGGGCAAGCGGTTGTATGAGTCATACTGC
TGTATAGGTATAAGGTTTCAAGCGATTCAGGCTTCTAATGTTGTTTCAGCGGTCAGAGCCCGCTTC
CACAACATTAGAGCCTGATGTCAGAAGACAAAGATGTCAGGAGGCTAGCAATGCTAGCCTCCTAA
AAAGATTTTAAATAATGGACTTTAAAGAATATAAACGGCTTTTTACAGAAAAAGCACAGAATAATG
GGAGGGATTC**

**RTDNA: None**

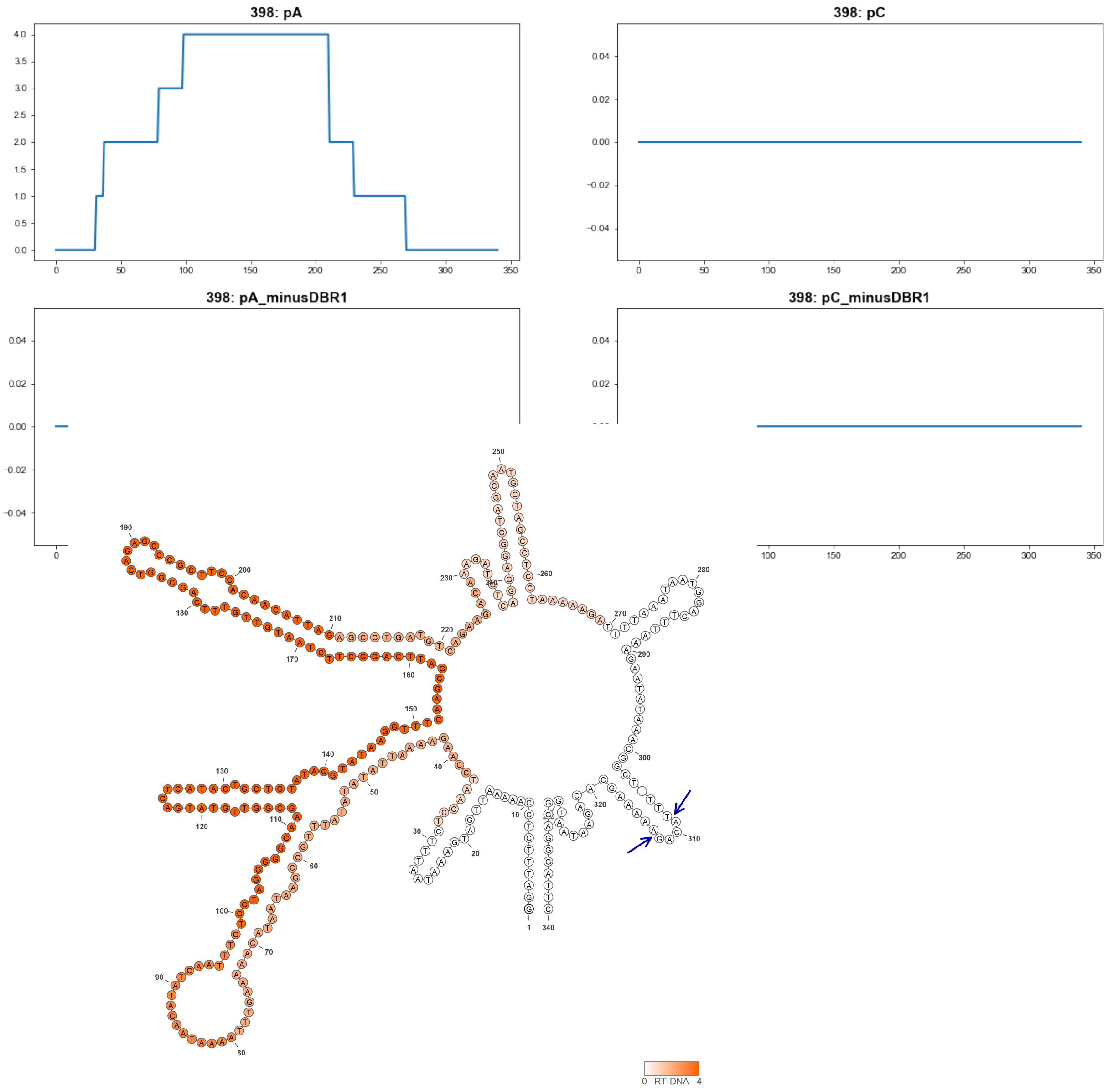


**398**

### Retron Terminal 400: Proteobacteria

**Bacterial Vector for RTDNA: pSLS.579**

**Bacterial Vector for Editing: pAGD283**

**Phage Vector for Editing: pAGD394**

**Human Vector for Editing: pSLS.812 (H1)**

**RTDNA Production (relative to Eco1): 0.94628144 by PAGE**

**Bacterial Editing: 2.15208712 percent precise (0.66222418x Eco1)**

**Phage Editing: 36.03613898 percent precise (0.888819556x Eco1)**

**Human Editing (demultiplexed): 0.847635 percent precise (7.301470398x Eco1)**

**ncRNA:**

**TTGGACATCAGTCATTCGCTCAGATTCATGAGAGAGTTAGACCCTAGCGGCCAGACCGCAGGATG AGCGAGTCGCGCATCTGGAACGGACCTTACTTTAGGCCCACCCAACGAAAACGAATTTTTCACAT AGTGAAAAATATAGTTTTCGTTGTAGCGCCCAAAGGGCGCTGTCACCCAAAGGGTGATGGATATT TTAGTAACTGGAATGGCTGATGTCCAA**

**RTDNA:**

**ATATCCATCACCCTTTGGGTGACAGCGCCCTTTGGGCGCTACAACGAAAACTATATTTTTCACTAT GTGAAAAATTCGTTTTCGTTGGGTGGGCCTAAAGTAAGG**

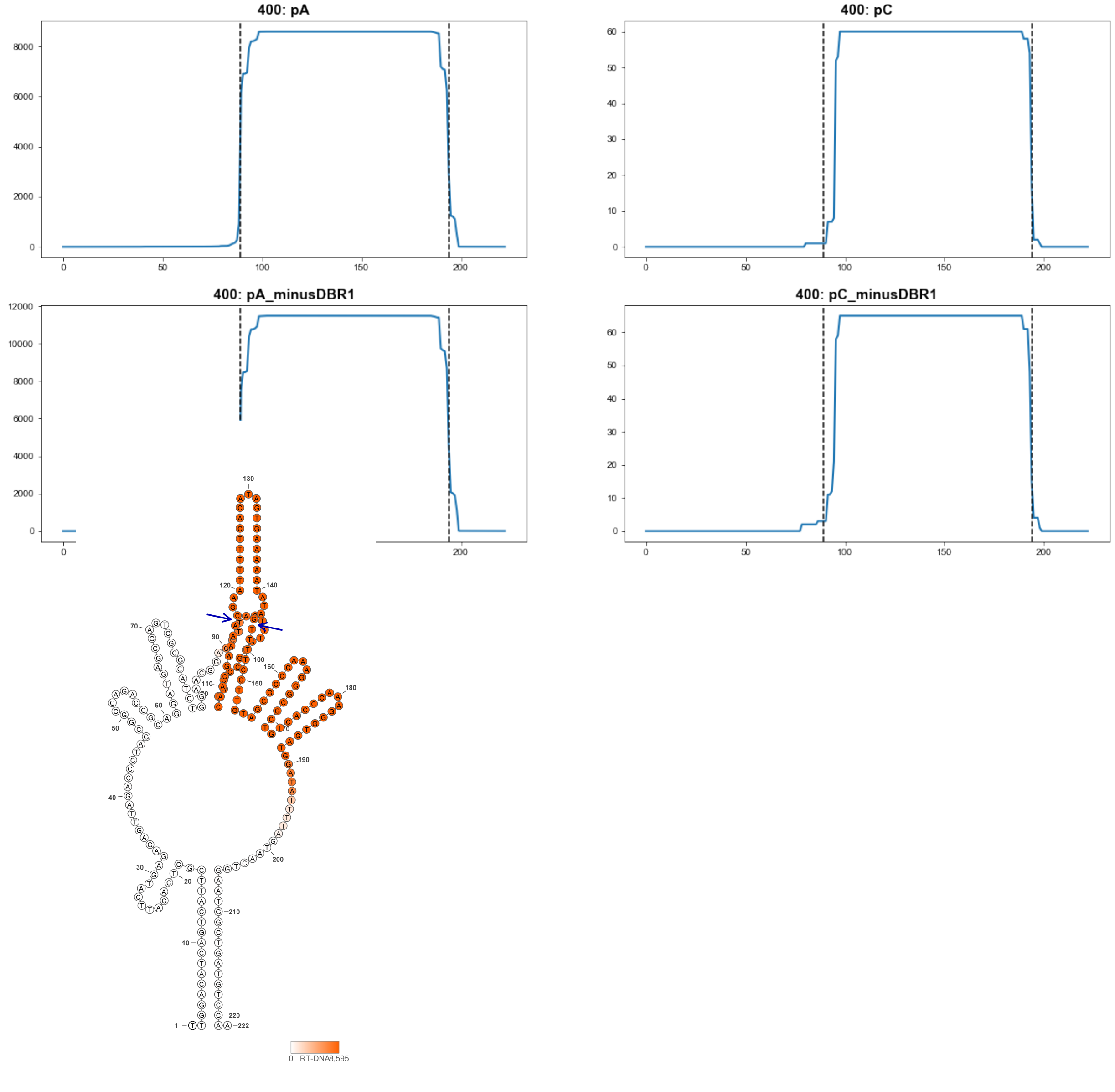


400

### Retron Terminal 401: Proteobacteria

**Bacterial Vector for RTDNA: None**

**Bacterial Vector for Editing: None**

**Phage Vector for Editing: None**

**Human Vector for Editing: pSLS.867**

**RTDNA Production (relative to Eco1): undetermined**

**Bacterial Editing: undetermined**

**Phage Editing: undetermined**

**Human Editing (demultiplexed): 0 percent precise (0x Eco1)**

**ncRNA:**

**TTGAGCATCGAACGTTCACTCAGATCAGAGACAGAGTTAGACCCCAGCGGTAATGCCGCAGGATT
GGCGAGTCGCCAATCTGGATTGGATGGCACTTCAGGCCCATTCAATGGAAATGAATTTTTCACGG
AGCGAAAAATGTAATTTCCATTGAAGCGCCTAAAGGCGCACTACCTGAAGGTAATGGTTATTTAGT
AACTGGAACGTTCGATGCTCAA**

**no RTDNA sequencing data**

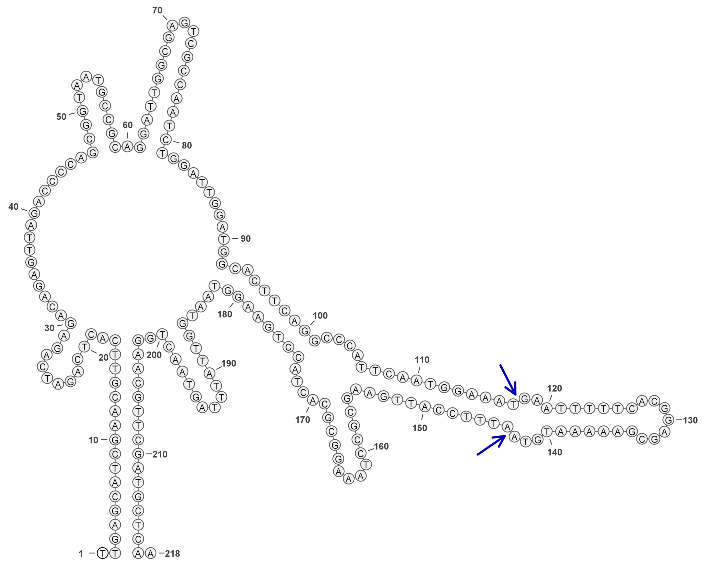

**401**

### Retron Terminal 402: Firmicutes

**Bacterial Vector for RTDNA: None**

**Bacterial Vector for Editing: None**

**Phage Vector for Editing: None**

**Human Vector for Editing: pSLS.868**

**RTDNA Production (relative to Eco1): undetermined**

**Bacterial Editing: undetermined**

**Phage Editing: undetermined**

**Human Editing (demultiplexed): 0 percent precise (0x Eco1)**

**ncRNA:**

**AGGTAGCCCTATTAGAACTTTATGTGGTTTTATGCCACAGGTTTGGCGAGTCGTCAAGCTAAGAAG**
**AACATTATTTACAAGCCTTGGTGATCATAACTTTTCCCTCCGACGGAGTCTACAGAAAAAGTTATG**
**ACCACCACTCCGGCTGATGCCTACGTGACACTTGATTGAAATTAAAGTAAGAAGCTAGACTGAAT**
**GTATTTCAAAAGGCAACTTTAATTTCAATCAAGTGAGAAGATAGAAAATCTGATTTGGTAATATGG**
**GCTACCT**

**no RTDNA sequencing data**

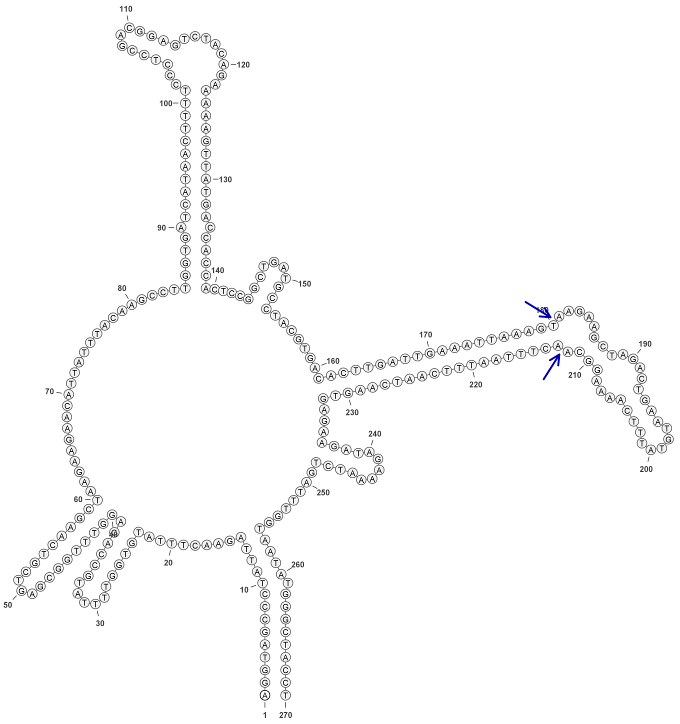

**402**

### Retron Terminal 406: Firmicutes

**Bacterial Vector for RTDNA: None**

**Bacterial Vector for Editing: None**

**Phage Vector for Editing: None**

**Human Vector for Editing: pSLS.869**

**RTDNA Production (relative to Eco1): undetermined**

**Bacterial Editing: undetermined**

**Phage Editing: undetermined**

**Human Editing (demultiplexed): 0.008431 percent precise (0.072624062x Eco1)**

**ncRNA:**

**ATTTCAGGTCTTTTACATCAATACCTACTAGACTTATGGTTTTTTTAAAAAAAAAACCAGGTATGGT**
**GAGTCACCATGCTAAGAAGAACTTAATTTCAAGCCCTGTGTAATACATTTAACGCGTCGGGAGAC**
**GCTATATAAATGTATTACACAACCTTAGGCTTGCGCCACGGTGTACAATGAACATTTTGACGAAAA**
**GAGTATCAAAATGTTCATTGTGGAGGATAGAGACGTGATAGTGGTAAGTGTATGGATGTAAGAGC**
**CTGTAAT**

**no RTDNA sequencing data**

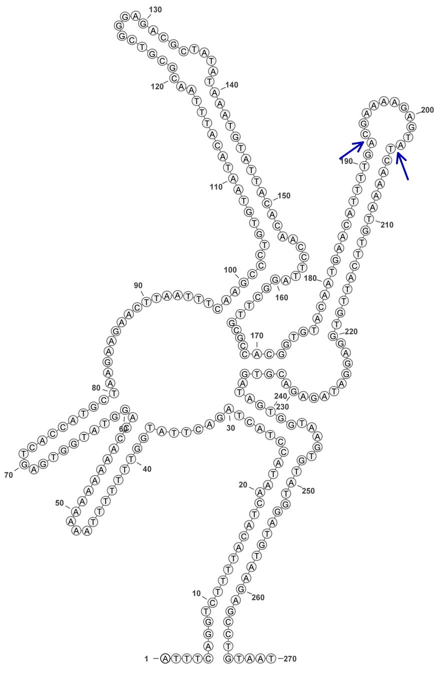

406

### Retron Terminal 412: Proteobacteria

**Bacterial Vector for RTDNA: pSLS.625**

**Bacterial Vector for Editing: None**

**Phage Vector for Editing: None**

**Human Vector for Editing: pSLS.852**

**RTDNA Production (relative to Eco1): 0.028892021 by PAGE**

**Bacterial Editing: undetermined**

**Phage Editing: undetermined**

**Human Editing (demultiplexed): 0.583286 percent precise (5.024386042x Eco1)**

**ncRNA:**

**CGGCCTCACTAGCCTTAGTGGCTTAGCCACGGAATGGCGAGTCGCCATTCTGGGGAGAACTTGCT**
**TTCAAGCCCAGGCGAAGATTCACCAACTCTTGGAAAGAGTTGGCGAATCTTCGCCCCGCTTCCGC**
**AAAAGCCTCCAGCCCCGAGATGCAGGACAAATCCTTGCATCTCGTCCTTGAGGGCCGAAATGCAA**
**GGATTTGTCCTGCATCTCGGTGATTATGTCCGTATGTAGTGAGGCCG**

**RTDNA: None**

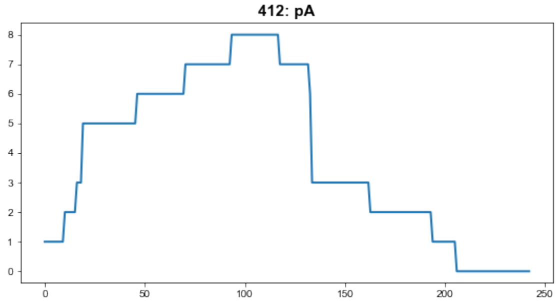


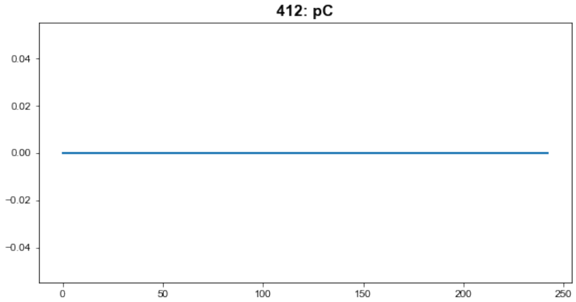


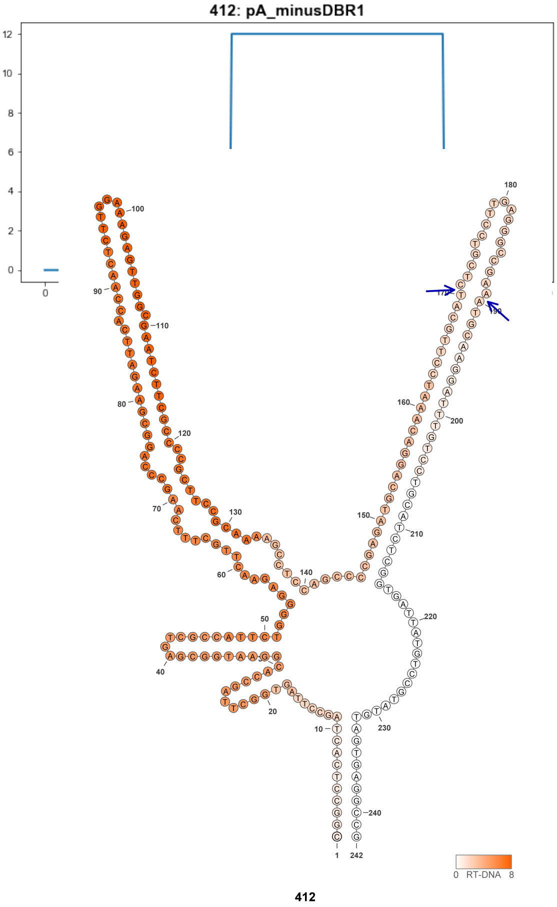


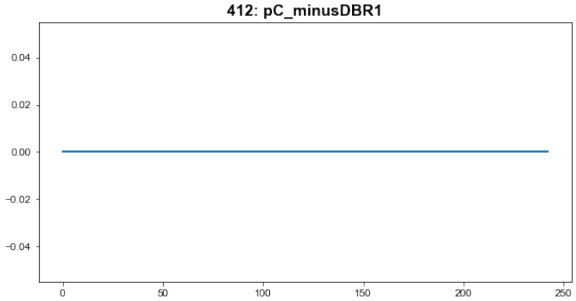

### Retron Terminal 412.1: Proteobacteria

**Bacterial Vector for RTDNA: None**

**Bacterial Vector for Editing: None**

**Phage Vector for Editing: None**

**Human Vector for Editing: pSLS.862 (8bp stem)**

**RTDNA Production (relative to Eco1): 0.028892021 by PAGE**

**Bacterial Editing: undetermined**

**Phage Editing: undetermined**

**Human Editing (demultiplexed): undetermined**

**ncRNA:**

**CGGCCTCACTAGCCTTAGTGGCTTAGCCACGGAATGGCGAGTCGCCATTCTGGGGAGAACTTGCT**
**TTCAAGCCCAGGCGAAGATTCACCAACTCTTGGAAAGAGTTGGCGAATCTTCGCCCCGCTTCCGC**
**AAAAGCCTCCAGCCCCGAGATGCAGGACAAATCCTTGCATCTCGTCCTTGAGGGCCGAAATGCAA**
**GGATTTGTCCTGCATCTCGGTGATTATGTCCGTATGTAGTGAGGCCG**

**RTDNA: None**

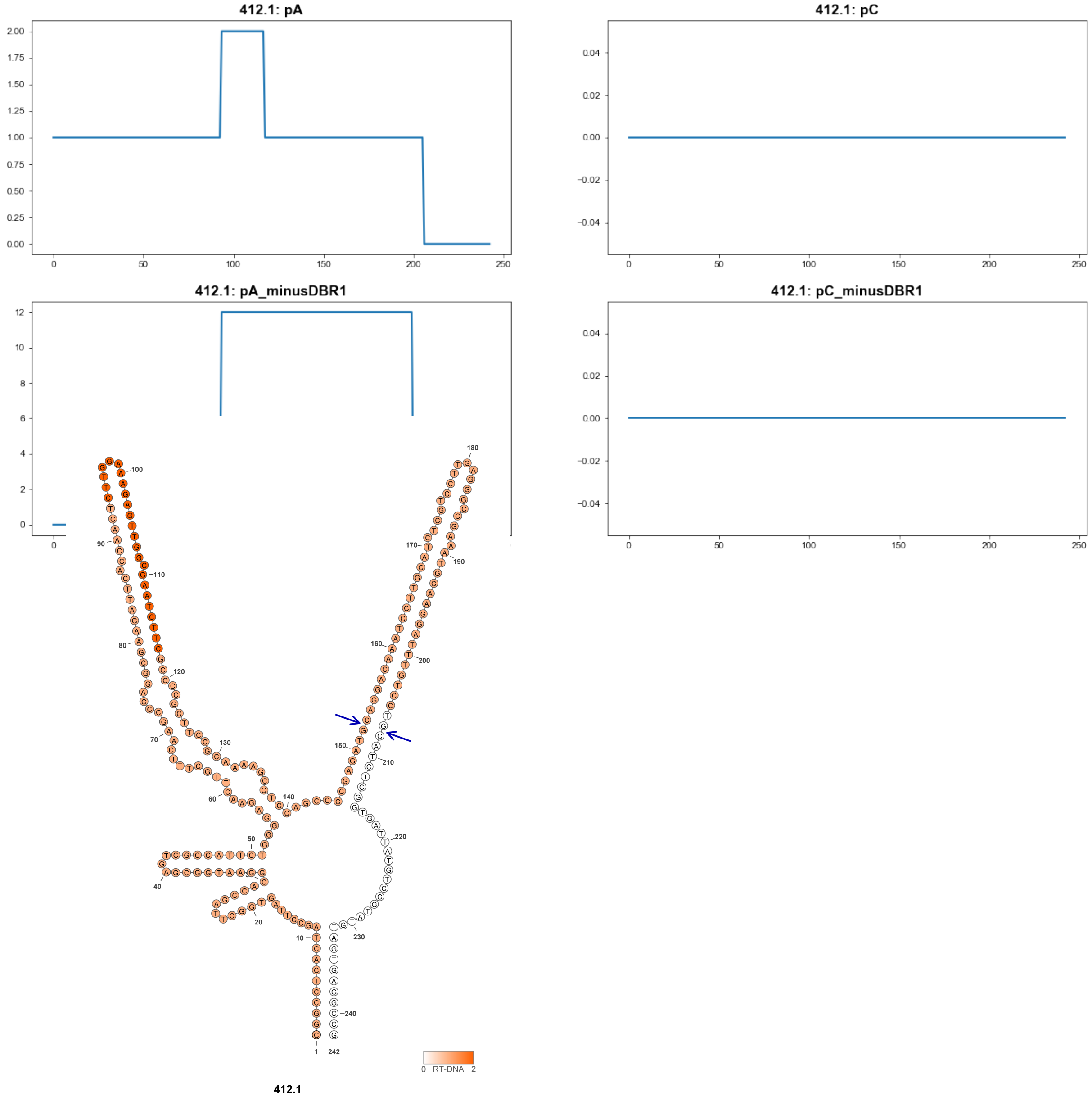

### Retron Terminal 418: Proteobacteria

**Bacterial Vector for RTDNA: None**

**Bacterial Vector for Editing: None**

**Phage Vector for Editing: None**

**Human Vector for Editing: pSLS.870**

**RTDNA Production (relative to Eco1): undetermined**

**Bacterial Editing: undetermined**

**Phage Editing: undetermined**

**Human Editing (demultiplexed): 0.065697 percent precise (0.565909502x Eco1)**

**ncRNA:**

**AGCATGTAGAGAGTGGGTGTTCAAGTTAGCACATGCCGAAAGGCAAGGCCGGCGAGTCGCCGAG**
**CCTGAAGCGAACTATCATTCAAGCCTCAGGCCAAAAGCGTCATTGCGAGCAGGAGCTCGCAACG**
**CCGTTTTGGCCTGCGCCATGAAACCGGCGCACTGAAGCCGAAAAGCACGGCTTCAGTGTCAGAA**
**GGAAGGAAATAGTAACTTGAACGCCCACTCCCTCCATGCT**

**no RTDNA sequencing data**

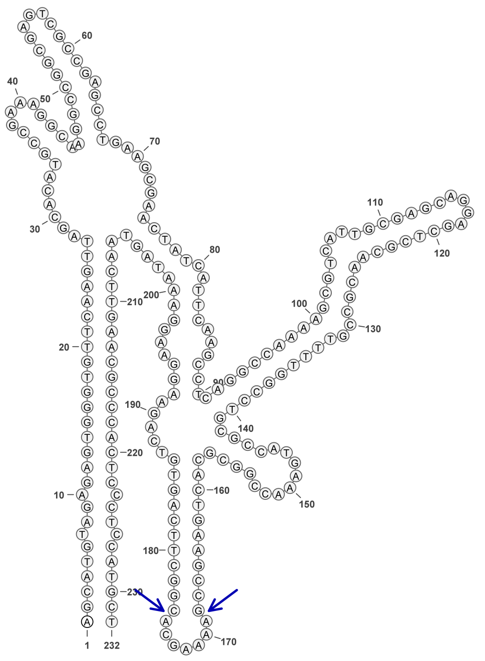

418

### Retron Terminal 430: Firmicutes

**Bacterial Vector for RTDNA: pSLS.638**

**Bacterial Vector for Editing: pAGD296**

**Phage Vector for Editing: pAGD395**

**Human Vector for Editing: pSLS.1010**

**RTDNA Production (relative to Eco1): 2.339048049 by PAGE**

**Bacterial Editing: 6.14085249 percent precise (1.889617277x Eco1)**

**Phage Editing: 3.880576222 percent precise (0.09571314x Eco1)**

**Human Editing (demultiplexed): undetermined**

**ncRNA:**

**GGCAGCATATCCGGGAAAATATTTTCCGGCCTGACGGGGAGTCCCCGCCGGGAGTTAGAAACAG**
**TGGGCATGCAGAATACTGCACCCGTAATCCGCTCCAAAGTACATCCGCACACTTCCGTGTGCACA**
**CCGCCTTCGGCTGTACATTTCCGCAGCTTACGGTATCGGTGTATTGTGCCGTCCGGCTTATCAAAG**
**CTATGAGCAACGCGTGAGGATAAAAGAGGAACTGCATGGGATGCTCACATGCGGTTCCTCTTTTT**
**GGTTTTTACCGTGAAAATATTTTCCATGCTGTC**

**RTDNA:**

**GGACGGCACAATACACCGATACCGTAAGCTGCGGAAATGTACAGCCGAAGGCGGTGTGCACACG**
**GAAGTGTGCGGATGTACTTTGGAGCGGATTACGGGTGCAGTATTCTGCATG**

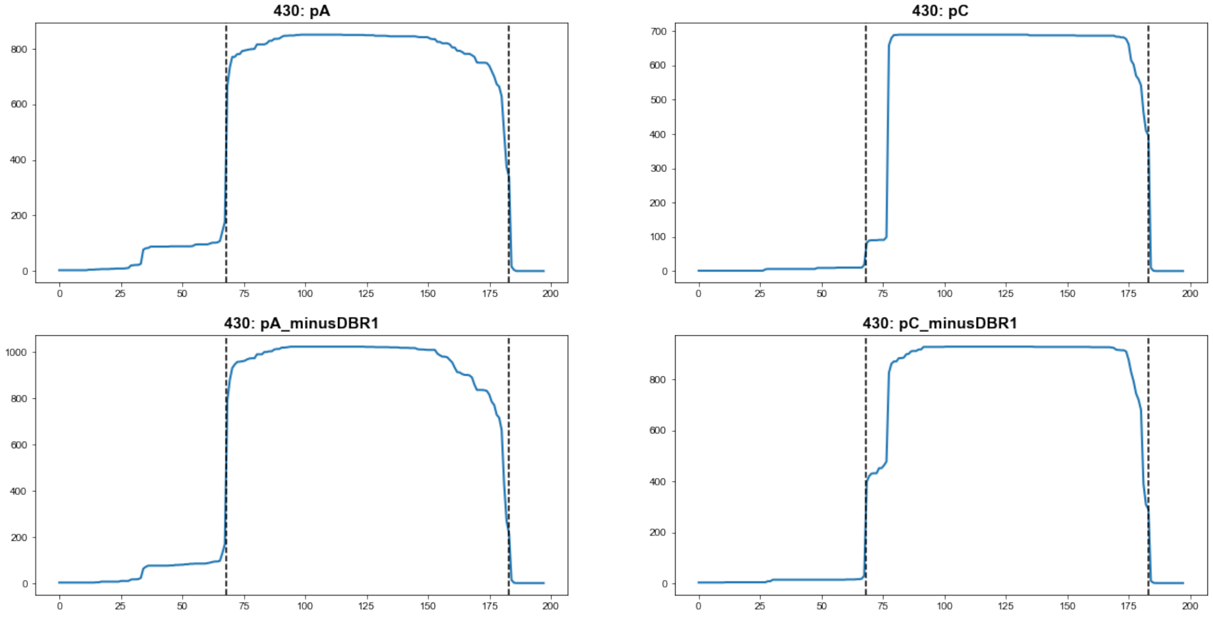


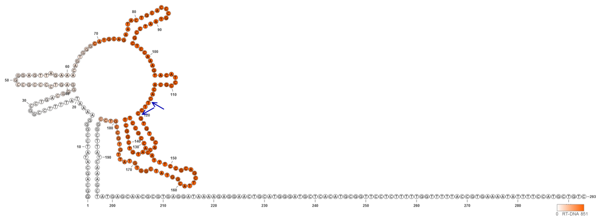

**430**

### Retron Terminal 443: Proteobacteria

**Bacterial Vector for RTDNA: pSLS.626**

**Bacterial Vector for Editing: None**

**Phage Vector for Editing: None**

**Human Vector for Editing: pSLS.853**

**RTDNA Production (relative to Eco1): 0.649121856 by PAGE**

**Bacterial Editing: undetermined**

**Phage Editing: undetermined**

**Human Editing (demultiplexed): 0.432163 percent precise (3.72262277x Eco1)**

**ncRNA:**

**GATAGAGGCAATAGCTTATCCGGGGAATCTTATCCGGATCCGCTGGCGAGTCGTCACCGGATAAT
CGATATCTGAGCTATAGACCCTAAGTTGCTCCGAAGTACCTCAGCACACTTCGTGTGCGACGCCTT
TGGCTGAGTACGTCTGCGCACCTTAGGGTATCGTTTTATAGTATTGCCTCTATC**

**RTDNA:**

**AAAACGATACCCTAAGGTGCGCAGACGTACTCAGCCAAAGGCGTCGCACACGAAGTGTGCTGAG
GTACTTCGGAGCAACTTAGGGTCTATAGCTCAGAT**

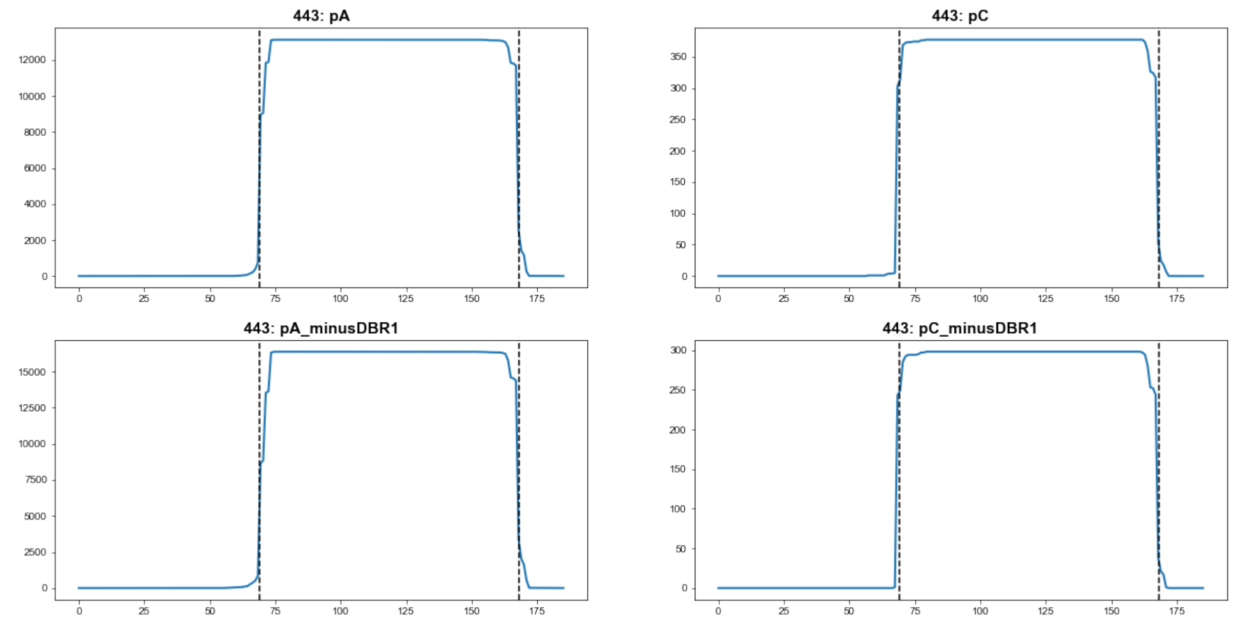


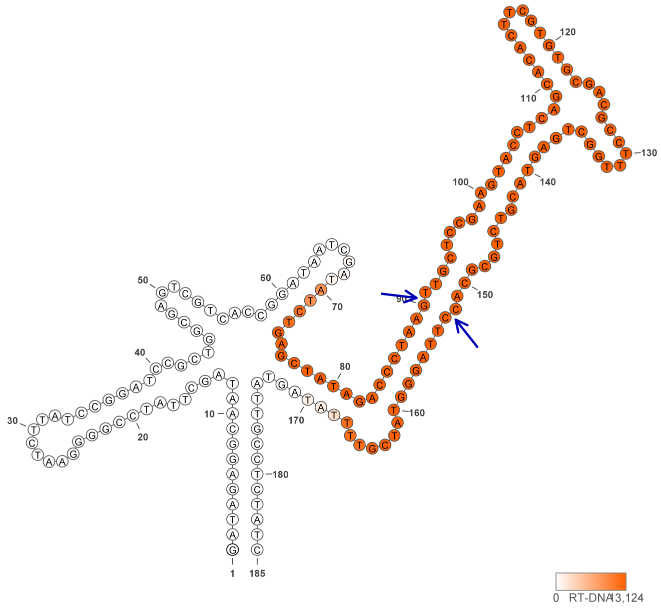


443

### Retron Terminal 466: Proteobacteria

**Bacterial Vector for RTDNA: None**

**Bacterial Vector for Editing: None**

**Phage Vector for Editing: None**

**Human Vector for Editing: pSLS.804 (H1)**

**RTDNA Production (relative to Eco1): undetermined**

**Bacterial Editing: undetermined**

**Phage Editing: undetermined**

**Human Editing (demultiplexed): 0.004326 percent precise (0.037263871x Eco1)**

**ncRNA:**

**CGCCAGCAGTGGCAATAGCGTTTCCGGCCTTTTGTGCCGGGAGGGTCGGCGAGTCGCTGACTTAA**
**CGCCAGTAGTATGTCCATATACCCAAAGTCGCTTCATTGTACCTGAGTACGCTTCGCGTACGTCGC**
**GCTGACGCGCTCAGTACAGTTACGCGCCTTCGGGATGGTTTAATGGTATTGCCGCTGTTGGCG**

**no RTDNA sequencing data**

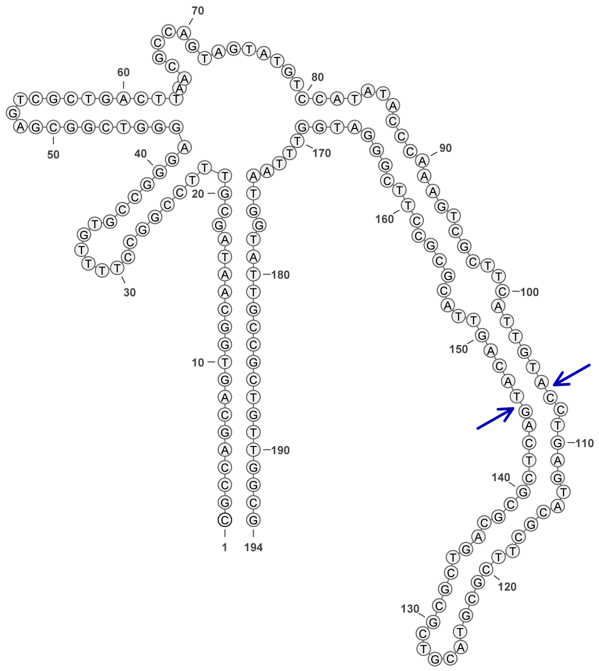


**466**

### Retron Terminal 482: Firmicutes

**Bacterial Vector for RTDNA: pSLS.661**

**Bacterial Vector for Editing: None**

**Phage Vector for Editing: None**

**Human Vector for Editing: pSLS.903**

**RTDNA Production (relative to Eco1): 1.111439727 by PAGE**

**Bacterial Editing: undetermined**

**Phage Editing: undetermined**

**Human Editing (demultiplexed): 6.252318 percent precise (53.85704318x Eco1)**

**ncRNA:**

**AAACAGCCGTCGGCAGCGAATTTGCACAAGCAAAGCTGCGGCGAGTCGCCGCGGATGATTTCAG GAATATTACCGATATAAAAAGTTCTCCACCGAATCATGAATTAACTTCGTTAATAGAAATTCGGCT CCGAACATTTTAATGCTTTTGTGCCGTCGGCTGTTT**

**RTDNA:**

**AAAAGCATTAAAATGTTCGGAGCCGAATTTCTATTAACGAAGTTAATTCATGATTCGGTGGAGAAC TTTTTATATCGGTAATATTC**

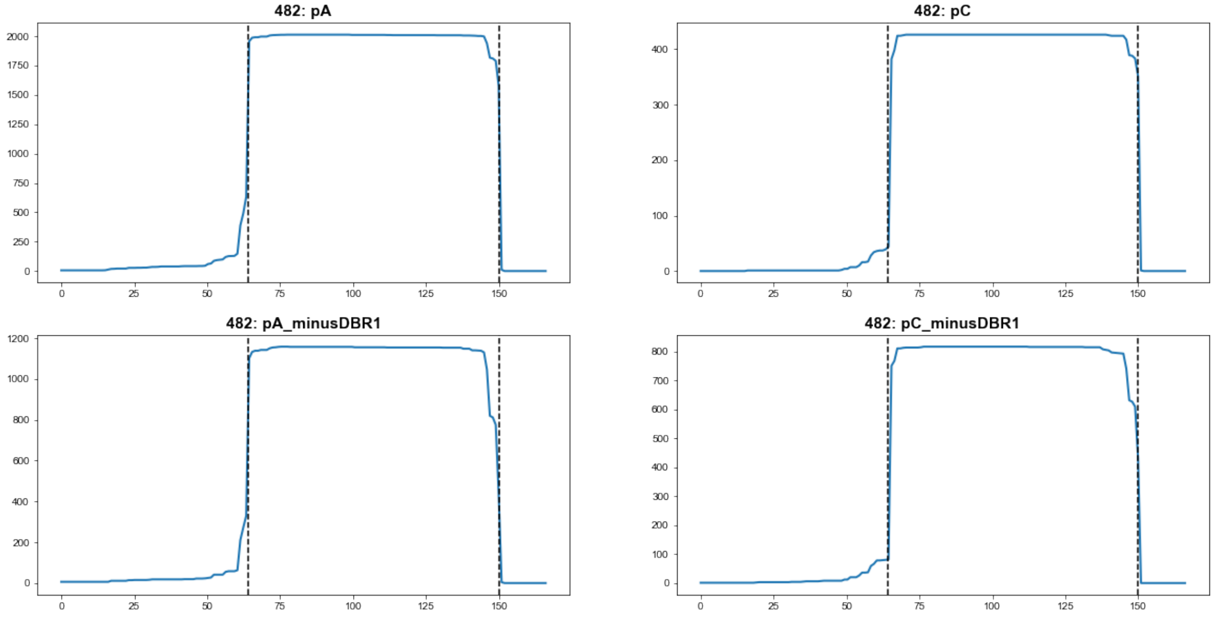


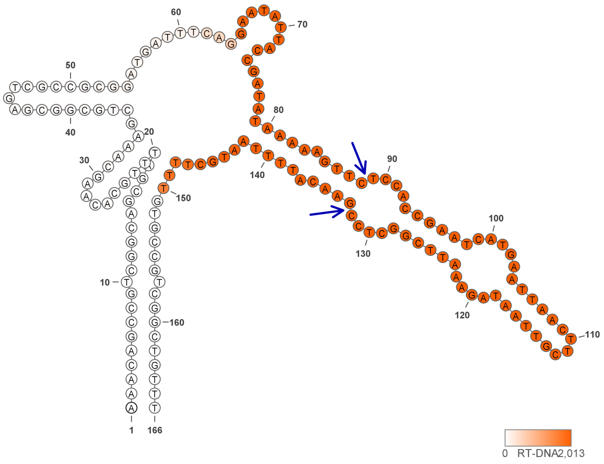


**482**

#### Retron Terminal 486: Firmicutes

**Bacterial Vector for RTDNA: pSLS.627**

**Bacterial Vector for Editing: None**

**Phage Vector for Editing: None**

**Human Vector for Editing: pSLS.851 (U6)**

**RTDNA Production (relative to Eco1): 0 by PAGE**

**Bacterial Editing: undetermined**

**Phage Editing: undetermined**

**Human Editing (demultiplexed): 2.586619 percent precise (22.28096063x Eco1)**

**ncRNA:**

**TTTATAAAGTAACTTTGCGATAAGCTAAATTTTTCTTCATTGCATTTCGTAAAATTCAGCTCATCGTT**
**AACTTACTTCACAAGTTTCGTAAGTTAAGTACTTCAAAACCACCAGCATAAAAGGTGGTTTTTTTAG**
**GAGGTAATATGAATCCAATATTGATAAGCGAACAGCAAATTAAGTTGCTTTTTAAA**

**no RTDNA sequencing data**

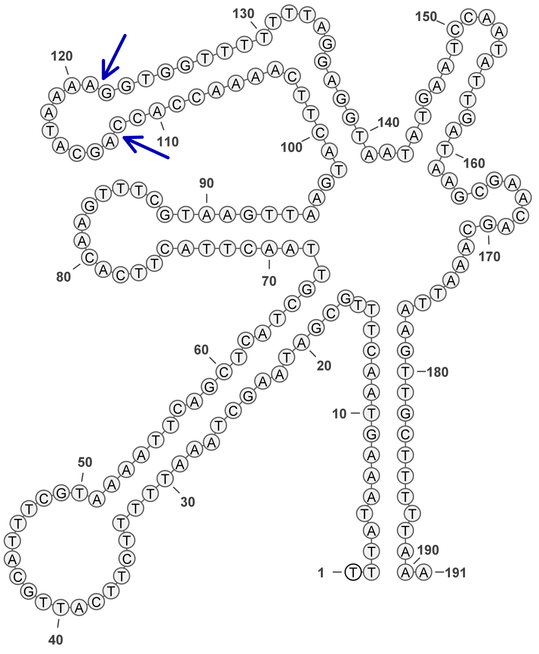


**486**

### Retron Terminal 488: Proteobacteria

**Bacterial Vector for RTDNA: pSLS.493**

**Bacterial Vector for Editing: pAGD150**

**Phage Vector for Editing: None**

**Human Vector for Editing: None**

**RTDNA Production (relative to Eco1): undetermined**

**Bacterial Editing: 0.65994758 percent precise (0.203074142x Eco1)**

**Phage Editing: undetermined**

**Human Editing (demultiplexed): undetermined**

**ncRNA:**

**TTTTTGTTGCATTATTCTTAGCGCACCAGCGATTCGCTTGGTATGCGGCAGTCAACAACATTAAATT**
**GTCAGTTTTTGCTTTTACAAAAAAGCGTCCACTATGGACCTATCAGCGAGGCCCCACCGTCGTCAA**
**CTACGATGGTGCCGTAGCCGACGACATTACGTACGGTCGAAAGATAGCAGAGG**

**RTDNA:**

**GTACGTAATGTCGTCGGCTACGGCACCATCGTAGTTGACGACGGTGGGGCCTCGCTGATAGGTCC**
**ATAGTGGACGCTTTTTTGTAAAAGCAAAAACTGACAATTTAATGTTGTTGACTGCCGCATACCAAG**
**CGAATCGCTGGTGCGCTAAGAATAATGCA**

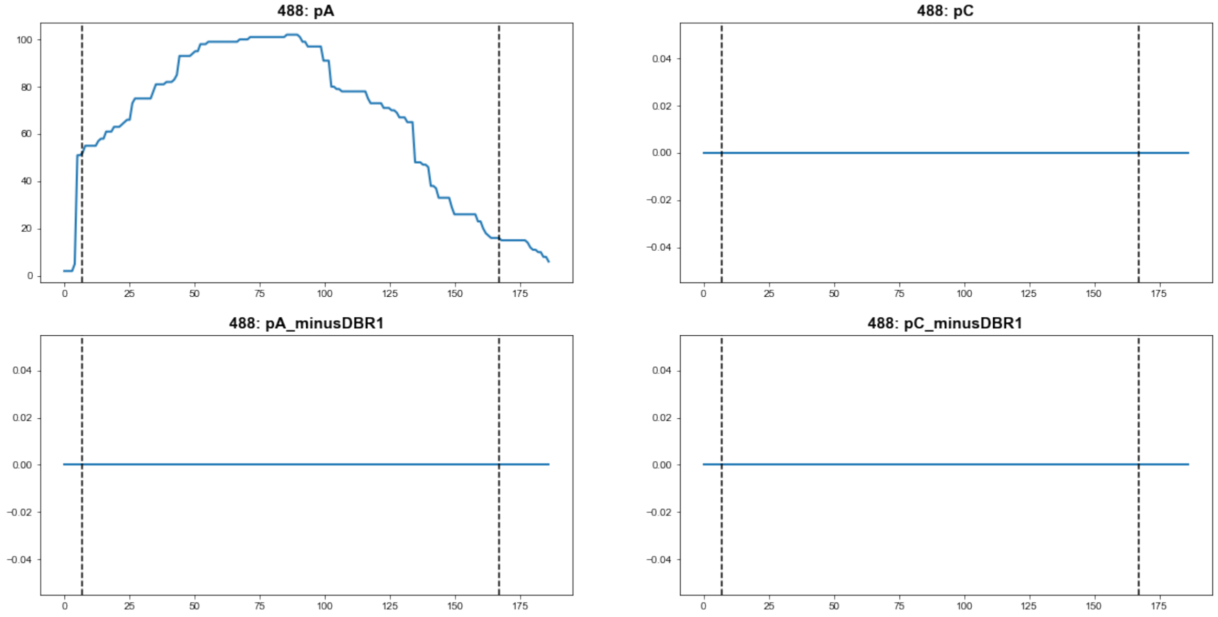


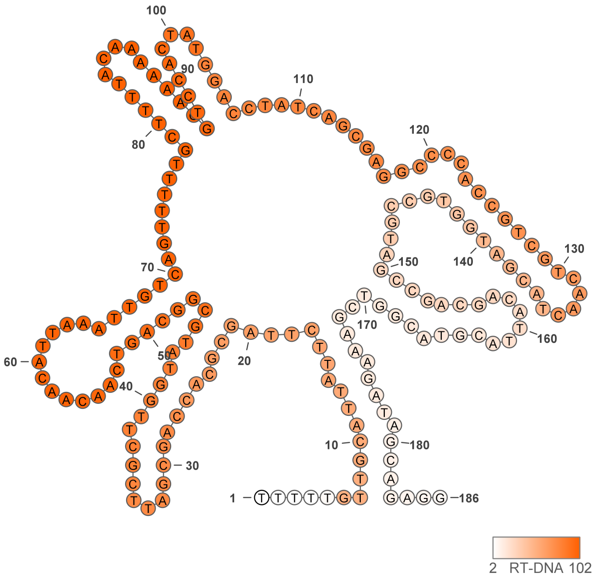


488

### Retron Terminal 506: Proteobacteria

**Bacterial Vector for RTDNA: None**

**Bacterial Vector for Editing: None**

**Phage Vector for Editing: None**

**Human Vector for Editing: pSLS.946**

**RTDNA Production (relative to Eco1): undetermined**

**Bacterial Editing: undetermined**

**Phage Editing: undetermined**

**Human Editing (demultiplexed): 0 percent precise (0x Eco1)**

**ncRNA:**

**TTTGTTTGATAGATTTACTTTCAGTGCAGCAAAGGCACCTATAACGAGTATCGATAAACTCTGTGA**

**GACTCTATCTGTGAGTCGCGAAGAGCTATACGACACACTCGATCTTCCAGACGAA**

**no RTDNA sequencing data**

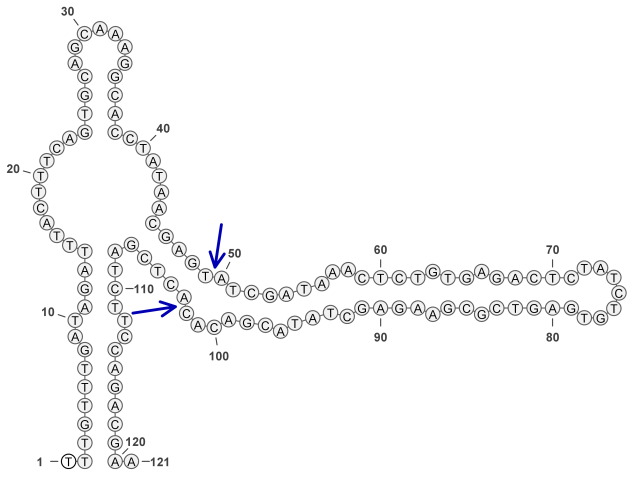


**506**

### Retron Terminal 530: Proteobacteria

**Bacterial Vector for RTDNA: pSLS.662**

**Bacterial Vector for Editing: None**

**Phage Vector for Editing: None**

**Human Vector for Editing: None**

**RTDNA Production (relative to Eco1): 0 by PAGE**

**Bacterial Editing: undetermined**

**Phage Editing: undetermined**

**Human Editing (demultiplexed): undetermined**

**ncRNA:**

**GGATGTTACTCAATTAATTTCAGCTCTTATTGGCTAGTTTTAACTAGTGCTATTGAATAGATAGCTT GTGGGGAGAAGTGTCGTTTCCGTCATTTGTCCCAAATAATCCTGATAGCCGAGGGTAGTAATCTCA GTGTTTATTTAAGTGCTGTTGAGTAACCAAGTAAATTGAGATTACAAGCTCTATGTATTGTTCTTTA GAGTTTATTATATCT**

**RTDNA: None**

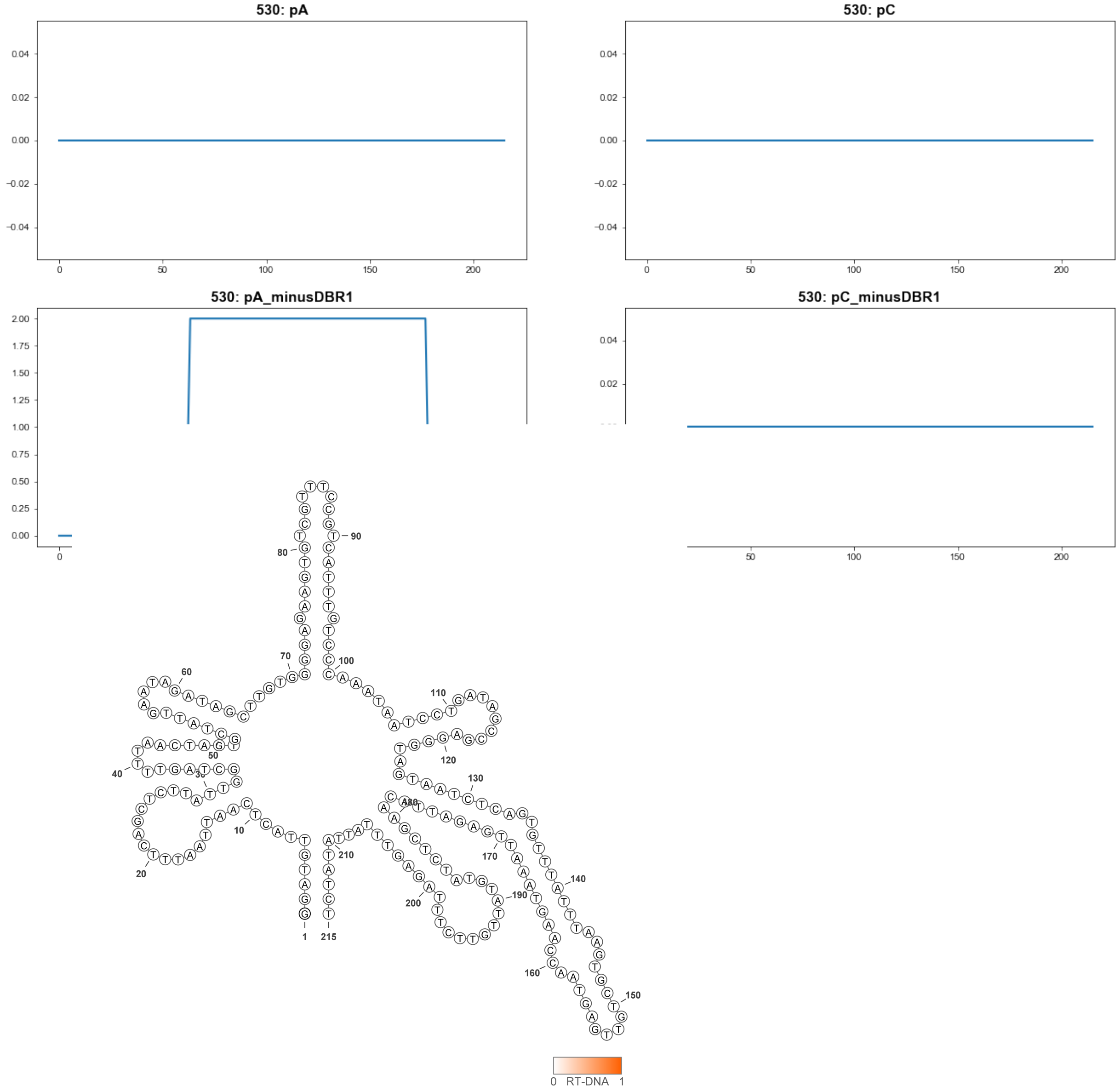


**530**

### Retron Terminal 544: Proteobacteria

**Bacterial Vector for RTDNA: pSLS.622**

**Bacterial Vector for Editing: None**

**Phage Vector for Editing: None**

**Human Vector for Editing: None**

**RTDNA Production (relative to Eco1): 0 by PAGE**

**Bacterial Editing: undetermined**

**Phage Editing: undetermined**

**Human Editing (demultiplexed): undetermined**

**ncRNA:**

**AATGAAGAACTTACTCTCTCTGTGCGTATGCTCTAATAAAAGCTCACCATTTGGTCTGGCACCCAA
TTCCGGGTAACTAAATGTGGAGGTGCAATATGCACATTAACTTTGAAATAAACTTAGCTATTTGTTG
TAGTAAAAAAAGAGATGACTATTCCACTAGAGATAGTGAACTAGTTCATAGACGTAAAGGAGAGA
GGGTTAAAGCTAGGAATAGATTGCCTATTGGGTATATCTTACTAGGTTTGACCGTTCTAGAAGGAG
TGTTAAAGCTTCTGACAGGATAAAATAAGTTCTTCATT**

**RTDNA: None**

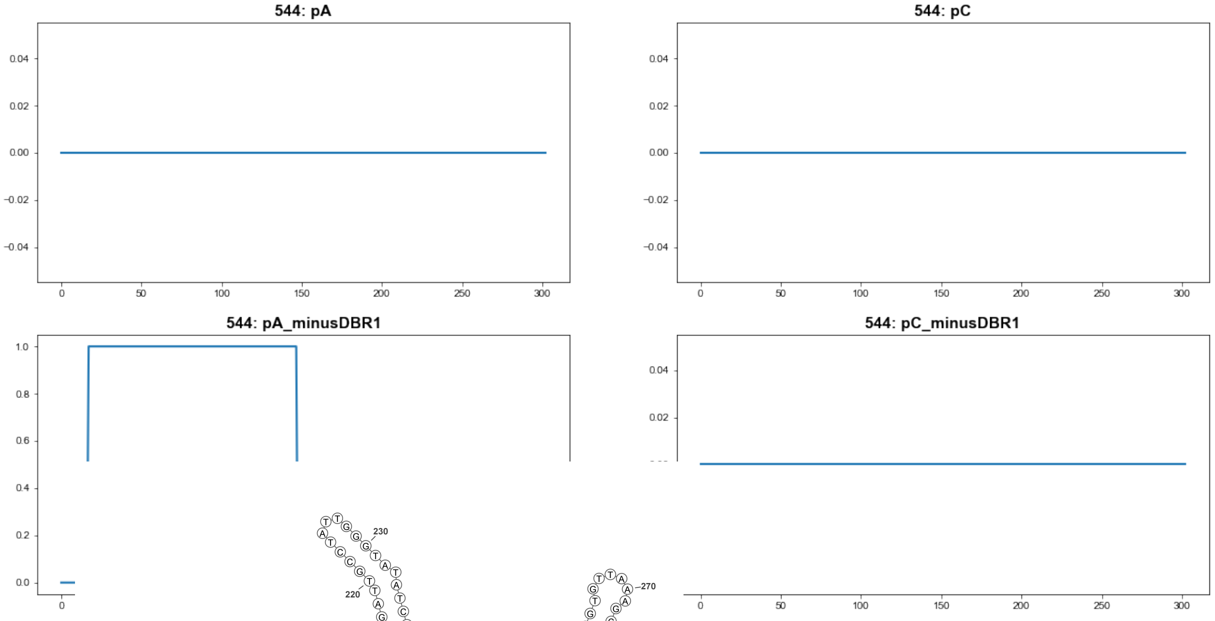


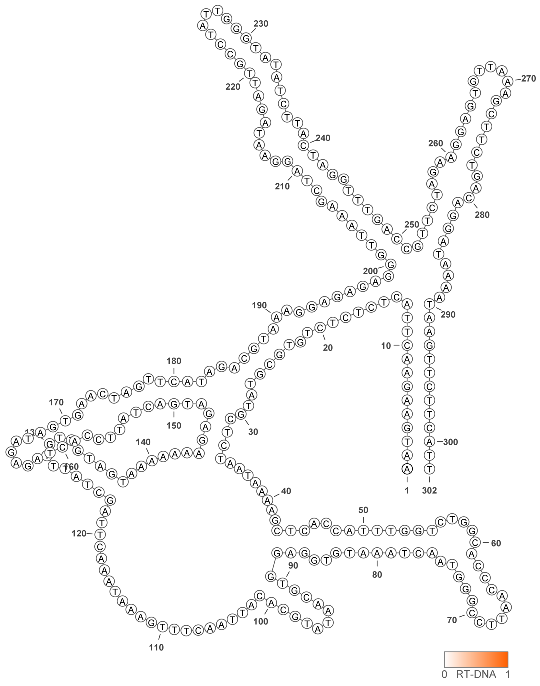


**544**

### Retron Terminal 553: Proteobacteria

**Bacterial Vector for RTDNA: None**

**Bacterial Vector for Editing: None**

**Phage Vector for Editing: None**

**Human Vector for Editing: pSLS.947**

**RTDNA Production (relative to Eco1): undetermined**

**Bacterial Editing: undetermined**

**Phage Editing: undetermined**

**Human Editing (demultiplexed): 0.198174 percent precise (1.707057395x Eco1)**

**ncRNA:**

**TGGTTACGTGTATGTTGCGCAAGGAAACCGATGCCTTGTTTTTAAGGCGATCTCGCATGAGATCAT**
**GACCGGCCTGGCCGCTAATTAATGGTGAGAACGCCCCTGAGCGTGTATCAGCACTAAAAGAATAG**
**CGACGATTGTCTGGCAATGTGAGTTAAAAGTCCCAAAGAGCCAATAGAGTTTTTCCTTGCTTTCCT**
**ACTGGTCTGGCAAAATTTCATAACCAATAGGAGTTGCATCATGTTTATTATCAACATCAACATTAAC**
**GTAACCA**

**no RTDNA sequencing data**

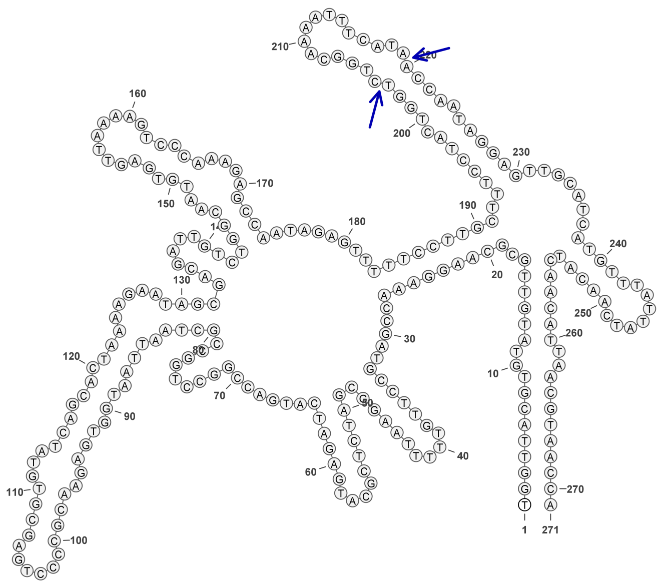

**553**

### Retron Terminal 604: Proteobacteria

**Bacterial Vector for RTDNA: pSLS.590**

**Bacterial Vector for Editing: None**

**Phage Vector for Editing: None**

**Human Vector for Editing: pSLS.825**

**RTDNA Production (relative to Eco1): 0 by PAGE**

**Bacterial Editing: undetermined**

**Phage Editing: undetermined**

**Human Editing (demultiplexed): 0.299257 percent precise (2.577779501x Eco1)**

**ncRNA:**

**TGGACACCTTCAAGTTCCTTTCGGCCTAGAAGCTTCGCGGCGGGGCAATTCCGCCTTGAGGAGCT**

**GTATCGATGAAATATTTGAAGAAAACTTGTCGCCCTAAAGCGGCAAATGATGTGCACAACAATGA**

**AGGTGTCCG**

**RTDNA: None**

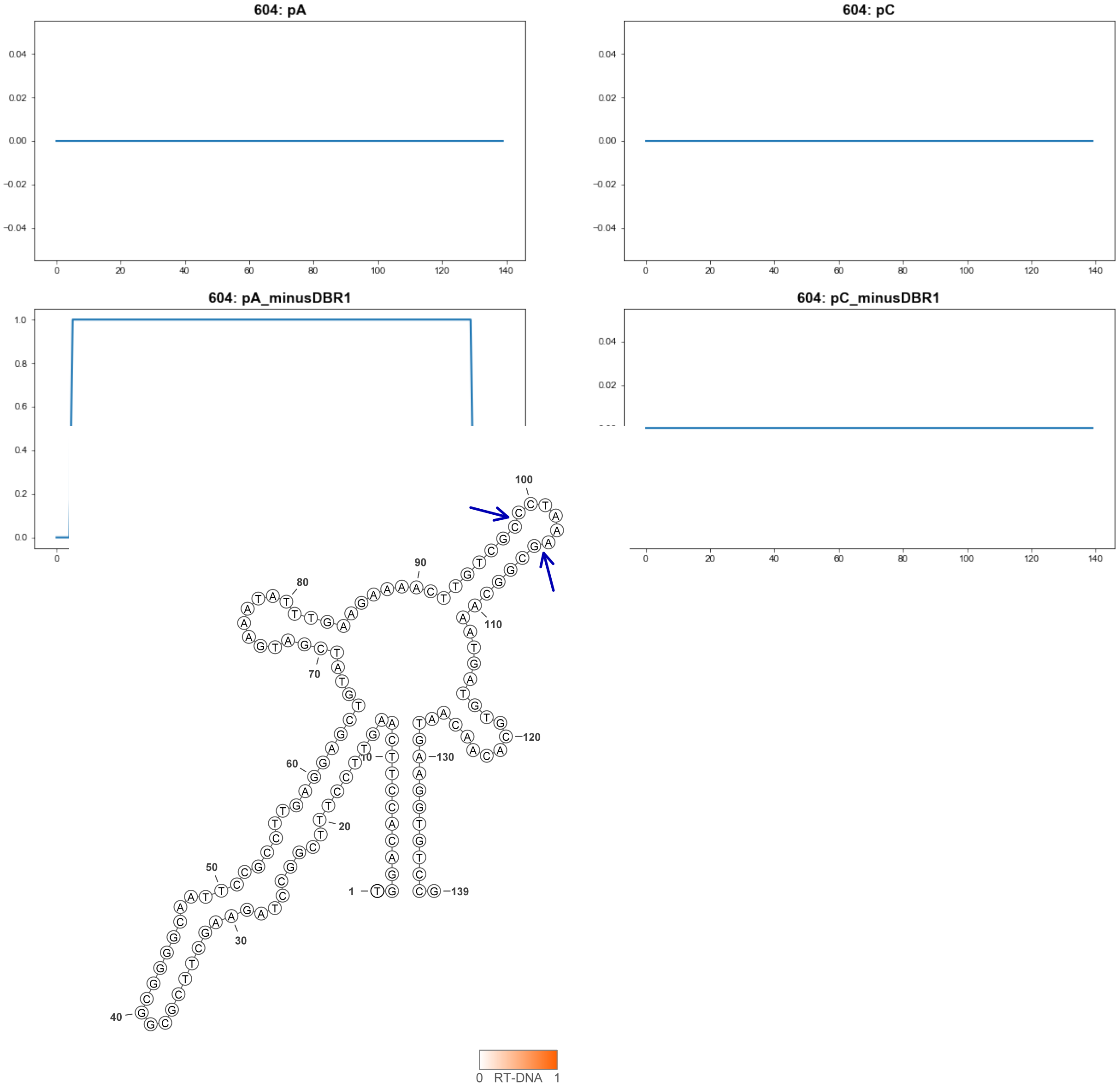


604

### Retron Terminal 613: Bacteroidetes

**Bacterial Vector for RTDNA: pSLS.591**

**Bacterial Vector for Editing: None**

**Phage Vector for Editing: None**

**Human Vector for Editing: pSLS.826**

**RTDNA Production (relative to Eco1): 0 by PAGE**

**Bacterial Editing: undetermined**

**Phage Editing: undetermined**

**Human Editing (demultiplexed): 0.197908 percent precise (1.704766089x Eco1)**

**ncRNA:**

**TGATTGTAACACAGGAGAAGAAGATAAAAAATTTGGCAAAATGGATTAAAGCCATTGCTGAATTTA TAAAGGCATTGATTCAATTATCAAATGCAATGAAGCCTTTGATAGAATGGTGCAAATCTTTTTTTGA ATTGTTATAATCA**

**RTDNA: None**

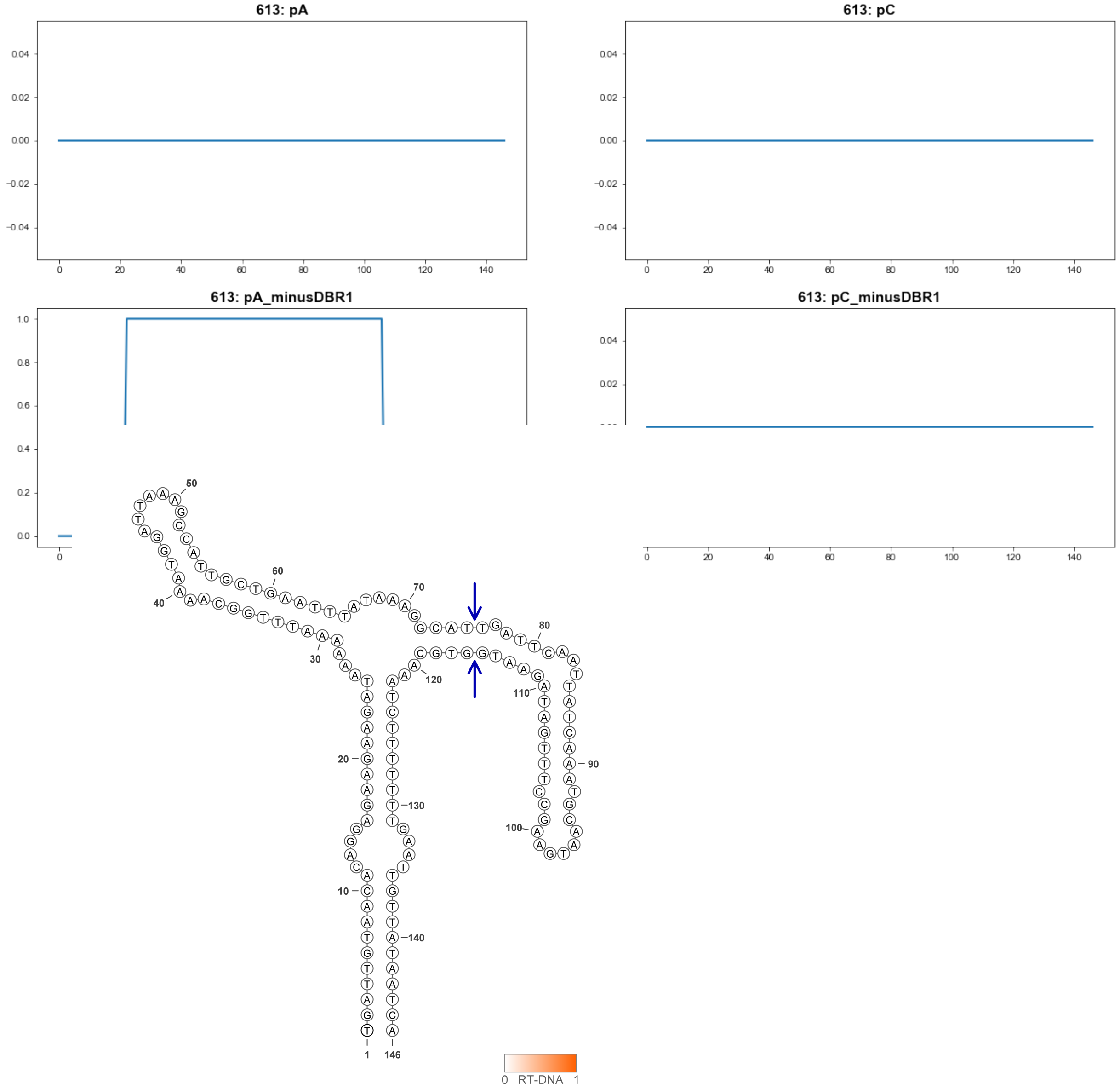


613

### Retron Terminal 662: Proteobacteria

**Bacterial Vector for RTDNA: pSLS.665**

**Bacterial Vector for Editing: None**

**Phage Vector for Editing: None**

**Human Vector for Editing: None**

**RTDNA Production (relative to Eco1): 0 by PAGE**

**Bacterial Editing: undetermined**

**Phage Editing: undetermined**

**Human Editing (demultiplexed): undetermined**

**ncRNA:**

**TTGTGCTGGGGTACGTTGGTACCTTGTGGGGCTGAAAGGCTCTAATTACTCGCTCGCTCTCCTCGG**

**TGGGGTGCCTTGCCTCGGCGCCCGGCCATGAGAATCGGCGCGA**

**RTDNA: None**

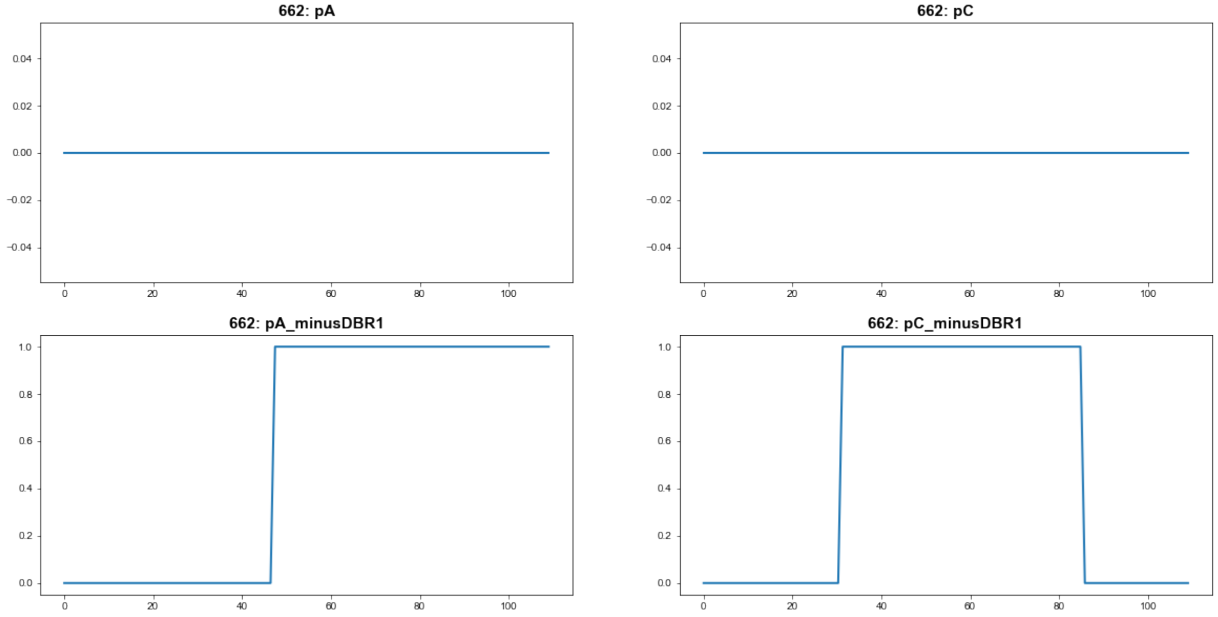


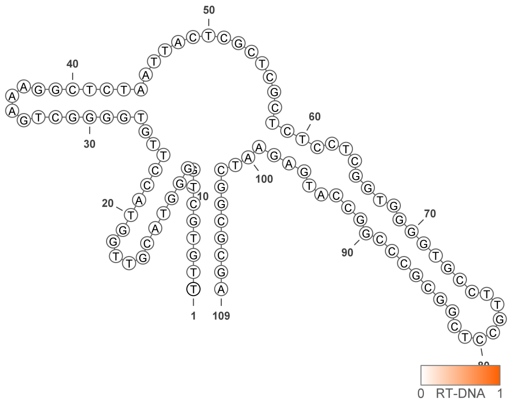


**662**

### Retron Terminal 685: Proteobacteria

**Bacterial Vector for RTDNA: pSLS.666**

**Bacterial Vector for Editing: None**

**Phage Vector for Editing: None**

**Human Vector for Editing: None**

**RTDNA Production (relative to Eco1): 0 by PAGE**

**Bacterial Editing: undetermined**

**Phage Editing: undetermined**

**Human Editing (demultiplexed): undetermined**

**ncRNA:**

**CCTCGTCACTTATTGAGCTTATTTAGCTCGTGGGAAAACGAAAGTTTTCTGCCTTTTTGACCCATAC ATGCCTAGCAGTGACCATGATGAGCACCGTTCAGAACATGTGGGAATTCGAACCTTGGGACGAAA CCAAGGTTCGTCGAGACGGGG**

**RTDNA:**

**TTCGTCCCAAGGTTCGAATTCCCACATGTTCTGAACGGTGCTCATCATGGTCACTGCTAGGCATGT ATGGGTCAAAAAGGCAGAAAACTTTCGTTTTCCCACGAGCTAAATAAGCTCAATAAGTGAC**

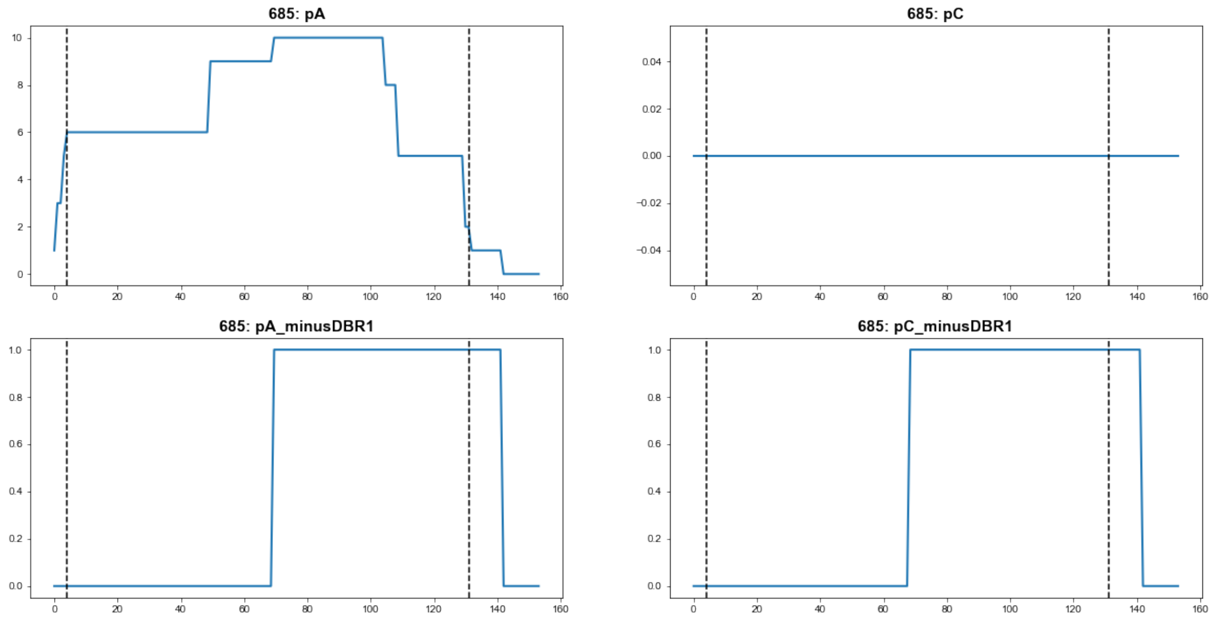


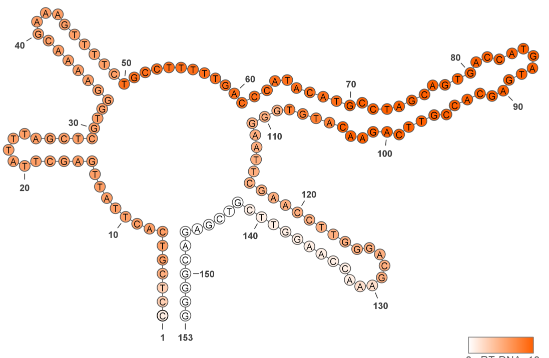


**685**

### Retron Terminal 703: Proteobacteria

**Bacterial Vector for RTDNA: pSLS.628**

**Bacterial Vector for Editing: None**

**Phage Vector for Editing: None**

**Human Vector for Editing: pSLS.854**

**RTDNA Production (relative to Eco1): 0 by PAGE**

**Bacterial Editing: undetermined**

**Phage Editing: undetermined**

**Human Editing (demultiplexed): 0.010568 percent precise (0.091032035x Eco1)**

**ncRNA:**

**TTAATATCTCTAACTCTTATCTTTCTGAAATTGAAAGTGGCAAAAAGCCTCCAAGTATTGAGCTCCT ACAAAGCTATGCGACAGTATTTGATATACCTGTCTCTTCTCTTTTGTATTTTTCTGAACAGTTAGAG AACGAAGGCAAAATCTCTAAAAGTTTTAGGATTAAATCGGCAGGATTTGTTTTAAAGCTTTTAGAC TGGAGTATTAA**

**RTDNA: None**

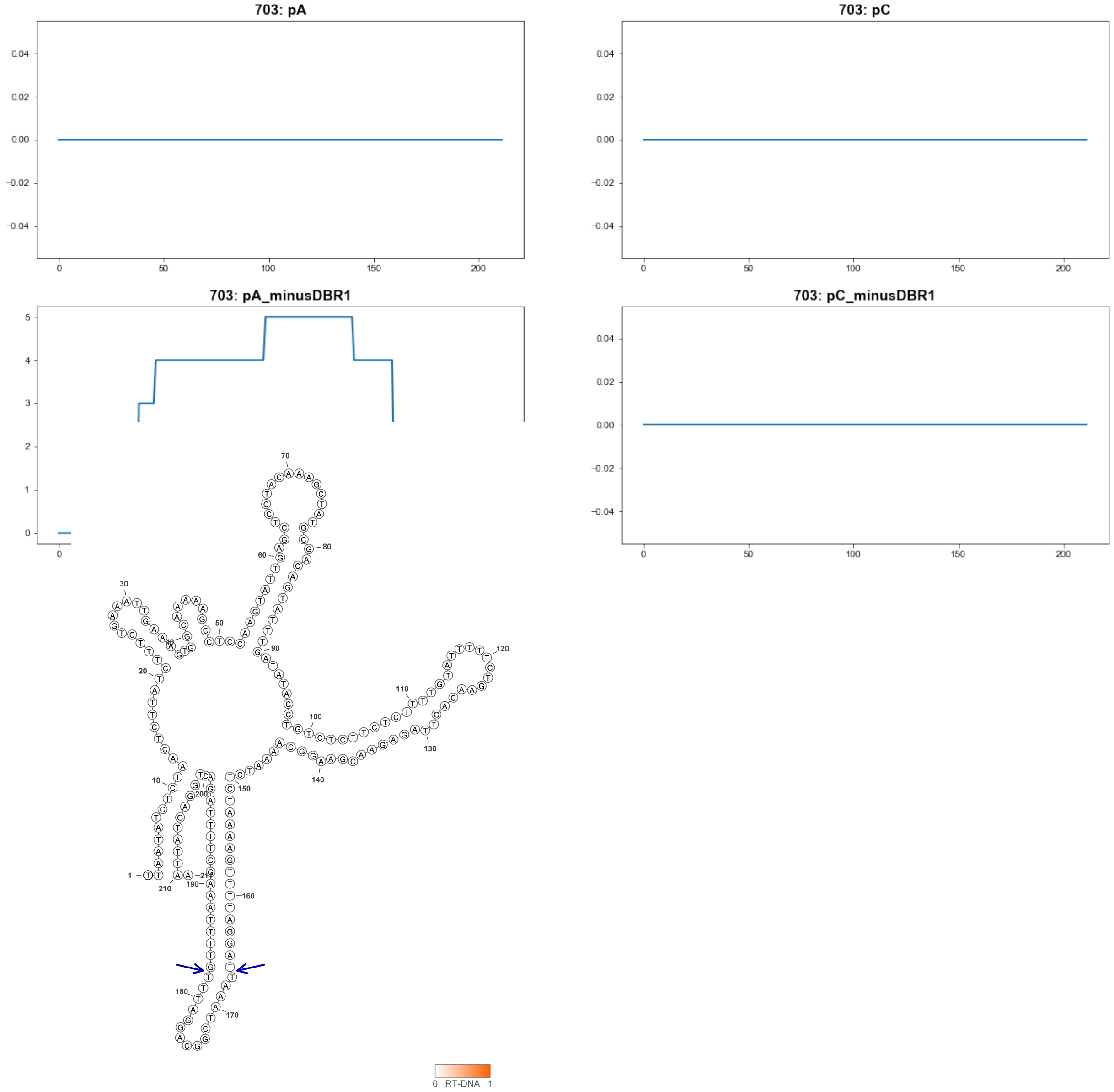


**703**

### Retron Terminal 729: Candidatus Levybacteria

**Bacterial Vector for RTDNA: pSLS.668**

**Bacterial Vector for Editing: None**

**Phage Vector for Editing: None**

**Human Vector for Editing: pSLS.904**

**RTDNA Production (relative to Eco1): 0 by PAGE**

**Bacterial Editing: undetermined**

**Phage Editing: undetermined**

**Human Editing (demultiplexed): 0.014489 percent precise (0.124807263x Eco1)**

**ncRNA:**

**TTAATTCTTTACTCAATCATTTACTGATTGACTTAAGACGTTCTTTTTTGAATGTCGCAAGACGTATT**

**AAAACTACGTTTTGCATTTTCCGGCGCCGGTATTCTTATATCGATAGTTGCAAGTAAGGTAACAAC**

**ACTACTTCTTCAAAAGAAATAACTATACTTCGGCACTAGCAAAAAGAATTAA**

**RTDNA: None**

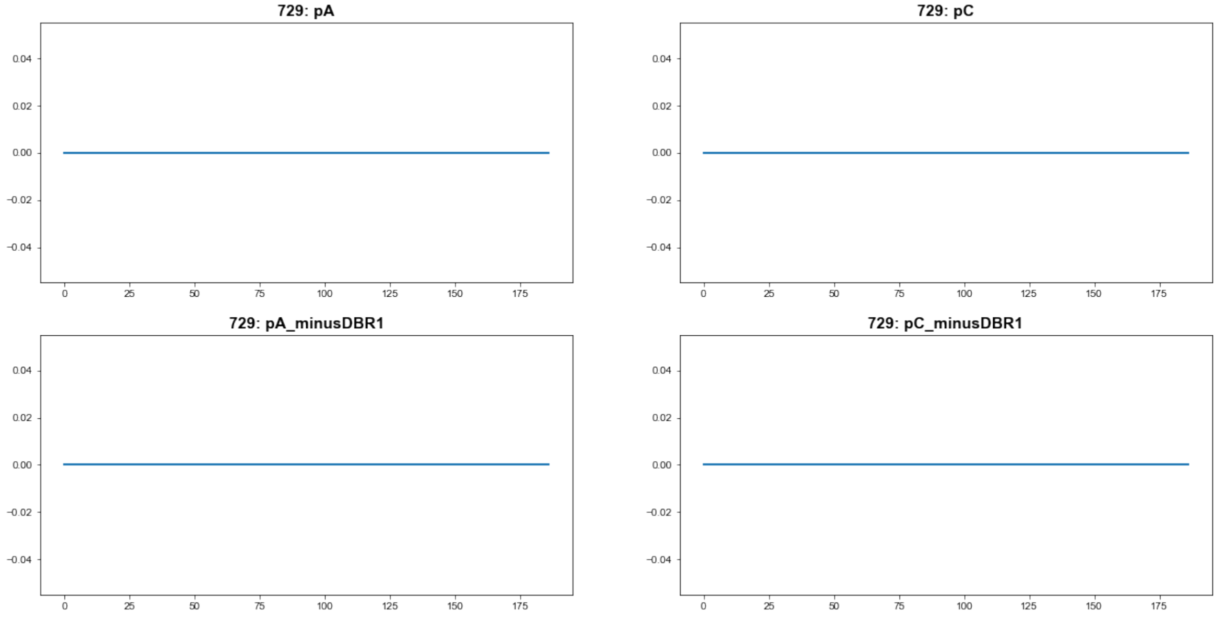


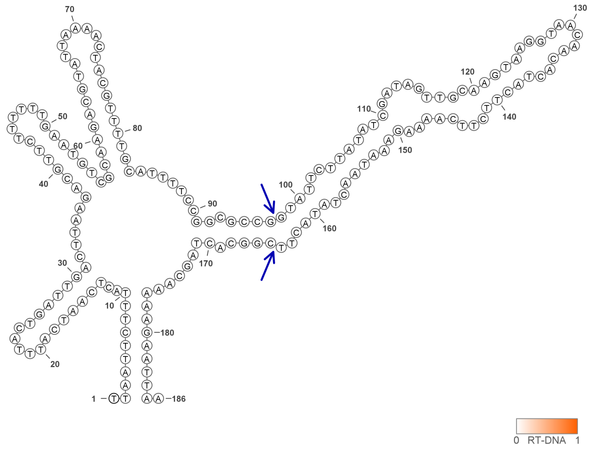


729

### Retron Terminal 733: Proteobacteria

**Bacterial Vector for RTDNA: None**

**Bacterial Vector for Editing: None**

**Phage Vector for Editing: None**

**Human Vector for Editing: pSLS.992**

**RTDNA Production (relative to Eco1): undetermined**

**Bacterial Editing: undetermined**

**Phage Editing: undetermined**

**Human Editing (demultiplexed): 0.031202 percent precise (0.268771912x Eco1)**

**ncRNA:**

**GATTTCTTCATCGATAGAGTAAAGGGTTTTGCTTGCAGAACGGAGCCTTGAGCTTCCTAGCCTATC**
**ATTTACTTGTTGCTGCTACAGTTCGTATCGCACTTTTAAGTACACGTAAACGTGAAAAGATGTTCTT**
**TGTATCGTGAAAGGAATC**

**no RTDNA sequencing data**

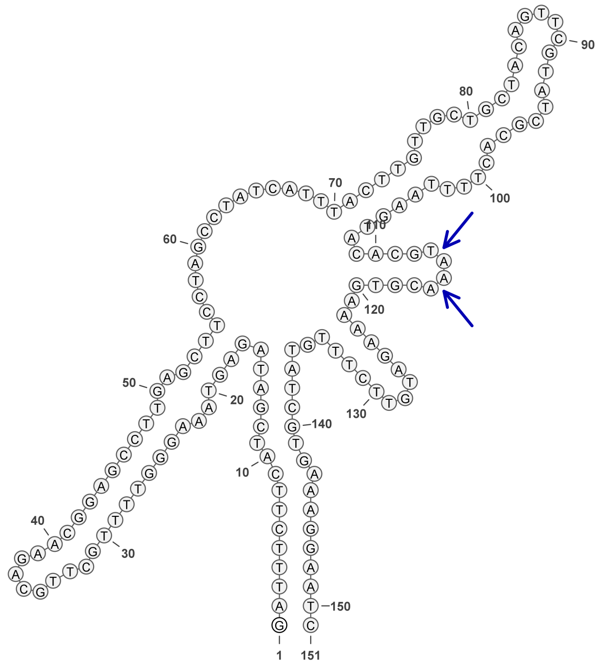

733

### Retron Terminal 735: Proteobacteria

**Bacterial Vector for RTDNA: None**

**Bacterial Vector for Editing: None**

**Phage Vector for Editing: None**

**Human Vector for Editing: pSLS.955**

**RTDNA Production (relative to Eco1): undetermined**

**Bacterial Editing: undetermined**

**Phage Editing: undetermined**

**Human Editing (demultiplexed): 0.007298 percent precise (0.062864477x Eco1)**

**ncRNA:**

**TTGGGTTTCTAGTTTTCTAGATTTCTCTATTAATGAACTAAGAAAGAGCGCTAAGAATGAAGATGAA TTAGTGCAAAAATTTAAGGGTACATTCCCTGAGTTATATAGTACAATAGAAGAGATTTCATCTTCGA TAGAATAATGGATCTCGCTTGCGAGATGGAGCCTGGAGCTCCCTAGCCTAA**

**no RTDNA sequencing data**

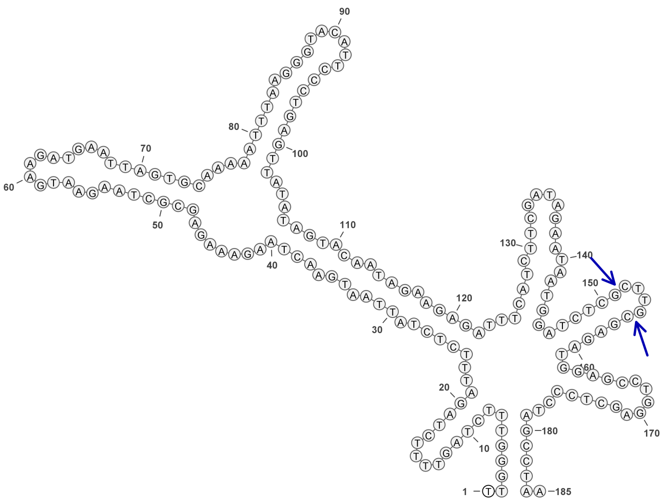

**735**

#### Retron Terminal 749: Proteobacteria

**Bacterial Vector for RTDNA: pSLS.580**

**Bacterial Vector for Editing: pAGD284**

**Phage Vector for Editing: pAGD396**

**Human Vector for Editing: pSLS.813 (H1)**

**RTDNA Production (relative to Eco1): 0.1855 by PAGE**

**Bacterial Editing: 1.63828543 percent precise (0.50412096x Eco1)**

**Phage Editing: 45.45809605 percent precise (1.121209038x Eco1)**

**Human Editing (demultiplexed): 0.004257 percent precise (0.036669509x Eco1)**

**ncRNA:**

**GCTTCTTCTTCGATAGAAGCTGGAGGGCTCAAATGAGCTGACGCATGCCGTCACTTGCTCCAGCA AACACTAGCTCGGATTGTCTTGCGTCCGCTAAGCCGCTTTAAGCCTCACGGGGTGTAAAGATGTTA TTGGTATCGAAGAGGAAGC**

**RTDNA:**

**CAATAACATCTTTACACCCCGTGAGGCTTAAAGCGGCTTAGCGGACGCAAGACAATCCGAGCTAG TGTTT**

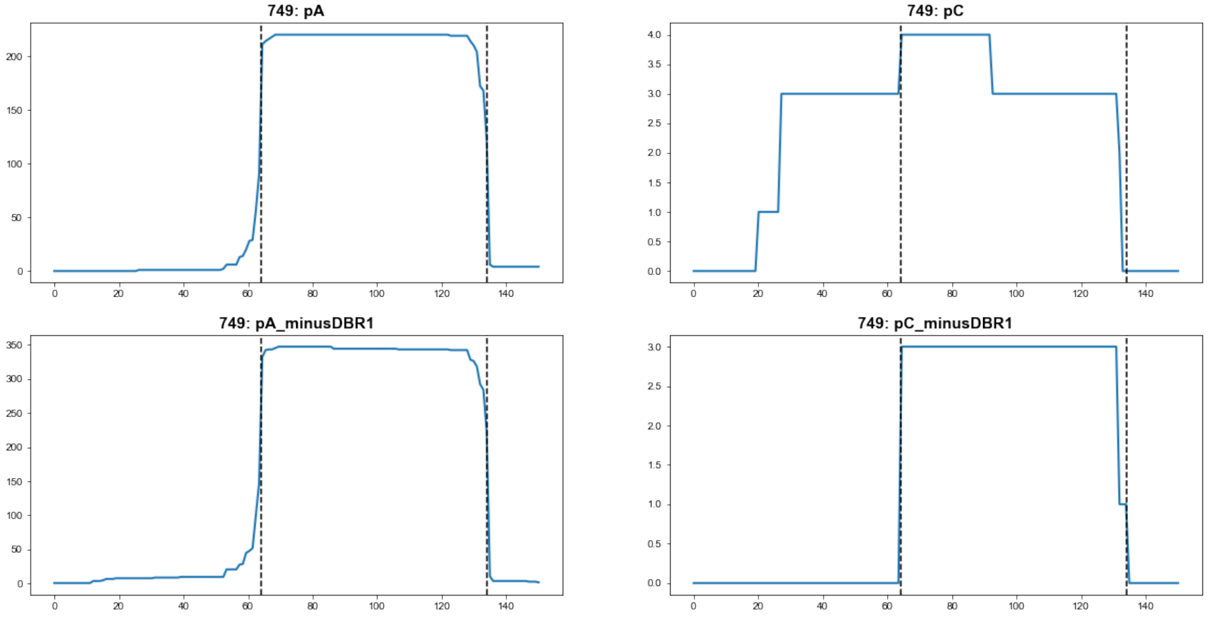


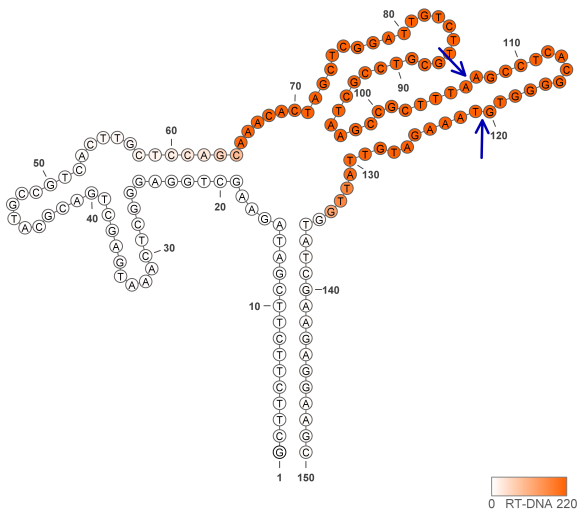


**749**

### Retron Terminal 763: Proteobacteria

**Bacterial Vector for RTDNA: None**

**Bacterial Vector for Editing: None**

**Phage Vector for Editing: None**

**Human Vector for Editing: pSLS.929**

**RTDNA Production (relative to Eco1): undetermined**

**Bacterial Editing: undetermined**

**Phage Editing: undetermined**

**Human Editing (demultiplexed): 0.131415 percent precise (1.131999897x Eco1)**

**ncRNA:**

**ATGTTTACATTCGGTAGTAATAGAGATTGCCTTGTGCAATGTAGGGAAACCTACCATCTTTCTTTGC GACAGTGTAGATTTTATATCATTAGTAAAATTTTATGCTTAAGCTTGGTTAGAACAGTCTCCACATA CCAGTGTGATTACTTTGATATTCGCGCATTATGTGTCACACGTGAATGTTTGTTTTTGTACCGGATT AAACAT**

**no RTDNA sequencing data**

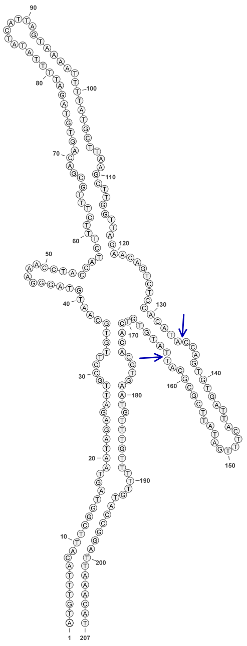

763

### Retron Terminal 781: Firmicutes

**Bacterial Vector for RTDNA: pSLS.669**

**Bacterial Vector for Editing: None**

**Phage Vector for Editing: None**

**Human Vector for Editing: pSLS.905**

**RTDNA Production (relative to Eco1): 0 by PAGE**

**Bacterial Editing: undetermined**

**Phage Editing: undetermined**

**Human Editing (demultiplexed): 0 percent precise (0x Eco1)**

**ncRNA:**

**AGTAAATTGATGAGTAGTAACATGTATCAATTATCATGATTGGTTGATATAACCTTGATGCCCTTAC**
**TAGAATTTAAGCAAAAAACTTGAATACTGAATAGTAAAAAGCCAAGGAGAATTTATATACAATTTG**
**CGATGTTTTATCTGCTCGTAAAGTTATCTAATATTTTTCATAGTTGTATACTCGCAGTGCGTTAATGT**
**GAGAAGTACTCATCAAAAGACT**

**RTDNA: None**

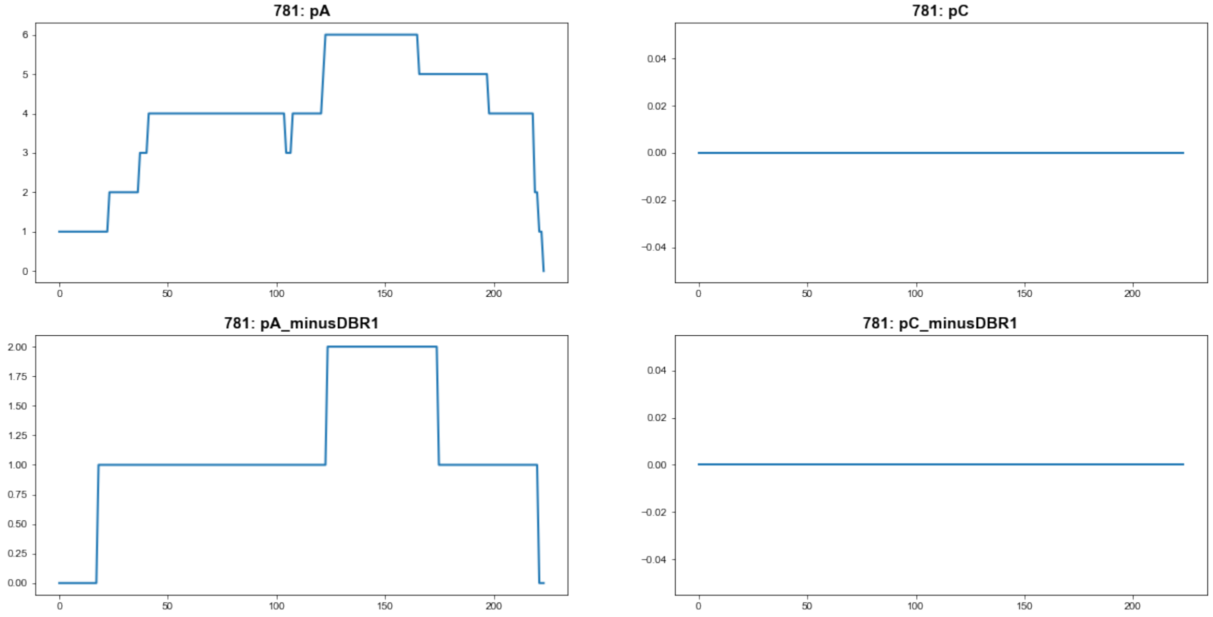


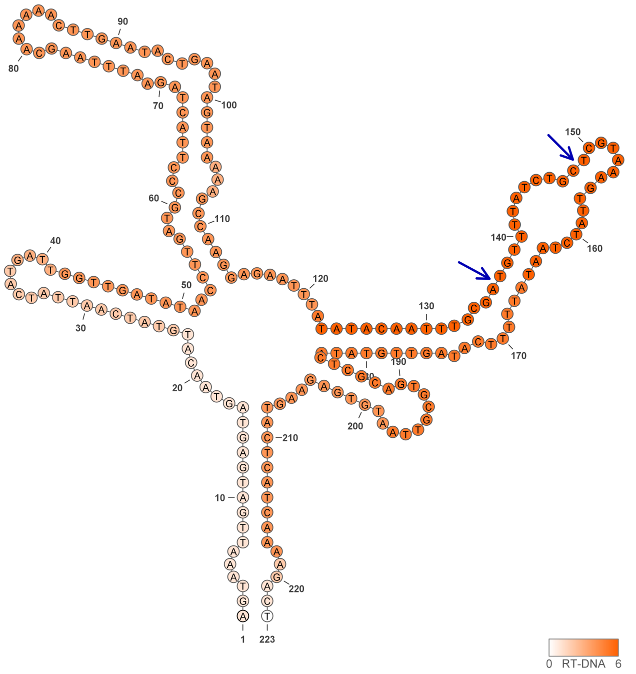

### Retron Terminal 783: Proteobacteria

**Bacterial Vector for RTDNA: pSLS.581**

**Bacterial Vector for Editing: None**

**Phage Vector for Editing: None**

**Human Vector for Editing: None**

**RTDNA Production (relative to Eco1): 0 by PAGE**

**Bacterial Editing: undetermined**

**Phage Editing: undetermined**

**Human Editing (demultiplexed): undetermined**

**ncRNA:**

**GAGAAGCTGATCAGCCCATGGTGAAGTTCAGGGCTACTTATGCTAGTCGACATGGGATTGTTCCA
AACAATGACAACCTGTGCTATGCCCAATAACGGAGCAGCTCGACGACAAGGTTAGAGGTGCGAG
TCGCATCTGCCTATTACGGAACCACTTGCTACGTGTGTCTTGCTGCTCCGCCTAAGAGATTTAAAA
TGGCATTTGATTACGATTTACGACGCATAAAGACAAAAGGTCAGCTTCTC**

**RTDNA: None**

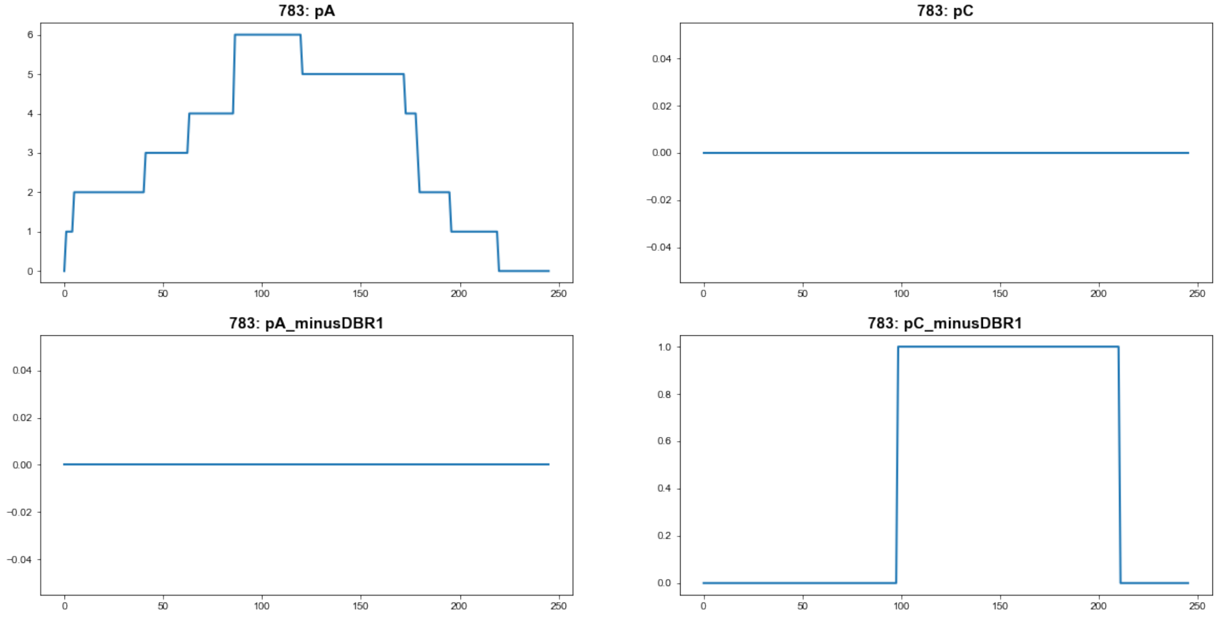


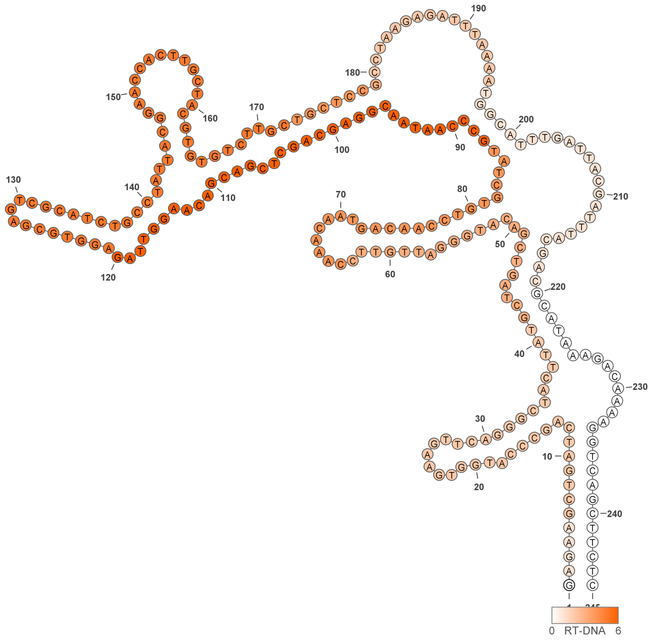


**783**

### Retron Terminal 789: Proteobacteria

**Bacterial Vector for RTDNA: pSLS.567**

**Bacterial Vector for Editing: None**

**Phage Vector for Editing: None**

**Human Vector for Editing: None**

**RTDNA Production (relative to Eco1): 0 by PAGE**

**Bacterial Editing: undetermined**

**Phage Editing: undetermined**

**Human Editing (demultiplexed): undetermined**

**ncRNA:**

**AGGGGTGGGCGTCCCTGCCTGTGGTTCAGGGCGCGAGTCGCGTCCGCCATCGCGGTATCACTTG CTTCTCCAGCACGGCGCCCACCCCT**

**RTDNA: None**

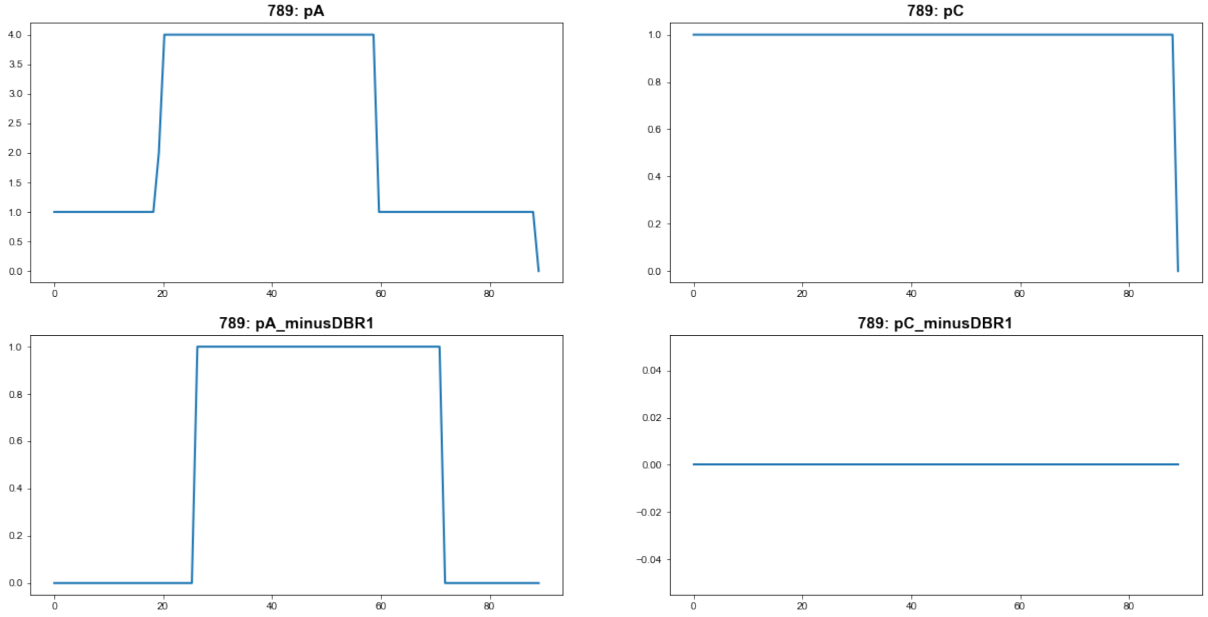


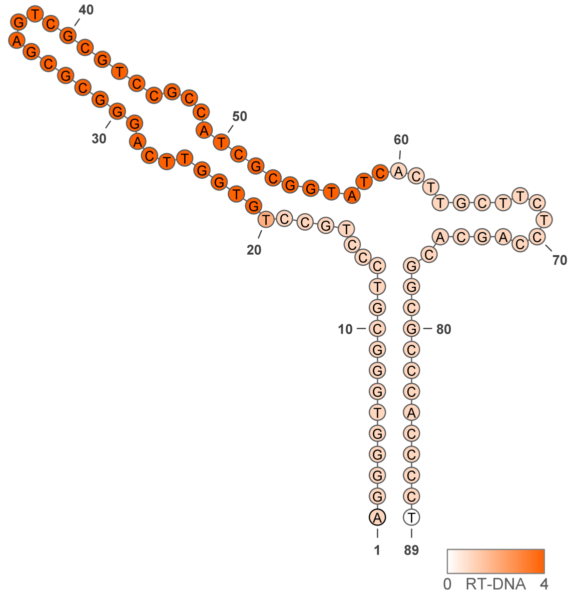

### Retron Terminal 796: Proteobacteria

**Bacterial Vector for RTDNA: pSLS.670**

**Bacterial Vector for Editing: None**

**Phage Vector for Editing: None**

**Human Vector for Editing: pSLS.906A**

**RTDNA Production (relative to Eco1): 0 by PAGE**

**Bacterial Editing: undetermined**

**Phage Editing: undetermined**

**Human Editing (demultiplexed): 0.072866 percent precise (0.627662782x Eco1)**

**ncRNA:**

**GGTGGTTGGATGACAGACTTCGGATTAGTCCGTGGGTACGAGTCCGTACTCGACAATTAAGATCT CACATATTCCATCCAGCCACCTC**

**RTDNA:**

**TGGAATATGTGAGATCTTAATTGTCGAGTACGGACTCGTACCCACGGACTAATCCGAAGTCTGTC**

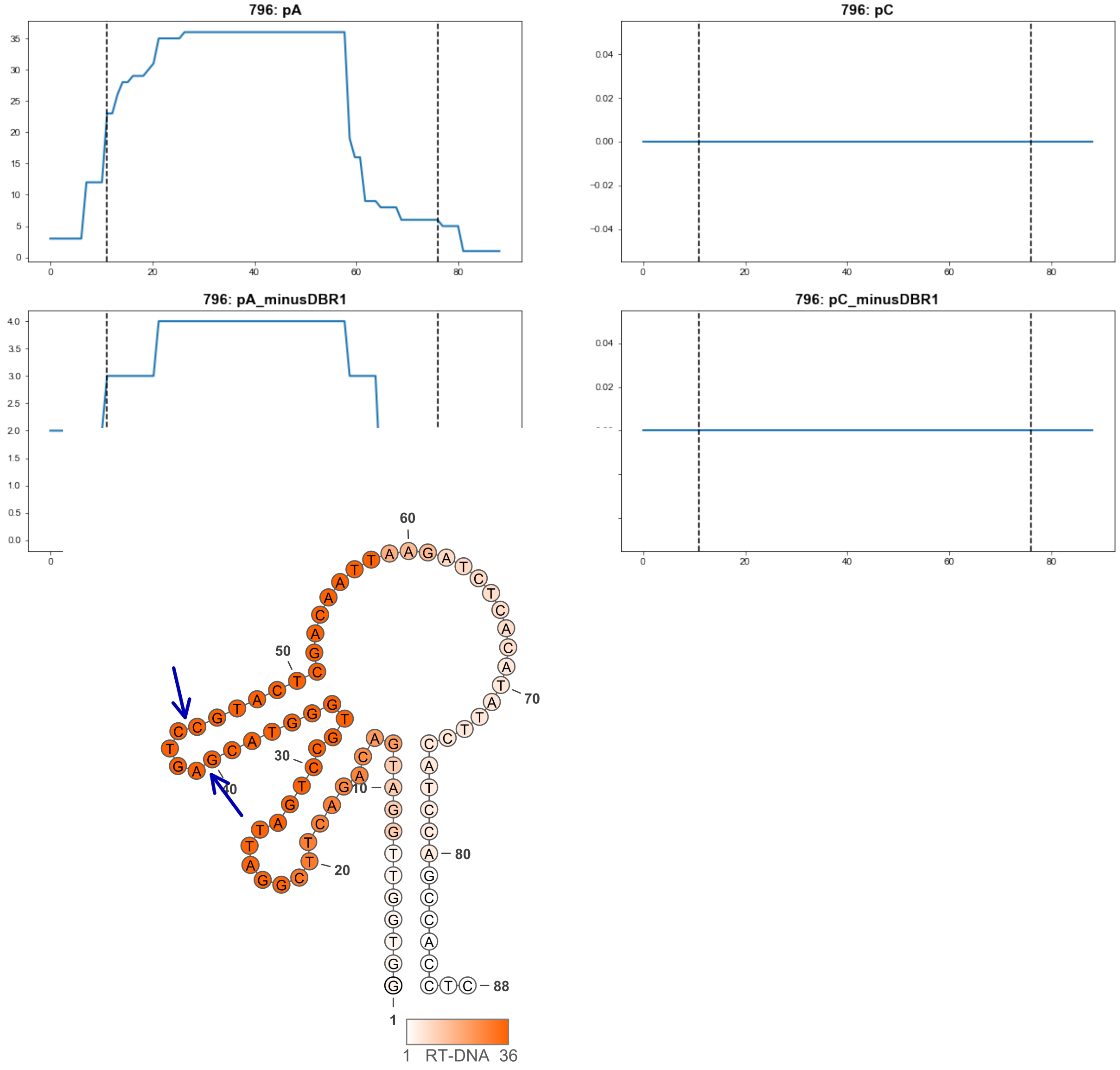


796

### Retron Terminal 803: Bacteroidetes

**Bacterial Vector for RTDNA: pSLS.640**

**Bacterial Vector for Editing: None**

**Phage Vector for Editing: None**

**Human Vector for Editing: pSLS.957 (U6)**

**RTDNA Production (relative to Eco1): 0.15625235 by PAGE**

**Bacterial Editing: undetermined**

**Phage Editing: undetermined**

**Human Editing (demultiplexed): 0 percent precise (0x Eco1)**

**ncRNA:**

**TGTAAGTTTCATATTGAACAAAATTTTATCATCTGAAGGAAATATTGAGAATTGGGCCAAAAGTGC
CTTTGATGTAATTAAAGACAAAATAGCCCTAATTCATTTTCAGAATGATGATTCATTTTTGAGTAGA
TTTGCTAATGAGGAAAACACGATAAACTTTAAGAATTTAAGAAAAAAACTTGCGGTGGTAAAGGG
ATAAATTTACGGATATGAGTTCGTATTCGGTATTAGAATAACAGTTAGTTCTTACCCTTGCCGCCGC
TTTTATTATATGAAACCTACA**

**RTDNA:**

**ATATCCGTAAATTTATCCCTTTACCACCGCAAGTTTTTTTCTTAAATTCTTAAAGTTTATCGTGTTTTC
CTCATTAGCAAATCTACTCAAAAATGAATCATCATTCTGAAAATGAATTAGGGCTATTTTGTCTTTA
ATTACATCAAAGGCACTTTTGGCCCAATTCTCAATATTTCCTTCAG**

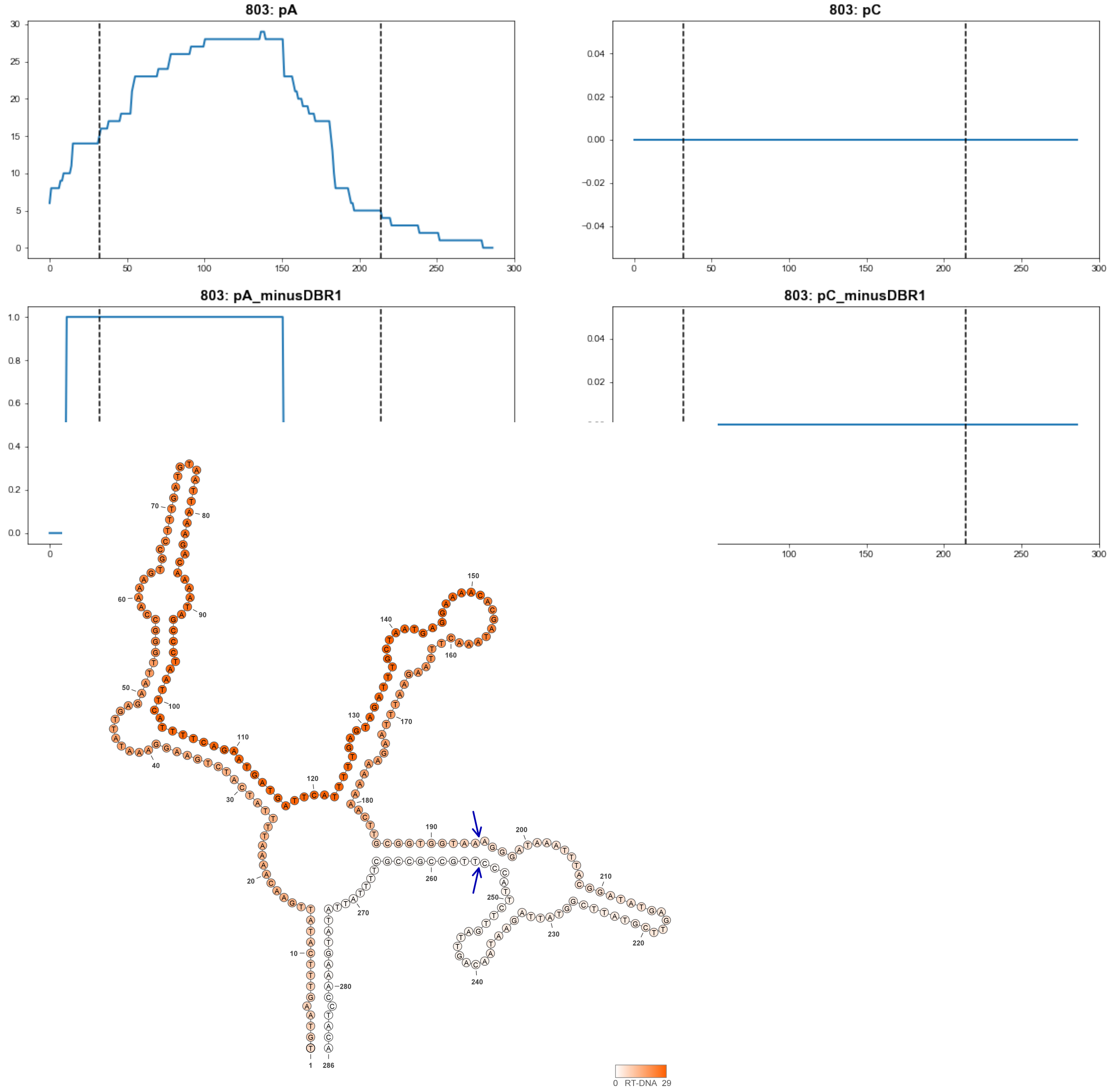


**803**

### Retron Terminal 807: Proteobacteria

**Bacterial Vector for RTDNA: pSLS.582**

**Bacterial Vector for Editing: None**

**Phage Vector for Editing: None**

**Human Vector for Editing: None**

**RTDNA Production (relative to Eco1): 0 by PAGE**

**Bacterial Editing: undetermined**

**Phage Editing: undetermined**

**Human Editing (demultiplexed): undetermined**

**ncRNA:**

**AGAGGTCAATTTTTTCAGTAAACTTGAGAGTATGAGTTCGTACTCGCACAGGCGTAAATTACAAAG TTCTACCTTACTGACAGATTGGCCTCT**

**RTDNA: None**

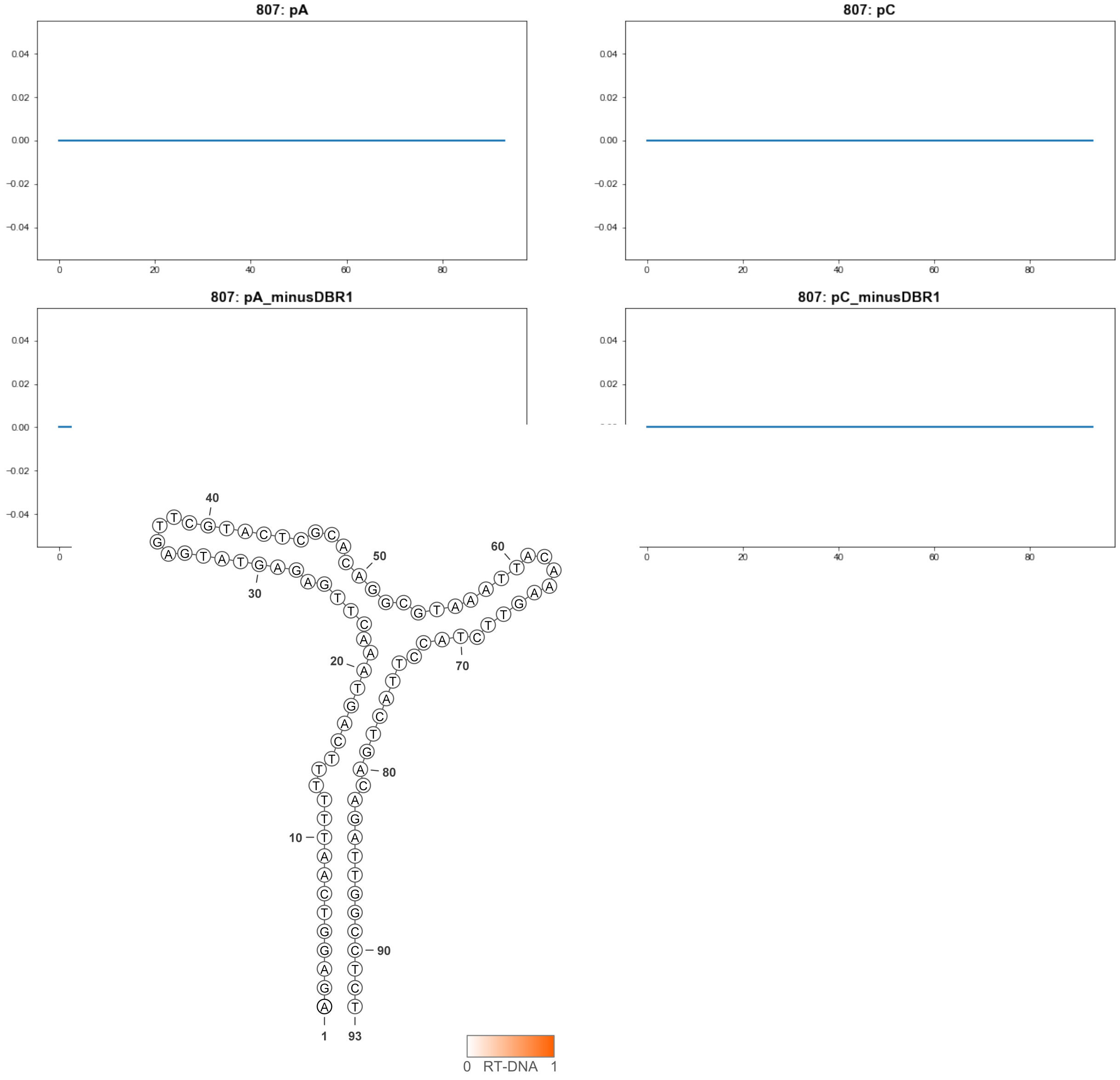


**807**

### Retron Terminal 808: Firmicutes

**Bacterial Vector for RTDNA: pSLS.568**

**Bacterial Vector for Editing: pAGD169**

**Phage Vector for Editing: pAGD397**

**Human Vector for Editing: pSLS.805 (U6)**

**RTDNA Production (relative to Eco1): 0.352 by PAGE**

**Bacterial Editing: 2.52302863 percent precise (0.776367532x Eco1)**

**Phage Editing: 31.41819369 percent precise (0.774919449x Eco1)**

**Human Editing (demultiplexed): 0.02724 percent precise (0.234643512x Eco1)**

**ncRNA:**

**ATGTTTAGTGGTGACTAGACTTTTAGCAAGTGCTAATCGAAGTAAGTATGTATTTAATTCTTATTTC AAAACCATAATGATGATTCTGCCCAAAATTGCAGAACCATCCTCACCTTTAAATTAAGAATTAAAA CCCTAAGAACTTCTTAATTTAAAGGTGAAGTAGTCACTAAACAT**

**RTDNA:**

**CACCTTTAAATTAAGAAGTTCTTAGGGTTTTAATTCTTAATTTAAAGGTGAGGATGGTTCTGCAATT TTGGGCAGAATCATCATTATG**

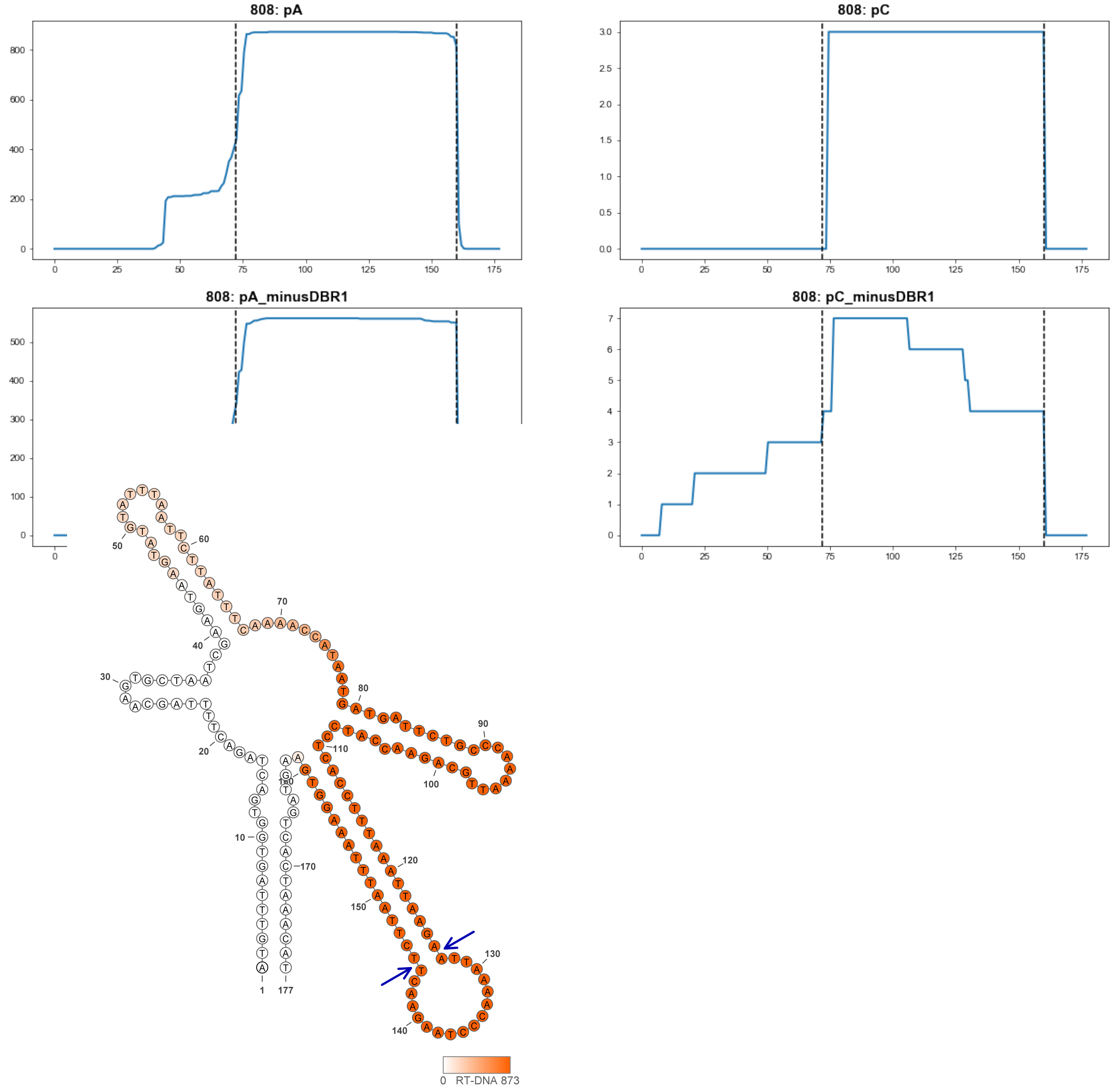


**808**

### Retron Terminal 810: Firmicutes

**Bacterial Vector for RTDNA: pSLS.569**

**Bacterial Vector for Editing: pAGD170**

**Phage Vector for Editing: None**

**Human Vector for Editing: pSLS.806 (H1)**

**RTDNA Production (relative to Eco1): 0.053647421 by PAGE**

**Bacterial Editing: 0.28861676 percent precise (0.088810994x Eco1)**

**Phage Editing: undetermined**

**Human Editing (demultiplexed): 0 percent precise (0x Eco1)**

**ncRNA:**

**TGGGTGACTAGACTTTTAGCAAGTGCTAATCGAAGTAAGTATGTATTGAATTCTTATTTCAAAAGA GAAATGGTAATTCTGTAGAAACCAACAGAATCACCATACCCCTTTAAATTAACAATTCATAATTTAA AGGGGAGGTAGTCACCTA**

**RTDNA:**

**CCCCTTTAAATTATGAATTGTTAATTTAAAGGGGTATGGTGATTCTGTTGGTTTCTACAGAATTACC ATTTCTCTTTTGAAATAAGAATTCAATACATACTTA**

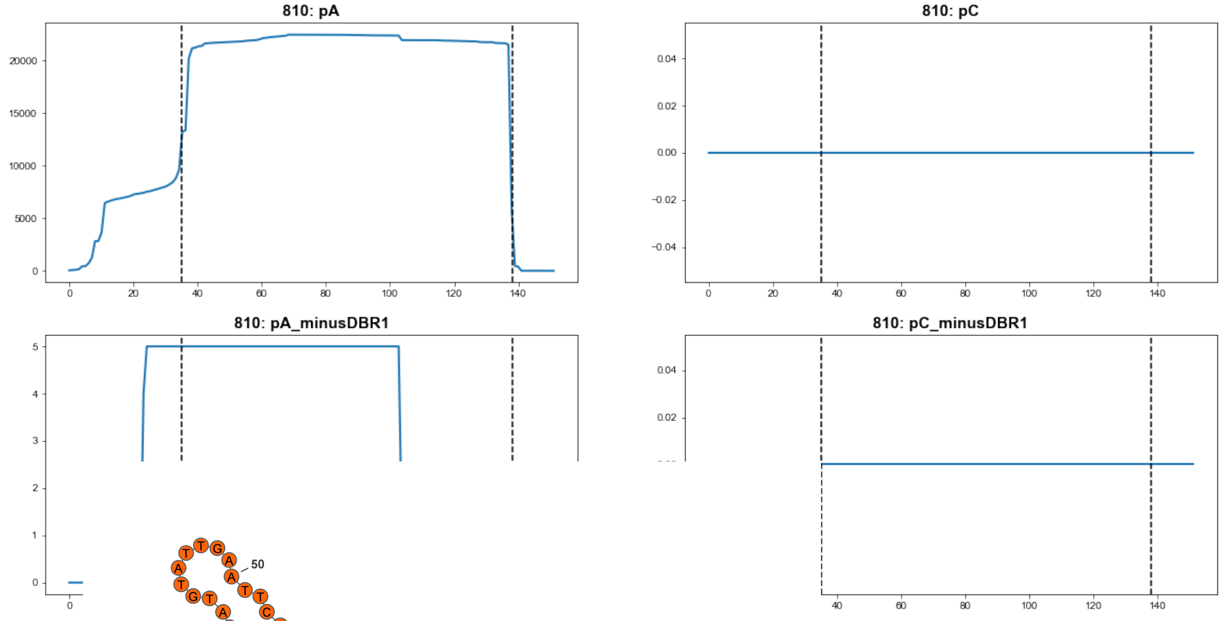


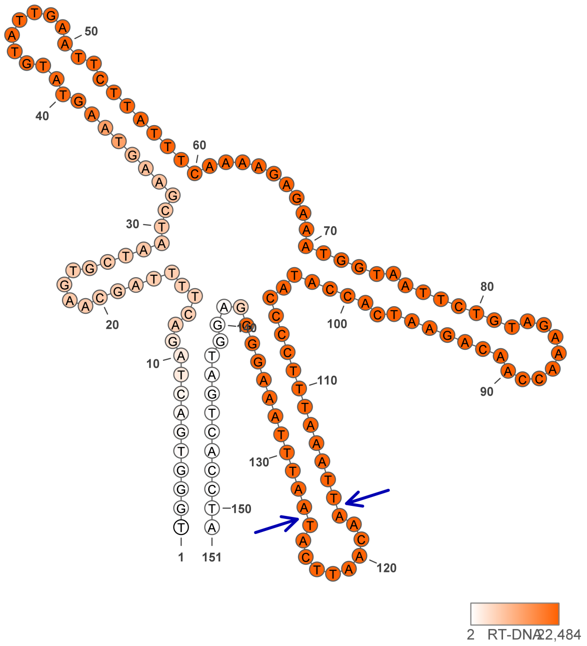


**810**

### Retron Terminal 812: Firmicutes

**Bacterial Vector for RTDNA: pSLS.583**

**Bacterial Vector for Editing: pAGD285**

**Phage Vector for Editing: None**

**Human Vector for Editing: pSLS.816 (H1)**

**RTDNA Production (relative to Eco1): 0.133844 by PAGE**

**Bacterial Editing: 1.21862471 percent precise (0.374986097x Eco1)**

**Phage Editing: undetermined**

**Human Editing (demultiplexed): undetermined**

**ncRNA:**

**ATAATGAGGGTAGTTAGCAATCTATAGCAATGTGCTATAGTAGGTAGTTTCTATCTTAATCAGTGAT**
**GTCATCAAACTGAAAGCGAGTTTGATGACGGCTCTGTTTAAGAAGGACGCTTAAACAGACCGTAA**
**CTACCCTTTTTGT**

**RTDNA:**

**GGTCTGTTTAAGCGTCCTTCTTAAACAGAGCCGTCATCAAACTCGCTTTCAGTTTGATGACATCACT**
**GATTAAGATAGAAACT**

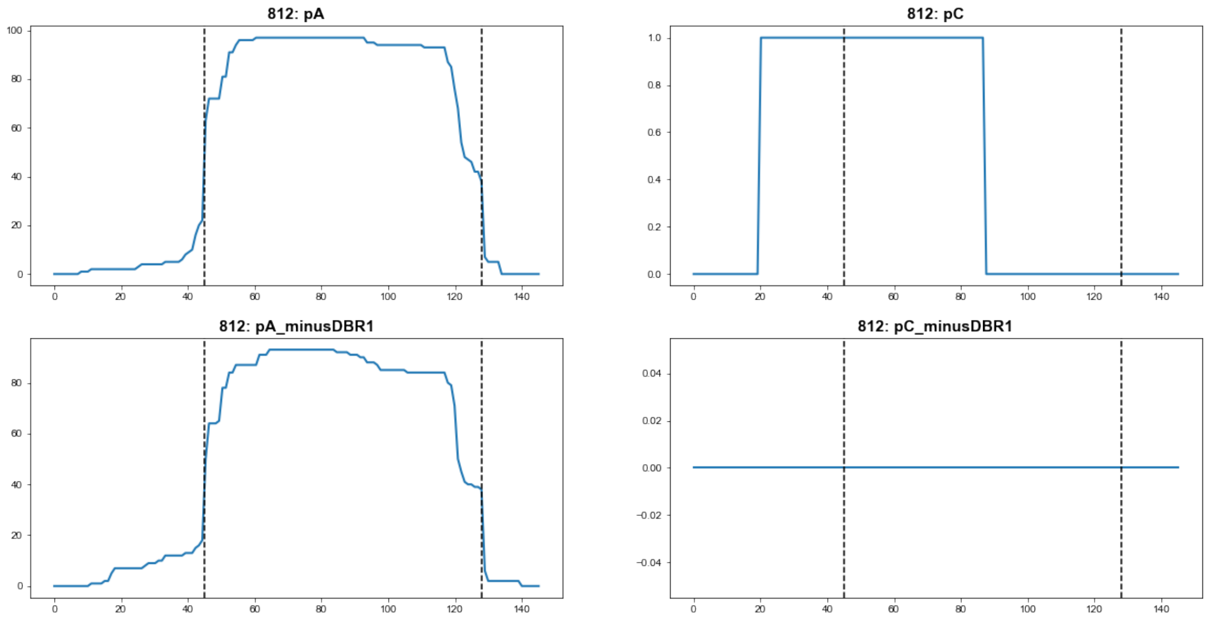


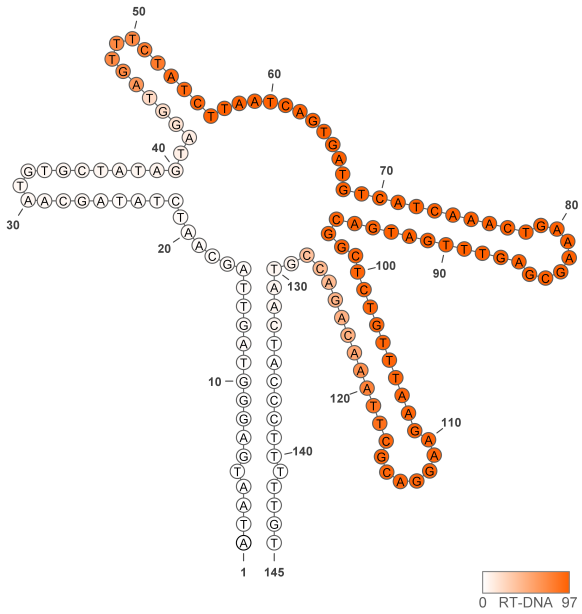


**812**

### Retron Terminal 813: Firmicutes

**Bacterial Vector for RTDNA: None**

**Bacterial Vector for Editing: None**

**Phage Vector for Editing: None**

**Human Vector for Editing: pSLS.871**

**RTDNA Production (relative to Eco1): undetermined**

**Bacterial Editing: undetermined**

**Phage Editing: undetermined**

**Human Editing (demultiplexed): 0.071522 percent precise (0.616085657x Eco1)**

**ncRNA:**

**CTCTGGACATTCAATAGCATGGGAGCGATTCGCCCCAATAAAGAAAGGCACTTACTTTATCTAACA**
**AATCCGTCACTACGTTCCGTATTTGTTAGATAAGCGTCGCAAACGACGCTTTGATTTTCGTATTGAA**
**TGTCCGGAG**

**no RTDNA sequencing data**

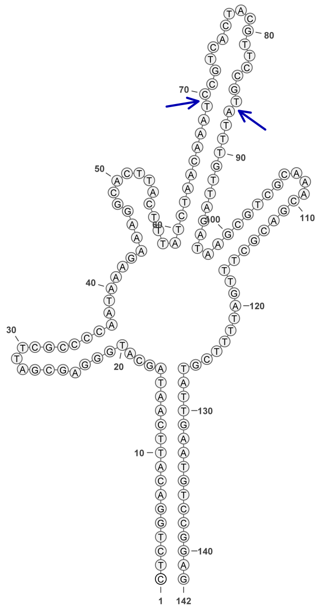

813

#### Retron Terminal 814: Firmicutes

**Bacterial Vector for RTDNA: pSLS.570**

**Bacterial Vector for Editing: None**

**Phage Vector for Editing: None**

**Human Vector for Editing: pSLS.807 (H1)**

**RTDNA Production (relative to Eco1): 0.010570244 by PAGE**

**Bacterial Editing: undetermined**

**Phage Editing: undetermined**

**Human Editing (demultiplexed): 15.93654 percent precise (137.2762746x Eco1)**

**ncRNA:**

**TAAAACCGGGAACGATCAGACCGGGGTGAATTCGCCCCCTTGATCAAACGGCACTAACCACTGTT TGCCGTGCGTGCGTCTTGTTCATTTCATTCTCTGCGACGCACGCACGGCAAACAGACAGATCCATT ATTATTACAATTTATTTAGTGATCGTTCCCGGTTTTA**

**RTDNA:**

**AAATAAATTGTAATAATAATGGATCTGTCTGTTTGCCGTGCGTGCGTCGCAGAGAATGAAATGAAC AAGACGCACGCACGG**

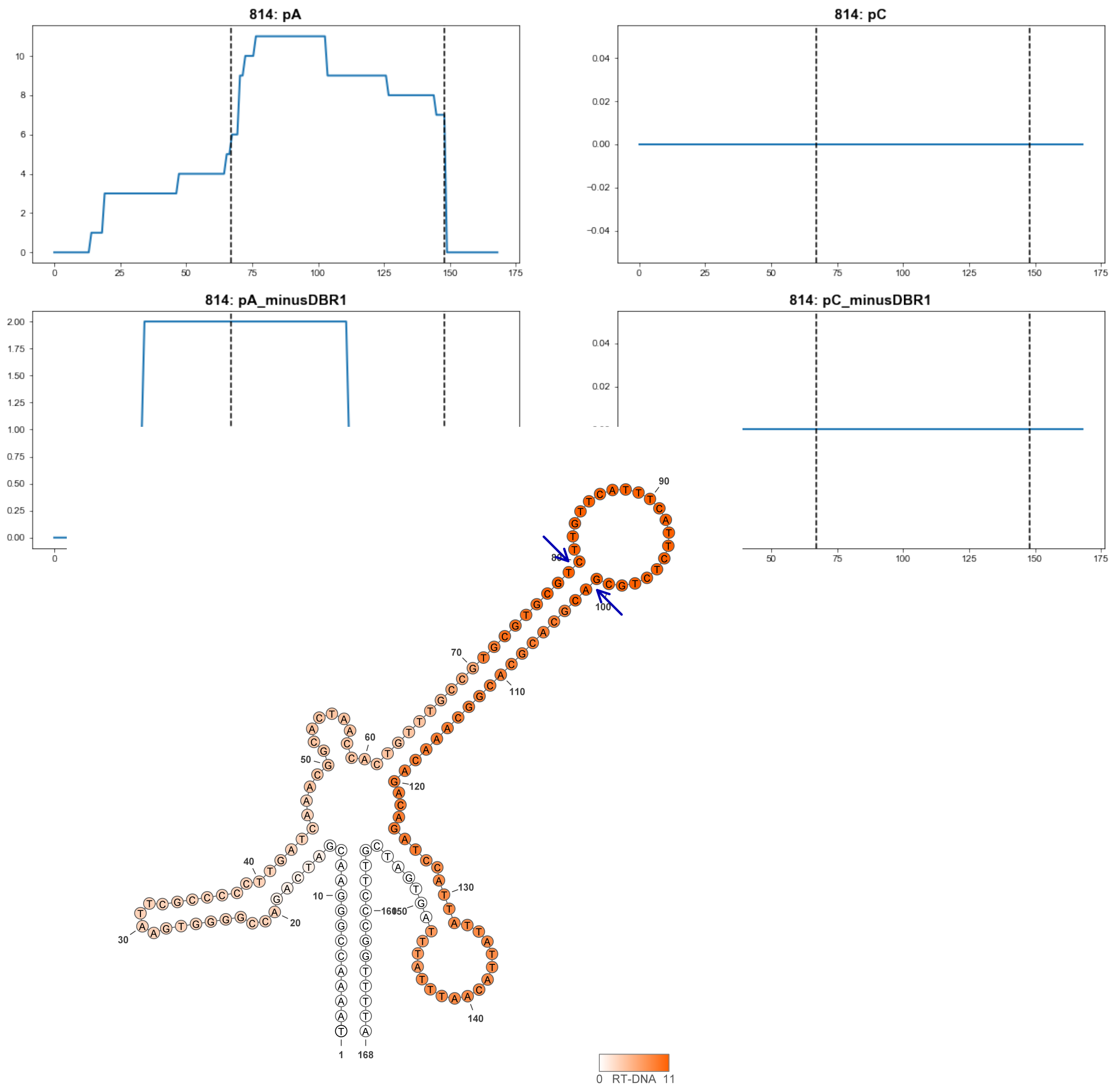


**814**

### Retron Terminal 814.2: Firmicutes

**Bacterial Vector for RTDNA: None**

**Bacterial Vector for Editing: None**

**Phage Vector for Editing: None**

**Human Vector for Editing: pSLS.975**

**RTDNA Production (relative to Eco1): 0.010570244 by PAGE**

**Bacterial Editing: undetermined**

**Phage Editing: undetermined**

**Human Editing (demultiplexed): undetermined**

**ncRNA:**

**TAAAACCGGGAACGATCAGACCGGGGTGAATTCGCCCCCTTGATCAAACGGCACTAACCACTGTT TGCCGTGCGTGCGTCTTGTTCATTTCATTCTCTGCGACGCACGCACGGCAAACAGACAGATCCATT ATTATTACAATTTATTTAGTGATCGTTCCCGGTTTTA**

**RTDNA:**

**AAATAAATTGTAATAATAATGGATCTGTCTGTTTGCCGTGCGTGCGTCGCAGAGAATGAAATGAAC AAGACGCACGCACGG**

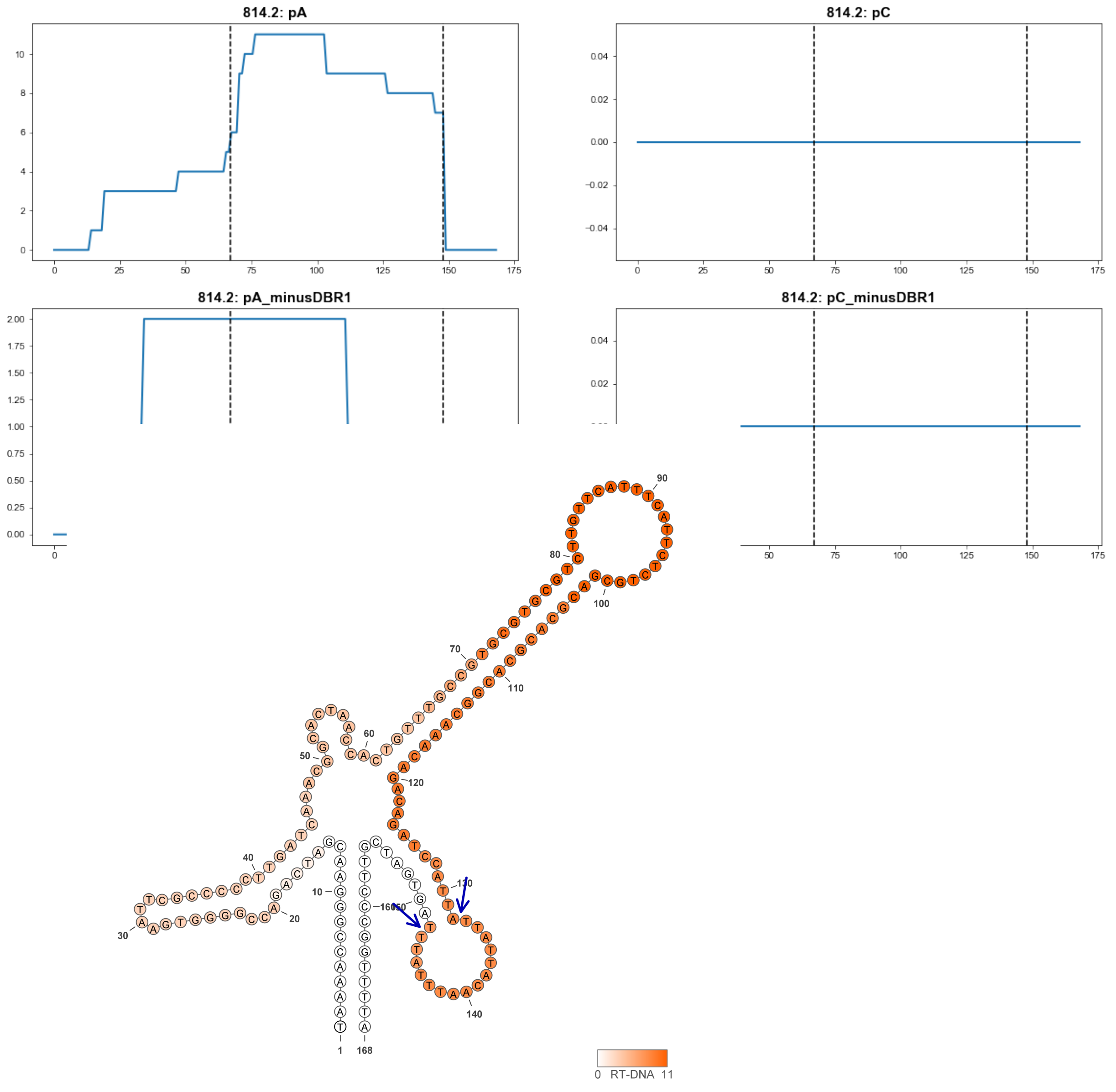


**814.2**

### Retron Terminal 814.9: Firmicutes

**Bacterial Vector for RTDNA: None**

**Bacterial Vector for Editing: None**

**Phage Vector for Editing: None**

**Human Vector for Editing: pSLS.974**

**RTDNA Production (relative to Eco1): 0.010570244 by PAGE**

**Bacterial Editing: undetermined**

**Phage Editing: undetermined**

**Human Editing (demultiplexed): undetermined**

**ncRNA:**

**TAAAACCGGGAACGATCAGACCGGGGTGAATTCGCCCCCTTGATCAAACGGCACTAACCACTGTT TGCCGTGCGTGCGTCTTGTTCATTTCATTCTCTGCGACGCACGCACGGCAAACAGACAGATCCATT ATTATTACAATTTATTTAGTGATCGTTCCCGGTTTTA**

**RTDNA:**

**AAATAAATTGTAATAATAATGGATCTGTCTGTTTGCCGTGCGTGCGTCGCAGAGAATGAAATGAAC AAGACGCACGCACGG**

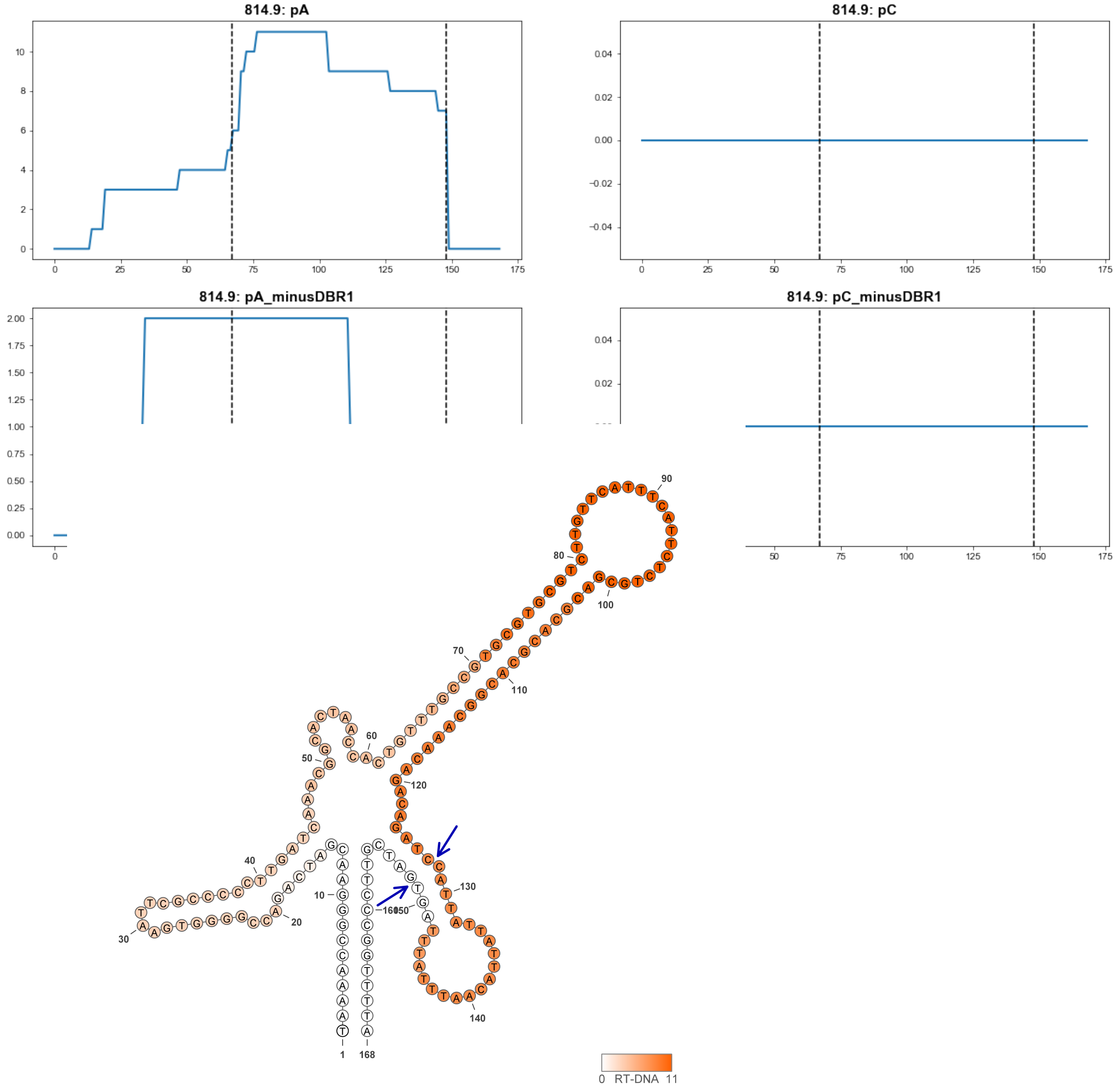


**814.9**

### Retron Terminal 815: Firmicutes

**Bacterial Vector for RTDNA: pSLS.641**

**Bacterial Vector for Editing: None**

**Phage Vector for Editing: None**

**Human Vector for Editing: pSLS.872**

**RTDNA Production (relative to Eco1): 0.326941875 by PAGE**

**Bacterial Editing: undetermined**

**Phage Editing: undetermined**

**Human Editing (demultiplexed): 5.324956 percent precise (45.86880981x Eco1)**

**ncRNA:**

**AGAAAGAGTGACTGGATGGAGGTGTATTCGCCTCCCTAAGTAAAGGCATTAACAGCGGCTGTCGA GCGAGTTACTCGTGCACTGCTTTTTCTCGCAACTCGCTCGACAGCCGCCTATAGTGTTAATCTTAG GAGATTTAAATTATGAGCAGTCACTCTTTTT**

**RTDNA:**

**TCATAATTTAAATCTCCTAAGATTAACACTATAGGCGGCTGTCGAGCGAGTTGCGAGAAAAAGCA GTGCACGAGTAACTCGCTCG**

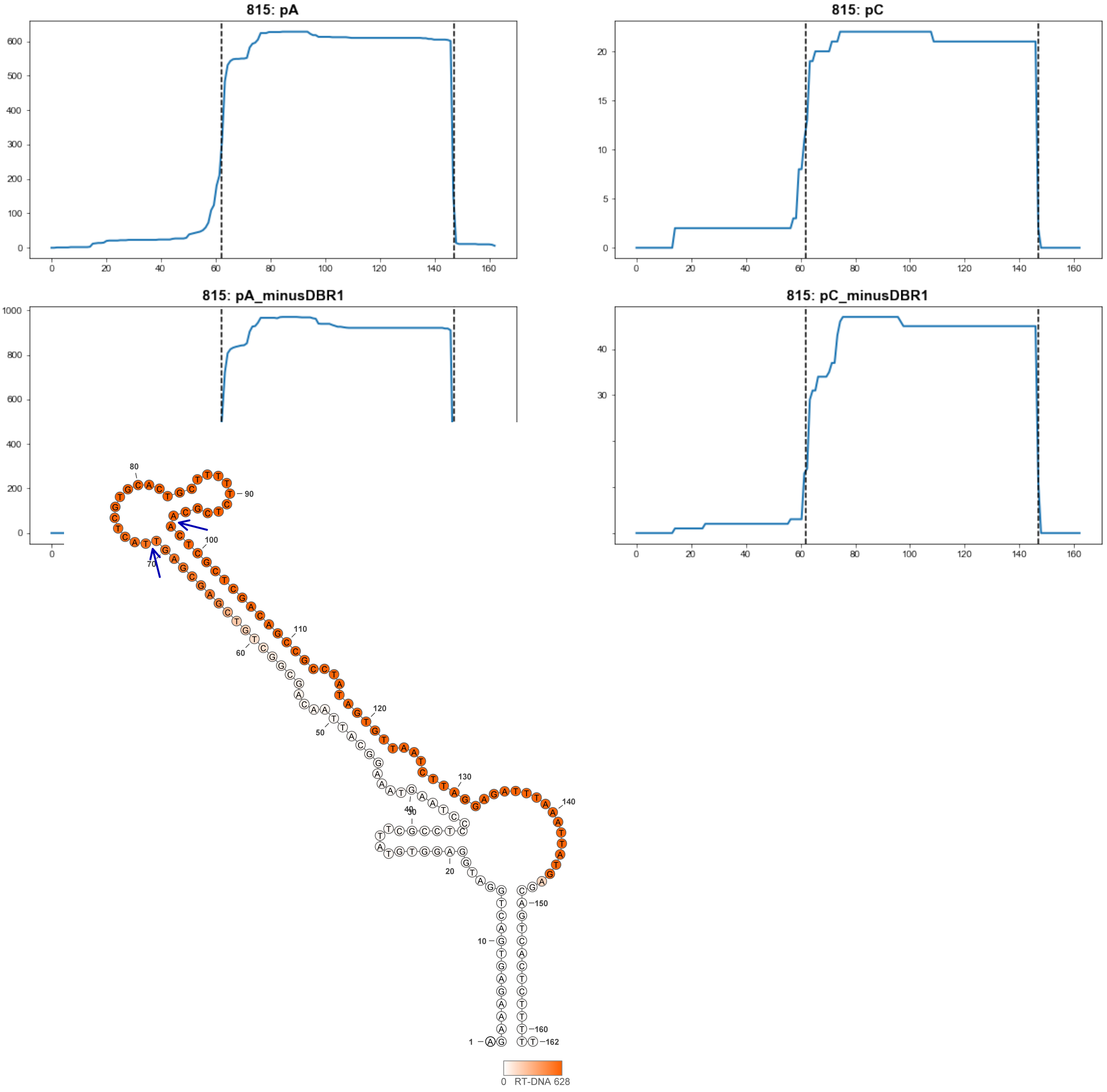


815

### Retron Terminal 816: Firmicutes

**Bacterial Vector for RTDNA: pSLS.571**

**Bacterial Vector for Editing: pAGD163**

**Phage Vector for Editing: pAGD398**

**Human Vector for Editing: pSLS.808 (H1)**

**RTDNA Production (relative to Eco1): 0.748840019 by PAGE**

**Bacterial Editing: 0.85961746 percent precise (0.264515066x Eco1)**

**Phage Editing: 3.461016098 percent precise (0.085364828x Eco1)**

**Human Editing (demultiplexed): 0.58818 percent precise (5.066542626x Eco1)**

**ncRNA:**

**AAGAGAACAACTAGAATGAGGTGATTCACCTCCTTGTTTAACGGCACTAATTTTTACACTGTAATTT ACTAACGTAAATTACAGTGTAAAAGATCATAAGTAGTTGTTCTCTT**

**RTDNA:**

**TATGATCTTTTACACTGTAATTTACGTTAGTAAATTACAGTGTAAAAATTAGTGCCGTTAAACAAGG AGGTGAATCACCTCATT**

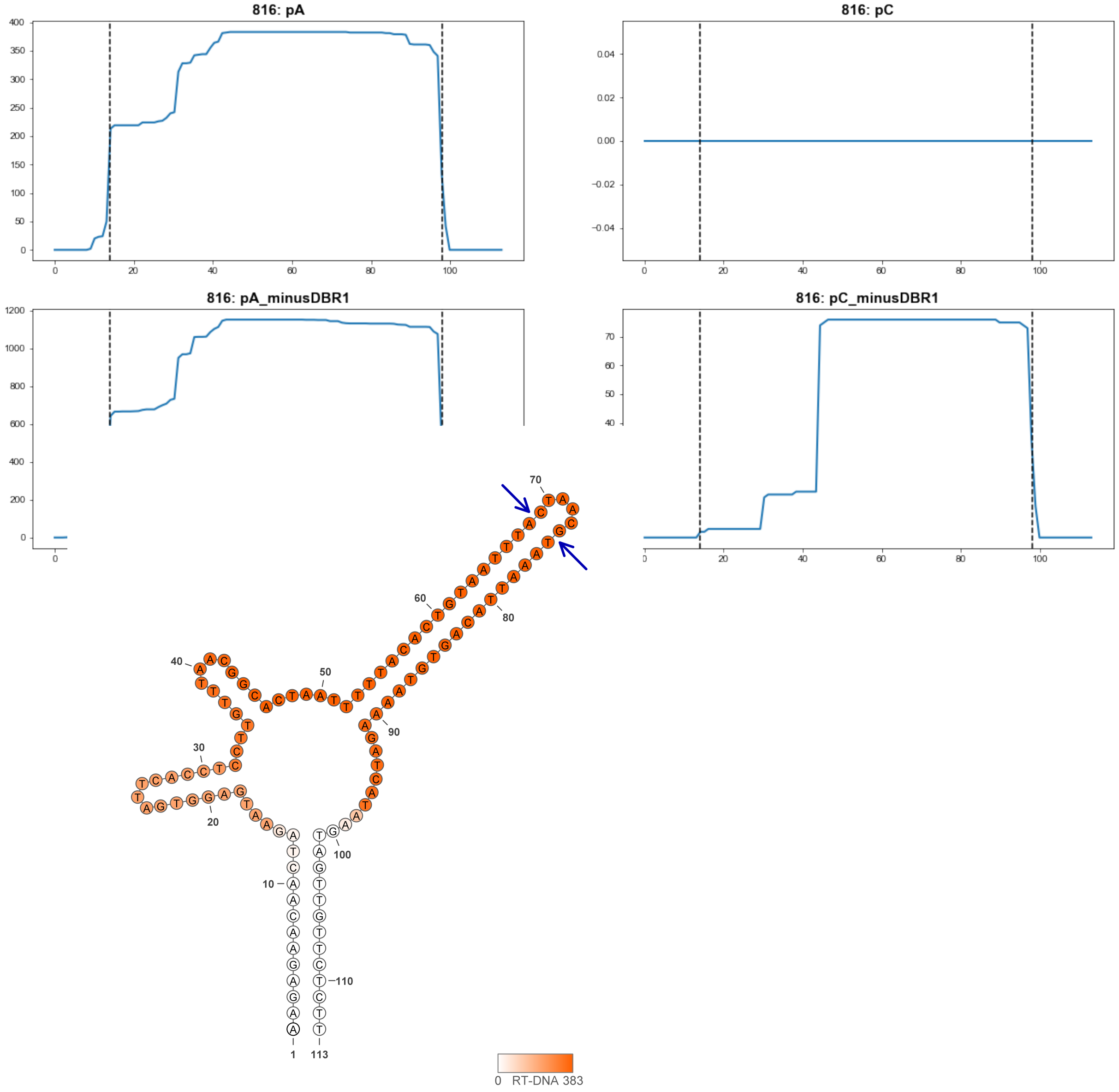


**816**

### Retron Terminal 817: Firmicutes

**Bacterial Vector for RTDNA: None**

**Bacterial Vector for Editing: None**

**Phage Vector for Editing: None**

**Human Vector for Editing: pSLS.959 (U6)**

**RTDNA Production (relative to Eco1): undetermined**

**Bacterial Editing: undetermined**

**Phage Editing: undetermined**

**Human Editing (demultiplexed): 1.085725 percent precise (9.35236151x Eco1)**

**ncRNA:**

**TGAGAATGTATAGTAATCTAACAATTCCAGAGGTAATAATTTTGAAATCTATAAAAGAAAATGGAA**
**GAATTGATTTTGTTAAATTGGGCATTGAATTAGGATTACCAGCTGGTATGATAGTAAATTTAGTTCA**
**GGAACTATATGATAGAAGATTTATAAAAGTTAAACAAAAAAGAATATATCTTA**

**no RTDNA sequencing data**

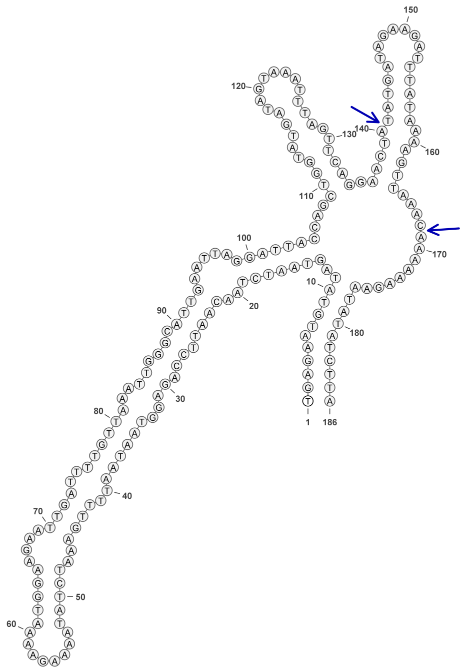


817

### Retron Terminal 818: Firmicutes

**Bacterial Vector for RTDNA: None**

**Bacterial Vector for Editing: None**

**Phage Vector for Editing: None**

**Human Vector for Editing: pSLS.995**

**RTDNA Production (relative to Eco1): undetermined**

**Bacterial Editing: undetermined**

**Phage Editing: undetermined**

**Human Editing (demultiplexed): 0.000945 percent precise (0.008140166x Eco1)**

**ncRNA:**

**TTCACTGAATATTATGAACAGCGTTATCTCTTCGGATATGATTAATTCGTAGGTGAAACAATGTGTC**
**GAGATAGAATTAAAAAGTAGGCTACAAAGTAACTATCAACGGTGAGGAATTATGTAGCTACAAGG**
**CTAAGTGAAGTAAAGGTAAAACTTTCGGTGGA**

**no RTDNA sequencing data**

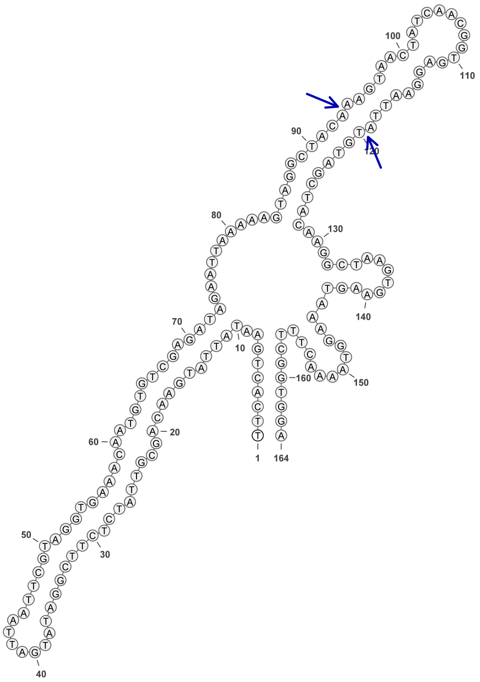


**818**

### Retron Terminal 824: Proteobacteria

**Bacterial Vector for RTDNA: pSLS.584**

**Bacterial Vector for Editing: pAGD286**

**Phage Vector for Editing: pAGD399**

**Human Vector for Editing: pSLS.817 (H1)**

**RTDNA Production (relative to Eco1): 0.719435522 by PAGE**

**Bacterial Editing: 2.64486484 percent precise (0.81385806x Eco1)**

**Phage Editing: 8.461875429 percent precise (0.208709384x Eco1)**

**Human Editing (demultiplexed): 0.549712 percent precise (4.735181883x Eco1)**

**ncRNA:**

**GTGGGAGCCTCAGGCGAGGGTGTGTATCATGCCCGTTCTGCCAAGACCCACCAAAGAAGGGCAC CGTGGAGGCACACGGCGCTCCTTCGAAGCACCGCGTACCTCCACGGTGCAATGCGAAAGCAACT TGAGGCTTTGCTTAGTATGAGGCTCCCAC**

**RTDNA:**

**GCATTGCACCGTGGAGGTACGCGGTGCTTCGAAGGAGCGCCGTGTGCCTCCACGGTGCCCTTCTT TGGTGGGT**

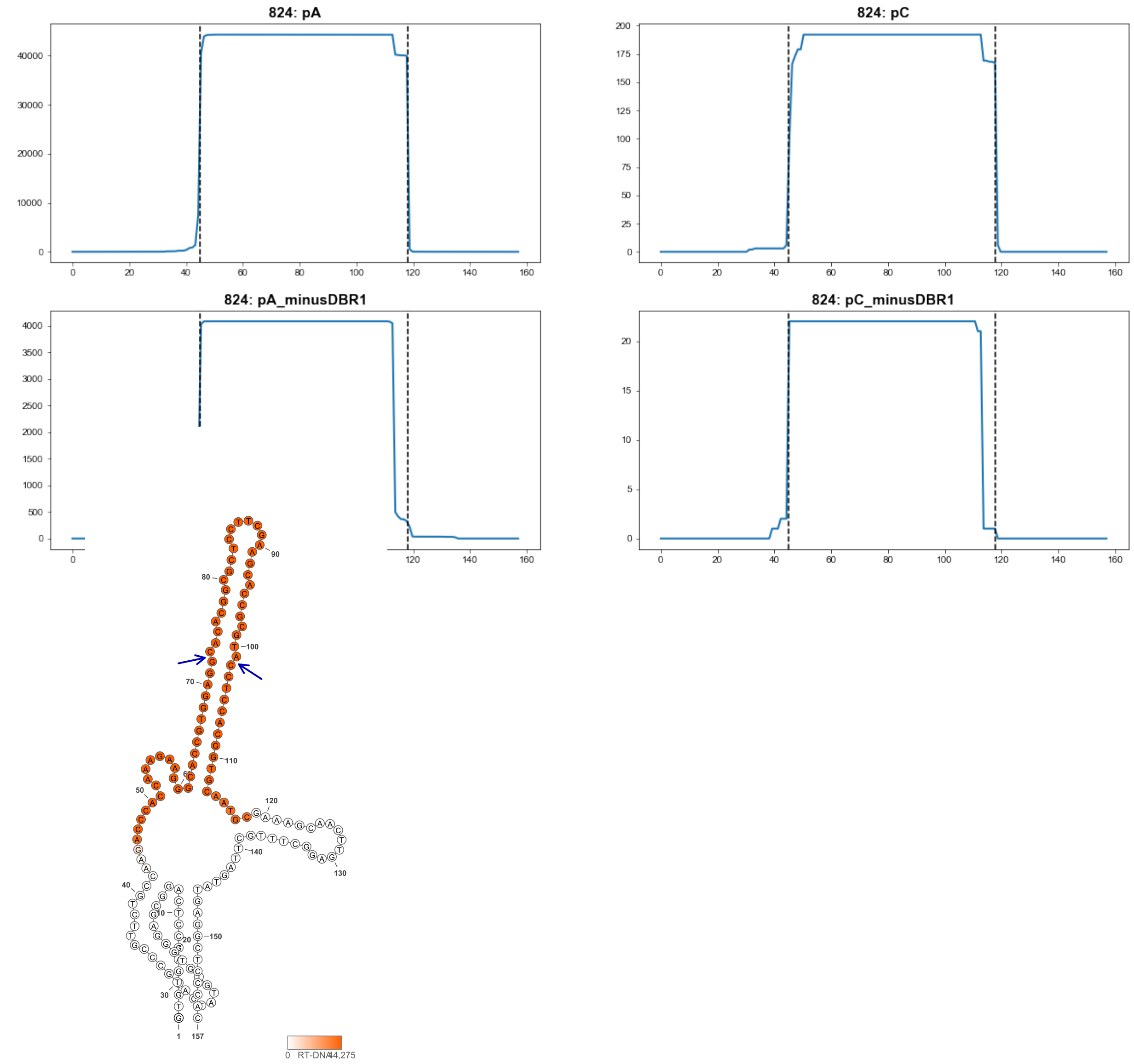


824

### Retron Terminal 825: Proteobacteria

**Bacterial Vector for RTDNA: None**

**Bacterial Vector for Editing: None**

**Phage Vector for Editing: None**

**Human Vector for Editing: pSLS.873**

**RTDNA Production (relative to Eco1): undetermined**

**Bacterial Editing: undetermined**

**Phage Editing: undetermined**

**Human Editing (demultiplexed): 0 percent precise (0x Eco1)**

**ncRNA:**

**GTGGGAGCCTCAGGCGAGGGTGTGTATCATGCCCATCTGCCAAGATCCTGCAAAGAAGGGCACC**
**GGTCACATGGCCGGCACTCCTTCGGAGCACCGGCCATGTGACCGGCGCAGTACGAAAGCAATTT**
**AAGGCTTTGCTAAGTATGAGGCTCCCAC**

**no RTDNA sequencing data**

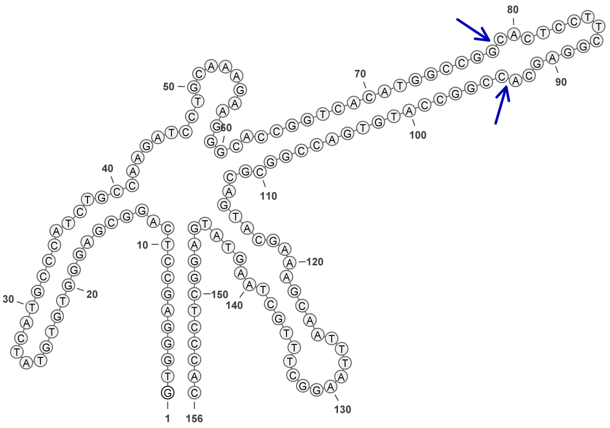


**825**

### Retron Terminal 839: Proteobacteria

**Bacterial Vector for RTDNA: None**

**Bacterial Vector for Editing: None**

**Phage Vector for Editing: None**

**Human Vector for Editing: pSLS.961**

**RTDNA Production (relative to Eco1): undetermined**

**Bacterial Editing: undetermined**

**Phage Editing: undetermined**

**Human Editing (demultiplexed): 0 percent precise (0x Eco1)**

**ncRNA:**

**GGTGGTGGCGGAATCACGAGAGTATGTATCATGCTCATTTGTGAAGATCATGCACAGAAGGGCGC**

**CTGCACACAGGCAGACGCTCCTTCGAATCGTCTGCCTGTGTGCAGGCGGCATCAACGAAAGCTAT**

**CAGCTTTGCTAAAGGTAGATCTATGAGAACGCCACCACC**

**no RTDNA sequencing data**

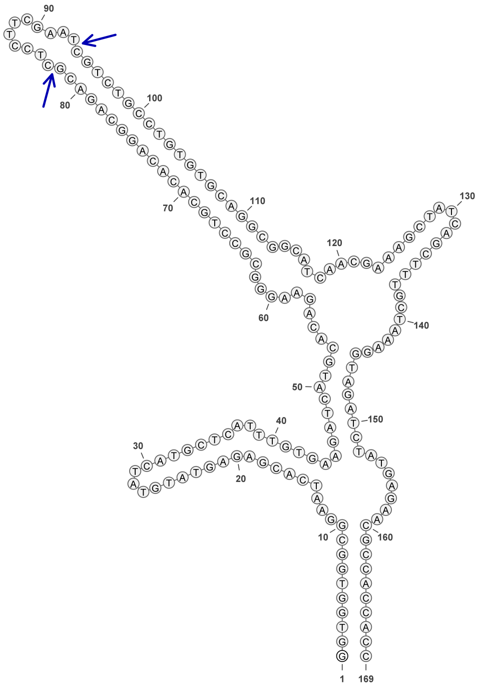

839

### Retron Terminal 842: Proteobacteria

**Bacterial Vector for RTDNA: pSLS.572**

**Bacterial Vector for Editing: None**

**Phage Vector for Editing: None**

**Human Vector for Editing: None**

**RTDNA Production (relative to Eco1): 0.079180737 by PAGE**

**Bacterial Editing: undetermined**

**Phage Editing: undetermined**

**Human Editing (demultiplexed): undetermined**

**ncRNA:**

**GGTAGTGGCGTTCACGAGGGTGTGTATCATACCCATTTGTGAAGGTCGCGTACAGAATGTGCGCC TGTGAAGCAGCAGGCGCTCCTTCGAAACGCCTGCTGCTTCACAGGCGTCATCAACGAAAGCCTCA TCAGCTTTGCTTAAGGACTGACCTGGATGAAAACGCCACTACC**

**RTDNA:**

**GTTGATGACGCCTGTGAAGCAGCAGGCGTTTCGAAGGAGCGCCTGCTGCTTCACAGGCGCACATT CTGTACGCGACCTTCAC**

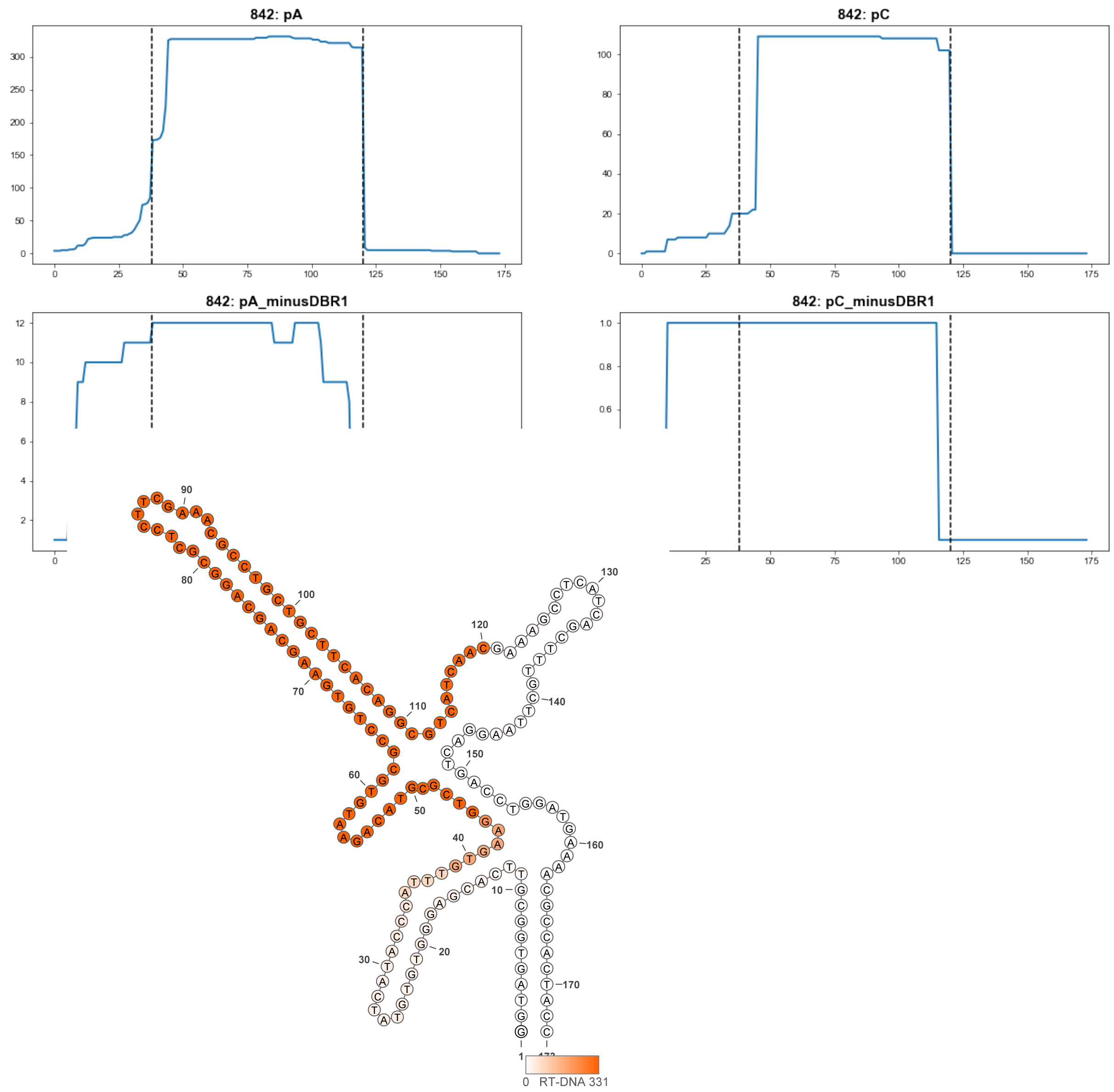


**842**

### Retron Terminal 844: Proteobacteria

**Bacterial Vector for RTDNA: pSLS.573**

**Bacterial Vector for Editing: None**

**Phage Vector for Editing: None**

**Human Vector for Editing: pSLS.809 (H1)**

**RTDNA Production (relative to Eco1): 0 by PAGE**

**Bacterial Editing: undetermined**

**Phage Editing: undetermined**

**Human Editing (demultiplexed): 0.022069 percent precise (0.190100869x Eco1)**

**ncRNA:**

**GGTAGTGGCGTTCGCGAGAGTGTGTATCACACTCATTTGCGAAGGTCGCGGAAAGAATGGGCGC CTGAGAAGCAGCAGGCGTTCCTTCGAAACGCCTGCTGCTTCTCAGGCGTCATCAACGAAAGTTTT CATCAGCTTTGCTAAAGGATGGACCTTGTGAAAACGCCACTACC**

**RTDNA: None**

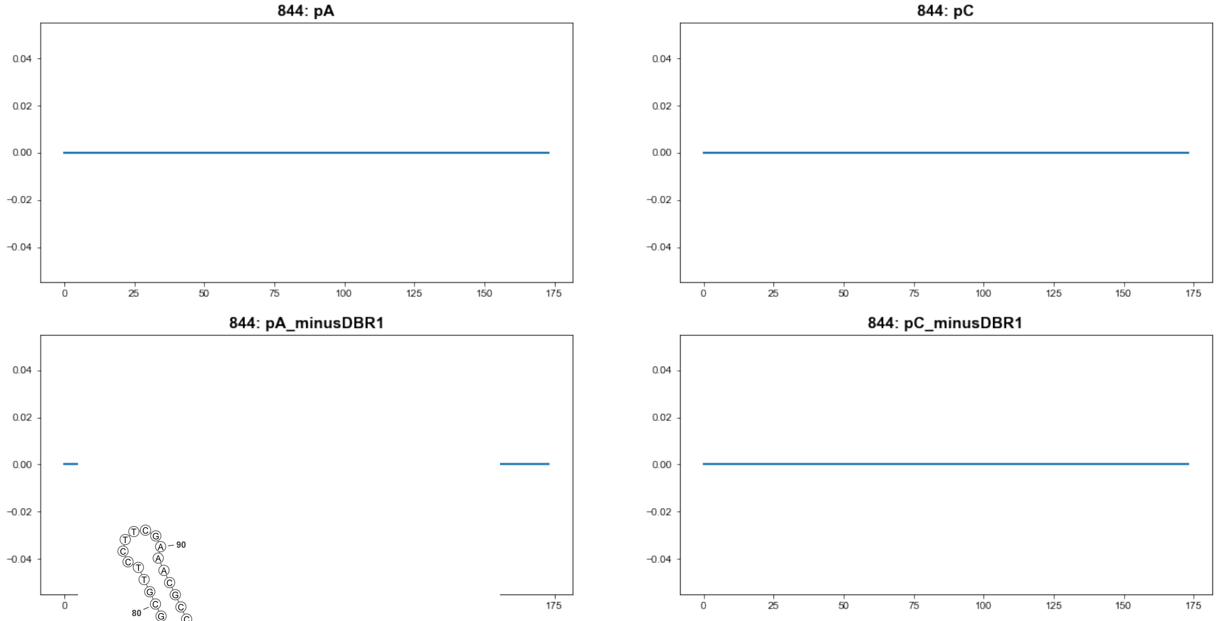


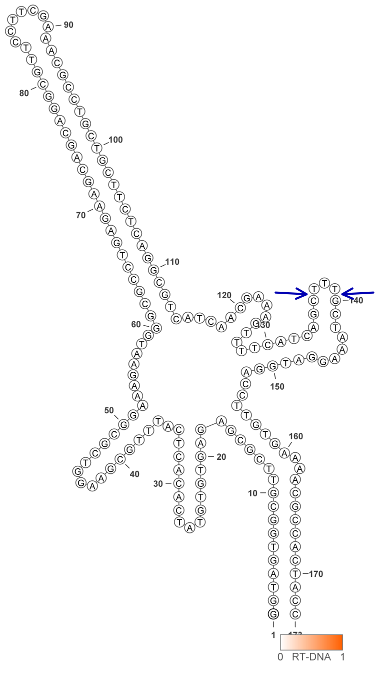


844

### Retron Terminal 848: Verrucomicrobia

**Bacterial Vector for RTDNA: pSLS.671**

**Bacterial Vector for Editing: None**

**Phage Vector for Editing: None**

**Human Vector for Editing: pSLS.906B**

**RTDNA Production (relative to Eco1): 0 by PAGE**

**Bacterial Editing: undetermined**

**Phage Editing: undetermined**

**Human Editing (demultiplexed): 0.025155 percent precise (0.216683464x Eco1)**

**ncRNA:**

**GTTTGACTGTTAGCTAGGTCCGATGGAAGCCCCACGACGGGTCCACTTTCCCAACAATCAACCAA AGGTGTCAAGTTTTCCACCGACTTATTTCTGGGTTGCGGTTGTTCGACGGTCAAGC**

**RTDNA:**

**TGACCGTCGAACAACCGCAACCCAGAAATAAGTCGGTGGAAAACTTGACACCTTTGGTTGATTGT TGGGAAAGTGGACCCGTCGTGGGGCTTCC**

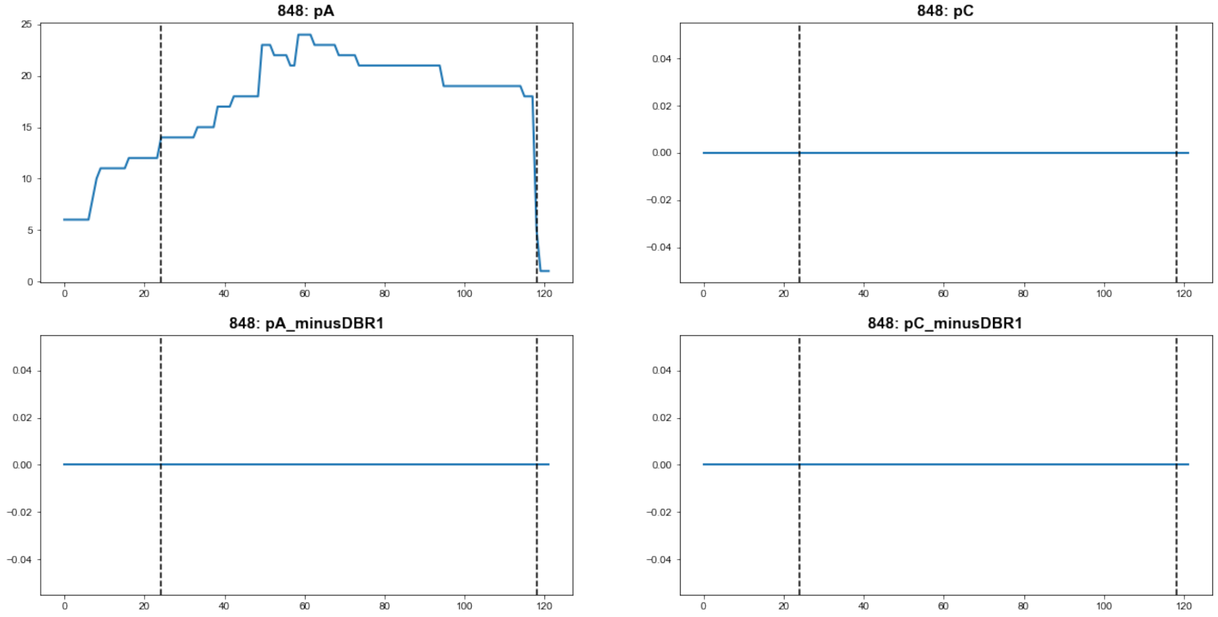


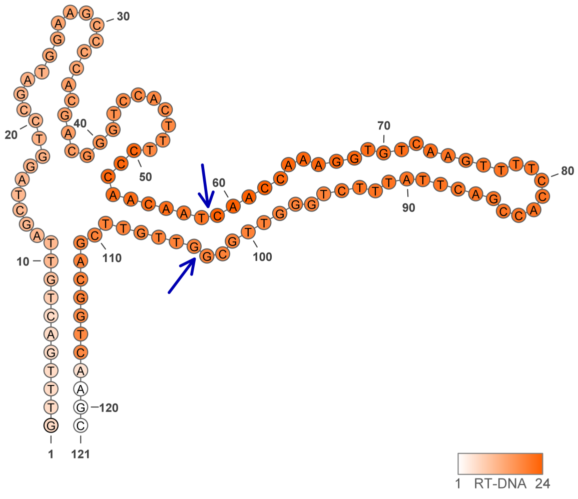


**848**

### Retron Terminal 864: Proteobacteria

**Bacterial Vector for RTDNA: pSLS.672**

**Bacterial Vector for Editing: pAGD260**

**Phage Vector for Editing: None**

**Human Vector for Editing: pSLS.907**

**RTDNA Production (relative to Eco1): 0.033586531 by PAGE**

**Bacterial Editing: 0.23321499 percent precise (0.071763175x Eco1)**

**Phage Editing: undetermined**

**Human Editing (demultiplexed): 0.030041 percent precise (0.258771136x Eco1)**

**ncRNA:**

**GGGAGCTTACAGGCAGATTTTTTTGTCGCGTTTCGCGACATTTGAAAACATCATGTGGGGGATTGC**
**TTCCTGACGTGACTCAACACGTTGACGCGAAATTCGTGTCGGCGGACGCATATGAGTGCCTGTAA**
**GCGTCCC**

**RTDNA:**

**TCATATGCGTCCGCCGACACGAATTTCGCGTCAACGTGTTGAGTCACGTCAGGAAGCAATCCCCC**
**ACAT**

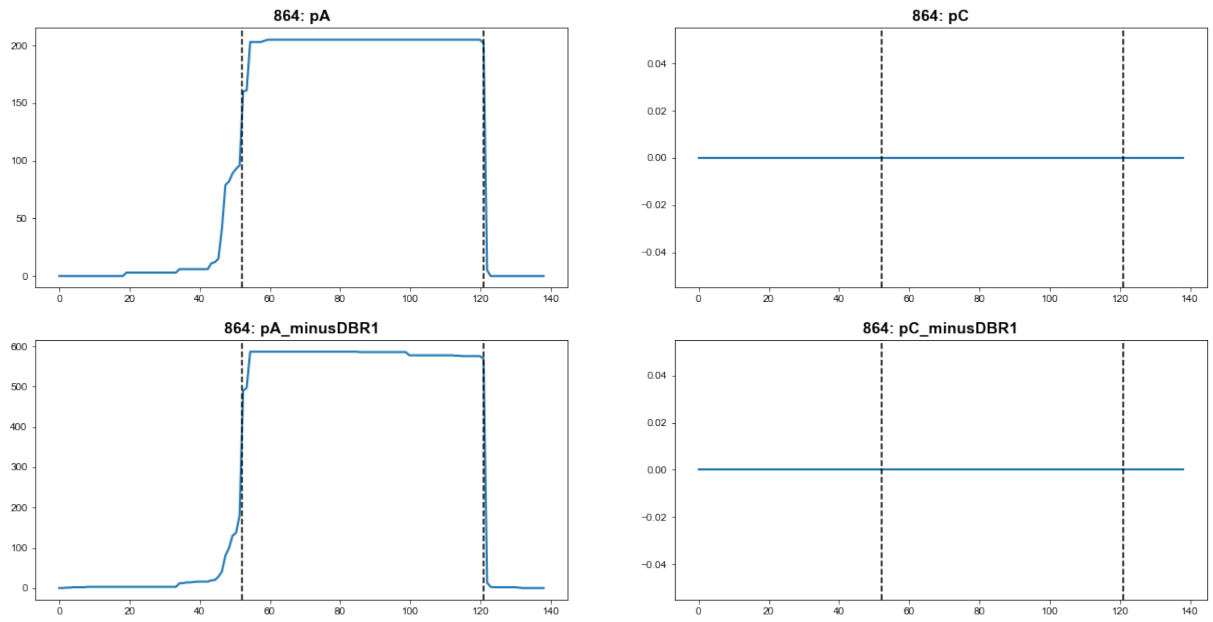


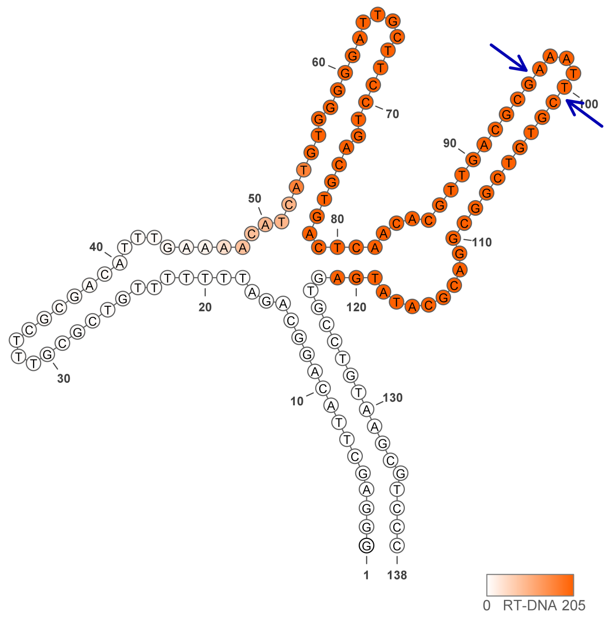


864

### Retron Terminal 866: Proteobacteria

**Bacterial Vector for RTDNA: None**

**Bacterial Vector for Editing: None**

**Phage Vector for Editing: None**

**Human Vector for Editing: pSLS.962**

**RTDNA Production (relative to Eco1): undetermined**

**Bacterial Editing: undetermined**

**Phage Editing: undetermined**

**Human Editing (demultiplexed): 0 percent precise (0x Eco1)**

**ncRNA:**

**GATGAGGATGAAAGCTATGGATGGGTTTCGGTTGTCAATGAAGGCAAAGCGTGAAGGTGGGTTGC**
**ATACCAAAAAAAATGTGATAAATTTCGCCTGGGCTTATAGACAGATTTTTTGTCATGCTTCTGTGAC**
**AAGTGGCACCAATGTTGGAGTCCATCGCTCTAATAGTTCTTCTCAGAATGGTCATCTAAGGTCATC**
**TCTGGTAACGCGTAAGAGTGTCTATAAGCCCTCCTCTTC**

**no RTDNA sequencing data**

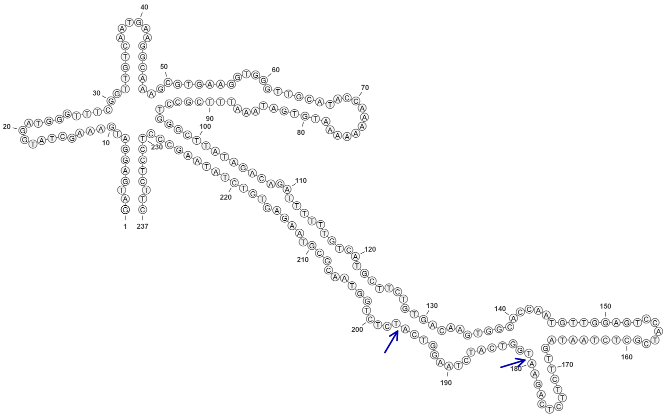

**866**

### Retron Terminal 888: Proteobacteria

**Bacterial Vector for RTDNA: pSLS.630**

**Bacterial Vector for Editing: None**

**Phage Vector for Editing: None**

**Human Vector for Editing: pSLS.855**

**RTDNA Production (relative to Eco1): 0 by PAGE**

**Bacterial Editing: undetermined**

**Phage Editing: undetermined**

**Human Editing (demultiplexed): 0 percent precise (0x Eco1)**

**ncRNA:**

**GAACTCCGTAGCAACTCGTCACAGAACGCCCGATTTTATCAATGTAACCATCAACAGCTAAGGCA CCAGAACAATAGATGGATAGATCGGCGCCCAACTGAAGCATAAGATCGCTACCGCAAGGTCCAC TGAGATGCTATGCTCGCCGTGGACTCGCGGACAGATTTTTTGCCATCGATTCATGGCATGTGAAAC AAATGTTCCGAGATATGCCCTCGGATGTCCTCTGTGTAGCAGTAGCGCGGTGTATC**

**RTDNA:**

**ATATCTCGGAACATTTGTTTCACATGCCATGAATCGATGGCAAAAAATCTGTCCGCGAGTCCACGG CGAGCATAGCATCTCAGTGGACCTTGCGGTAGCGATCTTATGCTTCAGTTGGGCGCCGATCTATC CATCTATTGTTCTGGTGCCTTAGCTGTTGATGGTTACATTGATAAAATCGGGCGTTCT**

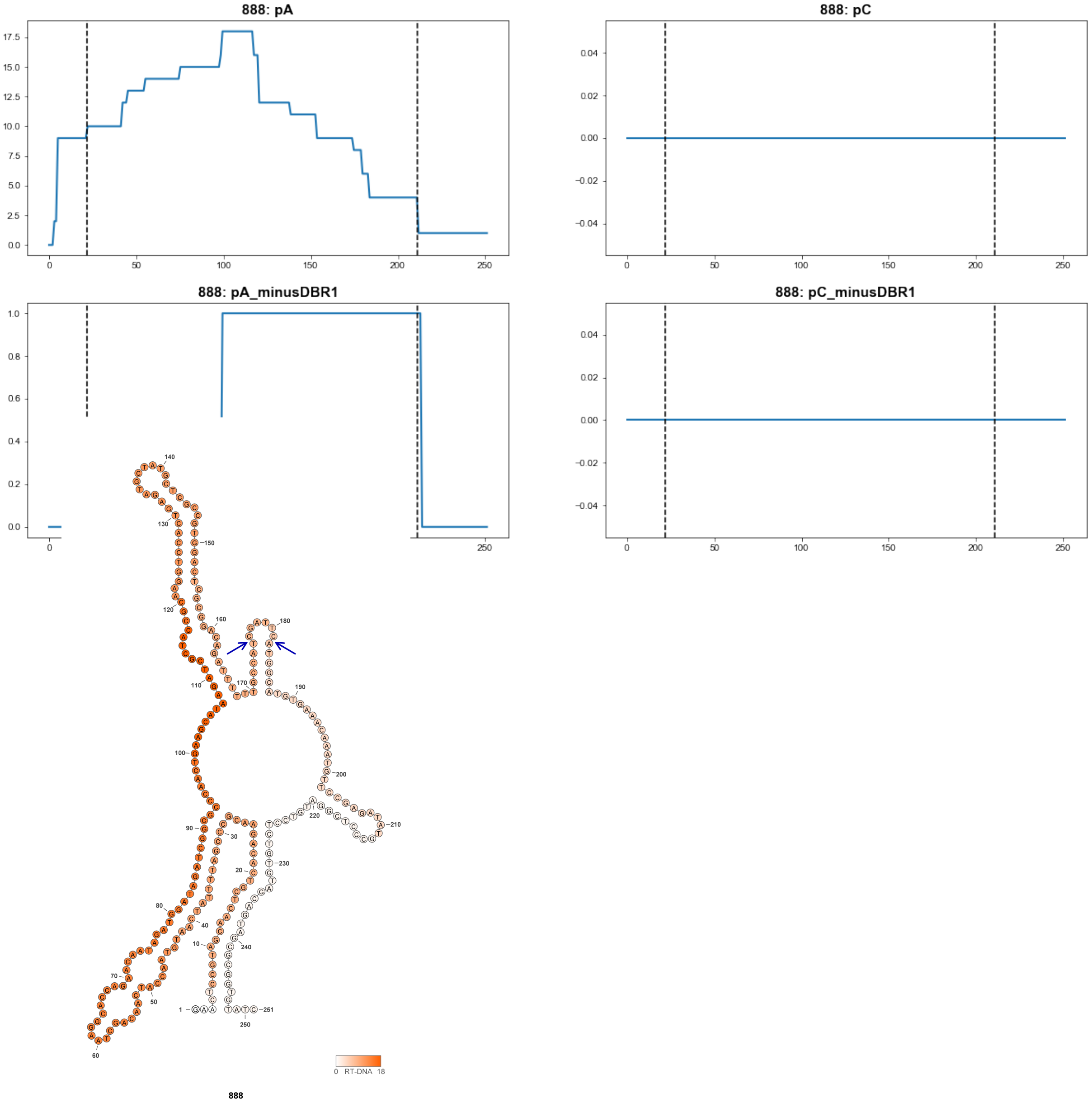

### Retron Terminal 915: Proteobacteria

**Bacterial Vector for RTDNA: pSLS.673**

**Bacterial Vector for Editing: None**

**Phage Vector for Editing: None**

**Human Vector for Editing: pSLS.844**

**RTDNA Production (relative to Eco1): 0 by PAGE**

**Bacterial Editing: undetermined**

**Phage Editing: undetermined**

**Human Editing (demultiplexed): undetermined**

**ncRNA:**

**AGGCTTGCGGATAGATTCTGCTATTGTGTTTCGCGATAGTTGATACCAGTGCGCGGGGCATTGCCC TACGCATTCCTTACGGACTGTTCACCCATGGTCCGATGACTTGGCAAACGAGTTTCCGCAAGCCT**

**RTDNA:**

**TTGCGGAAACTCGTTTGCCAAGTCATCGGACCATGGGTGAACAGTCCGTAAGGAATGCGTAGGGC AATGCCCCGCGCACTGGTATCAACTATCGCGAAAC**

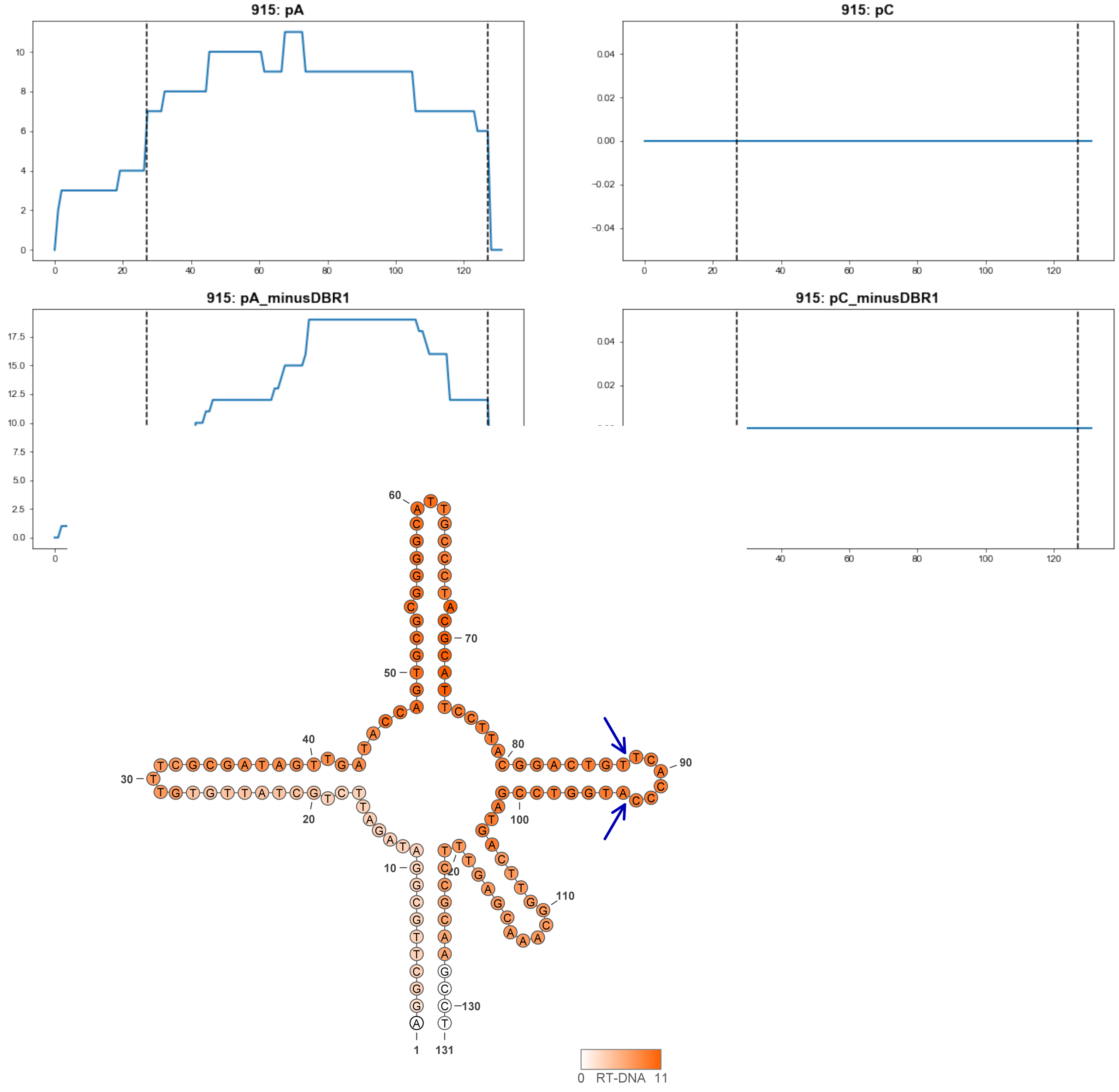


**915**

### Retron Terminal 925: Proteobacteria

**Bacterial Vector for RTDNA: None**

**Bacterial Vector for Editing: None**

**Phage Vector for Editing: None**

**Human Vector for Editing: pSLS.930**

**RTDNA Production (relative to Eco1): undetermined**

**Bacterial Editing: undetermined**

**Phage Editing: undetermined**

**Human Editing (demultiplexed): 0.161641 percent precise (1.392364611x Eco1)**

**ncRNA:**

**GGGGCTGCAGGAAGATTTGGTTGCCGTAGTTTCGCGGCATCAGAAACAATTGCAGCGTGGATTTG**

**CCTAATGGCTCGACTGCAGCGATACGCTCGCTGTGCGTGCCAATGAGTTCCTGCAGCCCC**

**no RTDNA sequencing data**

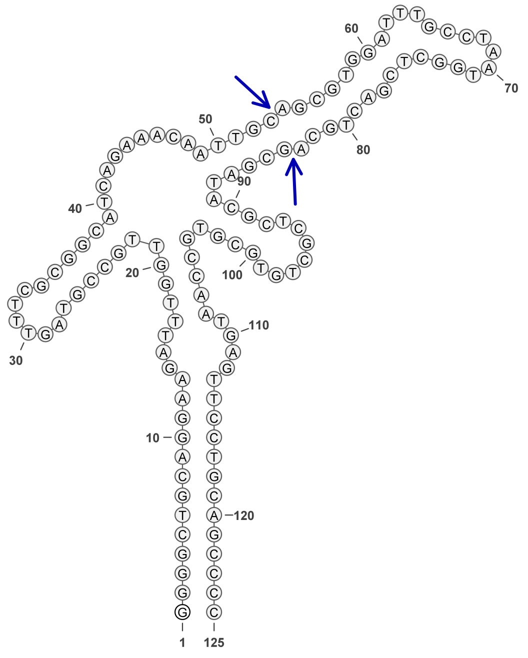

925

### Retron Terminal 939: Proteobacteria

**Bacterial Vector for RTDNA: pSLS.592**

**Bacterial Vector for Editing: None**

**Phage Vector for Editing: None**

**Human Vector for Editing: pSLS.827**

**RTDNA Production (relative to Eco1): 0 by PAGE**

**Bacterial Editing: undetermined**

**Phage Editing: undetermined**

**Human Editing (demultiplexed): 0.011486 percent precise (0.098939625x Eco1)**

**ncRNA:**

**GCTTACAGGCAGATTTATTATCGTGTTTCGCGATATATGATCCAAAGTCAGGCACGCTGCTCCCTA**
**TGATCCATGATTGAAACATATGGCCCATGGCCTATGCGACGCGTAGGAGTGCCTGTAAGC**

**RTDNA:**

**GTCGCATAGGCCATGGGCCATATGTTTCAATCATGGATCATAGGGAGCAGCGTGCCTGACTTTGG**
**ATCATATATCGCGAAACACGATAATAAATCTGCCTGTAAGC**

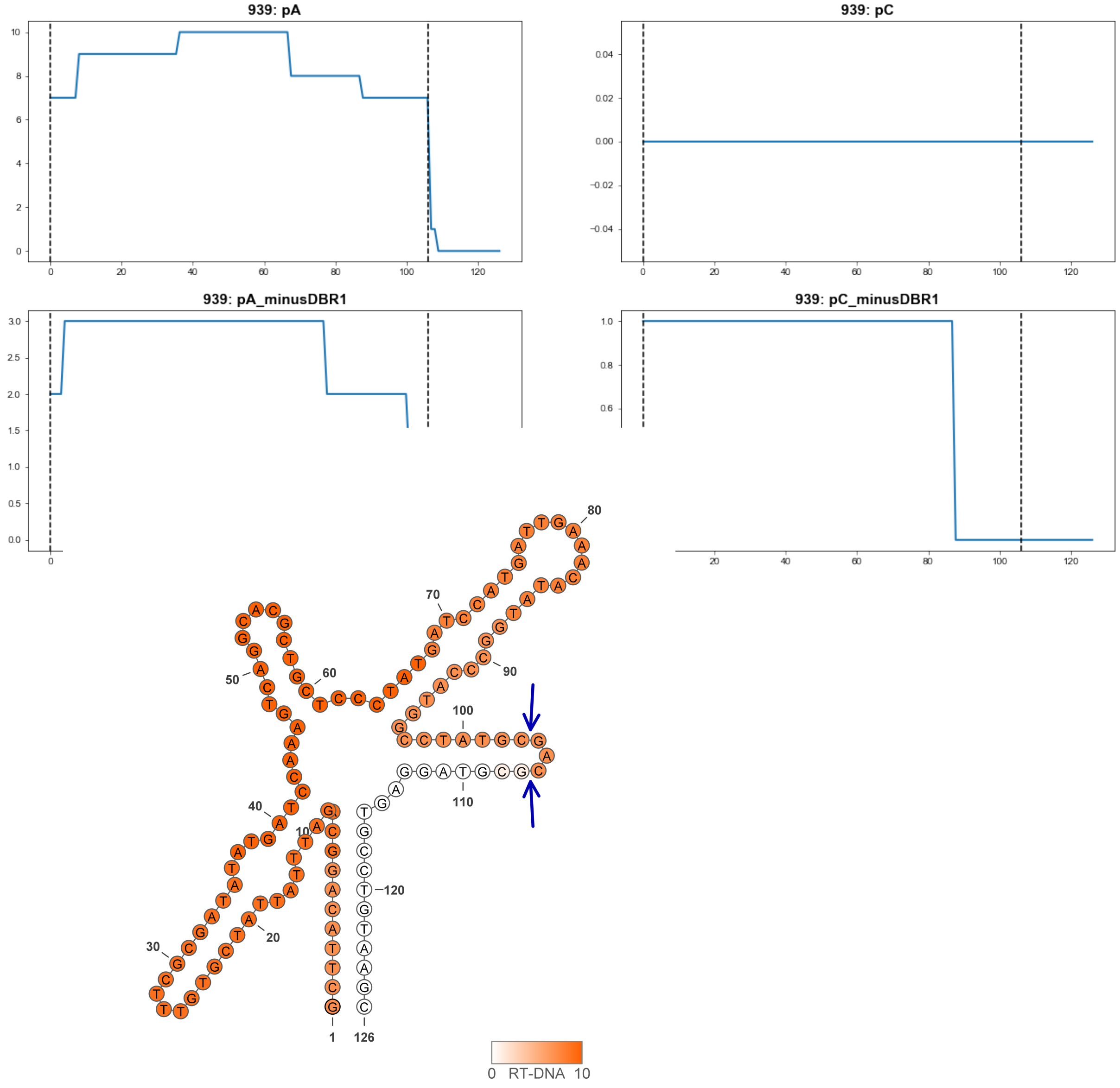


**939**

### Retron Terminal 953: Proteobacteria

**Bacterial Vector for RTDNA: pSLS.674**

**Bacterial Vector for Editing: None**

**Phage Vector for Editing: None**

**Human Vector for Editing: pSLS.909**

**RTDNA Production (relative to Eco1): 0.335754741 by PAGE**

**Bacterial Editing: undetermined**

**Phage Editing: undetermined**

**Human Editing (demultiplexed): 0 percent precise (0x Eco1)**

**ncRNA:**

**GTGCCTTACGGGCAGACTTCCGGCTGCGCATCGCGGCAAATGACAACAATGGTAGGCGTGCTGC TTACGTCTAACTTCTTTTGCGATATTTGGAGTTCTCTATCATGTATCGCTTACTTTCGCTTAAGAGTG CCCGTAAGCAC**

**RTDNA:**

**CTTAAGCGAAAGTAAGCGATACATGATAGAGAACTCCAAATATCGCAAAAGAAGTTAGACGTAAG CAGCACGCCTACC**

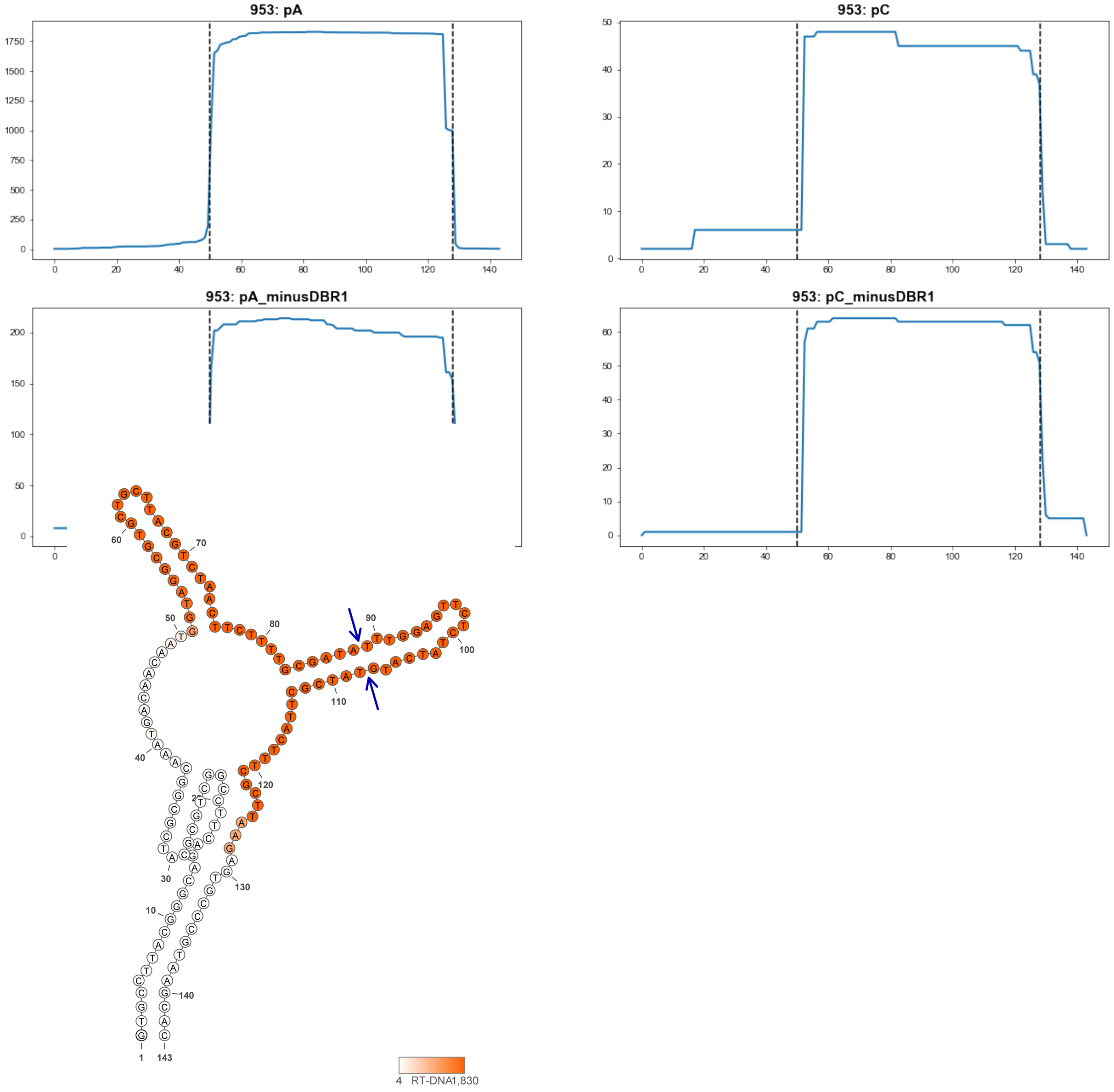


953

### Retron Terminal 962: Proteobacteria

**Bacterial Vector for RTDNA: None**

**Bacterial Vector for Editing: None**

**Phage Vector for Editing: None**

**Human Vector for Editing: pSLS.931**

**RTDNA Production (relative to Eco1): undetermined**

**Bacterial Editing: undetermined**

**Phage Editing: undetermined**

**Human Editing (demultiplexed): 0.061947 percent precise (0.533607256x Eco1)**

**ncRNA:**

**GGCTTGCAGGCAGATTCGTTGTTATGTTTCGTAACAAGTAATATAAATGGCGGGGGCATCGCCTTC**

**GCGACTCGTGATCCTGGTGACCTTGACTGTCATCCTGCGCATAGAGAGTGCCTGCAAGCT**

**no RTDNA sequencing data**

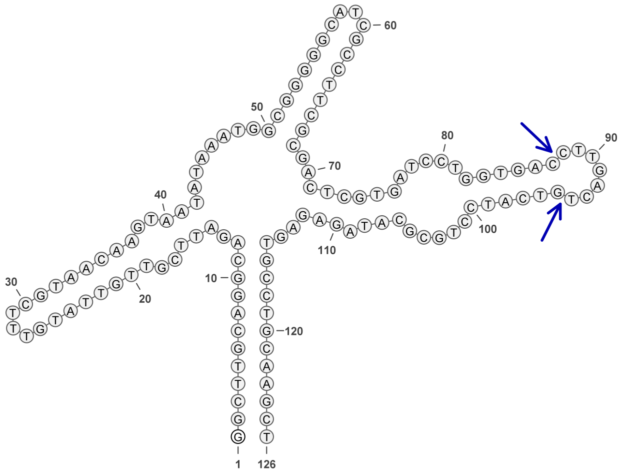


**962**

### Retron Terminal 975: Proteobacteria

**Bacterial Vector for RTDNA: pSLS.631**

**Bacterial Vector for Editing: None**

**Phage Vector for Editing: None**

**Human Vector for Editing: pSLS.856**

**RTDNA Production (relative to Eco1): 0 by PAGE**

**Bacterial Editing: undetermined**

**Phage Editing: undetermined**

**Human Editing (demultiplexed): 0.012411 percent precise (0.106907512x Eco1)**

**ncRNA:**

**GTCCTTTGGAAAGATCTTTAAATAGCGTATCGCTATTTGTTTTGAAATTGGGCTTAGGCTCTTCTTTT GTCCAGCAATACTTTGCGGTCAGTGCTTAGTCACTCTGCACGGTTGTTGACCATTGAGTTTCCAAA GGAC**

**RTDNA:**

**CAATGGTCAACAACCGTGCAGAGTGACTAAGCACTGACCGCAAAGTATTGCTGGACAAAAGAAG AGCCTA**

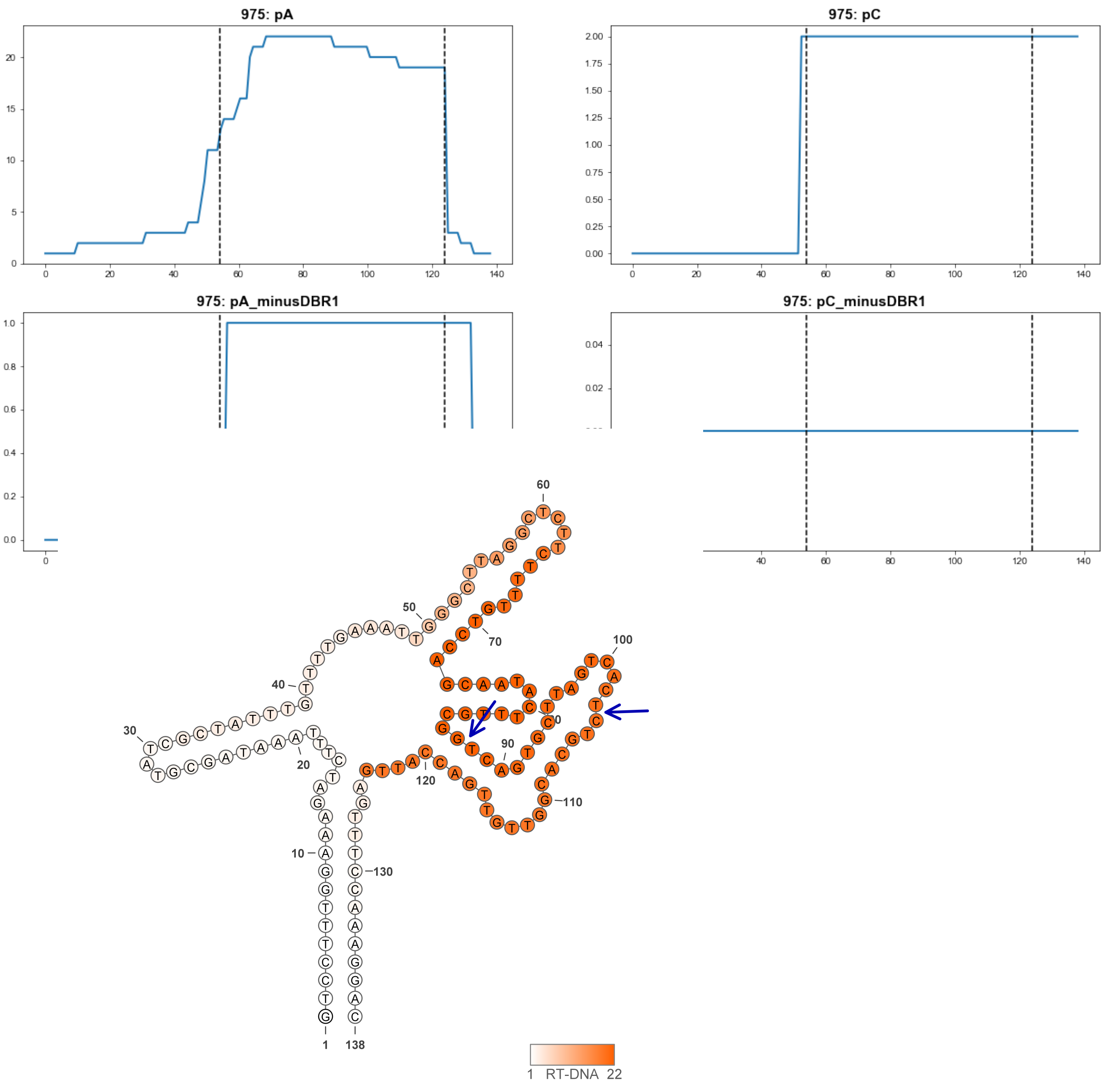


**975**

### Retron Terminal 991: Proteobacteria

**Bacterial Vector for RTDNA: pSLS.675**

**Bacterial Vector for Editing: None**

**Phage Vector for Editing: None**

**Human Vector for Editing: pSLS.910**

**RTDNA Production (relative to Eco1): 0 by PAGE**

**Bacterial Editing: undetermined**

**Phage Editing: undetermined**

**Human Editing (demultiplexed): 0.014152 percent precise (0.121904368x Eco1)**

**ncRNA:**

**GGTTTATGGGCAGATTTTGGGTCGCGTTTCGCGATCAGTGAAAAAAATGTCGTTGACATTACCTTT**

**ATGATGATTAGCTGTACTTTATGATCATTACTATTGTTAAATGTGTAGGAGTGCCCATAAAC**

**RTDNA: None**

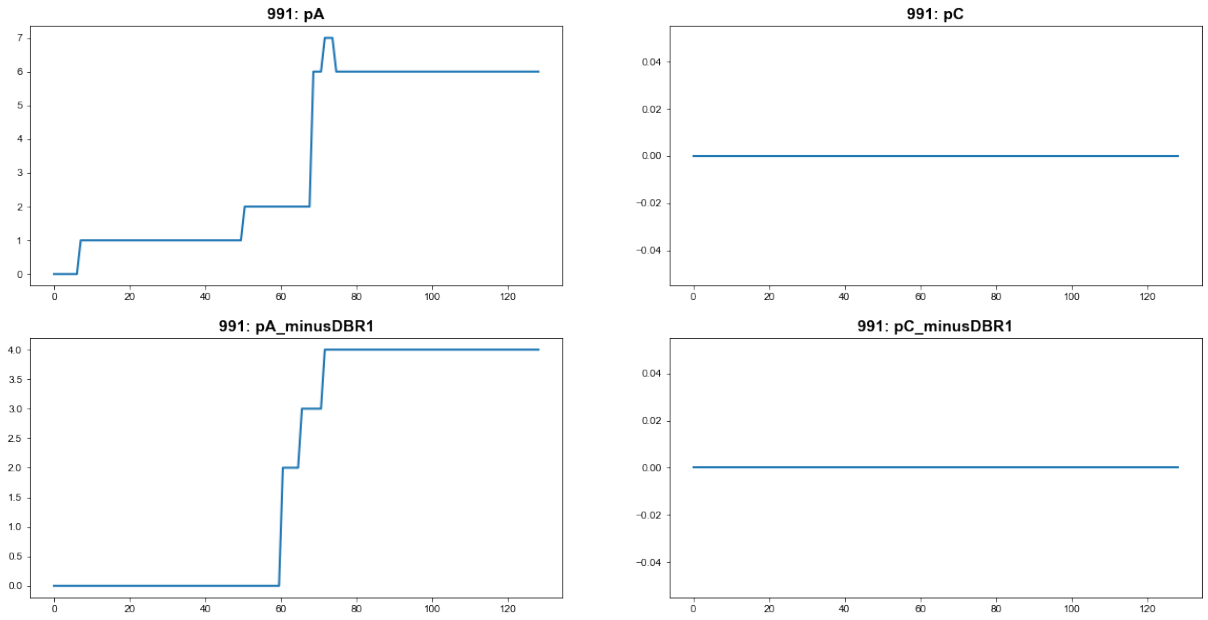


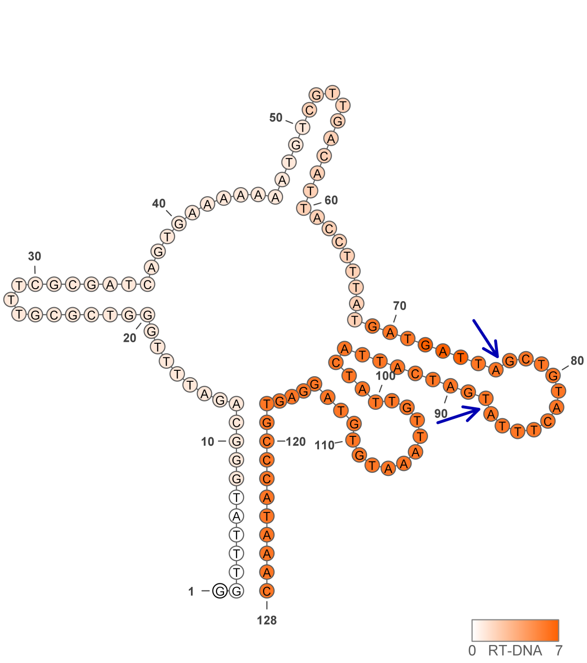


991

### Retron Terminal 1000: Proteobacteria

**Bacterial Vector for RTDNA: None**

**Bacterial Vector for Editing: None**

**Phage Vector for Editing: None**

**Human Vector for Editing: pSLS.845**

**RTDNA Production (relative to Eco1): undetermined**

**Bacterial Editing: undetermined**

**Phage Editing: undetermined**

**Human Editing (demultiplexed): 0 percent precise (0x Eco1)**

**ncRNA:**

**GATGGTTTATGGGCAGATTTTTGGTCGCGTTTCGCGACCTGTGAAAAAATTGTTCGAGAGAACTGC**
**CTATTAGTAGATTATCATCATTATGAGGCTACCTGAGAAAAGTGCCTAGGAGTGCCCATAAACCAT**
**T**

**no RTDNA sequencing data**

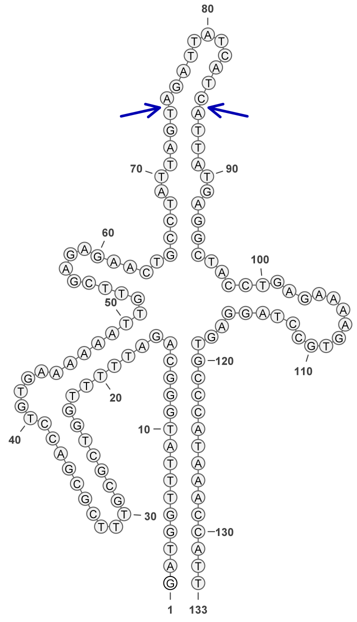

1000

### Retron Terminal 1016: Proteobacteria

**Bacterial Vector for RTDNA: pSLS.676**

**Bacterial Vector for Editing: None**

**Phage Vector for Editing: None**

**Human Vector for Editing: pSLS.911**

**RTDNA Production (relative to Eco1): 0 by PAGE**

**Bacterial Editing: undetermined**

**Phage Editing: undetermined**

**Human Editing (demultiplexed): 0.16609 percent precise (1.430687995x Eco1)**

**ncRNA:**

**TGGCTTATGGGCAGATTTTGGATTGTGATATCGCAATCAGTGAAAAAATTGTCCAATGGACTCGCC CCACGGGTACGTTAATTTAGCGGATCTGTGTAACTTAAGAATGCGTAAGAGTGCCCATAAGCCA**

**RTDNA:**

**CTTACGCATTCTTAAGTTACACAGATCCGCTAAATTAACGTACCCGTGGGGCGAGTCCATTGGACA**

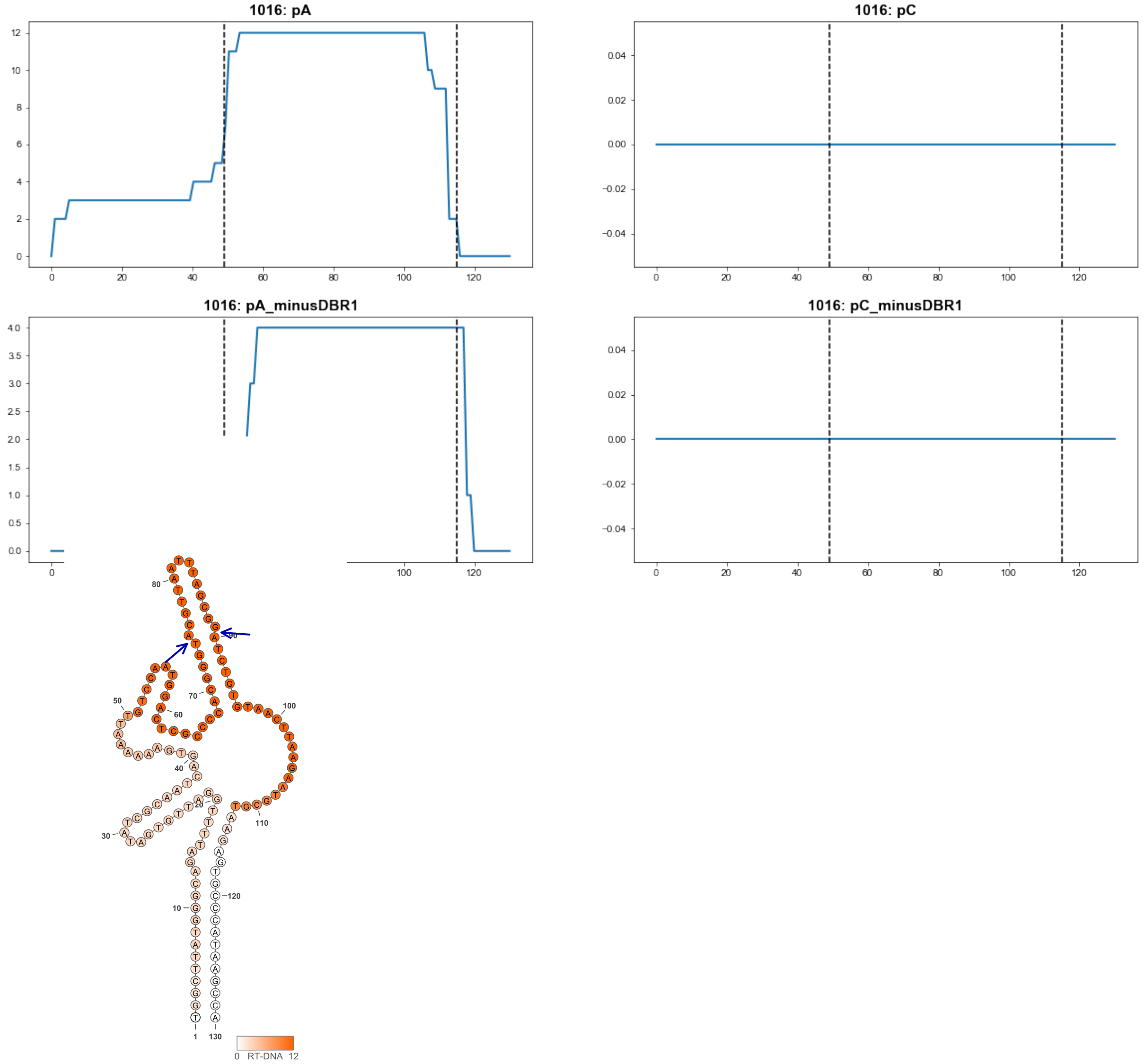


1016

### Retron Terminal 1034: Proteobacteria

**Bacterial Vector for RTDNA: None**

**Bacterial Vector for Editing: None**

**Phage Vector for Editing: None**

**Human Vector for Editing: pSLS.932**

**RTDNA Production (relative to Eco1): undetermined**

**Bacterial Editing: undetermined**

**Phage Editing: undetermined**

**Human Editing (demultiplexed): 0.057434 percent precise (0.49473258x Eco1)**

**ncRNA:**

**CTAAATAGCTTCGCACATTGATGCTTCACTACGCATCGAACGAGGCCAACGCGGACGGAAGGGC**

**GCGCACTTCCAAATGTAGTGCTATTACGTGAAGCTATTTAG**

**no RTDNA sequencing data**

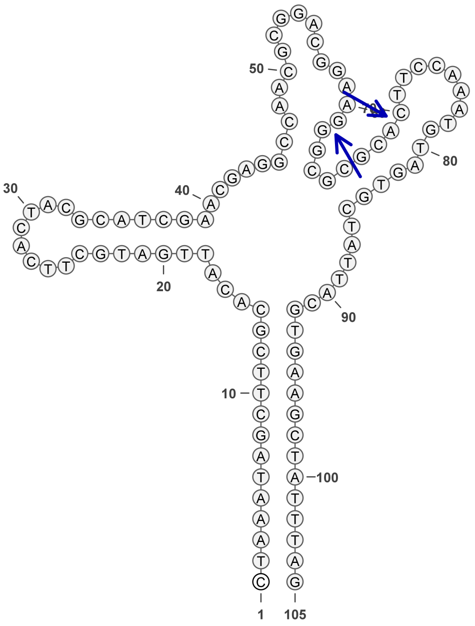


**1034**

### Retron Terminal 1036: Proteobacteria

**Bacterial Vector for RTDNA: pCF.6**

**Bacterial Vector for Editing: pAGD148**

**Phage Vector for Editing: pAGD400**

**Human Vector for Editing: None**

**RTDNA Production (relative to Eco1): undetermined**

**Bacterial Editing: 2.89469787 percent precise (0.890734815x Eco1)**

**Phage Editing: 8.72477538 percent precise (0.215193725x Eco1)**

**Human Editing (demultiplexed): undetermined**

**ncRNA:**

**ACGCATGTAGGCAGATTTGTTGGTTGTGAATCGCAACCAGTGGCCTTAATGGCAGGAGGAATCGC
CTCCCTAAAATCCTTGATTCAGAGCTATACGGCAGGTGTGCTGTGCGAAGGAGTGCCTGCATGCG
T**

**RTDNA:**

**TCCTTCGCACAGCACACCTGCCGTATAGCTCTGAATCAAGGATTTTAGGGAGGCGATTCCTCCTGC
C**

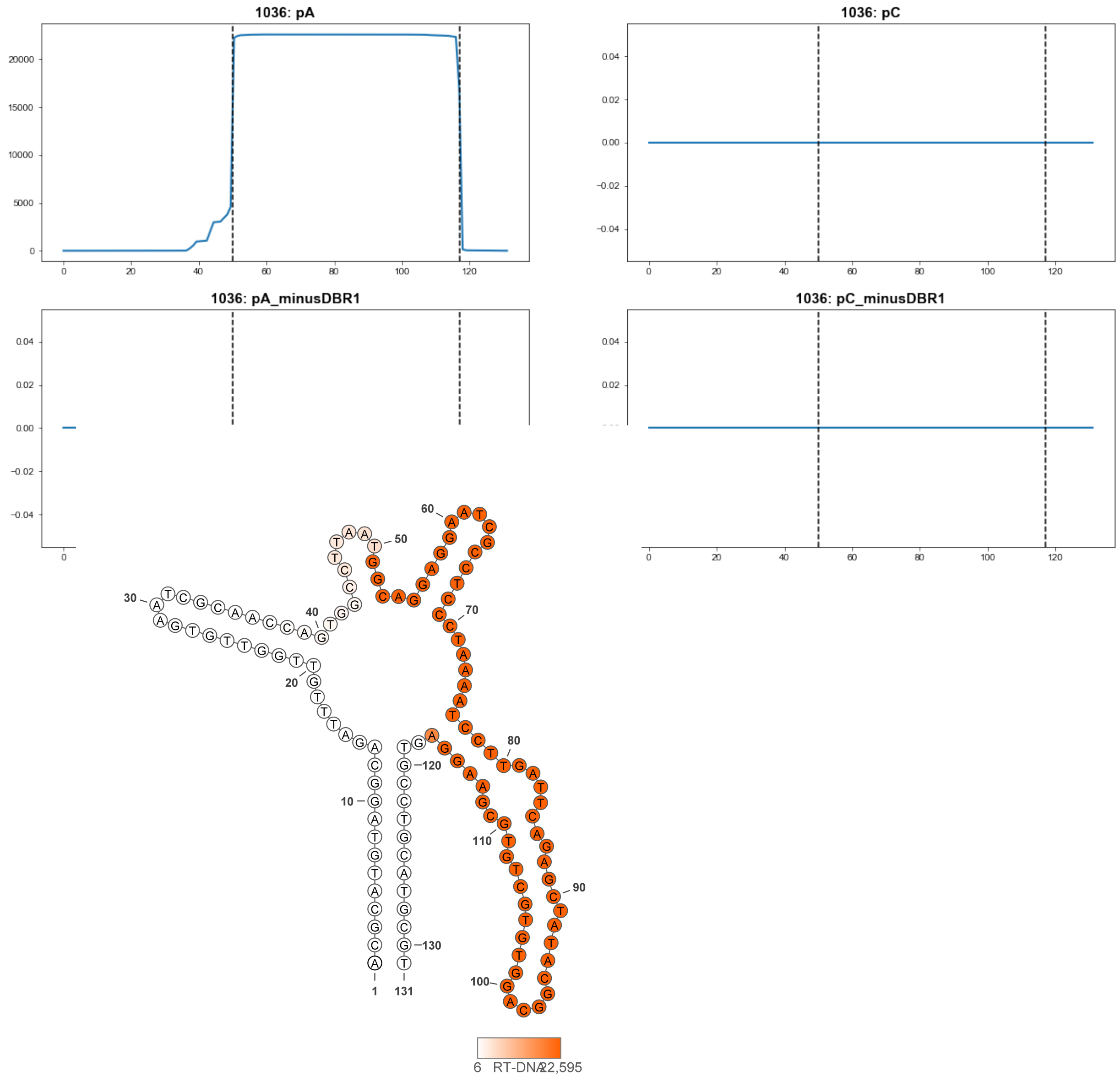


**1036**

### Retron Terminal 1049: Proteobacteria

**Bacterial Vector for RTDNA: pSLS.677**

**Bacterial Vector for Editing: None**

**Phage Vector for Editing: None**

**Human Vector for Editing: None**

**RTDNA Production (relative to Eco1): 0.837832433 by PAGE**

**Bacterial Editing: undetermined**

**Phage Editing: undetermined**

**Human Editing (demultiplexed): undetermined**

**ncRNA:**

**AGGGCTATAGGCAGATTTTTTGTCACGAAGCGTGATATATGAAAAGAATGTCCGGTGGAATGCCG ATCAAAGTGTTAAGTTACTTGATGATTAGATTGCAAATGAGCTTAATGAGTGCCTATAGACCC**

**RTDNA:**

**TCATTAAGCTCATTTGCAATCTAATCATCAAGTAACTTAACACTTTGATCGGCATTCCACCGGACAT TCTTT**

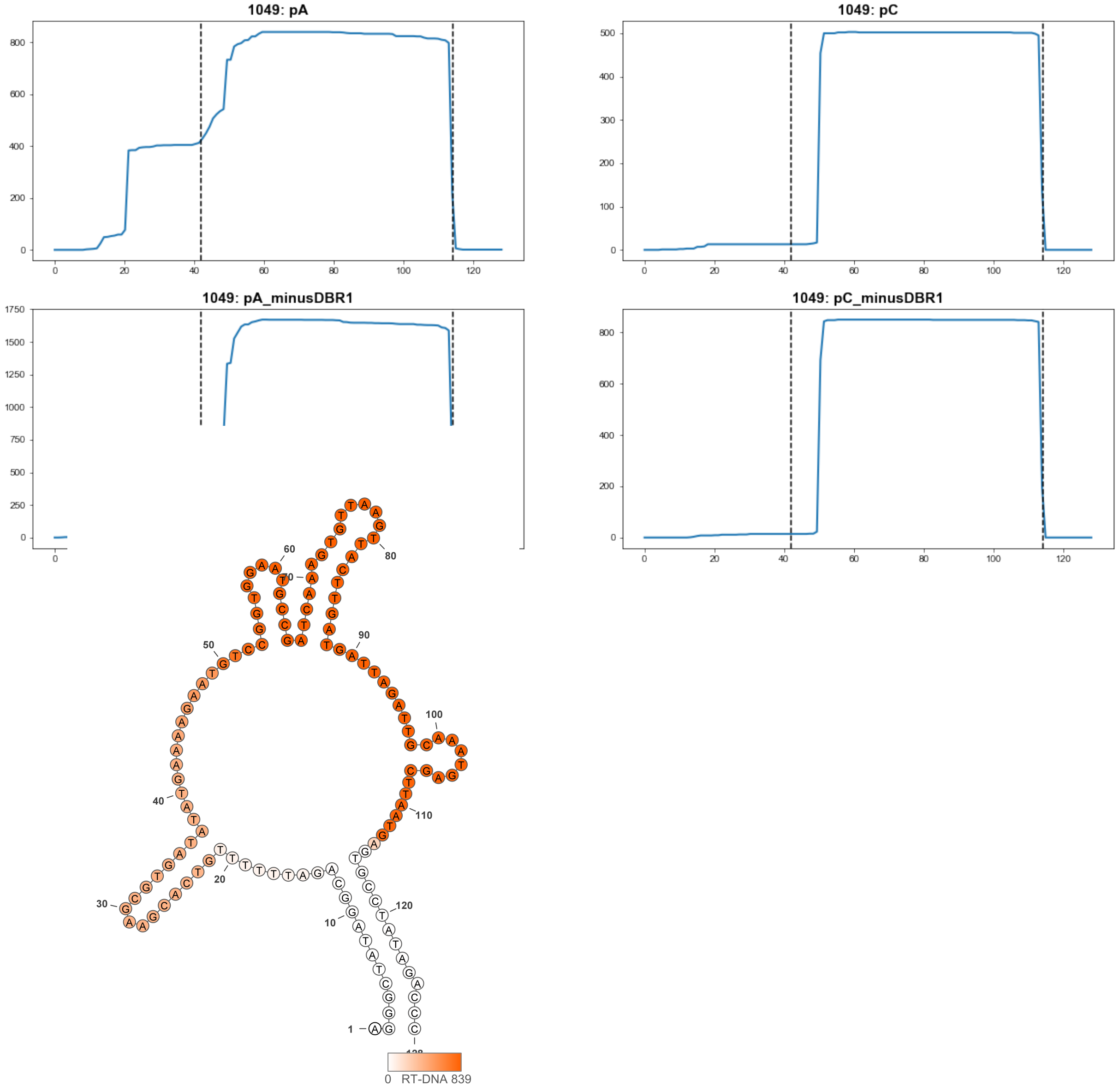


1049

### Retron Terminal 1054: Firmicutes

**Bacterial Vector for RTDNA: pSLS.678**

**Bacterial Vector for Editing: None**

**Phage Vector for Editing: None**

**Human Vector for Editing: pSLS.912**

**RTDNA Production (relative to Eco1): 0 by PAGE**

**Bacterial Editing: undetermined**

**Phage Editing: undetermined**

**Human Editing (demultiplexed): 0.007967 percent precise (0.068627198x Eco1)**

**ncRNA:**

**AACGTGTGGCTAGATTATTCGCTTGTGTTTCGCAAGTGTTGACCAAAATGGTAGGAGCATTAGCTT**
**CTGCTATTCGTATTACTATTGGTGTATTTACGTTAAATTTTCAATGTTTCTGCATCGCATAATGAGTA**
**GCCACACGTT**

**RTDNA: None**

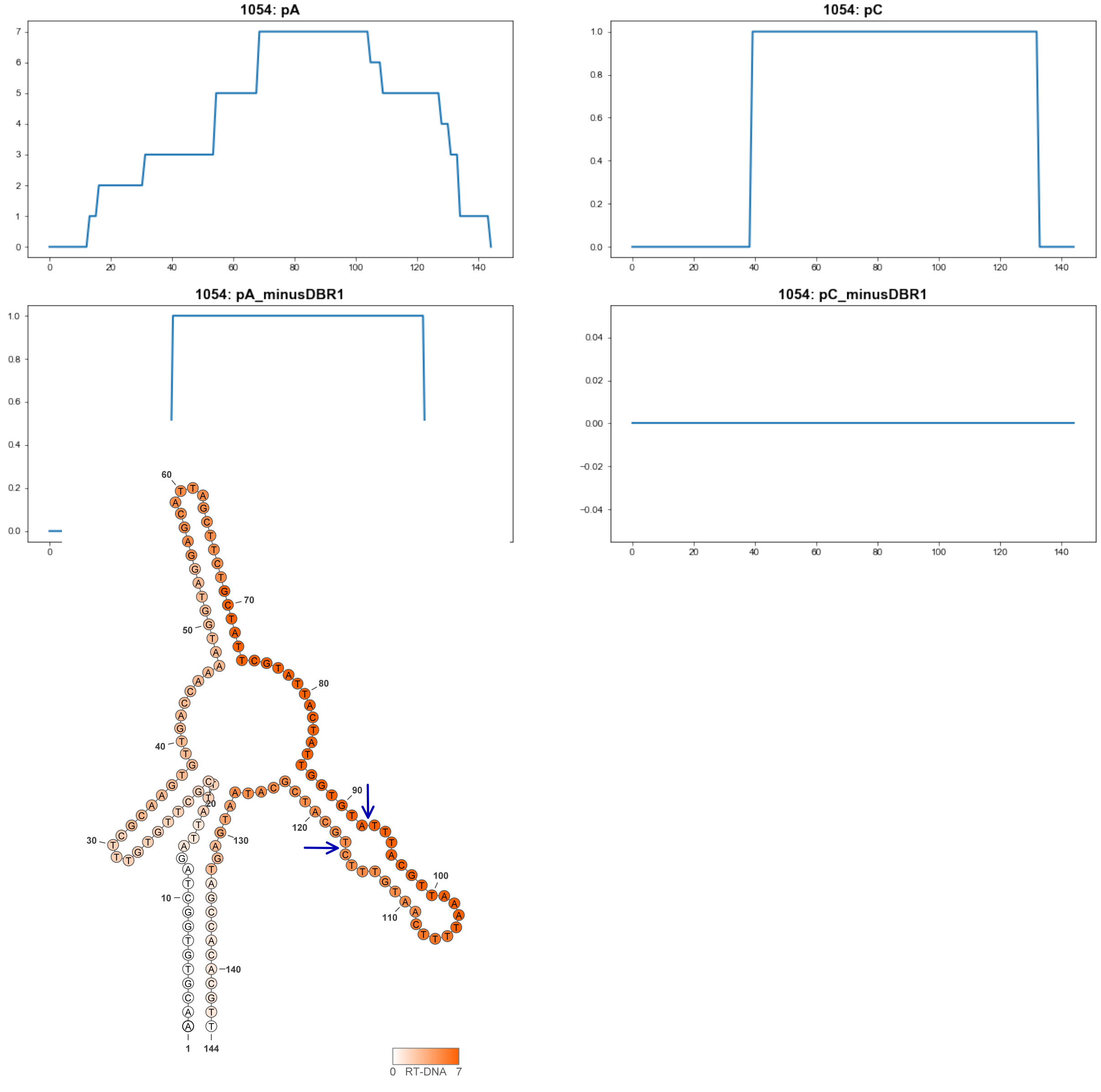


**1054**

### Retron Terminal 1079: Firmicutes

**Bacterial Vector for RTDNA: None**

**Bacterial Vector for Editing: None**

**Phage Vector for Editing: None**

**Human Vector for Editing: pSLS.933**

**RTDNA Production (relative to Eco1): undetermined**

**Bacterial Editing: undetermined**

**Phage Editing: undetermined**

**Human Editing (demultiplexed): 0 percent precise (0x Eco1)**

**ncRNA:**

**GCGTGTGAATAGATTTTTGACTTGTGTGTTCGCAAGTATTAACAATAATGTTACAAGAGCGTTTTGC**

**TCCCTCCAGTAGTCAATAATGTTCAATTCGACAATTGCTGCTGATGAATGATAAACTACATCAATA**

**GGCTGAACTAATTGGCTATATCTGTTTTACTGTCGTTCGCTTGAGAGTTTTCACACGC**

**no RTDNA sequencing data**

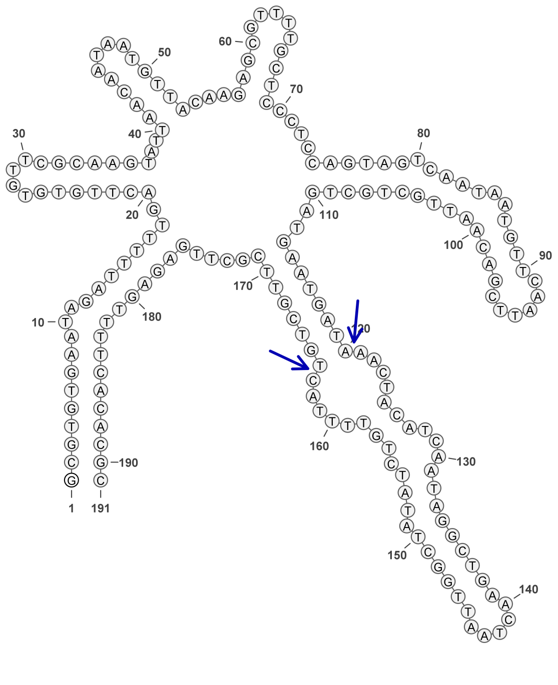


**1079**

### Retron Terminal 1100: Firmicutes

**Bacterial Vector for RTDNA: pSLS.616**

**Bacterial Vector for Editing: None**

**Phage Vector for Editing: None**

**Human Vector for Editing: pSLS.837**

**RTDNA Production (relative to Eco1): 0.414806916 by PAGE**

**Bacterial Editing: undetermined**

**Phage Editing: undetermined**

**Human Editing (demultiplexed): 0 percent precise (0x Eco1)**

**ncRNA:**

**TTGTGCACGGTTAGATTTTTTACTCATGTTTCGTGAGTATTGACAAGAACGGTACGATATGTGCTCT
TTACGCGTCTCCCGAACACTCACGCTATACTTTTGGGAGATTACGGGTTACACTGCCTAAAGAGTA
ACCGTGCACAA**

**RTDNA:**

**CTTTAGGCAGTGTAACCCGTAATCTCCCAAAAGTATAGCGTGAGTGTTCGGGAGACGCGTAAAGA
GCACATATCGTACC**

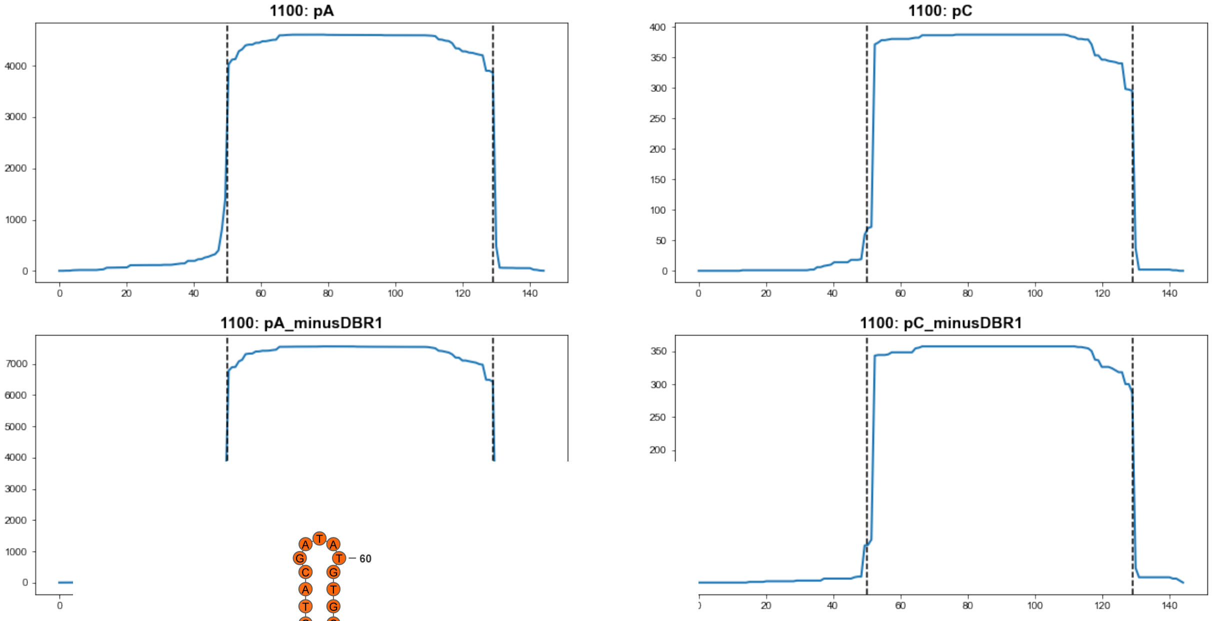


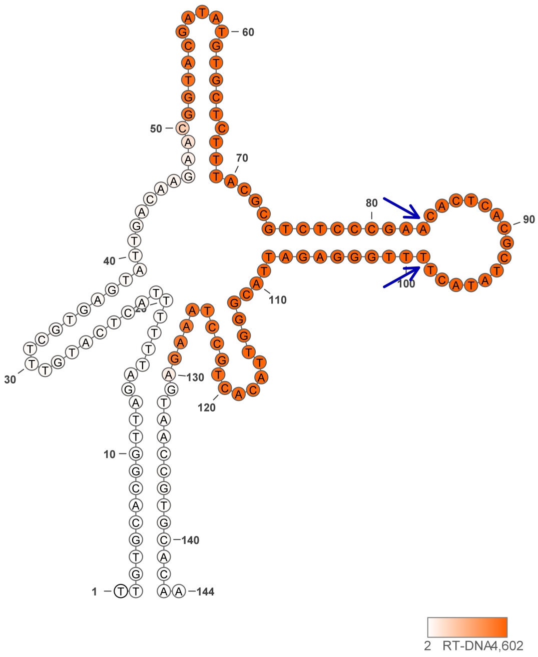


1100

### Retron Terminal 1119: Firmicutes

**Bacterial Vector for RTDNA: pSLS.679**

**Bacterial Vector for Editing: None**

**Phage Vector for Editing: None**

**Human Vector for Editing: pSLS.913**

**RTDNA Production (relative to Eco1): 0 by PAGE**

**Bacterial Editing: undetermined**

**Phage Editing: undetermined**

**Human Editing (demultiplexed): 0 percent precise (0x Eco1)**

**ncRNA:**

**ACGTATGATTAGATTTTGACTTATGTTTCGTAAGTTTTTATGTTAAGGGTAGGAGTTTTTGCCTTTCT ATTGCTTAGCCGAAAAAGTAGTCAGTTTAGAATCTGCTGAAGGTTAAGCAGACCGCGTCGGCGTA ATGAGTTATCATACGT**

**RTDNA:**

**TAACCTTCAGCAGATTCTAAACTGACTACTTTTTCGGCTAAGCAATAGAAAGGCAAAAACTCCTAC CCTTAACATAAAAACTTACGAAACATAAGTC**

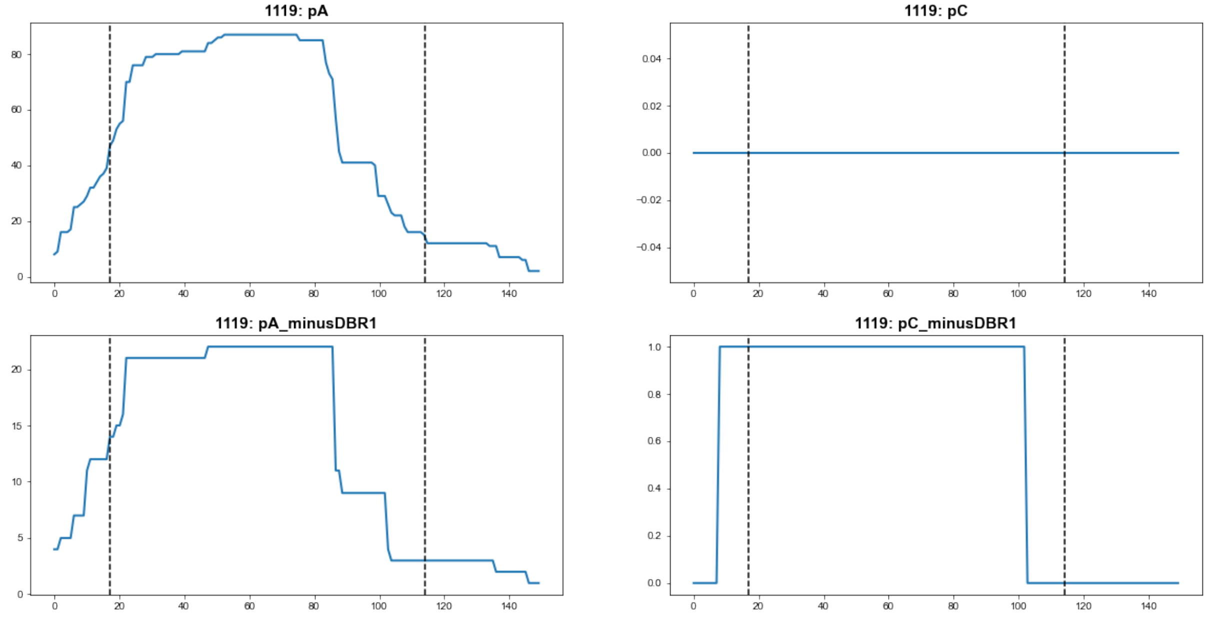


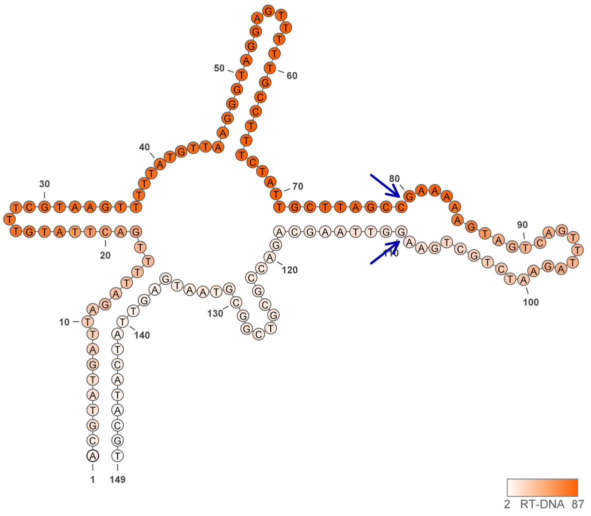


**1119**

### Retron Terminal 1122: Proteobacteria

**Bacterial Vector for RTDNA: pSLS.632**

**Bacterial Vector for Editing: None**

**Phage Vector for Editing: None**

**Human Vector for Editing: pSLS.857**

**RTDNA Production (relative to Eco1): 0.021555336 by PAGE**

**Bacterial Editing: undetermined**

**Phage Editing: undetermined**

**Human Editing (demultiplexed): 0 percent precise (0x Eco1)**

**ncRNA:**

**ACGGGGTCTCCCCTCGCATACGCGCTGTAGAAACTGCGGCGTAAGAGTCAAAGTCTGCACCTAT**

**CGTGCCCGCCGAATGCCCTTGGGTGCACCGGCGTGATTGGAAGGGGACCCCGT**

**RTDNA: CAATCACGCCGGTGCACCCAAGGGCATTCGGCGGGCACGATAGGTGCAGACTTTG**

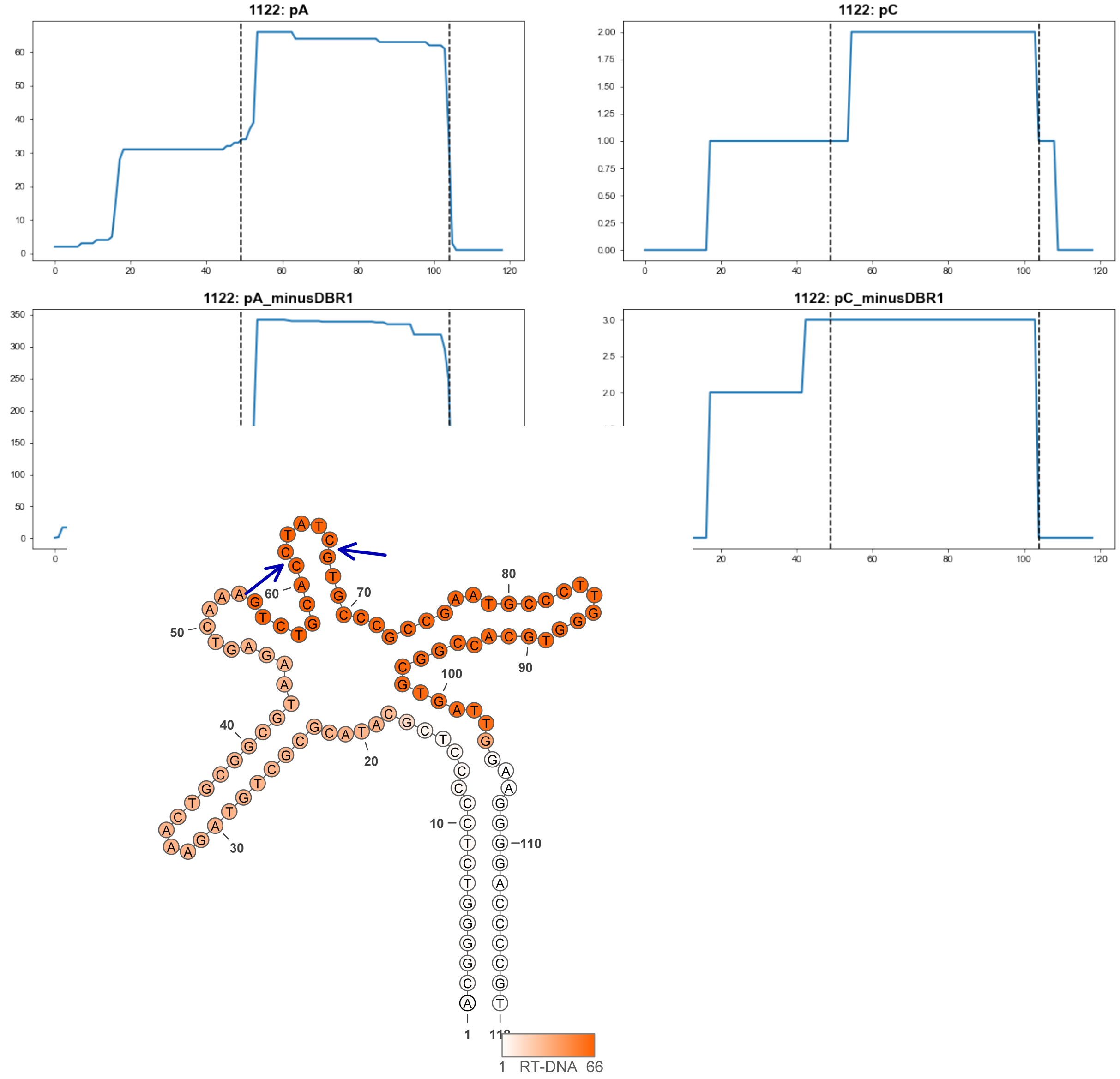


**1122**

### Retron Terminal 1143: Nitrospirae

**Bacterial Vector for RTDNA: pSLS.680**

**Bacterial Vector for Editing: None**

**Phage Vector for Editing: None**

**Human Vector for Editing: pSLS.914**

**RTDNA Production (relative to Eco1): 0.7864295 by PAGE**

**Bacterial Editing: undetermined**

**Phage Editing: undetermined**

**Human Editing (demultiplexed): 0.108618 percent precise (0.935628085x Eco1)**

**ncRNA:**

**GGGCTTGCGGGATGTATTGAAGTCTGTTGTTGATCCTGCGACAGGTCAAATCAAGGGGCCATATA TGCCTCTTGTGCTTTACGTGTATAGCTTACAGTGTTTACCTCTTGTTCTTGGCAGAAGGCGGAATAC CCGCAAGCCC**

**RTDNA:**

**CGCCTTCTGCCAAGAACAAGAGGTAAACACTGTAAGCTATACACGTAAAGCACAAGAGGCATATA TGGCCC**

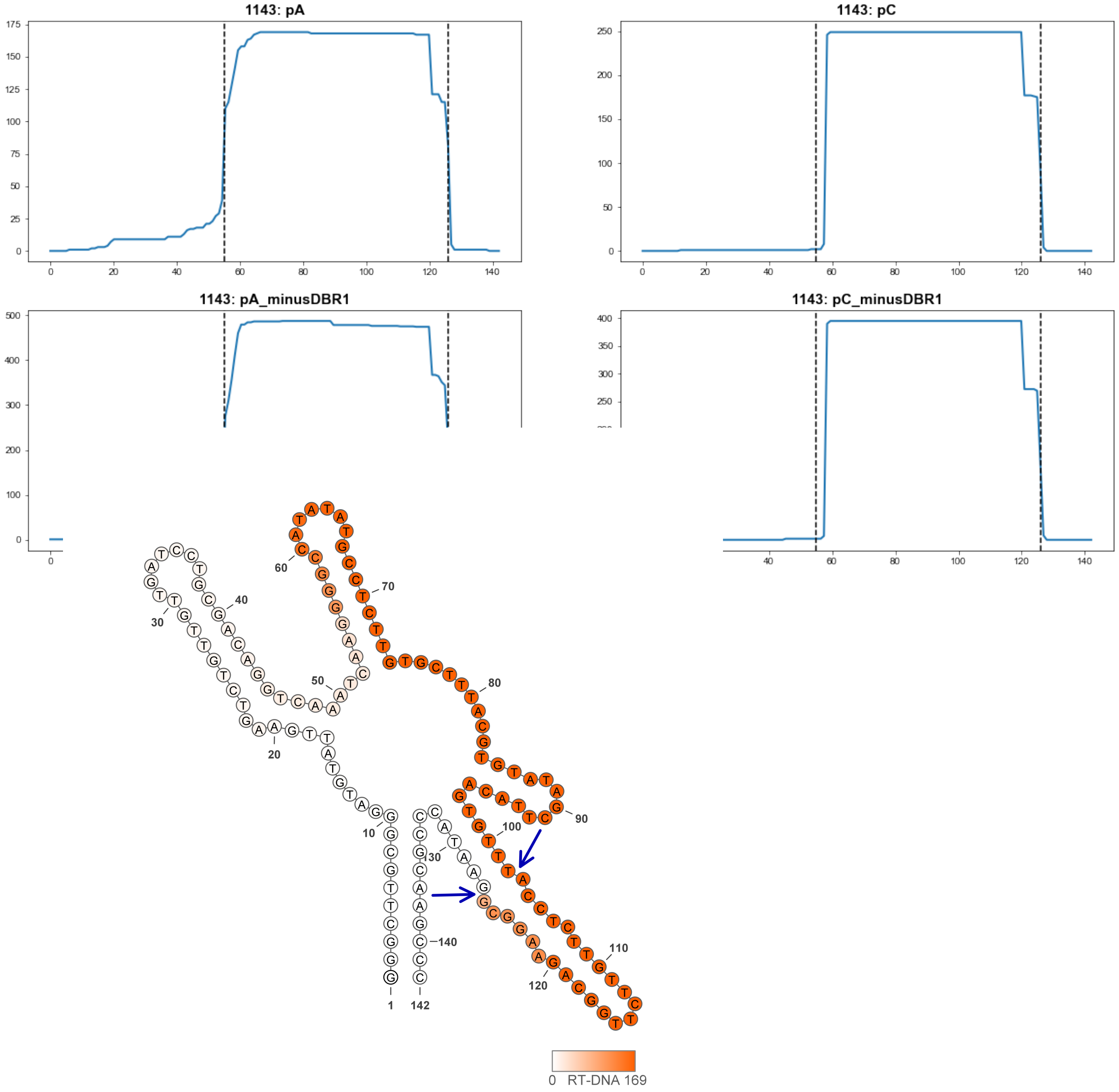


**1143**

### Retron Terminal 1156: Proteobacteria

**Bacterial Vector for RTDNA: pSLS.681**

**Bacterial Vector for Editing: None**

**Phage Vector for Editing: None**

**Human Vector for Editing: pSLS.875**

**RTDNA Production (relative to Eco1): 0.024610922 by PAGE**

**Bacterial Editing: undetermined**

**Phage Editing: undetermined**

**Human Editing (demultiplexed): 0.408723 percent precise (3.5207122x Eco1)**

**ncRNA:**

**TACGGCTCTGCCTTAAATCTGTGAGGTTGTTTCGCCTCGAAGTATCTTATGTTAGTACATCACGCTA CCAATCAGCGGTTAGTTACTTGACGTAACTGTTAATTGGCTAAAGTTTGCATAAAGTGATTGGGCG GAGCCGTA**

**RTDNA:**

**CACTTTATGCAAACTTTAGCCAATTAACAGTTACGTCAAGTAACTAACCGCTGATTGGTAGCGTGA TGTACTAAC**

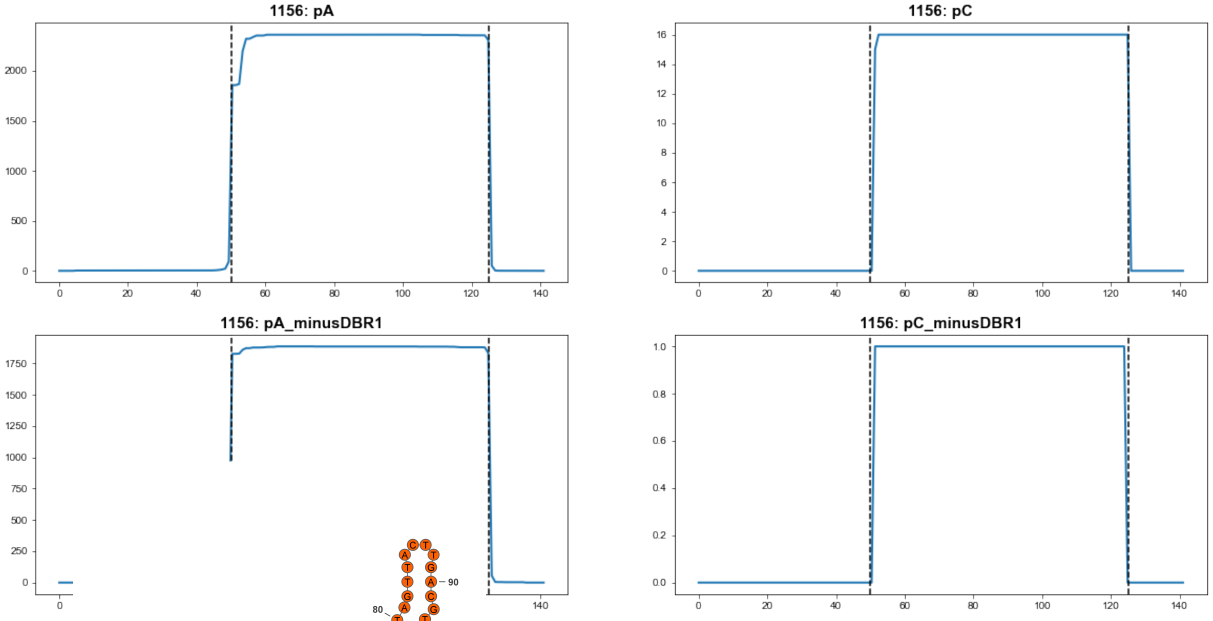


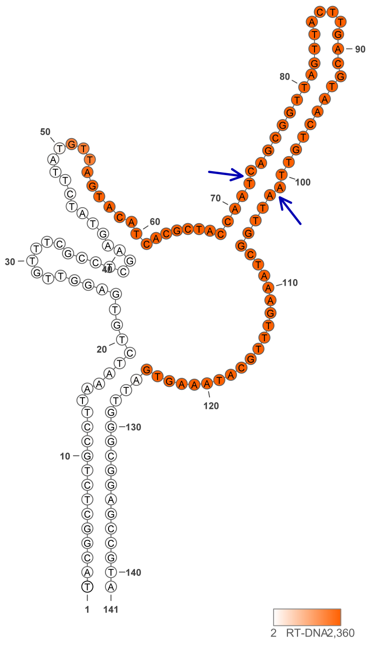


1156

### Retron Terminal 1163: Firmicutes

**Bacterial Vector for RTDNA: pSLS.593**

**Bacterial Vector for Editing: pAGD291**

**Phage Vector for Editing: pAGD401**

**Human Vector for Editing: pSLS.828**

**RTDNA Production (relative to Eco1): 0.26216772 by PAGE**

**Bacterial Editing: 1.13304372 percent precise (0.348651754x Eco1)**

**Phage Editing: 8.175623125 percent precise (0.201649064x Eco1)**

**Human Editing (demultiplexed): 0.055155 percent precise (0.475101429x Eco1)**

**ncRNA:**

**GAAGATATTCAGGGTCTCTGTTTTTCGTTAGCGTTACGCAACGGAGTTTTTAATGTGTGAGGCATAT GCTGCGTGATCGCTTTTGCGTTGTCCTCGTATACACGCCTTTATTATTCTGTATATGTCGCAGAAGG TAGAGTTTCCCTGTATATCTTC**

**RTDNA:**

**TACCTTCTGCGACATATACAGAATAATAAAGGCGTGTATACGAGGACAACGCAAAAGCGATCACG CAGCATATGCCTCACAC**

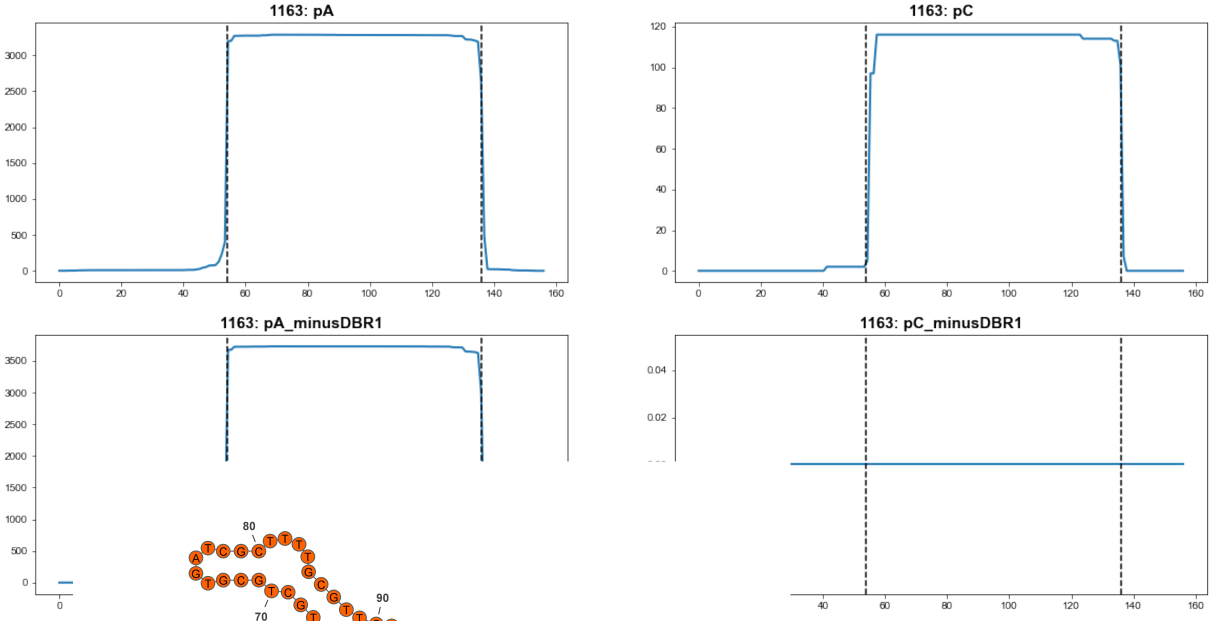


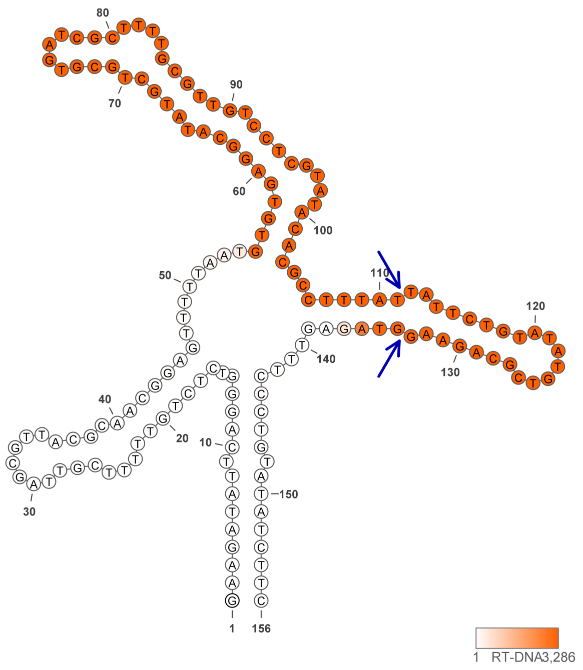


**1163**

### Retron Terminal 1165: Firmicutes

**Bacterial Vector for RTDNA: None**

**Bacterial Vector for Editing: None**

**Phage Vector for Editing: None**

**Human Vector for Editing: pSLS.876**

**RTDNA Production (relative to Eco1): undetermined**

**Bacterial Editing: undetermined**

**Phage Editing: undetermined**

**Human Editing (demultiplexed): 0 percent precise (0x Eco1)**

**ncRNA:**

**AAAATGGGTGGATTGTTTAGCGTGAGCGTTACAGCAACGATGGAATTAAATGTGAGAGGGATGTC**
**CGCTGGCTCTTTACATCACATTTACTGTTATTGATATATTGTATTAGATTATTGAAAATAATAATATA**
**TGGAATAGGTAGAATCCACCCATTTT**

**no RTDNA sequencing data**

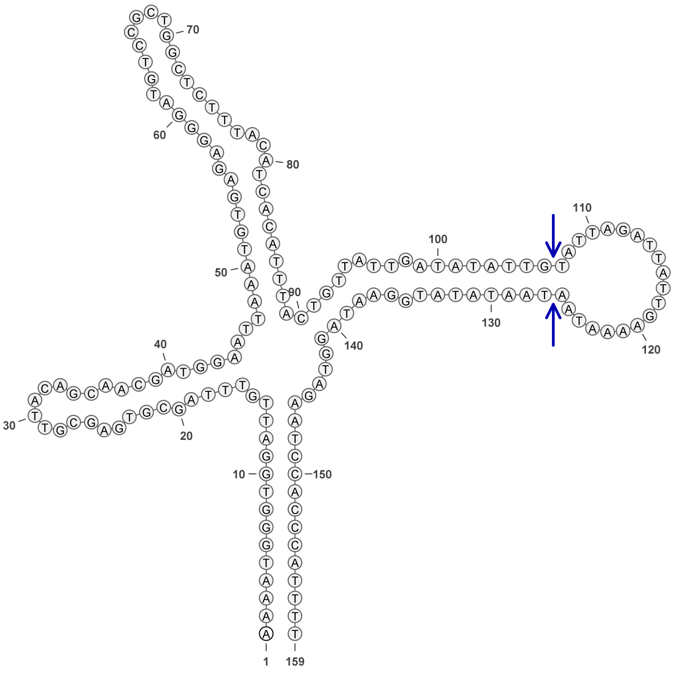


**1165**

### Retron Terminal 1168: Firmicutes

**Bacterial Vector for RTDNA: None**

**Bacterial Vector for Editing: None**

**Phage Vector for Editing: None**

**Human Vector for Editing: pSLS.877**

**RTDNA Production (relative to Eco1): undetermined**

**Bacterial Editing: undetermined**

**Phage Editing: undetermined**

**Human Editing (demultiplexed): 0.003049 percent precise (0.026263879x Eco1)**

**ncRNA:**

**GCAAACAAGTTGCATTTGAACTTTTGATGGTGATAATCGCTATCATATATTTTAATGTATCTATACA
TCTCTCTGCACTATTCTCATAGTTGATCCAATTGTCTCATATGGTATCTACGCGATGATTTAGAAAG
GTAGAAATGCACTTGTTTGT**

**no RTDNA sequencing data**

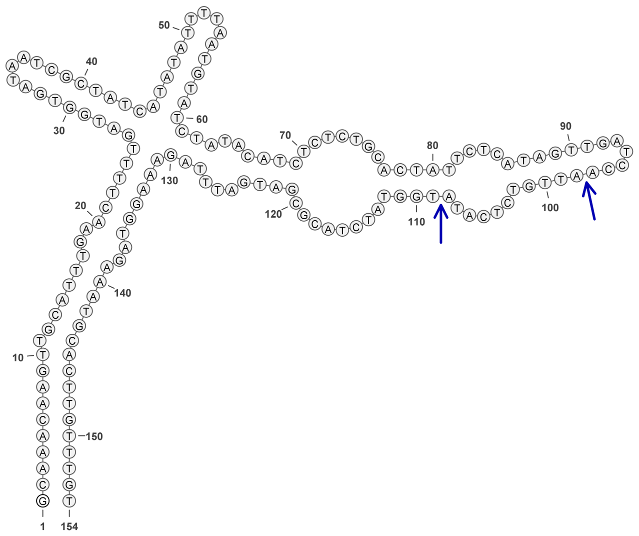

**1168**

### Retron Terminal 1184: Bacteroidetes

**Bacterial Vector for RTDNA: pSLS.682**

**Bacterial Vector for Editing: None**

**Phage Vector for Editing: None**

**Human Vector for Editing: pSLS.916**

**RTDNA Production (relative to Eco1): 0 by PAGE**

**Bacterial Editing: undetermined**

**Phage Editing: undetermined**

**Human Editing (demultiplexed): 0 percent precise (0x Eco1)**

**ncRNA:**

**GATATGCATCTGCTGAATGATTGATTTTTCGCTGTATTCTTACGGCGATTATGACAATGTCAGAACA
TTGTTCGTTCCCGGTTTTCGACTGTGTGCTTTCGATGTCGTTTGACACTTTAAGTAGAATATTCAGC
AGATGCTGTT**

**RTDNA:**

**ATATTCTACTTAAAGTGTCAAACGACATCGAAAGCACACAGTCGAAAACCGGGAACGAACAATGT
TCTGAC**

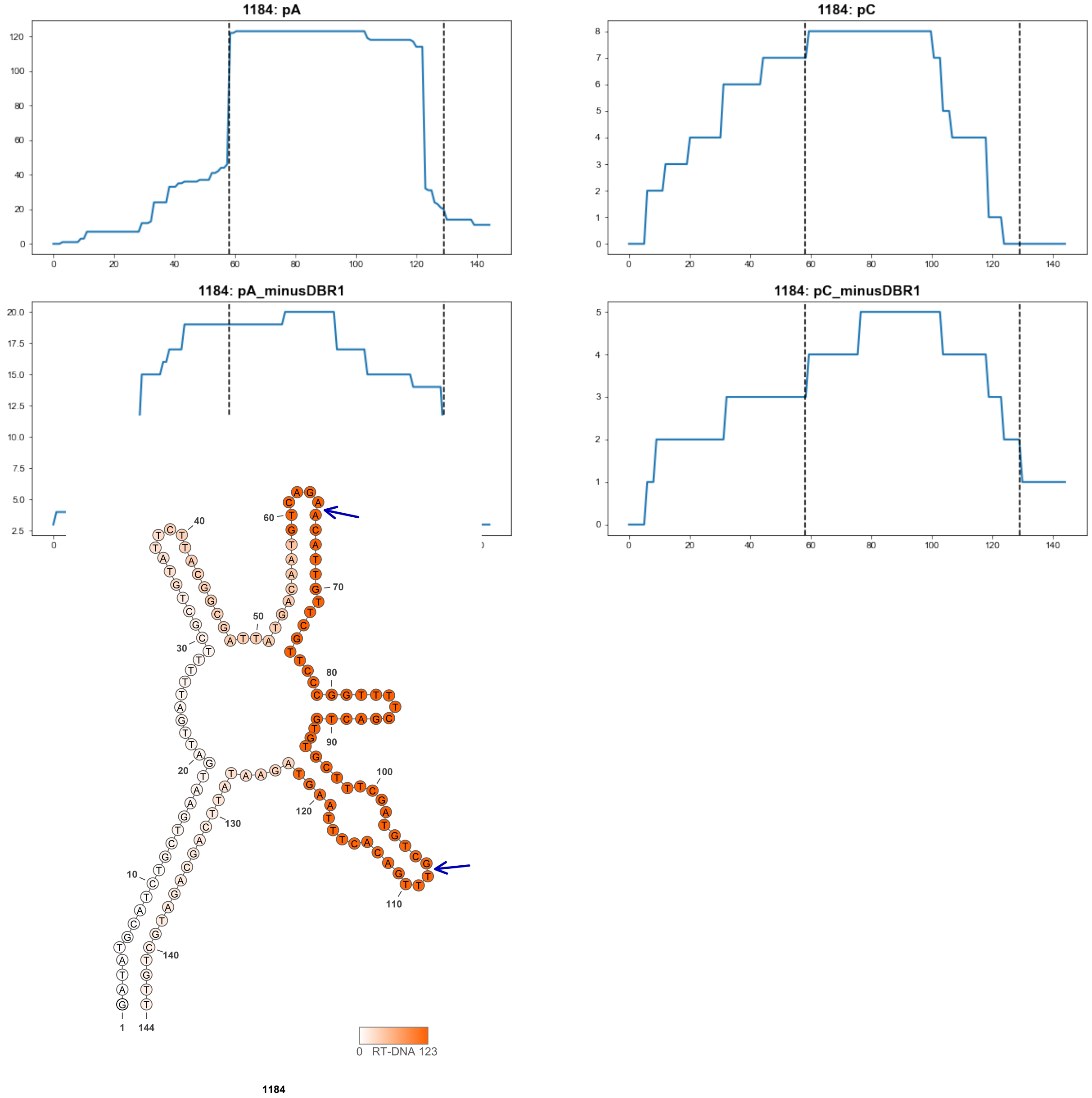

### Retron Terminal 1190: Proteobacteria

**Bacterial Vector for RTDNA: pSLS.642**

**Bacterial Vector for Editing: None**

**Phage Vector for Editing: None**

**Human Vector for Editing: None**

**RTDNA Production (relative to Eco1): 0.709527032 by PAGE**

**Bacterial Editing: undetermined**

**Phage Editing: undetermined**

**Human Editing (demultiplexed): undetermined**

**ncRNA:**

**AAAAACCTTATTATTTAACTTTTTGTTGTTGTTCCTGCAACAAGTAGAAAAAGAATAAAAGGCATTC
TTCTTCAAAATGATTCTTTCTGAAACCACAAGGTGAGCAGAATGAGGCTTAACGAAACATAGTTGG
CTGAAGATTAAGTGAAAAAAATAAGGTTTTT**

**RTDNA:**

**CACTTAATCTTCAGCCAACTATGTTTCGTTAAGCCTCATTCTGCTCACCTTGTGGTTTCAGAAAGAA
TCATTTTGAAGAAGAATGCCTTTT**

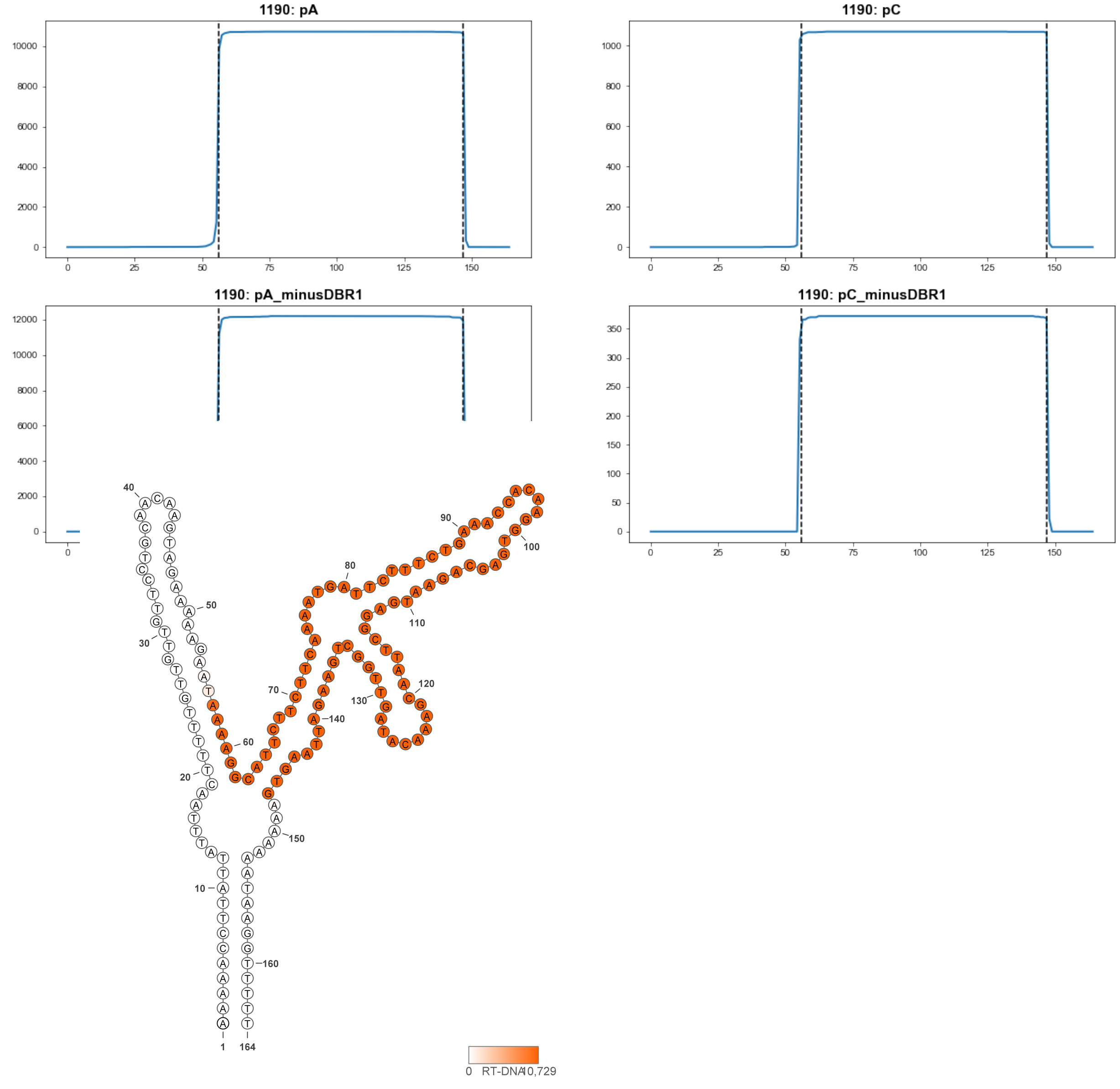


**1190**

### Retron Terminal 1191: Proteobacteria

**Bacterial Vector for RTDNA: pSLS.643**

**Bacterial Vector for Editing: None**

**Phage Vector for Editing: None**

**Human Vector for Editing: None**

**RTDNA Production (relative to Eco1): 0 by PAGE**

**Bacterial Editing: undetermined**

**Phage Editing: undetermined**

**Human Editing (demultiplexed): undetermined**

**ncRNA:**

**ATAGATTCAGGTACATGTTATAGTATGACTCAGCCCTTGAAAGAATGAGAAATGGTTATATTAGAA ATGAATGTTAAAACTTCGTAATAAAATAGAAAAACAATAAGTGAAAATAATAAGGTGTTTTGTGTTT TTTAAAAACTTTTTACGTAATACAACGAAACATGTTCTGAATCTAT**

**RTDNA: None**

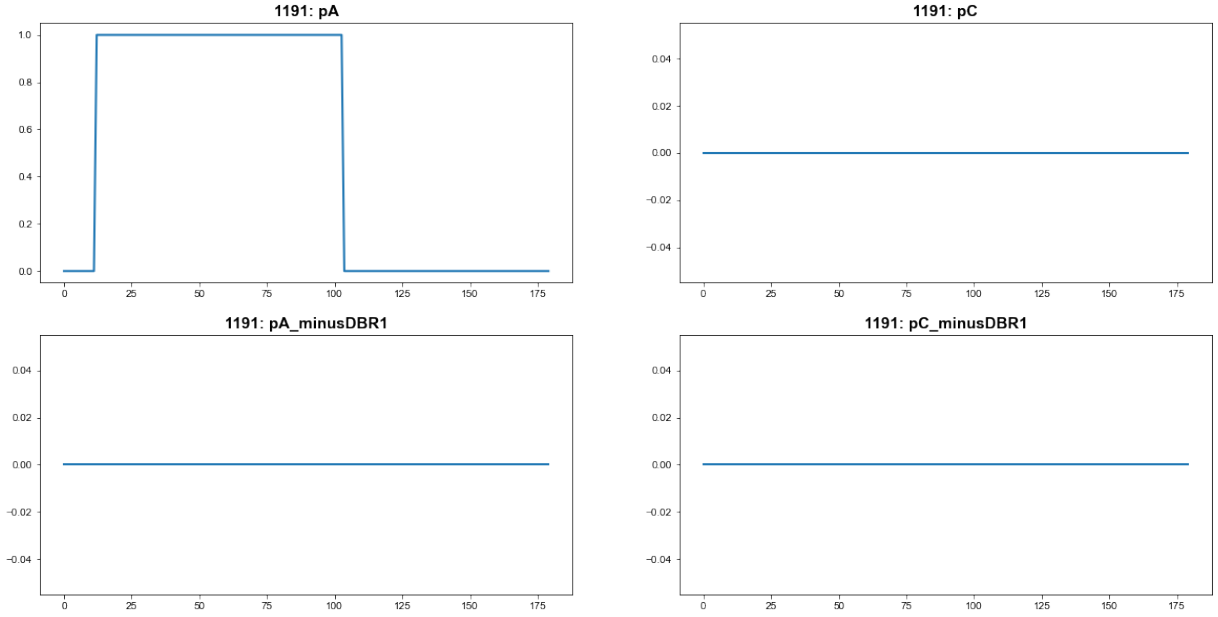


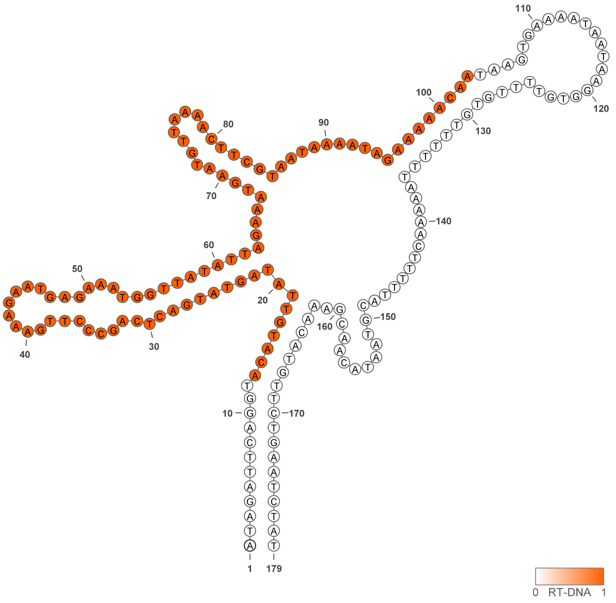


**1191**

### Retron Terminal 1195: Proteobacteria

**Bacterial Vector for RTDNA: pSLS.683**

**Bacterial Vector for Editing: pAGD261**

**Phage Vector for Editing: None**

**Human Vector for Editing: None**

**RTDNA Production (relative to Eco1): 0.285722494 by PAGE**

**Bacterial Editing: 0.41320126 percent precise (0.127147207x Eco1)**

**Phage Editing: undetermined**

**Human Editing (demultiplexed): undetermined**

**ncRNA:**

**GCAGCACCCCTCCAAGCTTTAACTATGCAGTGCTACCACTGCAATCATCGTTAATGTAAGGATATA TGCGACTGTGGATTTCGATGGGACTAATGATTCGCAATGTTATGAGTCTCGCGAGGCGTAAGGTA AATAGCGGGGGGTGCTGC**

**RTDNA:**

**TACCTTACGCCTCGCGAGACTCATAACATTGCGAATCATTAGTCCCATCGAAATCCACAGTCGCAT ATATCCTTAC**

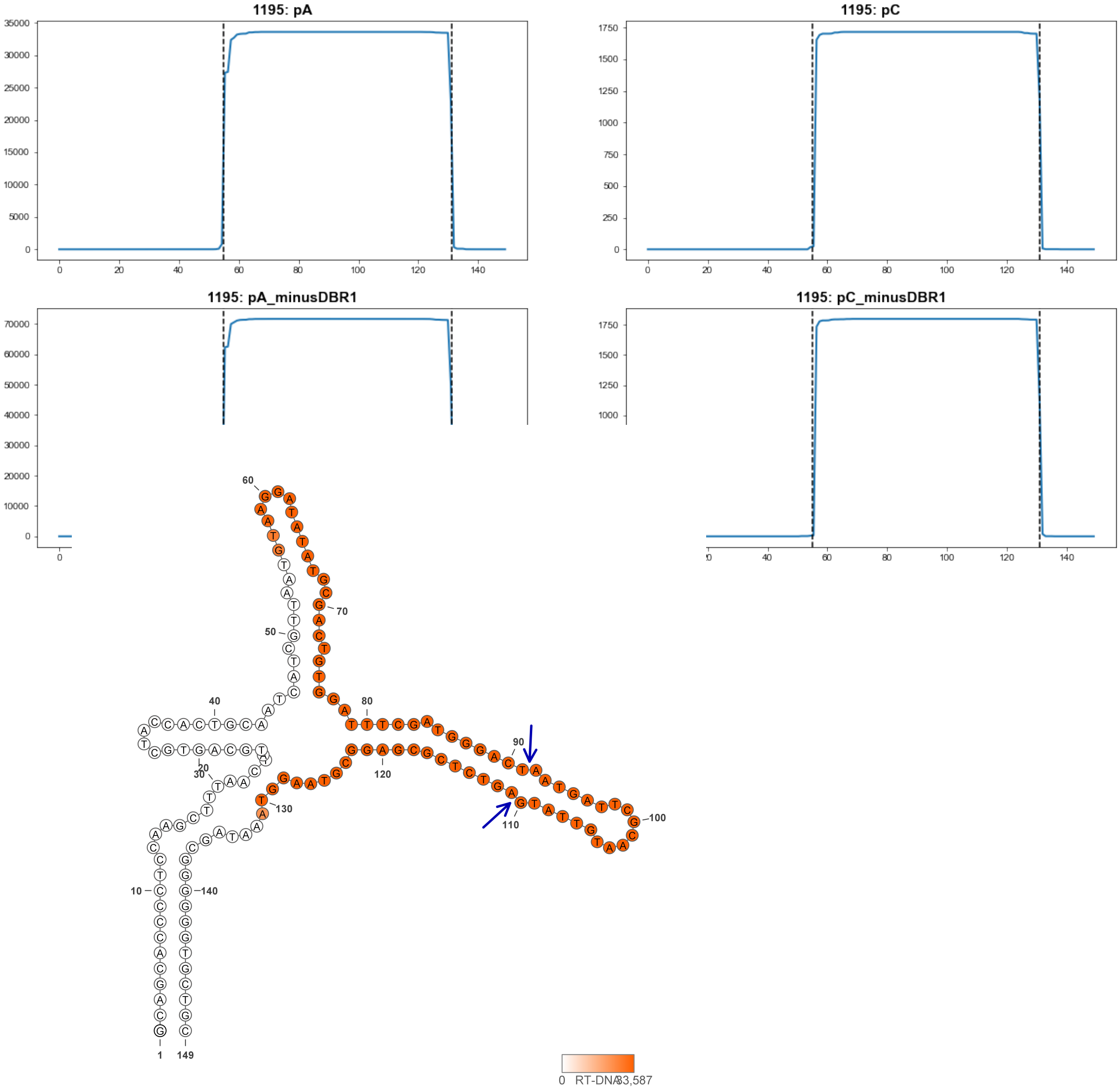


**1195**

### Retron Terminal 1235: Proteobacteria

**Bacterial Vector for RTDNA: pSLS.617**

**Bacterial Vector for Editing: None**

**Phage Vector for Editing: None**

**Human Vector for Editing: pSLS.838**

**RTDNA Production (relative to Eco1): 0.089953538 by PAGE**

**Bacterial Editing: undetermined**

**Phage Editing: undetermined**

**Human Editing (demultiplexed): 0 percent precise (0x Eco1)**

**ncRNA:**

**GCCTCCATTCAGGTATTAAATAGGGTTGTTCAAATGCGACCCGATTCAAAAAAGCCACAAGCAAC GGCACTCAGCGACTCTGAGACGTTACGCTCAAAGGAATAGGGTGAAATATGAATGGAGGC**

**RTDNA:**

**CCATTCATATTTCACCCTATTCCTTTGAGCGTAACGTCTCAGAGTCGCTGAGTGCCGTTGCTTGTG GCT**

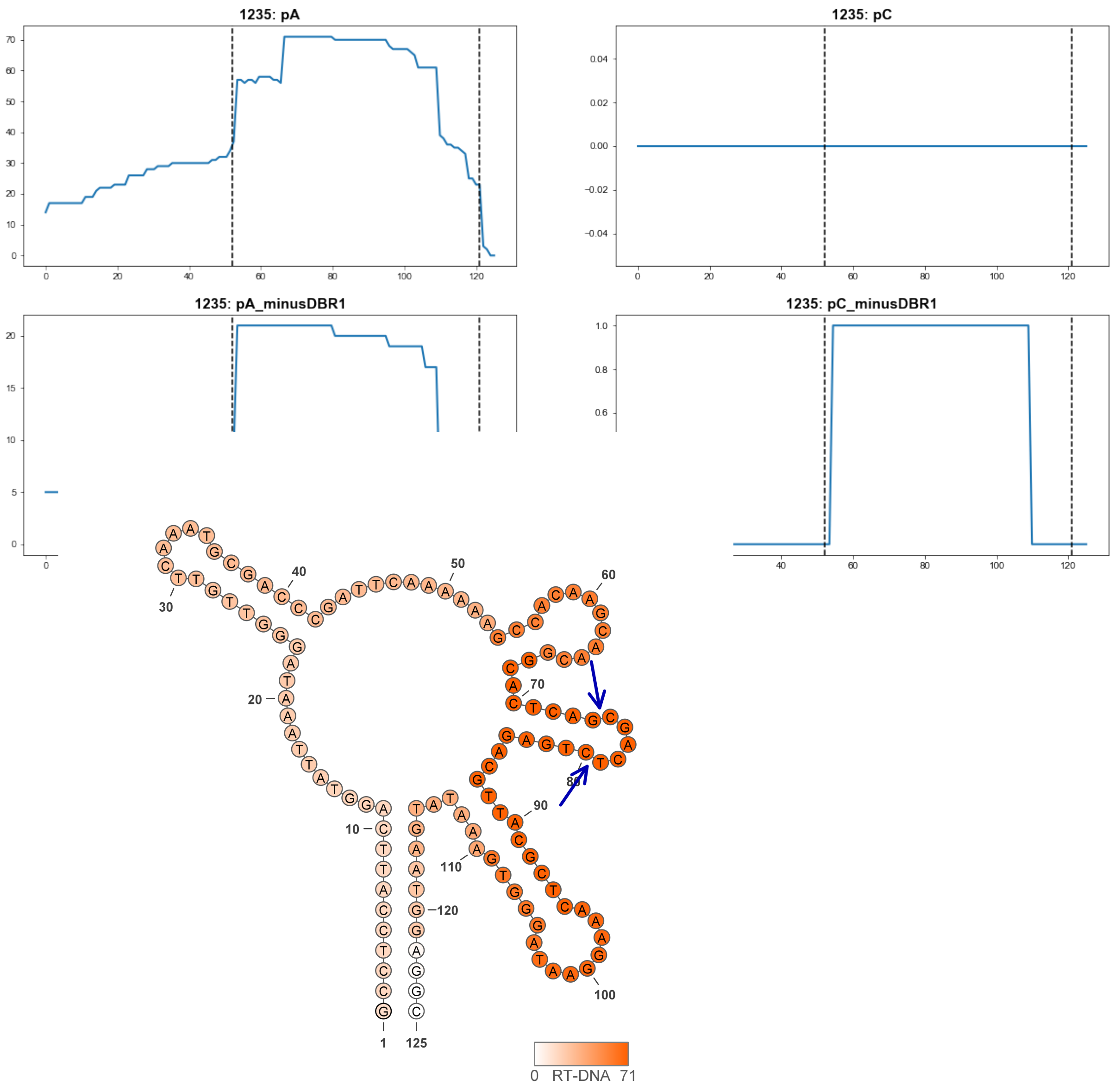


**1235**

### Retron Terminal 1265: Proteobacteria

**Bacterial Vector for RTDNA: None**

**Bacterial Vector for Editing: None**

**Phage Vector for Editing: None**

**Human Vector for Editing: pSLS.934**

**RTDNA Production (relative to Eco1): undetermined**

**Bacterial Editing: undetermined**

**Phage Editing: undetermined**

**Human Editing (demultiplexed): 0.177862 percent precise (1.532091204x Eco1)**

**ncRNA:**

**GCGAGGCGAGAGTTTAACTTTCTGCTGTTAATCTGCGGCGGTCATTGGCATGTTTCATTTATATCC**

**GTTTCTGATGCTCTGCTCCTTGGTGTATCAATGTATAAGAGAAAACAAAGTCGCCTCGC**

**no RTDNA sequencing data**

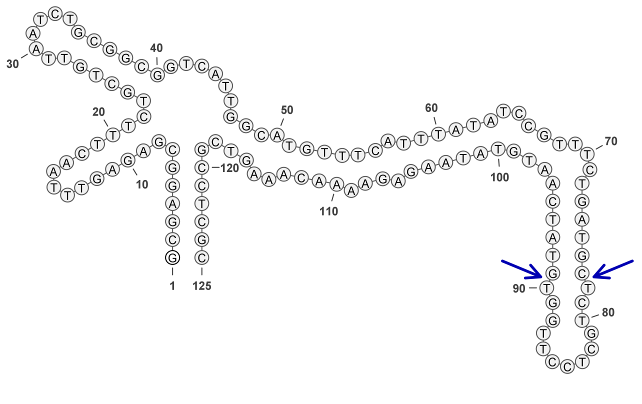


**1265**

### Retron Terminal 1278: Proteobacteria

**Bacterial Vector for RTDNA: pSLS.684**

**Bacterial Vector for Editing: None**

**Phage Vector for Editing: None**

**Human Vector for Editing: None**

**RTDNA Production (relative to Eco1): 0 by PAGE**

**Bacterial Editing: undetermined**

**Phage Editing: undetermined**

**Human Editing (demultiplexed): undetermined**

**ncRNA:**

**CCGGGCTCGGGGTACGATTTAACTTTCCACTGACTATATGCGGTGGCCAATGCAAGGGTAGCTTT**
**GCCTATTCGCGCTTAACGATGTTGTTAACGTTGGCGATGTGTAAAAGAGAAAATGACCCCGAGAC**
**CCG**

**RTDNA: CTCTTTTACACATCGCCAACGTTAACAACATCGTTAAGCGCGAATAGGCAAAGCTACCC**

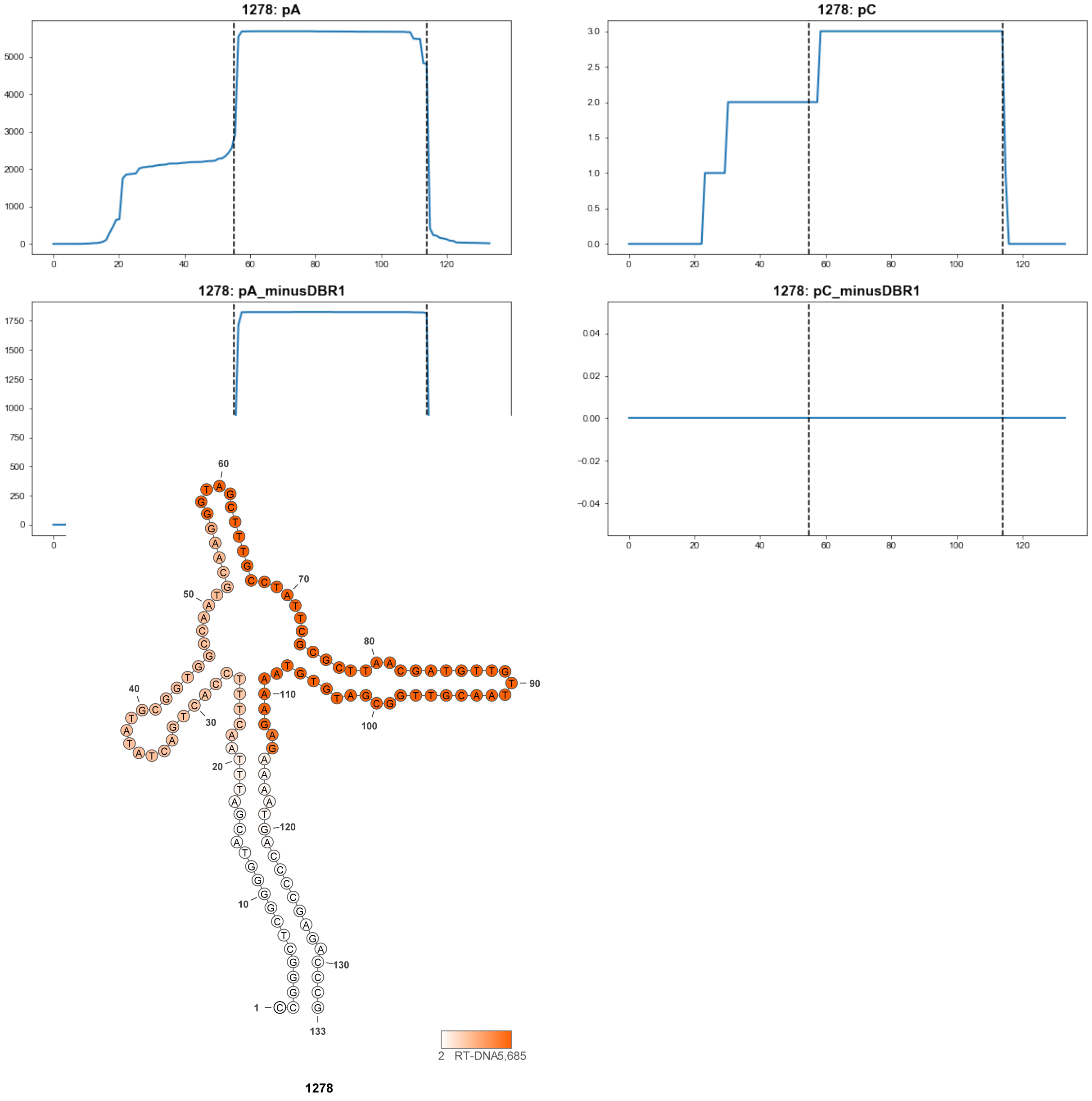

### Retron Terminal 1291: Proteobacteria

**Bacterial Vector for RTDNA: None**

**Bacterial Vector for Editing: None**

**Phage Vector for Editing: None**

**Human Vector for Editing: pSLS.965**

**RTDNA Production (relative to Eco1): undetermined**

**Bacterial Editing: undetermined**

**Phage Editing: undetermined**

**Human Editing (demultiplexed): 0 percent precise (0x Eco1)**

**ncRNA:**

**TTTAGCTTGAACCAATGTAAAATGCATTCGTCTGTAGACCAATGTGTTAAGCATTTCTCTCATAATT**
**TTATTATGAGTAGAGGGGAGTCAGAGTCTGTCCTCCTTGGAAATAGCATCCGATTGATGCTAAGCA**
**TGACCACGCAATTCCGACTGCGTCGGATGGCGTCGAGGCCTTGCCTAATCGCGCGGTCCTCCTCA**
**CCCTGCGGGTGTCGAAGTAAAT**

**no RTDNA sequencing data**

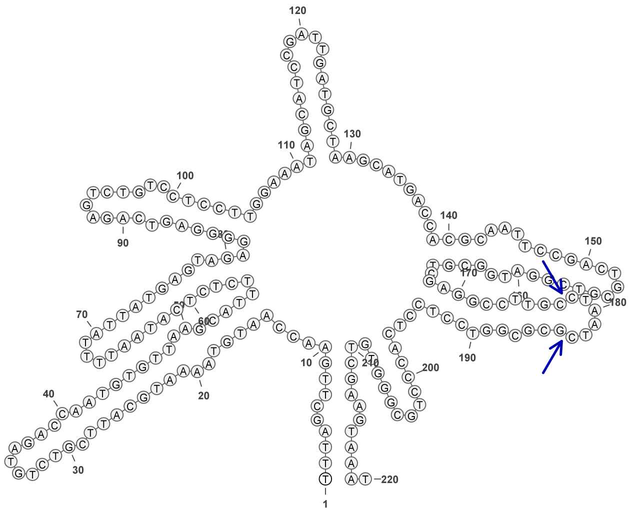

**1291**

### Retron Terminal 1294: Proteobacteria

**Bacterial Vector for RTDNA: pSLS.594**

**Bacterial Vector for Editing: pAGD292**

**Phage Vector for Editing: None**

**Human Vector for Editing: pSLS.829**

**RTDNA Production (relative to Eco1): 0.273897479 by PAGE**

**Bacterial Editing: 1.41124266 percent precise (0.434257054x Eco1)**

**Phage Editing: undetermined**

**Human Editing (demultiplexed): 0.099951 percent precise (0.860971135x Eco1)**

**ncRNA:**

**GCTGAGAATGCCGCGCGATGTCTAGTGGTTGCCGGTTGCGGCACCGTTGGGGCCATCGTTCAGC**
**GATGGACGAGGTCTCTGGGCCGTCGATTGCGACGTGCTCACGGTCAAGCTGGTGCTTGATGCTAA**
**CCTCGATCGAAGCACACGACCTGCGGTCGGCGCTGCGCGTGCCTCGCTTGTGCTCCGATCGAGG**
**TCACCCTGTCGGGTGAATAGTAGATGAACGATTGGCATTCTCAGC**

**RTDNA:**

**ATTCACCCGACAGGGTGACCTCGATCGGAGCACAAGCGAGGCACGCGCAGCGCCGACCGCAGG**
**TCGTGTGCTTCGATCGAGGTTAGCATCAAGCACCAGCTTGACCG**

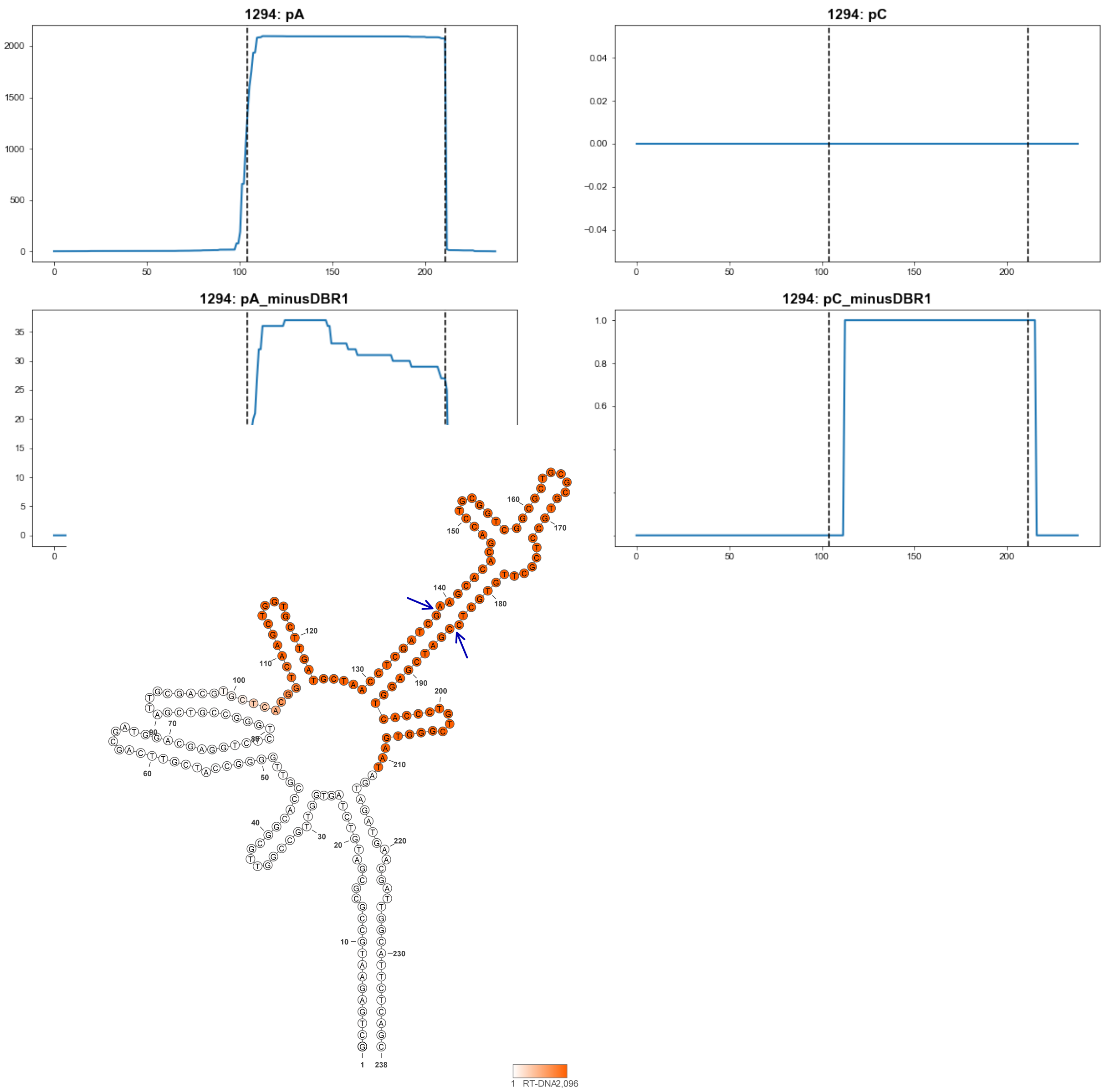


**1294**

### Retron Terminal 1320: Proteobacteria

**Bacterial Vector for RTDNA: None**

**Bacterial Vector for Editing: None**

**Phage Vector for Editing: None**

**Human Vector for Editing: pSLS.878**

**RTDNA Production (relative to Eco1): undetermined**

**Bacterial Editing: undetermined**

**Phage Editing: undetermined**

**Human Editing (demultiplexed): 1.161248 percent precise (10.00291151x Eco1)**

**ncRNA:**

**GCTGAGGGCTCTATGAAAGAGATTAGATATTTGCCACGAAAGTGGCAAGGGGTTCCTTCTGCGGA GCCCAGGGTATGGGAGTTCCTGTGCTCAGATCGCGGAGCAGTCTTTTCATGCTGCGCTTCAGTGC GATCTAGCCCGGGCGATGTCTTAGGGCTAGATCGCACTGATCCCCACCGAGGTGGGTCAAAGTAA TGTCTTTCTGGAGTCCTCAGC**

**no RTDNA sequencing data**

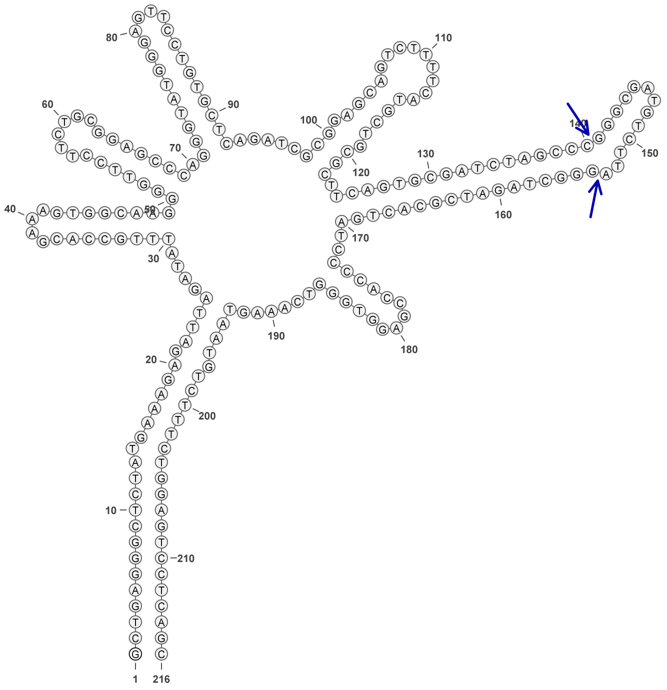

**1320**

### Retron Terminal 1321: Proteobacteria

**Bacterial Vector for RTDNA: None**

**Bacterial Vector for Editing: None**

**Phage Vector for Editing: None**

**Human Vector for Editing: pSLS.879**

**RTDNA Production (relative to Eco1): undetermined**

**Bacterial Editing: undetermined**

**Phage Editing: undetermined**

**Human Editing (demultiplexed): 0.013498 percent precise (0.116270856x Eco1)**

**ncRNA:**

**GCTGAGGACTTCATGATCGAGATTAGACATTTGTCACGAGAGTGGCAAGGGGTTCTGTTTTTGGG**
**ACCCTGGATGCGGGAGTTCCTGCATTCGTTTCGCGGAGCTGTATTTTCATGCTGCGCATTAGTGCG**
**CGTTAGCCCAGGCGACGTCTTCAGGCTAACGCGCACTAATCCCCACCGAGGTGGGTCAAAGTAAT**
**GTCTTCATGGAGTTCTCAGC**

**no RTDNA sequencing data**

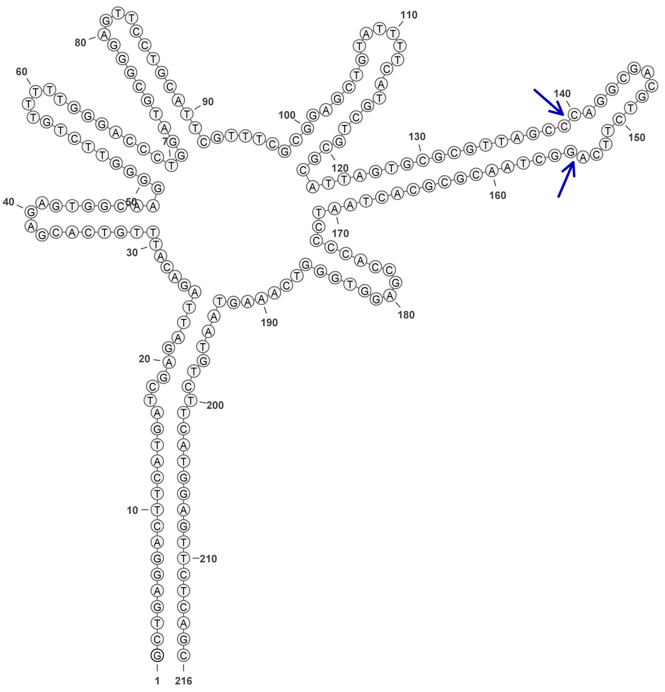

**1321**

### Retron Terminal 1323: Proteobacteria

**Bacterial Vector for RTDNA: pSLS.685**

**Bacterial Vector for Editing: None**

**Phage Vector for Editing: None**

**Human Vector for Editing: pSLS.918**

**RTDNA Production (relative to Eco1): 0.855035911 by PAGE**

**Bacterial Editing: undetermined**

**Phage Editing: undetermined**

**Human Editing (demultiplexed): 0.277885 percent precise (2.393682542x Eco1)**

**ncRNA:**

**TGAGGGCGCCGTGAGGCCGTCTAGGAATGCACTGCGCAGCCAGTGCTGGGTCGGTTTCGACCCA GGGTGTTGGATTTCCGACACTCGCAAAGCGGAGCAGGTTCTTCGCGCTGCGCGGTATGGCAAGA AAAGCCTGTCGAAGTCTCCGGCAGTTCTCGCCATACCCCCCACCCTTGGTGGGCCGAAGTAGATG CGTCATACGAGCCCTCA**

**RTDNA:**

**TTCGGCCCACCAAGGGTGGGGGGTATGGCGAGAACTGCCGGAGACTTCGACAGGCTTTTCTTGC CATACCGCGCAGCGCGAAGAACCTGCTCCGCTTTGCGAGTGTCGGAAATCCAACACCCTGGGTC GAAACCGACCCAGCACTGGCTGCGCAGTGCATTC**

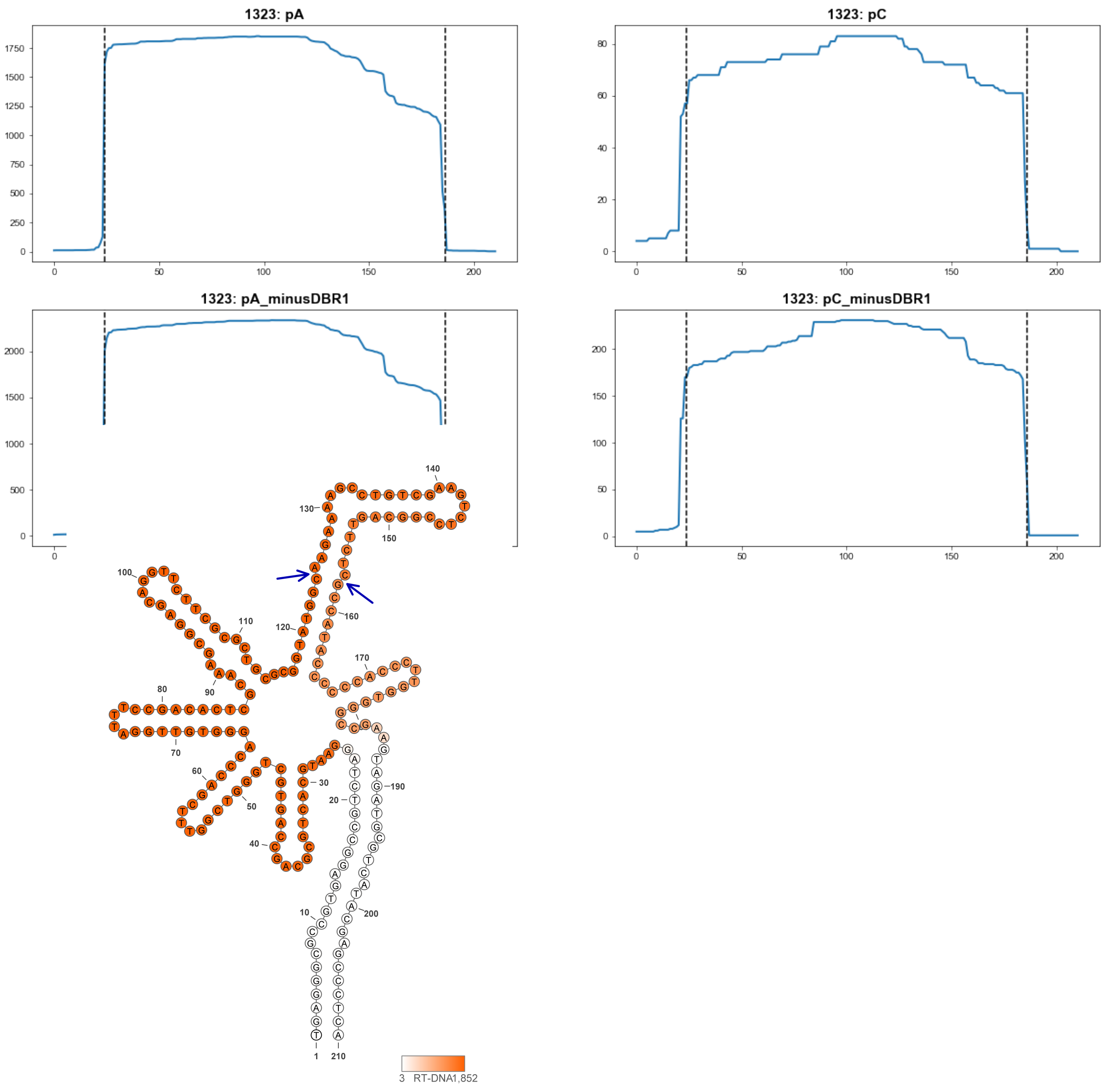


1323

### Retron Terminal 1336: Proteobacteria

**Bacterial Vector for RTDNA: None**

**Bacterial Vector for Editing: None**

**Phage Vector for Editing: None**

**Human Vector for Editing: pSLS.935**

**RTDNA Production (relative to Eco1): undetermined**

**Bacterial Editing: undetermined**

**Phage Editing: undetermined**

**Human Editing (demultiplexed): 0.19304 percent precise (1.662833467x Eco1)**

**ncRNA:**

**GCTGAGGGCTCCGCAGTAGCGGAATCTAGTATCCCGCACGATGTGCGGGCGTTAAGGGAGCAAA**
**TTATGCTTGCCGTCGAACGATCTCACGCTTCCAGGAGGTATAAGCCTTCTACGGCTCGCTATCGAA**
**GTCTTGCGGCAAGAAGGCTTATATCCGCCAGGGCGGAGCTTGAAGGTAGATGACTCGCTGGAGC**
**TCTCAGC**

**no RTDNA sequencing data**

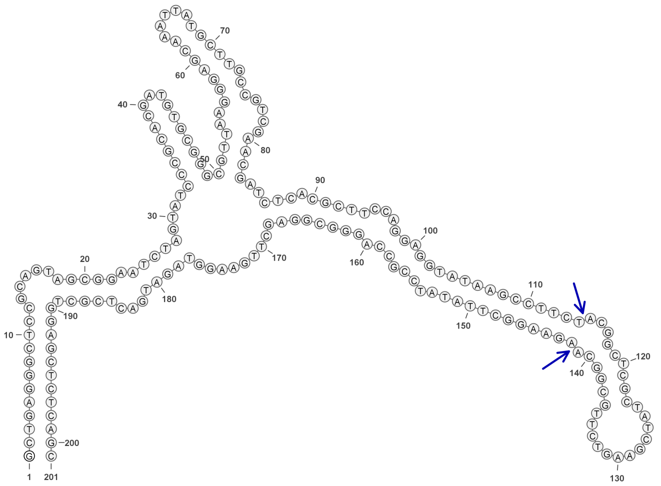

**1336**

### Retron Terminal 1352: Proteobacteria

**Bacterial Vector for RTDNA: pSLS.618**

**Bacterial Vector for Editing: None**

**Phage Vector for Editing: None**

**Human Vector for Editing: pSLS.839**

**RTDNA Production (relative to Eco1): 0.990479092 by PAGE**

**Bacterial Editing: undetermined**

**Phage Editing: undetermined**

**Human Editing (demultiplexed): 0 percent precise (0x Eco1)**

**ncRNA:**

**AACGTTGAGGAGGCCATAATGTTTCATTTAGCGTTAACACAAGTAACTACCATTACTTGTCAGGAG AAGTGTATACCTTTCCTGCGGGCCATATATTTTATGTGCTCATGTATTAGCACTCGCAATAGTGTAA GCAAAATCACGACGATCCCGACTTCGTCGGCTCGATCGTGGTCTTTCTTATCGTCGCGATTTTACT TACCCCTCCGGGGTCAAAGTAAATGAAACAATGGTCTCCTCAGCGTT**

**RTDNA:**

**TTGACCCCGGAGGGGTAAGTAAAATCGCGACGATAAGAAAGACCACGATCGAGCCGACGAAGTC GGGATCGTCGTGATTTTGCTTACACTATTGCGAGTGCTAATACATGAGCACATAAAATATATGGCC CGCAGGAAAGGTATACACTTCTCCTGACAAGT**

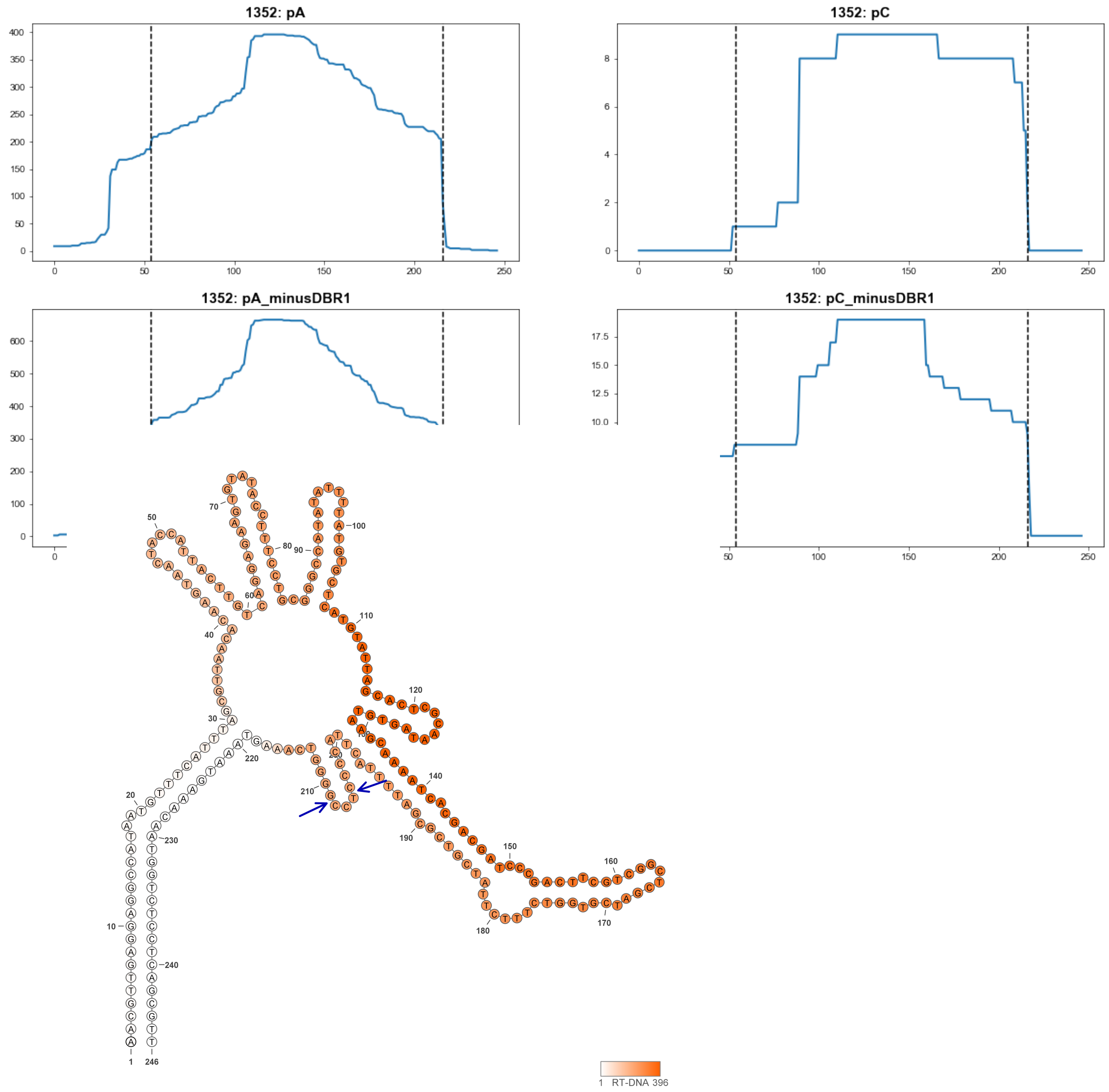


**1352**

### Retron Terminal 1353: Proteobacteria

**Bacterial Vector for RTDNA: pSLS.619**

**Bacterial Vector for Editing: None**

**Phage Vector for Editing: None**

**Human Vector for Editing: pSLS.840**

**RTDNA Production (relative to Eco1): 0.529590982 by PAGE**

**Bacterial Editing: undetermined**

**Phage Editing: undetermined**

**Human Editing (demultiplexed): 0.37742 percent precise (3.251070281x Eco1)**

**ncRNA:**

**GTTGAGGAGACCATAATGTTTCATTTAGCGTTCGCTCAAGTAACTATTACAGTTACTTGTCTGATTA
GTTTTAAACTCAATCAGTCGGCTATATTCTATATAAGCCCAAGTAATAGCTCTCGCAAAAGAGTAA
CCAAGATCACGACAATAGCGACTAAAGTCGCTCGATTGTGTCCTCACTTATTGTCGCGATCTTACT
TACCCCTCCGGGGCCAAAGTAAATGAAATCATGGTCTCCTCAGC**

**RTDNA:**

**TTGGCCCCGGAGGGGTAAGTAAGATCGCGACAATAAGTGAGGACACAATCGAGCGACTTTAGTC
GCTATTGTCGTGATCTTGGTTACTCTTTTGCGAGAGCTATTACTTGGGCTTATATAGAATATAGCCG
ACTGATTG**

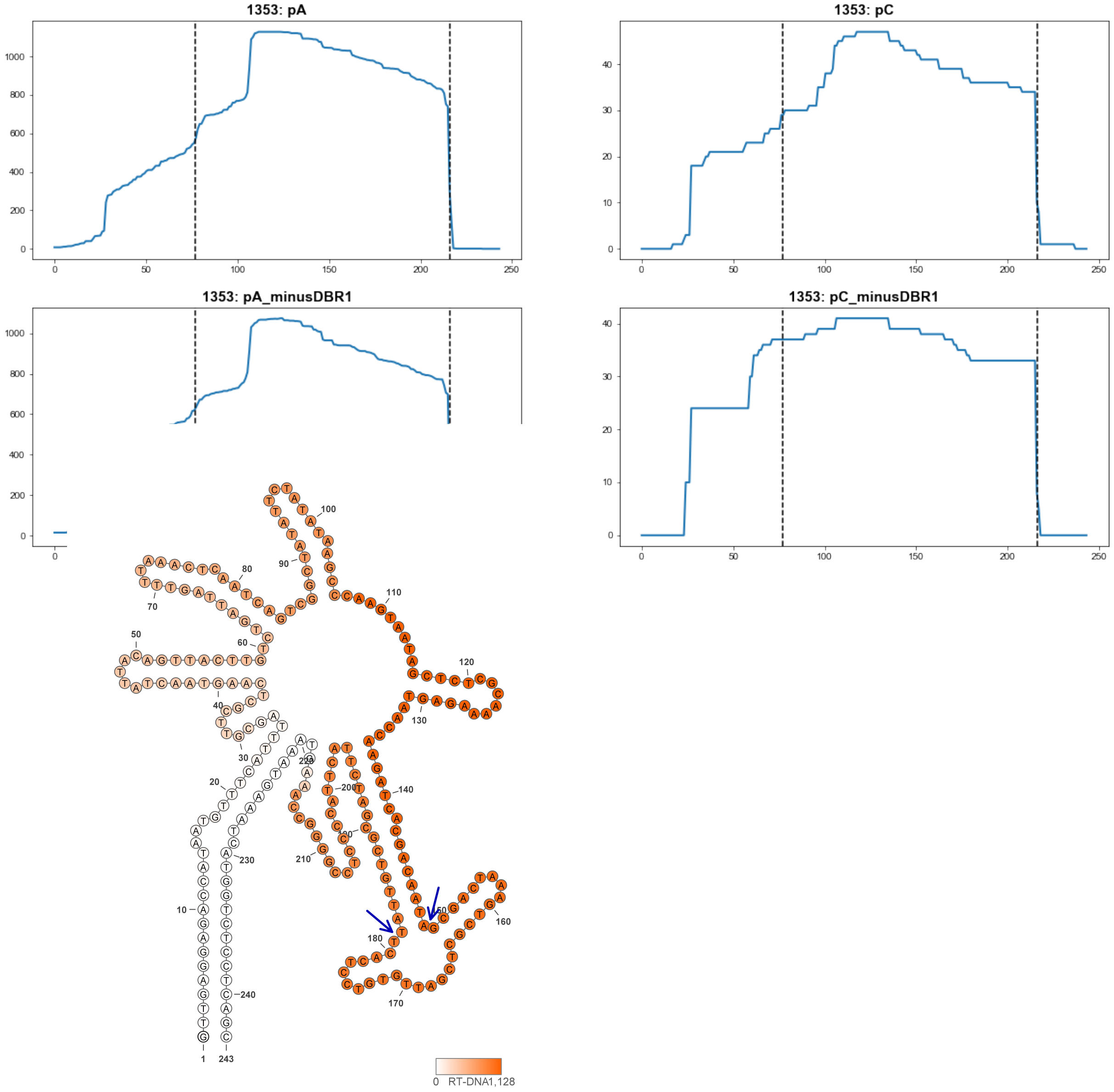


1353

### Retron Terminal 1373: Proteobacteria

**Bacterial Vector for RTDNA: None**

**Bacterial Vector for Editing: None**

**Phage Vector for Editing: None**

**Human Vector for Editing: pSLS.880**

**RTDNA Production (relative to Eco1): undetermined**

**Bacterial Editing: undetermined**

**Phage Editing: undetermined**

**Human Editing (demultiplexed): 0.52234 percent precise (4.499401332x Eco1)**

**ncRNA:**

**GGTGTGCTGATCAGCACTCCGTTCAGCCGTTAAACAATGATGAGTGAAATTCACCGCTTGTTTGAC**
**AGAATCTGTTGTGGCTTTTGTTACAACGGATGGGGATGTTGGAGAAACATCTCTGGAAAGATCTGA**
**CGCAAAGGCGGCTACGCCGCCCTTCAATGTCATACATTAACGGCCATCGAGGACTCGGAAAATGT**
**ATGACACTGATGGCCCCCTGGGCTCAAAGGATAGGTGAATGGAAATCTGGTCAGCACACC**

**no RTDNA sequencing data**

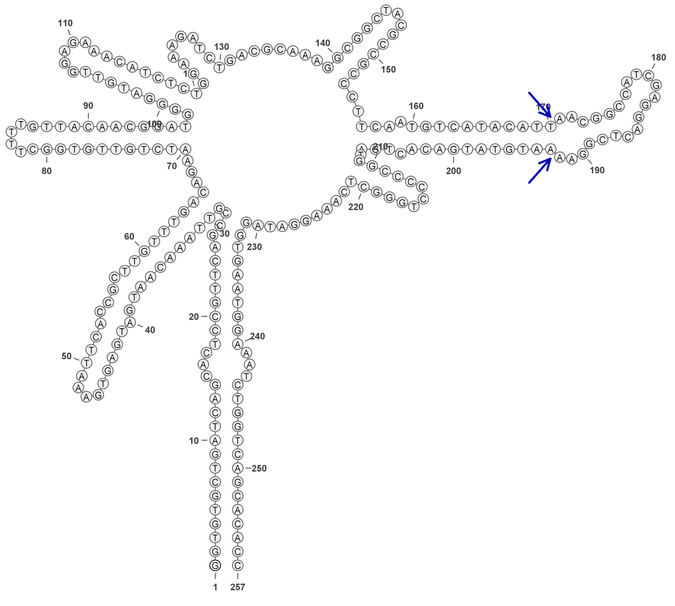

**1373**

### Retron Terminal 1374: Proteobacteria

**Bacterial Vector for RTDNA: pSLS.644**

**Bacterial Vector for Editing: None**

**Phage Vector for Editing: None**

**Human Vector for Editing: pSLS.881**

**RTDNA Production (relative to Eco1): 1.087299797 by PAGE**

**Bacterial Editing: undetermined**

**Phage Editing: undetermined**

**Human Editing (demultiplexed): undetermined**

**ncRNA:**

**GGTGTGCTGATCGGTATTCTATTCAGCCGTTAAACAAGAATGAGTGATTTATCACCACTTGTTTGA CAGATTCTGTTGTAGCTTCTGTTACAATAGATGGGGATGTTGGAGAAACGTCTCTGGCAAGATCTA ACGCTAAGGCGGCTACGCCGCCCTTCAATGTCATACCTTATGGGCCTTCGAAGACTCGGAAAATG TACGACACTGATGGCCCCCTGGGCTCAAAGGATAGGTGAATGGAAATCTGGTCAGTACACC**

**RTDNA:**

**CTATCCTTTGAGCCCAGGGGGCCATCAGTGTCGTACATTTTCCGAGTCTTCGAAGGCCCATAAGGT ATGACATTGAAGGGCGGCGTAGCCGCCTTAGCGTTAGATCTTGCCAGA**

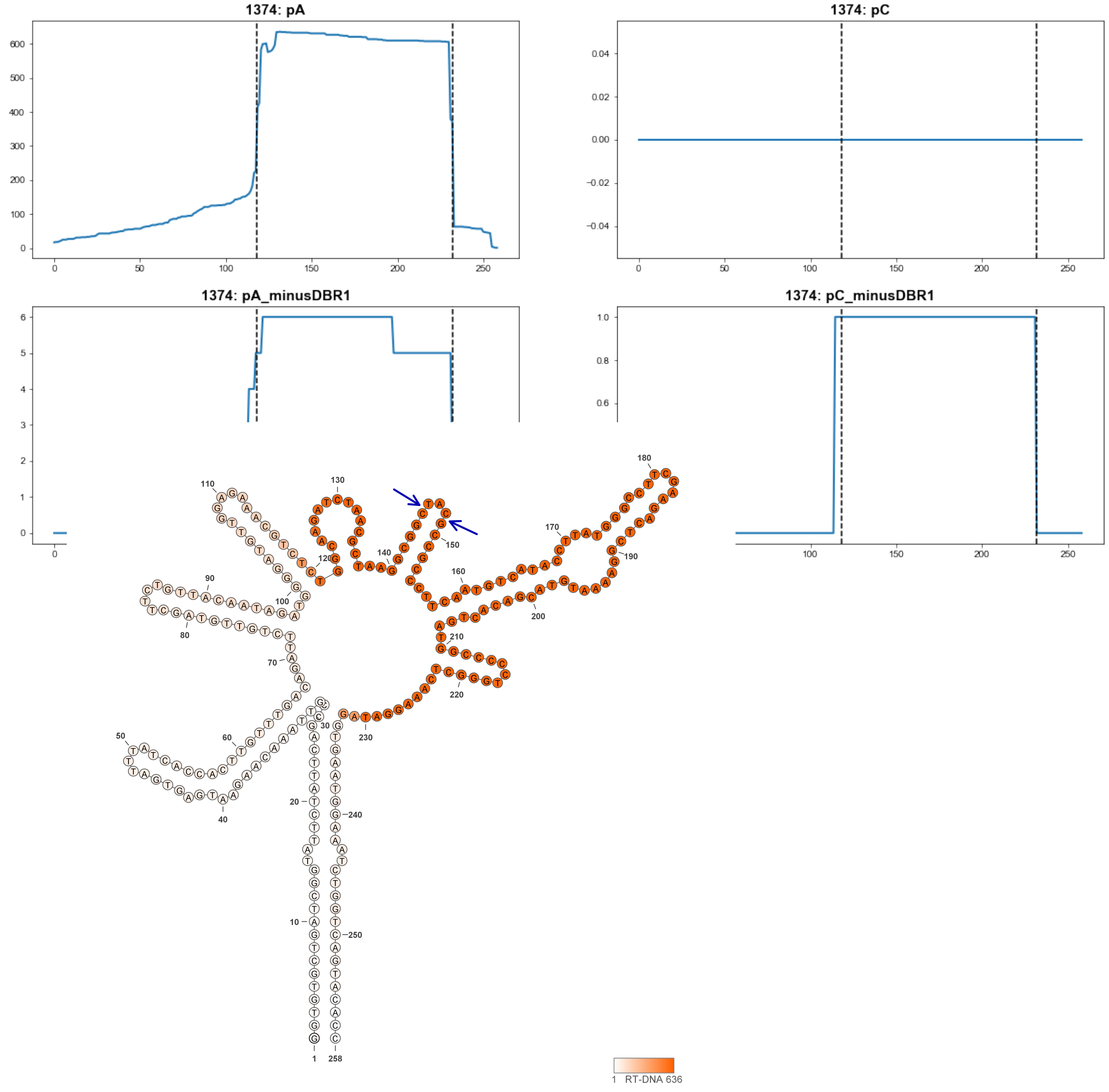


**1374**

### Retron Terminal 1391: Proteobacteria

**Bacterial Vector for RTDNA: pSLS.623**

**Bacterial Vector for Editing: None**

**Phage Vector for Editing: None**

**Human Vector for Editing: pSLS.846**

**RTDNA Production (relative to Eco1): 0.66128418 by PAGE**

**Bacterial Editing: undetermined**

**Phage Editing: undetermined**

**Human Editing (demultiplexed): 0.040212 percent precise (0.346383441x Eco1)**

**ncRNA:**

**GTATGAGTCGGGCCATGAGGCATGAAGACTAGTGTTAACCATGCGGCTTACGTATGGTGCATTAC
AGGGAAACCTGTAGTGTTCTCATTTAACTTTTTGTTAAATGAACTGTATTCCAAAGTTCGCTTTGTA
ATACAGTTTTGCTTTAAGGTTAATTTAAAGTAATGGAAAAGGCCTCGCTTTAATTCGCGTTTTACTG
CTAAAGCAATTTCGGTCGTAGCCTTGGTCATTTGCTTTAGCAGTAAAATGCAAGCATCAAAGGTAG
TTTTATGGCAATTGGACCGGCTCATAT**

**RTDNA:**

**TAAAGCGAGGCCTTTTCCATTACTTTAAATTAACCTTAAAGCAAAACTGTATTACAAAGCGAACTTT
GGAATACAGTTCATTTAACAAAAAGTTAAATGAGAACACTACAGGTTTCCCTGTAATGCACCATAC
GTAAGCCGCATGGTTAACACTA**

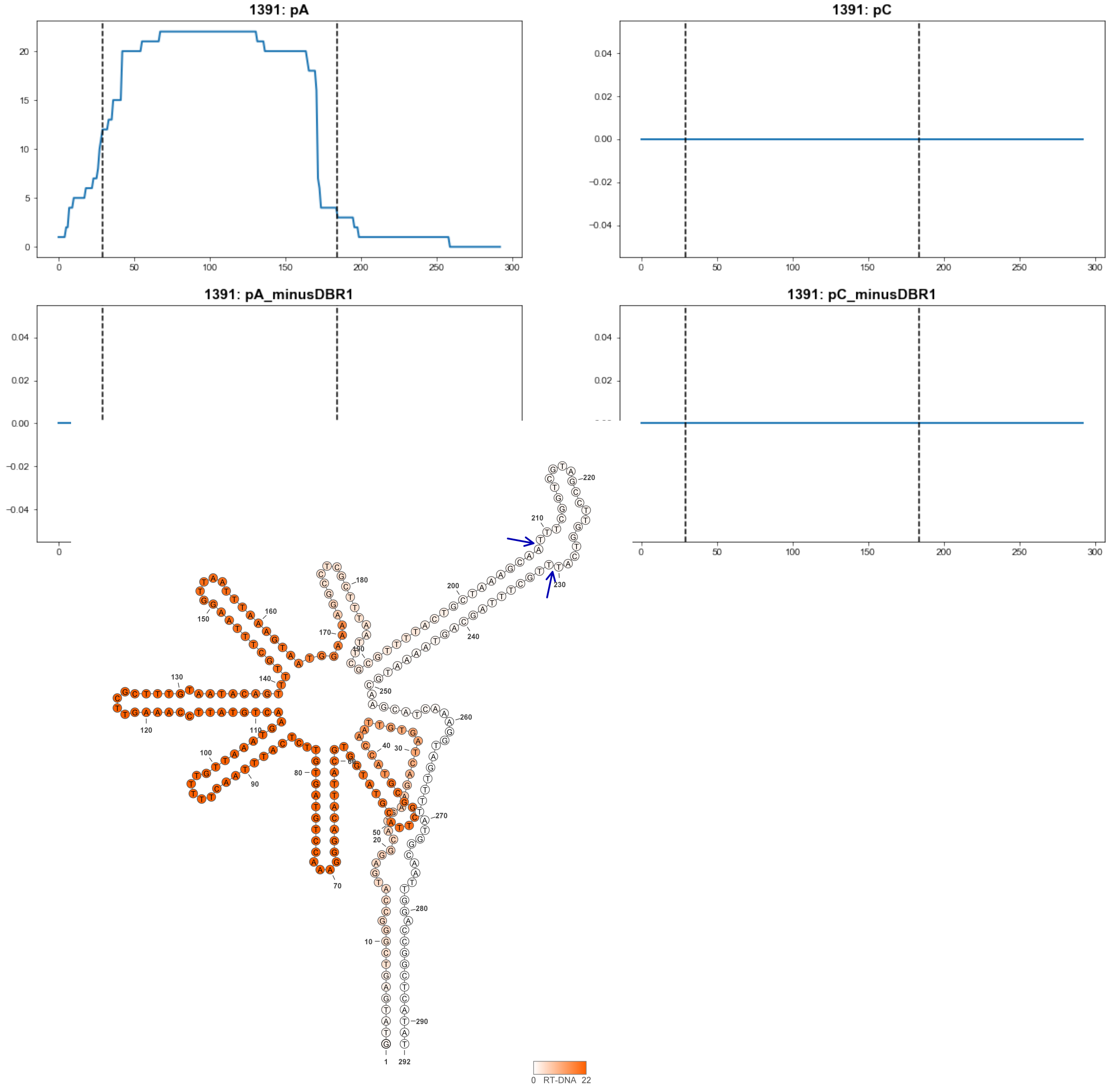


**1391**

#### Retron Terminal 1400: Actinobacteria

**Bacterial Vector for RTDNA: None**

**Bacterial Vector for Editing: None**

**Phage Vector for Editing: None**

**Human Vector for Editing: pSLS.882**

**RTDNA Production (relative to Eco1): undetermined**

**Bacterial Editing: undetermined**

**Phage Editing: undetermined**

**Human Editing (demultiplexed): 0.558756 percent precise (4.813086286x Eco1)**

**ncRNA:**

**TGAGGGGGGTCCGTGACGTCTAGCGTCTGTCGATGCCGTTGGTGTCGACACCGCCGGGTTGGAC**
**CCGGGGCTTATGCAGGCGGCGGGATACCCGCAAGCCTGGACCAAACCCGCGCTTCGCCTCCTGC**
**CGTGAACCGCGCTGCTCCGGCGATGTCTTGGGCGCGGCTCACGGCACGACTGGCGTCGCGGAGG**
**CTCAAAGGTAGATGATCACTGGCTCCCCTCA**

**no RTDNA sequencing data**

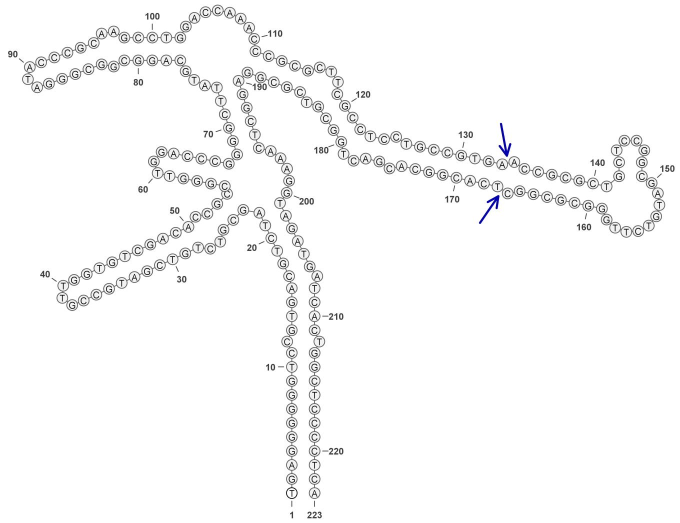

**1400**

### Retron Terminal 1410: Proteobacteria

**Bacterial Vector for RTDNA: None**

**Bacterial Vector for Editing: None**

**Phage Vector for Editing: None**

**Human Vector for Editing: pSLS.883 (U6, mod)**

**RTDNA Production (relative to Eco1): undetermined**

**Bacterial Editing: undetermined**

**Phage Editing: undetermined**

**Human Editing (demultiplexed): 0.051368 percent precise (0.442480468x Eco1)**

**ncRNA:**

**GCTTAAAATGACGTTATTAGGAAAATGTGTTCTACAAAAATATAAGAAAACAAAATTAAAAGATAT**
**AAATTCAAGAAAAAGTATTGATTCCAATCAGTTTATTTACTATCCTCATCAATGTGATGGACATTTT**
**CGATTTCTGGTAGACTGTTGATAAAATACTTGATTAGTAGTTATCAACTGTACAGCTATAACTTGAT**
**TGTAGTAGTTGTTCGGAGCTCAGAGTCTGAAGCTCTATGAAATAGCTTAATTTTAGGC**

**no RTDNA sequencing data**

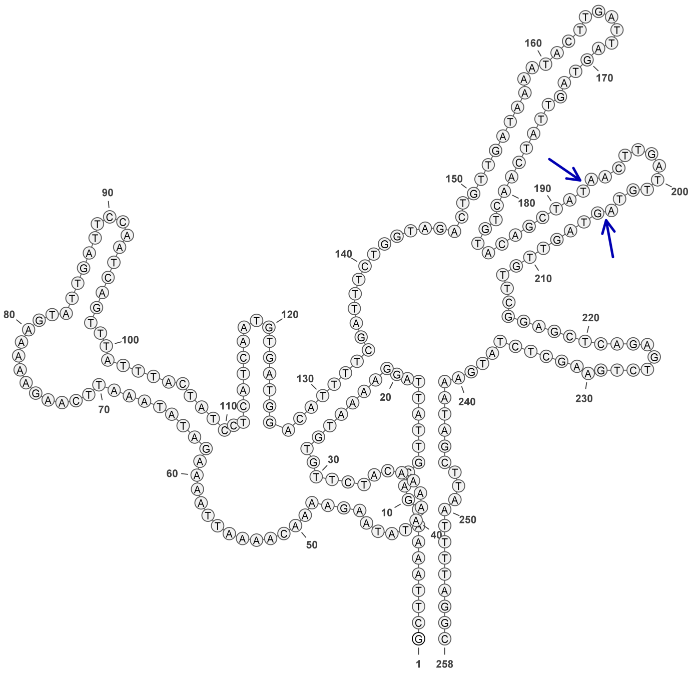


**1410**

### Retron Terminal 1421: Proteobacteria

**Bacterial Vector for RTDNA: pSLS.595**

**Bacterial Vector for Editing: pAGD293**

**Phage Vector for Editing: pAGD402**

**Human Vector for Editing: pSLS.830**

**RTDNA Production (relative to Eco1): 3.168862823 by PAGE**

**Bacterial Editing: 6.83033622 percent precise (2.101780062x Eco1)**

**Phage Editing: 24.27984944 percent precise (0.598854528x Eco1)**

**Human Editing (demultiplexed): 0 percent precise (0x Eco1)**

**ncRNA:**

**TGGGGTTCCGTCGTGAAACTTAGTTGTGGTAGATACATATATAGTGTATCAACTGCCATTGGCAGC
CTACGATTCCGTTTGAATGGTTTGACAGGATGCCAACGAGCACGGATTCCGTCGCTCTTTGTAGTA
GCCTTCGCCAGAGTGAGTAGTGATGTGCGCGTTTGCGCGCGTTCTTTTGCTCTGGGGCAACGAAG
CCTGTGCGCAAACGCGCGCATCACGACCTCACTTACGTGAGGTGAAGTAAGTGACTCAATGGAAC
CCCA**

**RTDNA:**

**TTCACCTCACGTAAGTGAGGTCGTGATGCGCGCGTTTGCGCACAGGCTTCGTTGCCCCAGAGCAA
AAGAACGCGCGCAAACGCGCACATCACTACTCACTCTGGCGAAGGCTACTAC**

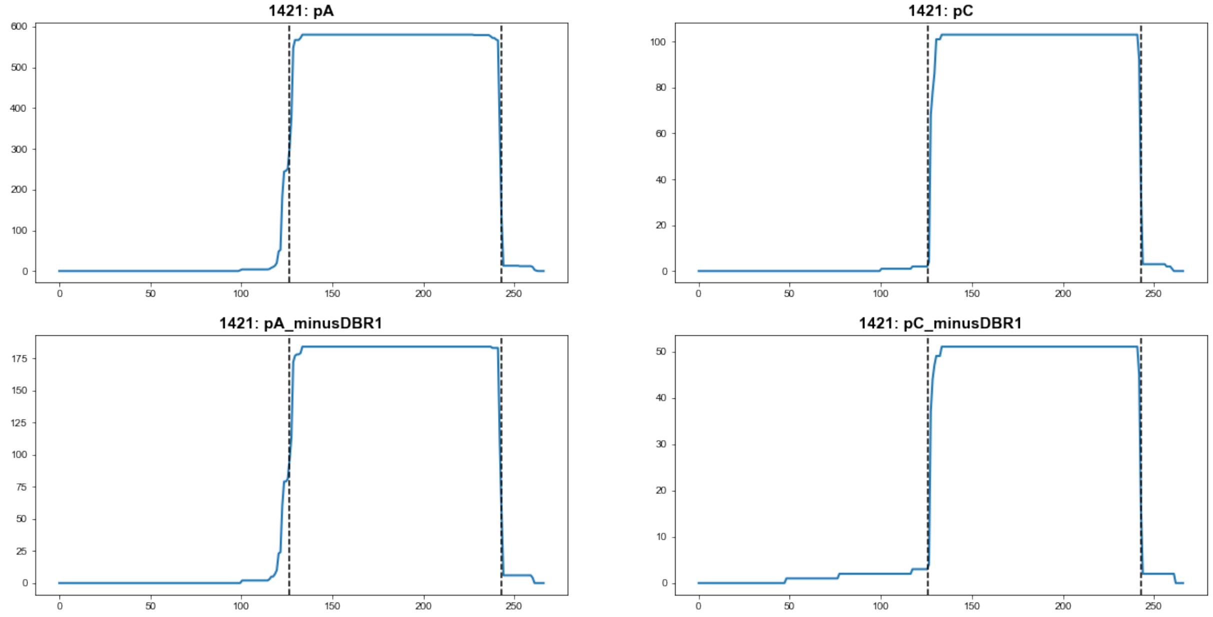


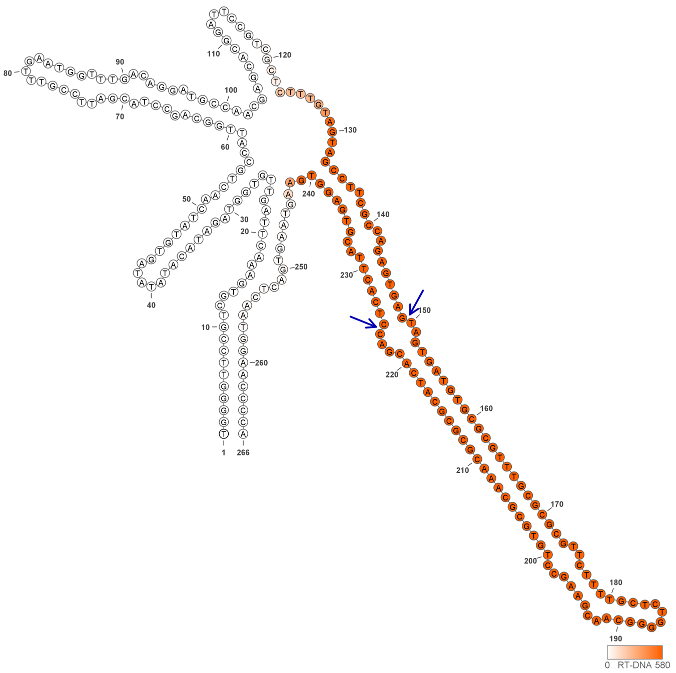


**1421**

### Retron Terminal 1422: Proteobacteria

**Bacterial Vector for RTDNA: pSLS.596**

**Bacterial Vector for Editing: None**

**Phage Vector for Editing: None**

**Human Vector for Editing: pSLS.831**

**RTDNA Production (relative to Eco1): 0.204204433 by PAGE**

**Bacterial Editing: undetermined**

**Phage Editing: undetermined**

**Human Editing (demultiplexed): undetermined**

**ncRNA:**

**GGCTTCAGCAGGGCTCGGATTTCCGCAGTCCAATGGCATAGCTTTCGATTGAGTGGCGCACAGGC TACCGAATTTGAAGATCGCGCTTCGCTTGAGGCTGCGGCCTCCTCTCCAAATTGGCAGCCTGCCC CTCACTGCGTGAGGTAAAGTAAATGACTCGTTGGAGTC**

**RTDNA: None**

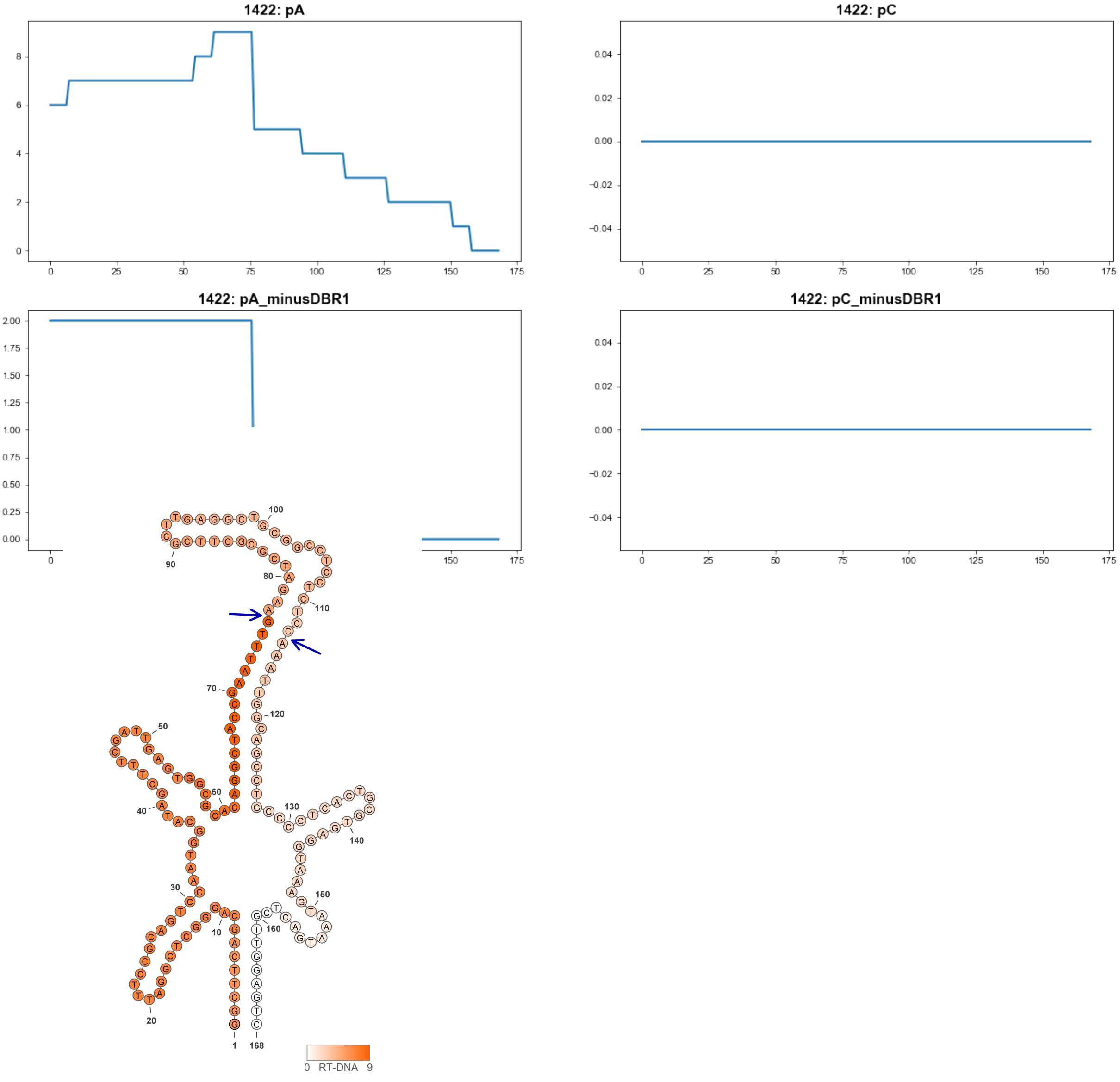


1422

### Retron Terminal 1429: Proteobacteria

**Bacterial Vector for RTDNA: pSLS.645**

**Bacterial Vector for Editing: pAGD299**

**Phage Vector for Editing: pAGD403**

**Human Vector for Editing: pSLS.884**

**RTDNA Production (relative to Eco1): 1.900218062 by PAGE**

**Bacterial Editing: 5.24661509 percent precise (1.614449221x Eco1)**

**Phage Editing: 33.31991159 percent precise (0.821824697x Eco1)**

**Human Editing (demultiplexed): 20.76062 percent precise (178.8305726x Eco1)**

**ncRNA:**

**GACTTGTAGAAACTAGCACGTTGGAGTGTAGTCGCTCCTTGGTTAAGGGTCACTTTAACCACGACT
ATACGGATAACTCCGTATAGTCGCTTCGAATAGCGTTGCTATTCTCGCGGTTCTTTCAAGGTCGAG
ATGTCGCGTCACGCAACATCTCGCTCTTGAAGAGAACTTAGGTGTCAGGTATAGAAAGTAGTTTCT
ACAAGTC**

**RTDNA:**

**TCTATACCTGACACCTAAGTTCTCTTCAAGAGCGAGATGTTGCGTGACGCGACATCTCGACCTTGA
AAGAACCGCGAGAATAGCAACGCTATTCGAAGCGACTATACGGAGTTATCCGTATAGTCGTGGTT**

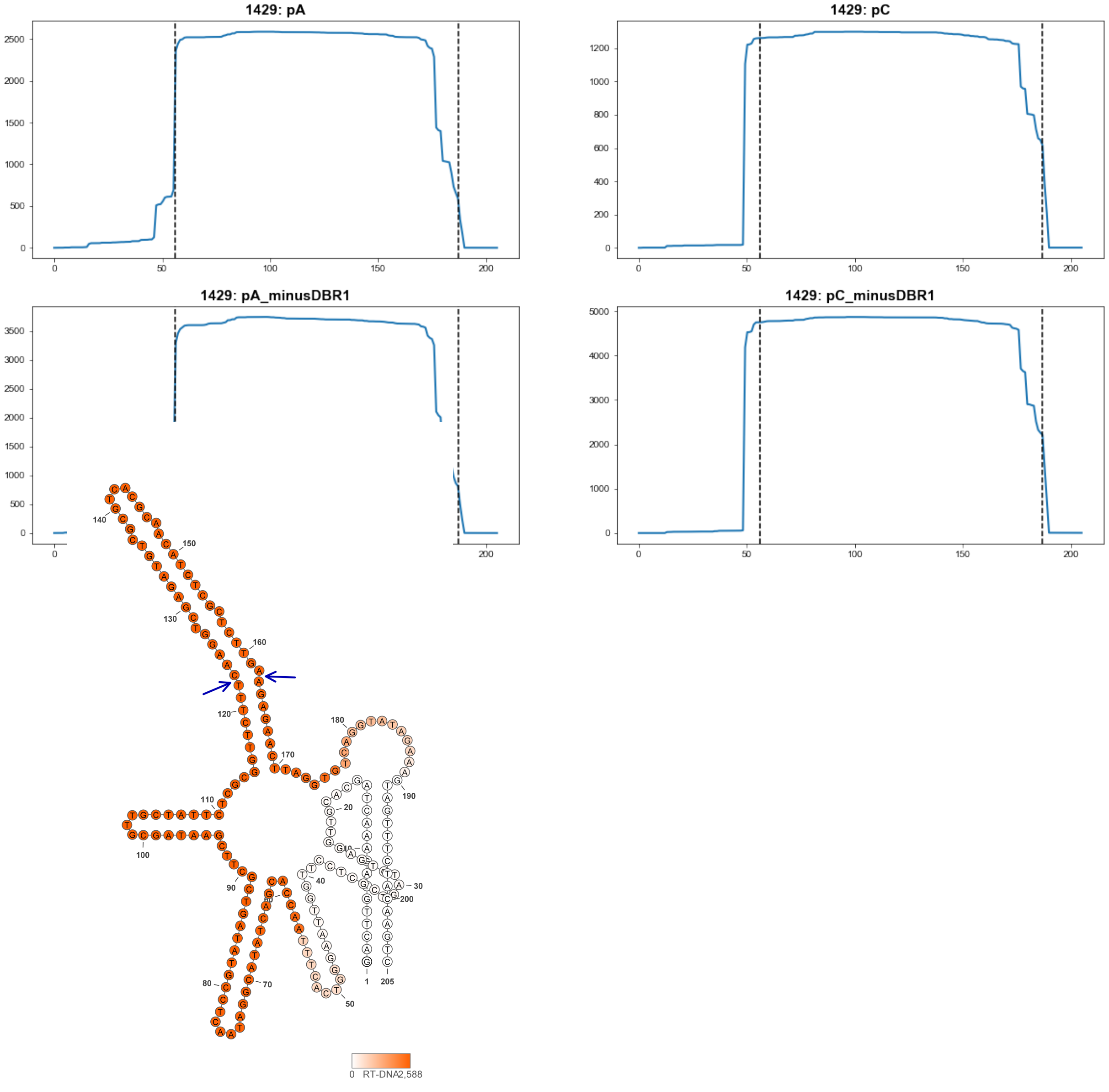


1429

### Retron Terminal 1438: Proteobacteria

**Bacterial Vector for RTDNA: None**

**Bacterial Vector for Editing: None**

**Phage Vector for Editing: None**

**Human Vector for Editing: pSLS.936**

**RTDNA Production (relative to Eco1): undetermined**

**Bacterial Editing: undetermined**

**Phage Editing: undetermined**

**Human Editing (demultiplexed): 5.705304 percent precise (49.14510169x Eco1)**

**ncRNA:**

**GGAATGGCTTACAGAAGCTAGTATGTTAGAGTGGAGTCGCTCTCATGATAAGGTCTGGGTTAACC
CAGTTCATGGAAATTACTGTAACCAGTAATTTCCATGATGTGGATAATAAAATCTCACAGCGAATT
TTTTCGCTGCTCGATTTTATTATCTGGATTGGACCAGCAAATTTTTGCGTTTTCGCAAAATTTTGCCG
TCCCAATCCACAGGCTAAAGTATTGAAGGTATCTTCTGTAAGCCATTTC**

**no RTDNA sequencing data**

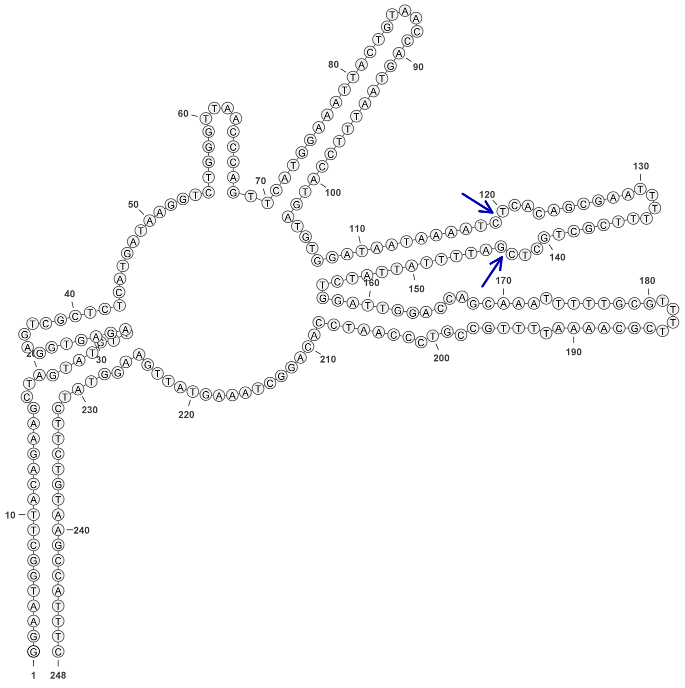


**1438**

### Retron Terminal 1450: Proteobacteria

**Bacterial Vector for RTDNA: pSLS.646**

**Bacterial Vector for Editing: None**

**Phage Vector for Editing: None**

**Human Vector for Editing: pSLS.885**

**RTDNA Production (relative to Eco1): 0.804059071 by PAGE**

**Bacterial Editing: undetermined**

**Phage Editing: undetermined**

**Human Editing (demultiplexed): 0 percent precise (0x Eco1)**

**ncRNA:**

**GCCTGCAGAGGCTAGTACGTTGGAGTGTAGTCGCTCCCCGGGTAAGGTTCCCGATAACCAAGTTG
TAGTAACTTCAGCGATGAAGCGTGCTGAAGTTACTACACTTACAAAACGCTTCGCGTTTTGACGTG
GTCCACCCATCCGCAATTTCGCTGCGCGAAATTACAGATGGGTAAAACAAGTCAAAAGCAAAGCT
GCTAAAGTAAAGAAGGTAGCCTCTGCAGGT**

**RTDNA:**

**CTTCTTTACTTTAGCAGCTTTGCTTTTGACTTGTTTTACCCATCTGTAATTTCGCGCAGCGAAATTGC
GGATGGGTGGACCACGTCAAAACGCGAAGCGTTTTGTAAGTGTAGTAACTTCAGCACGCTTCATC
GCTGAAGTTACTACAACTTGGTTATCGGGAAC**

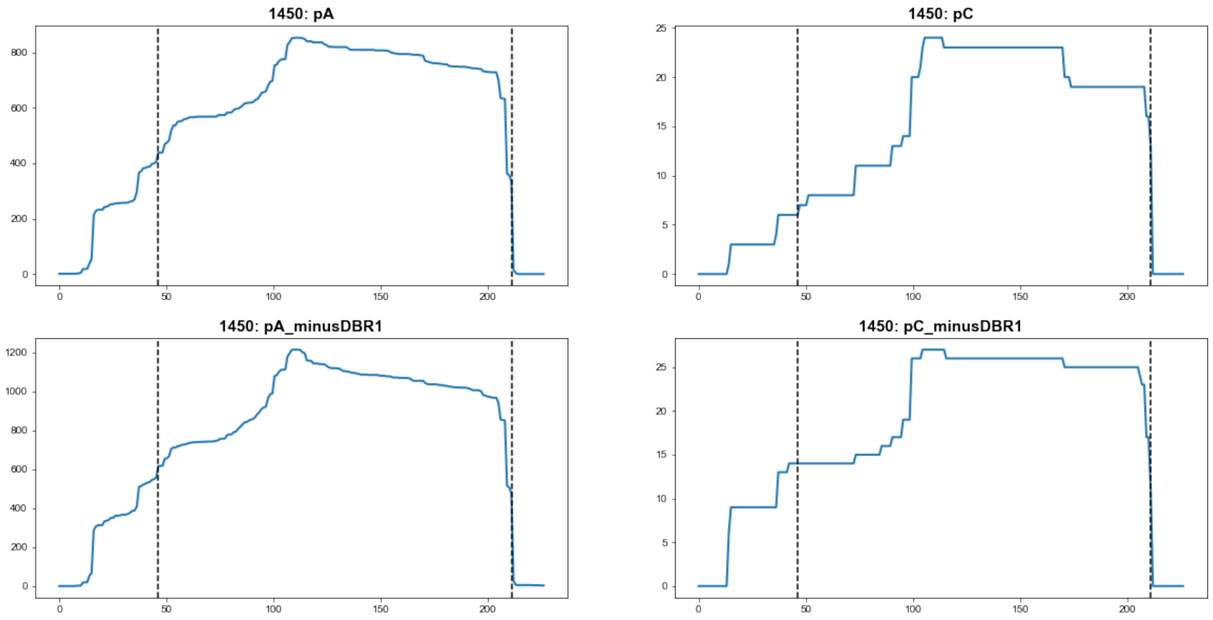


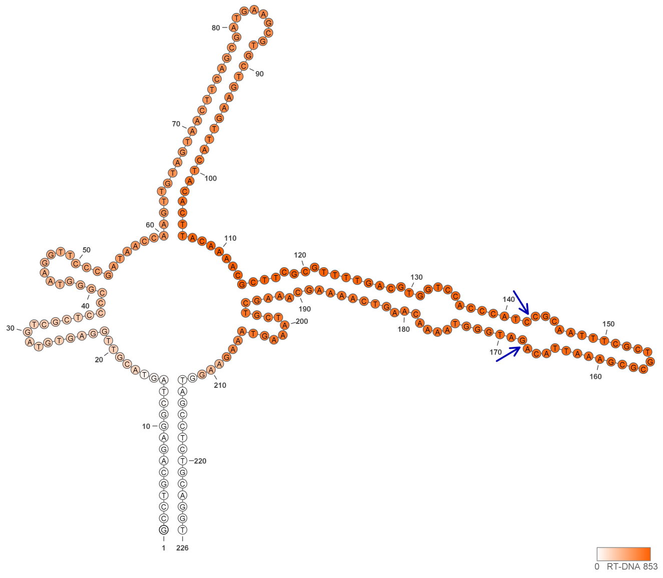


**1450**

### Retron Terminal 1453: Proteobacteria

**Bacterial Vector for RTDNA: pSLS.633**

**Bacterial Vector for Editing: None**

**Phage Vector for Editing: None**

**Human Vector for Editing: pSLS.864 (U6)**

**RTDNA Production (relative to Eco1): 1.409049692 by PAGE**

**Bacterial Editing: undetermined**

**Phage Editing: undetermined**

**Human Editing (demultiplexed): 0.196724 percent precise (1.694567193x Eco1)**

**ncRNA:**

**AGCTTACAGAAGCTAGTACGTTGGAGTGTAGTCGCTCCCCGGGCTAAGGTGCCCGTTAGCCAAGA CATACAAAAGCAAAAGCGAAAGGAAGCTTTTATTTTTGTATGTACCTAAAGATTTGAAAAAAGACT CAAATCTTTAGGTAGTACTAAAATCAATCACTTTTTGCGCACGCGCAAAAAGCAATTGATTTATAGT ACGCTTGCAAGTACAGAAGGTAGTTTCTGTAAGCT**

**RTDNA:**

**CTTCTGTACTTGCAAGCGTACTATAAATCAATTGCTTTTTGCGCGTGCGCAAAAAGTGATTGATTTT AGTACTACCTAAAGATTTGAGTCTTTTTTCAAATCTTTAGGTACATACAAAAATAAAAGCTTCCTTT CGCTTTTGCTTTTGTATGTCTTGGCTAACGGGCAC**

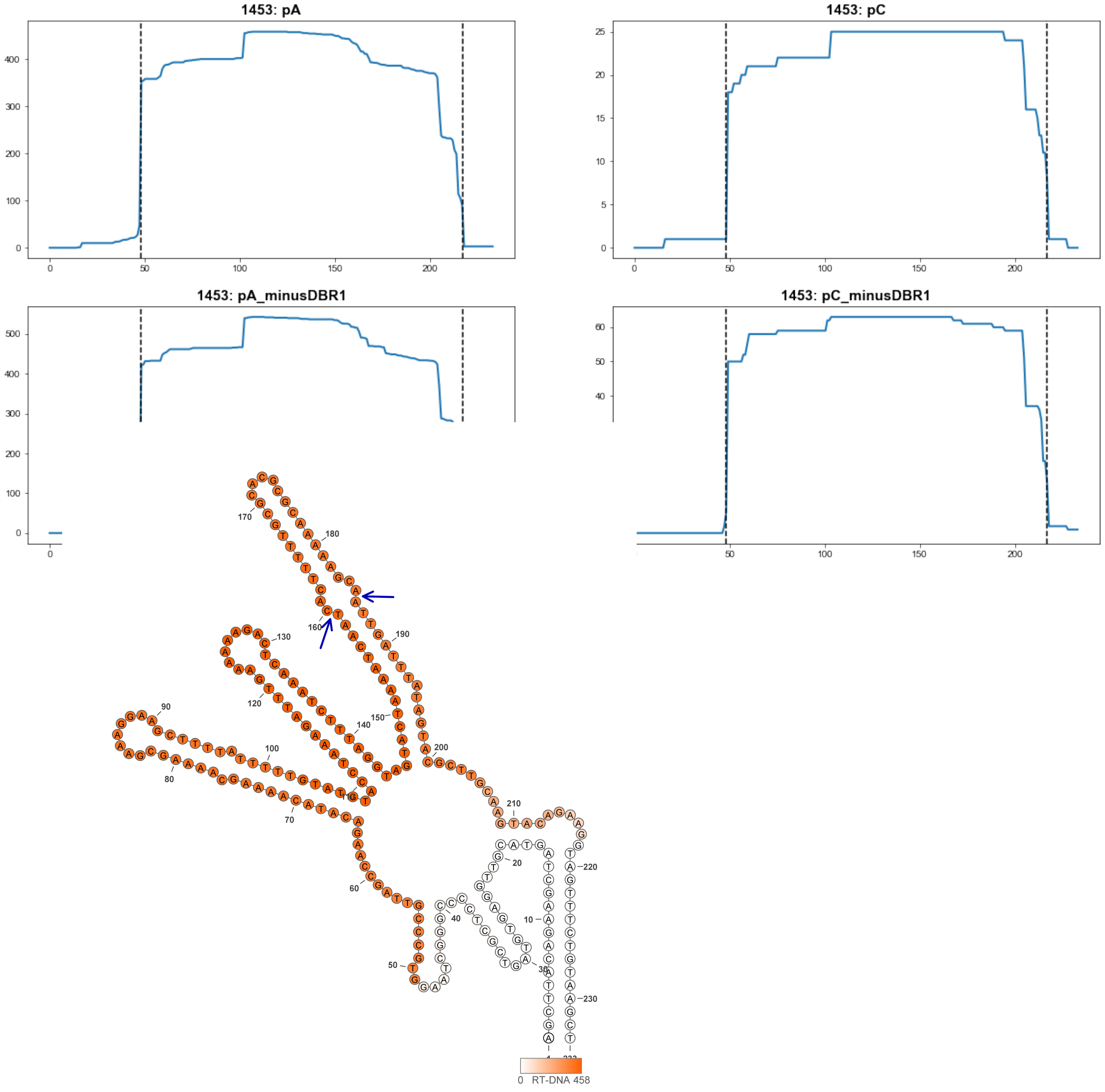


**1453**

### Retron Terminal 1467: Bacteroidetes

**Bacterial Vector for RTDNA: None**

**Bacterial Vector for Editing: None**

**Phage Vector for Editing: None**

**Human Vector for Editing: pSLS.937**

**RTDNA Production (relative to Eco1): undetermined**

**Bacterial Editing: undetermined**

**Phage Editing: undetermined**

**Human Editing (demultiplexed): 17.27088 percent precise (148.7701889x Eco1)**

**ncRNA:**

**GAGGATGGGGAATGAAACTAGCACGTTGGAGGTGTATTCGCCTCTTAGATTAACGCTGGCGTTAA**
**CCATACTAGTGCCTTAAAAGGCACTAGTATAGTTCCTAACGGAACTATACTACTAAATCTACTTCG**
**TAGTATTTAGCAGTTAGAATAAATTACAATGAAAGTAGTTTTCCCATCCTC**

**no RTDNA sequencing data**

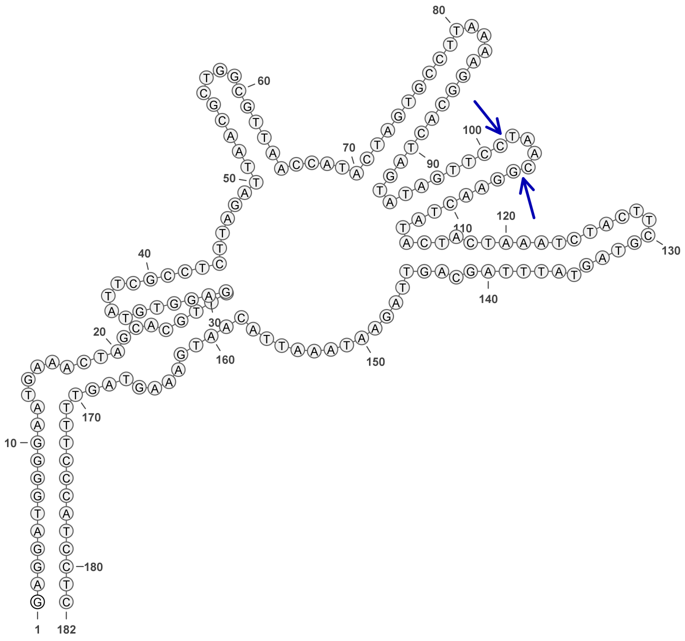

**1467**

### Retron Terminal 1484: Proteobacteria

**Bacterial Vector for RTDNA: pSLS.620**

**Bacterial Vector for Editing: None**

**Phage Vector for Editing: None**

**Human Vector for Editing: pSLS.841**

**RTDNA Production (relative to Eco1): 0 by PAGE**

**Bacterial Editing: undetermined**

**Phage Editing: undetermined**

**Human Editing (demultiplexed): 0.005402 percent precise (0.046532462x Eco1)**

**ncRNA:**

**GGCTGCTCACTGCTAGTGAGTATAGGTGTATCGCCTATCATTATGTTGAGGTTTATTTTAACCTCAT TACATTCTAAAAAGACGTTGTTGTTCGTGGCGGTGAGTAGCC**

**RTDNA:**

**ACTCACCGCCACGAACAACAACGTCTTTTTAGAATGTAATGAGGTTAAAATAAACCTCAACATAAT GATAGGCGATACACCTATACTC**

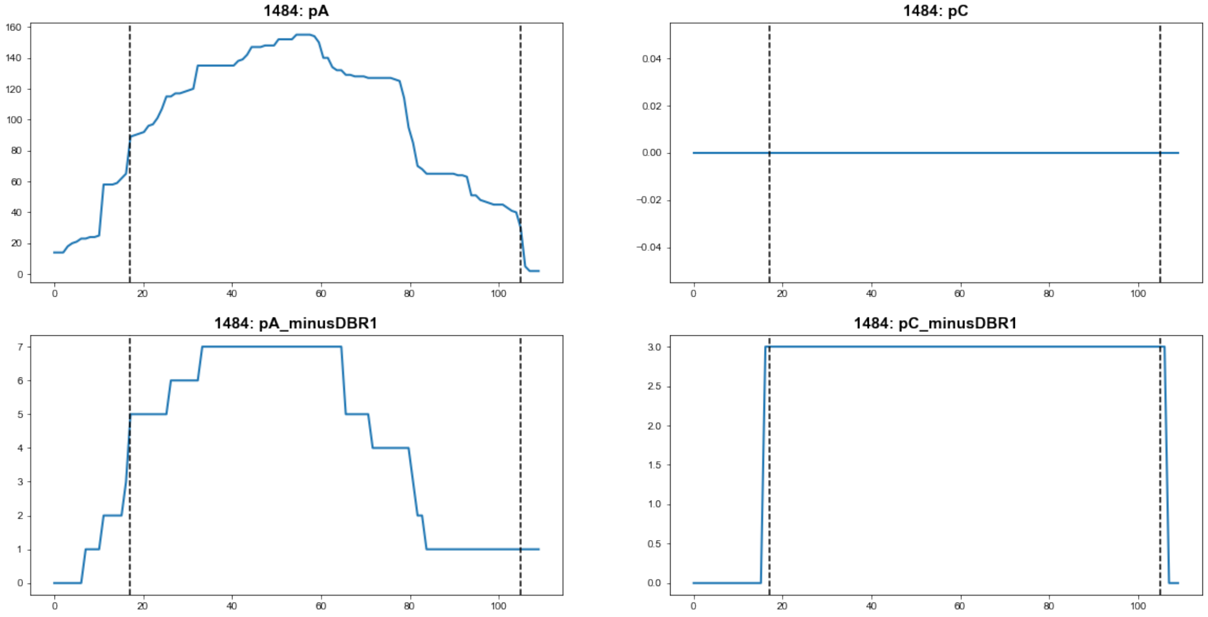


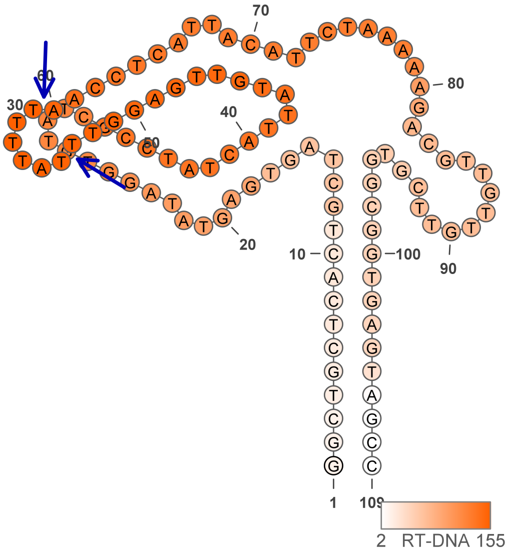


1484

### Retron Terminal 1485: Proteobacteria

**Bacterial Vector for RTDNA: pSLS.621**

**Bacterial Vector for Editing: None**

**Phage Vector for Editing: None**

**Human Vector for Editing: pSLS.842**

**RTDNA Production (relative to Eco1): 0.009793254 by PAGE**

**Bacterial Editing: undetermined**

**Phage Editing: undetermined**

**Human Editing (demultiplexed): 0.010758 percent precise (0.092668682x Eco1)**

**ncRNA:**

**GCTACTCACCGTTAGTTAGAGCGGGTGATTCGCCCGCCATCAATAAGGCTCACTTTAACCTCATTG ATTCTCTACCGGGTTTCGTGACGGTGGGTAGC**

**RTDNA:**

**GAAACCCGGTAGAGAATCAATGAGGTTAAAGTGAGCCTTATTGATGGCGGGCGAATCACCCGCTC TAACTAAC**

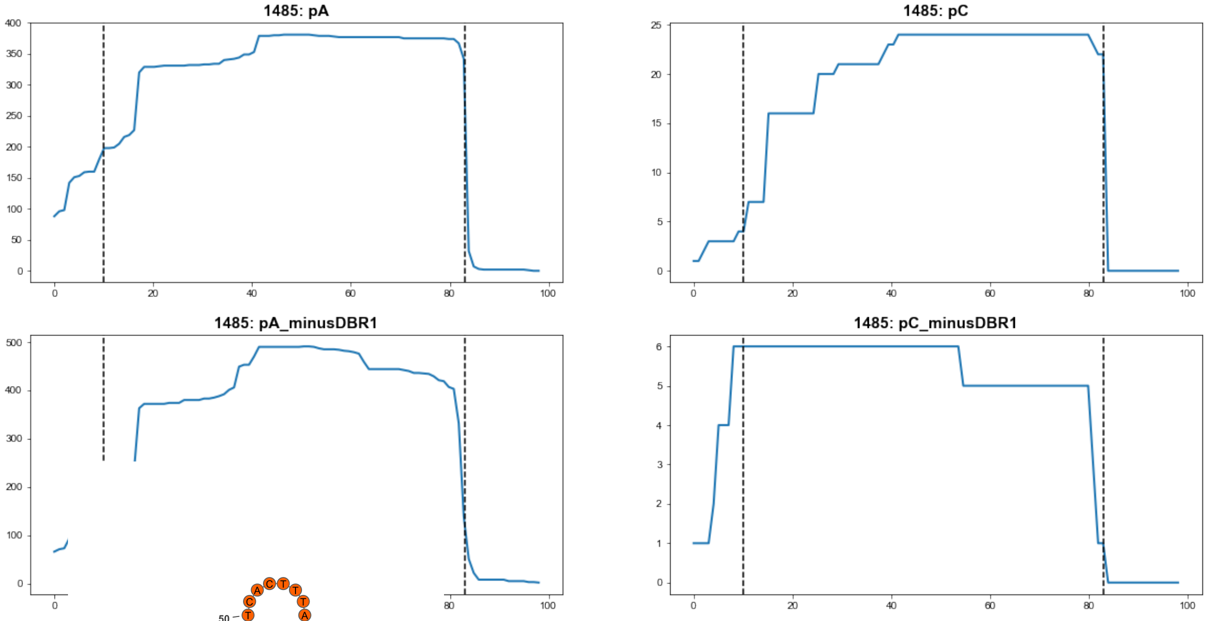


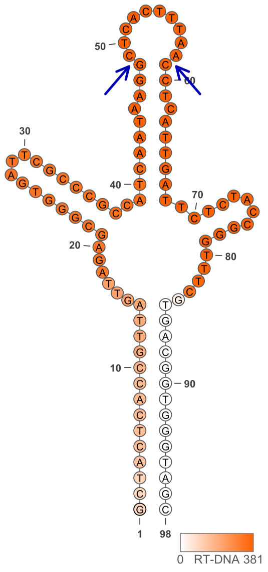


1485

### Retron Terminal 1488: Proteobacteria

**Bacterial Vector for RTDNA: None**

**Bacterial Vector for Editing: None**

**Phage Vector for Editing: None**

**Human Vector for Editing: pSLS.966**

**RTDNA Production (relative to Eco1): undetermined**

**Bacterial Editing: undetermined**

**Phage Editing: undetermined**

**Human Editing (demultiplexed): 0 percent precise (0x Eco1)**

**ncRNA:**

**TAGCTACTCAGTATTAGCGTTGGAGCTAAGCTCCCATGGGTGATTCGCCTATGTCGTTAAGGGTTT**

**TTTTAACCTCACTACGTTCTATTAAATATTTAAAAGACTTTCGTGATACTGAGTAGCTA**

**no RTDNA sequencing data**

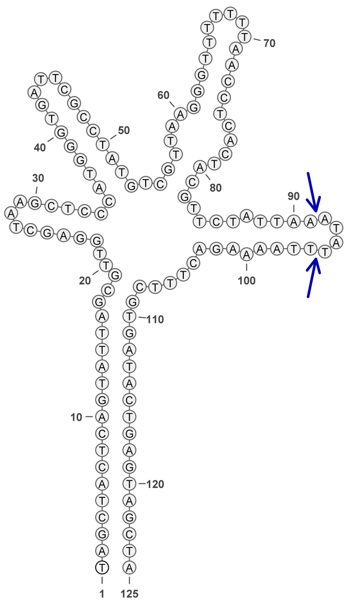


1488

### Retron Terminal 1498: Bacteroidetes

**Bacterial Vector for RTDNA: pSLS.634**

**Bacterial Vector for Editing: None**

**Phage Vector for Editing: None**

**Human Vector for Editing: None**

**RTDNA Production (relative to Eco1): 0.014145811 by PAGE**

**Bacterial Editing: undetermined**

**Phage Editing: undetermined**

**Human Editing (demultiplexed): undetermined**

**ncRNA:**

**TGTTCAAAGGGCTGACCTAAAGAGTTAGCTATGTTATCTGGAAACAGATATTCTCGGTGCCTTGTG
CCGAGAGATTTTAAGGATCCTTTAGTACCAATATTGGTCTACATTCTGCTTGAAAAATTATCATTGC
AGCCGCCGGATTCCTTCCAATAACAATTCTCCAAGAAGAATACATATCATACACCCAAGTAAGAA
GTAACTATGAAGTCATTCCCCTTTGAACA**

**RTDNA:**

**TCTTACTTGGGTGTATGATATGTATTCTTCTTGGAGAATTGTTATTGGAAGGAATCCGGCGGCTGC
AATGATAATTTTTCAAGCAGAATGTAGACCAATATTGGTACTAAAGGATC**

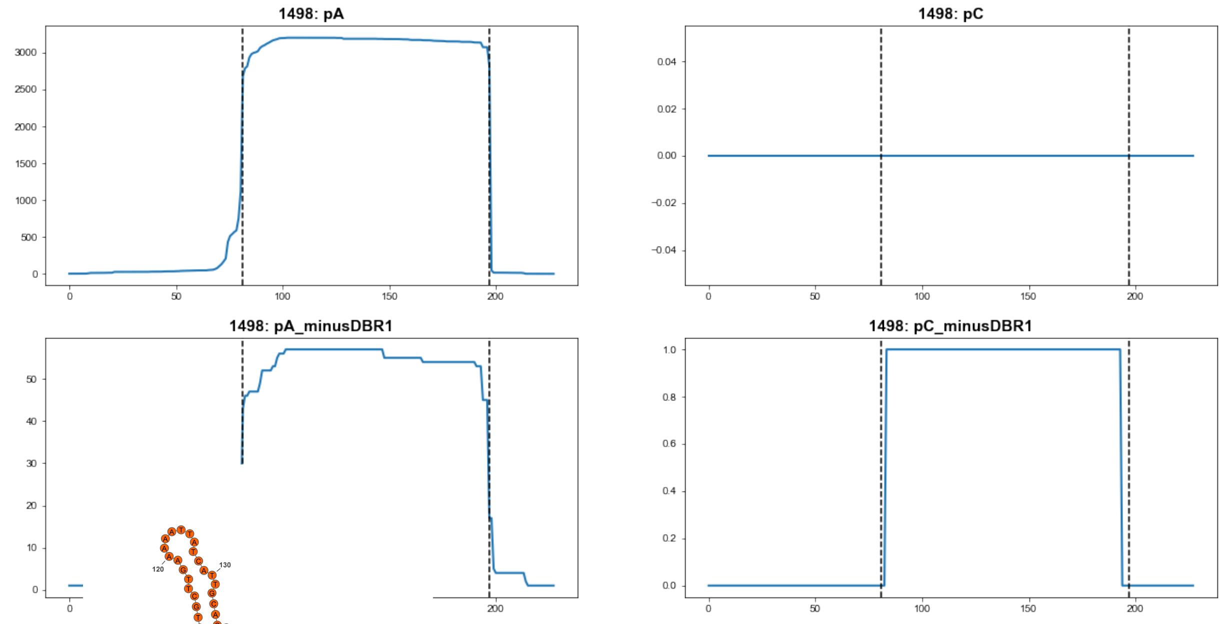


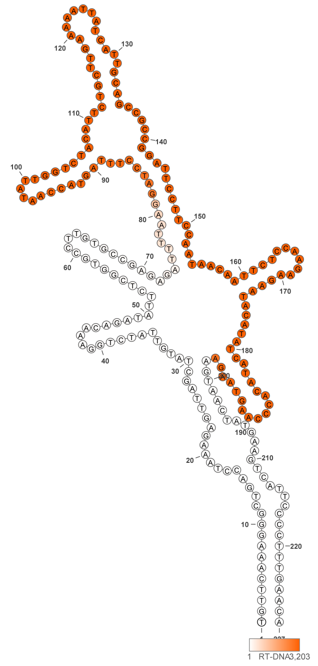


1498

### Retron Terminal 1499: Bacteroidetes

**Bacterial Vector for RTDNA: None**

**Bacterial Vector for Editing: None**

**Phage Vector for Editing: None**

**Human Vector for Editing: pSLS.967 (U6)**

**RTDNA Production (relative to Eco1): undetermined**

**Bacterial Editing: undetermined**

**Phage Editing: undetermined**

**Human Editing (demultiplexed): 0 percent precise (0x Eco1)**

**ncRNA:**

**ATTTGTTAGCATTTAACTGTAAAAGGTTTTCTCAGTGCCTTGTTGCTGAGAATTATTAAGGAGTCTT**
**TTTAAAGCAGCATAAAAATACTATGCATAATAGATATTAAATGTTGCCATTTCTAATCCTTACTAAC**
**ATTTAATATCCGTTATGAATATTATCAAACAACCGAGTAAGAGTAACAAAT**

**no RTDNA sequencing data**

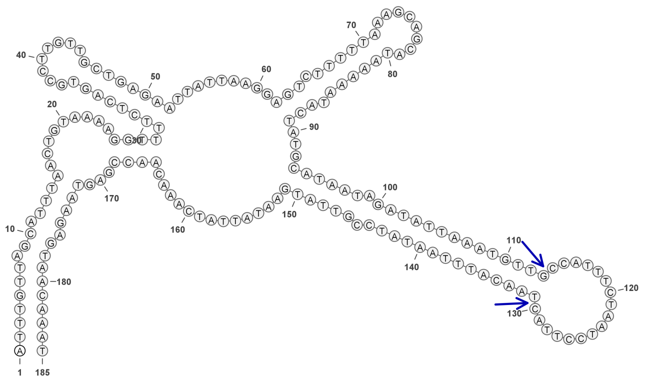

**1499**

### Retron Terminal 1502: Bacteroidetes

**Bacterial Vector for RTDNA: None**

**Bacterial Vector for Editing: None**

**Phage Vector for Editing: None**

**Human Vector for Editing: pSLS.938**

**RTDNA Production (relative to Eco1): undetermined**

**Bacterial Editing: undetermined**

**Phage Editing: undetermined**

**Human Editing (demultiplexed): 0 percent precise (0x Eco1)**

**ncRNA:**

**AATGGAAGGTCATAAATGGCAAGCAATTTCATGTCAATACCTATTAGGGAAGGACCTGATTTCCTG**

**GACAGTTGTCCAAGATTTTTGAAGTAACTTTTCAACCTTTACTACACAAAGTCCTCGCAGGCTGCG**

**GTTCGTGTAGAAAACGCTTCGCTAATTATATAAAAGAGTTGCCTTTATGACTATTCCATT**

**no RTDNA sequencing data**

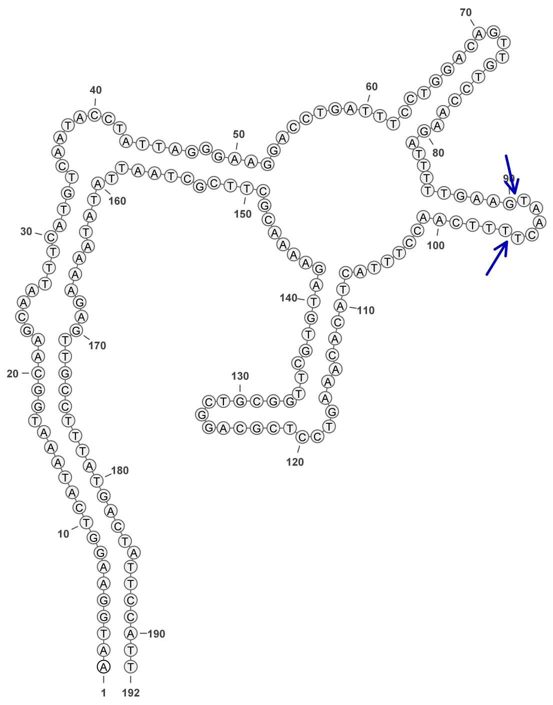

**1502**

### Retron Terminal 1513: Bacteroidetes

**Bacterial Vector for RTDNA: None**

**Bacterial Vector for Editing: None**

**Phage Vector for Editing: None**

**Human Vector for Editing: pSLS.999**

**RTDNA Production (relative to Eco1): undetermined**

**Bacterial Editing: undetermined**

**Phage Editing: undetermined**

**Human Editing (demultiplexed): 0.235361 percent precise (2.02738369x Eco1)**

**ncRNA:**

**TTCTTTAGATTAGCAGTATCTGTACAGACAGAAGCCTGCGTCCGTTTGATGCAGGAAAGTTAAAGC**

**CACATTACATGATTTCTATGTCACACCCCATGGGCGCGACACAGAAGTCGCCGAAAGGCGCTTGA**

**TAAAAGAGTAATTTGAAGAA**

**no RTDNA sequencing data**

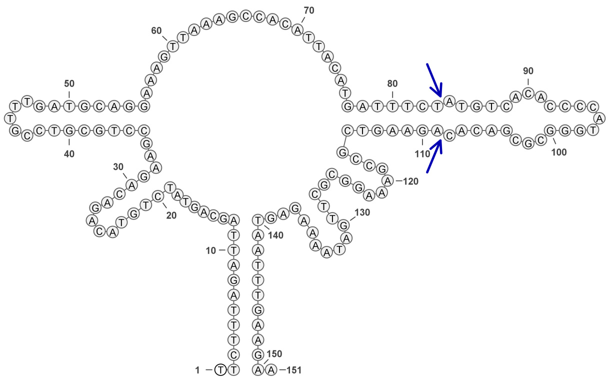


**1513**

### Retron Terminal 1520: Firmicutes

**Bacterial Vector for RTDNA: pSLS.686**

**Bacterial Vector for Editing: pAGD301**

**Phage Vector for Editing: pAGD404**

**Human Vector for Editing: pSLS.919**

**RTDNA Production (relative to Eco1): 1.624349676 by PAGE**

**Bacterial Editing: 5.03160505 percent precise (1.548287937x Eco1)**

**Phage Editing: 33.40254033 percent precise (0.823862708x Eco1)**

**Human Editing (demultiplexed): 0 percent precise (0x Eco1)**

**ncRNA:**

**ATTATTGACAACATCGTTATCCTTAGATGCTACTACTTTTGTATAAACGTGCACAATATTTATTTATA TGAAAGTAGTGGGAAGTGGTTCGCTTTCCGAGGTGCTATGTCAGGACTATATTTAATGAAGTAAGC GTCCAACTTCATATATATGAAGTTAGCCACTTACTTCATCAAAAGGAATAAAGGTAAGGAGGACG GTGTTGTCAATGAT**

**RTDNA:**

**TGACAACACCGTCCTCCTTACCTTTATTCCTTTTGATGAAGTAAGTGGCTAACTTCATATATATGAA GTTGGACGCTTACTTCATTAAATATAGTCCTGACATAGCACCTCGGAAAGCGAACCACTTCCCACT ACTTTCATATAAATAAATATTGTGCACGTTTATAC**

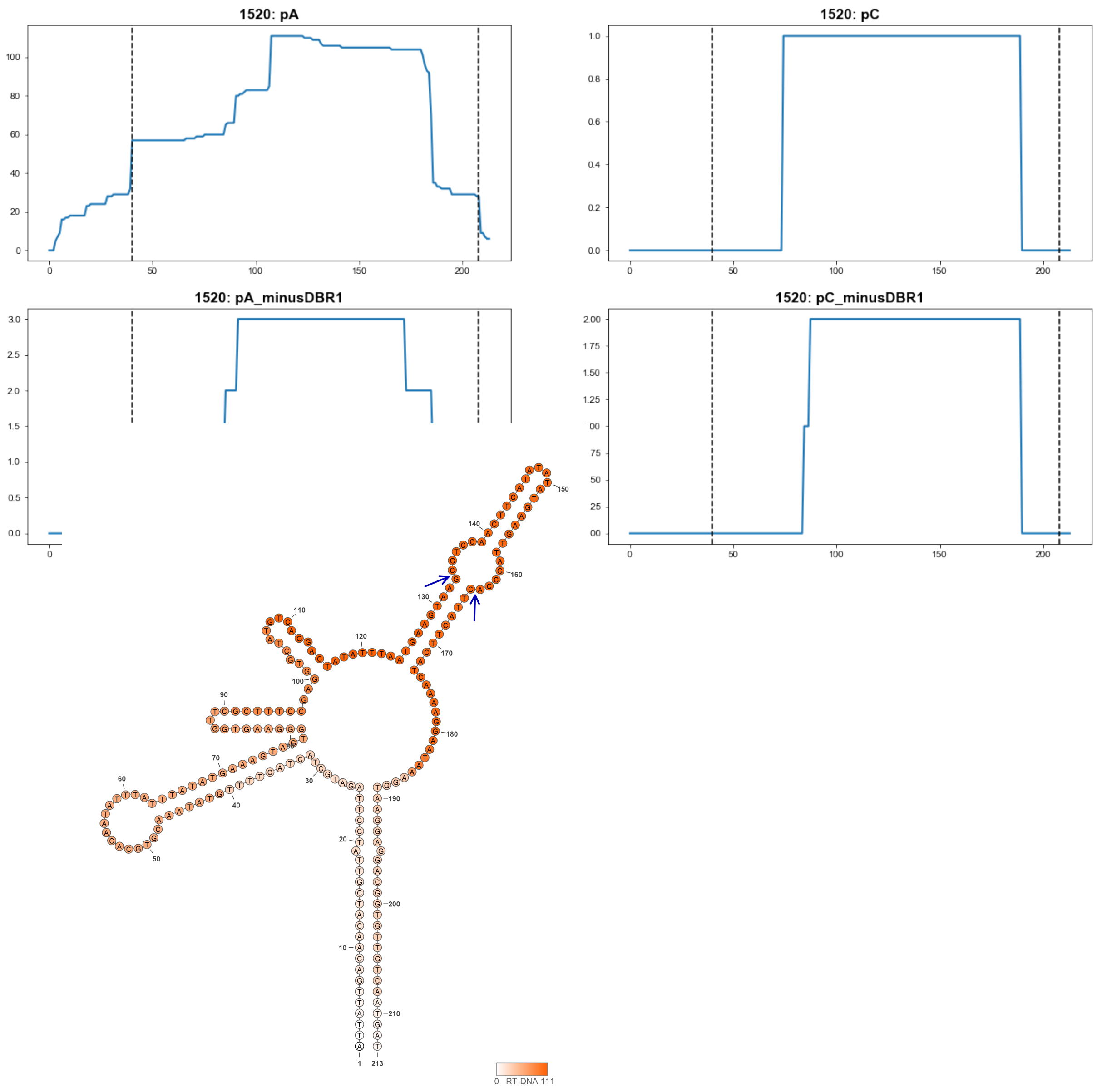


**1520**

### Retron Terminal 1525: Firmicutes

**Bacterial Vector for RTDNA: None**

**Bacterial Vector for Editing: None**

**Phage Vector for Editing: None**

**Human Vector for Editing: pSLS.1000**

**RTDNA Production (relative to Eco1): undetermined**

**Bacterial Editing: undetermined**

**Phage Editing: undetermined**

**Human Editing (demultiplexed): 0.080013 percent precise (0.689226555x Eco1)**

**ncRNA:**

**ATTTCTATAAATCATCCCATTCTTAGAATTGTTGTTAATATGCATTACATAACAACTAGGGGTTGGT**
**TCGACCTCGGAAGAATAATAAACACATGAAATTAGCTAAATCAACGGTGAACTTCCACGACAGTG**
**AAACTCCCCCGTTGATTTACCCAAAAACATATTGAGTAAGAATGGGGTGGTTTGTATGAAT**

**no RTDNA sequencing data**

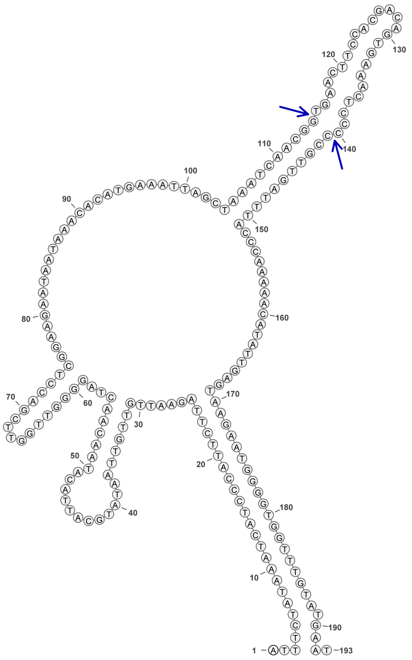

1525

### Retron Terminal 1531: Firmicutes

**Bacterial Vector for RTDNA: None**

**Bacterial Vector for Editing: None**

**Phage Vector for Editing: None**

**Human Vector for Editing: pSLS.939**

**RTDNA Production (relative to Eco1): undetermined**

**Bacterial Editing: undetermined**

**Phage Editing: undetermined**

**Human Editing (demultiplexed): 9.808371 percent precise (84.48864253x Eco1)**

**ncRNA:**

**ATTAGTACTGCTTTCTTAGAACGTTACAGATACATATCTGTTTACGGAAGATGGTTCTGTCTTCCTA**
**AGAAAATTAACAAACACAGAGTTACTCCAATCAATGGAGTAACTCCACGACAGTGAACTTACTCC**
**ATTGATTGGAGACAAAGAGATTTAAGGTAAGAAAGCGGTGCTAAT**

**no RTDNA sequencing data**

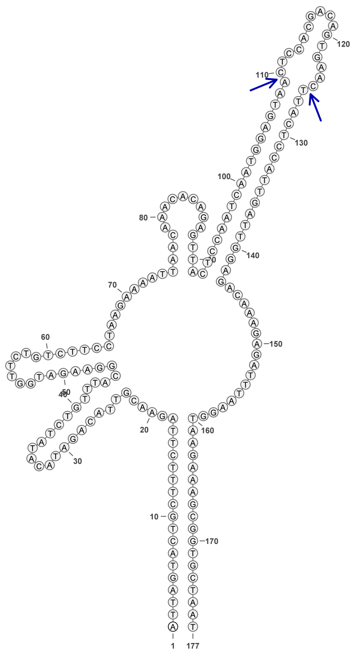

**1531**

### Retron Terminal 1548: Elusimicrobia

**Bacterial Vector for RTDNA: None**

**Bacterial Vector for Editing: None**

**Phage Vector for Editing: None**

**Human Vector for Editing: pSLS.1001**

**RTDNA Production (relative to Eco1): undetermined**

**Bacterial Editing: undetermined**

**Phage Editing: undetermined**

**Human Editing (demultiplexed): 0.047284 percent precise (0.407301169x Eco1)**

**ncRNA:**

**GAGAGCCGTGCCAATTTAGAAAGAGAGGGGGTGGTGCCTTCGCCGCCCTAGGTGCAGAACATCC**

**TAGAAGTTCCGCTAGAATTCAGCCAATCTCGCAGTAGCAGCGATTGGATGAATTCTAGCAGTATGT**

**TTGTGAGTAAATTGGCACGGCTCTC**

**no RTDNA sequencing data**

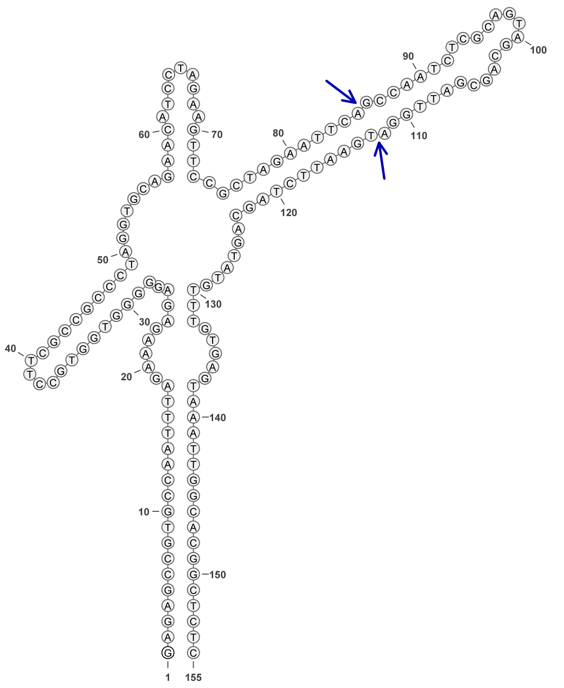


**1548**

### Retron Terminal 1550: Proteobacteria

**Bacterial Vector for RTDNA: pSLS.487**

**Bacterial Vector for Editing: pSLS.491**

**Phage Vector for Editing: pAGD405**

**Human Vector for Editing: pSLS.801 (H1)**

**RTDNA Production (relative to Eco1): 1 by PAGE**

**Bacterial Editing: 3.24978638 percent precise (1x Eco1)**

**Phage Editing: 40.54381877 percent precise (1x Eco1)**

**Human Editing (demultiplexed): 0.116091 percent precise (1x Eco1)**

**ncRNA:**

**TGCGCACCCTTAGCGAGAGGTTTATCATTAAGGTCAACCTCTGGATGTTGTTTCGGCATCCTGCAT TGAATCTGAGTTACTGTCTGTTTTCCTTGTTGGAACGGAGAGCATCGCCTGATGCTCTCCGAGCCA ACCAGGAAACCCGTTTTTTCTGACGTAAGGGTGCGCA**

**RTDNA:**

**GTCAGAAAAAACGGGTTTCCTGGTTGGCTCGGAGAGCATCAGGCGATGCTCTCCGTTCCAACAAG GAAAACAGACAGTAACTCAG**

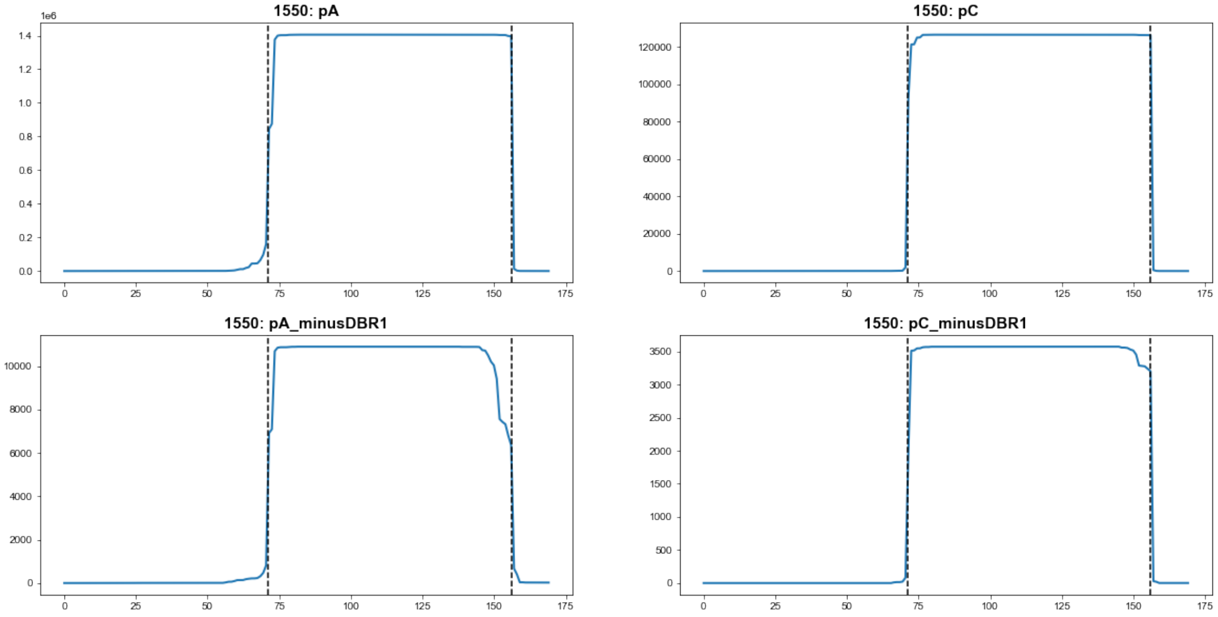


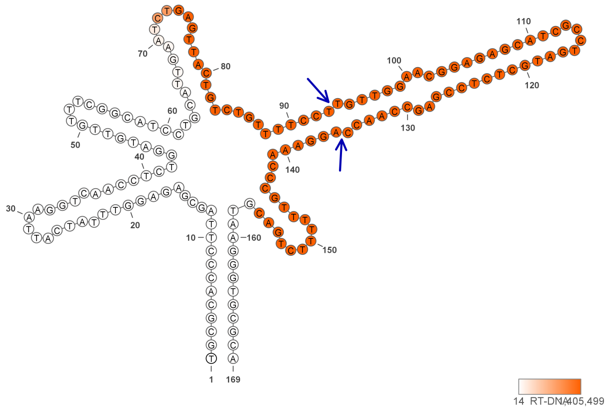

**1550**

### Retron Terminal 1550.1: Proteobacteria

**Bacterial Vector for RTDNA: None**

**Bacterial Vector for Editing: None**

**Phage Vector for Editing: None**

**Human Vector for Editing: pSLS.802 (U6)**

**RTDNA Production (relative to Eco1): 1 by PAGE**

**Bacterial Editing: undetermined**

**Phage Editing: undetermined**

**Human Editing (demultiplexed): undetermined**

**ncRNA:**

**TGCGCACCCTTAGCGAGAGGTTTATCATTAAGGTCAACCTCTGGATGTTGTTTCGGCATCCTGCAT TGAATCTGAGTTACTGTCTGTTTTCCTTGTTGGAACGGAGAGCATCGCCTGATGCTCTCCGAGCCA ACCAGGAAACCCGTTTTTTCTGACGTAAGGGTGCGCA**

**RTDNA:**

**GTCAGAAAAAACGGGTTTCCTGGTTGGCTCGGAGAGCATCAGGCGATGCTCTCCGTTCCAACAAG GAAAACAGACAGTAACTCAG**

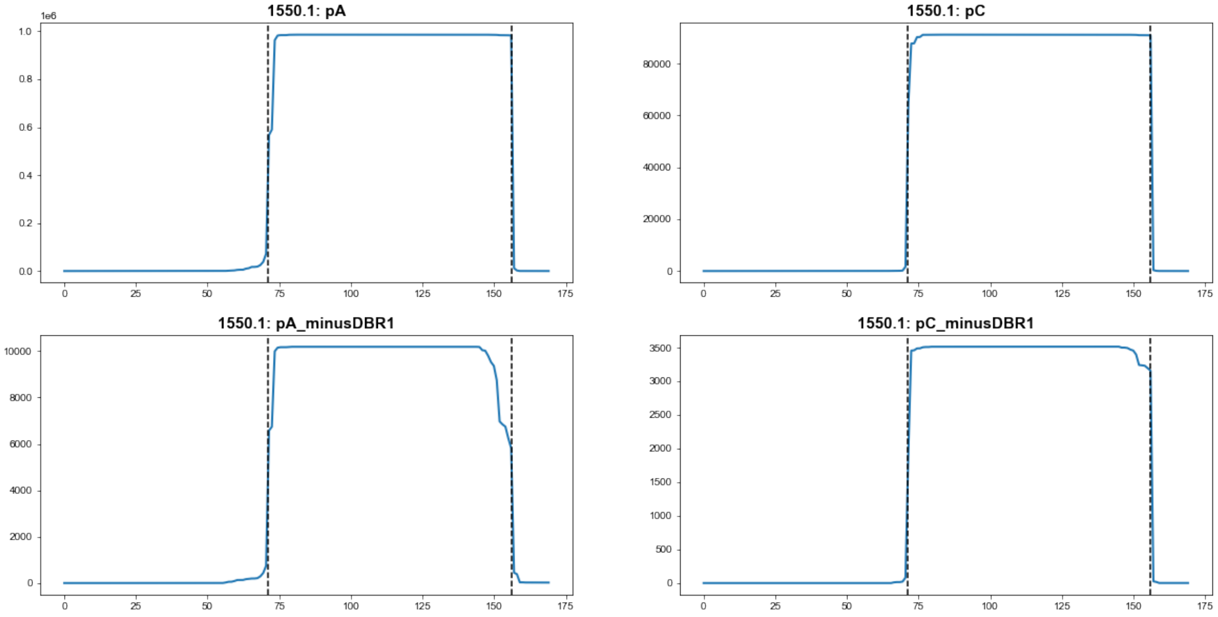


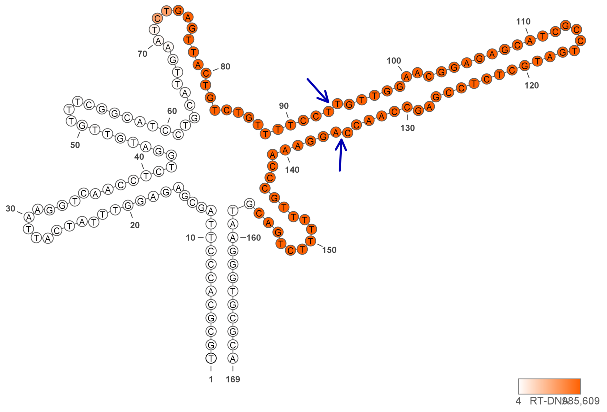

**1550.1**

### Retron Terminal 1552: Proteobacteria

**Bacterial Vector for RTDNA: None**

**Bacterial Vector for Editing: None**

**Phage Vector for Editing: None**

**Human Vector for Editing: pSLS.1002**

**RTDNA Production (relative to Eco1): undetermined**

**Bacterial Editing: undetermined**

**Phage Editing: undetermined**

**Human Editing (demultiplexed): 0.001801 percent precise (0.015513692x Eco1)**

**ncRNA:**

**GTTCAAAAAAAATCACCCGTAAAGAAAAGTATAGATAATATATTGATGGCGGATAGATTTATCTTG
CCATGCATATACCTTATGGATGGTATCAATAATGTATTGCTTTATGACTTGTTAAGTAAAGCTACAG
GAAAGGCAAATACATTGTGCGAAATTATTGTAAAATCTGCATTATCAAAACTGATGAAAGAGCGA
GTTATTGTTAGGACATCAGAAGGTTATCACATTACAGACGCTGGTATGGAATATGTACTACTTAGG
TTTGATCGGCTAGTTTTAGACAAGCTAAGACTTGAAATGATGAACTTTGAAAATCGTAGCAATATA
TCATTGAAC**

**no RTDNA sequencing data**

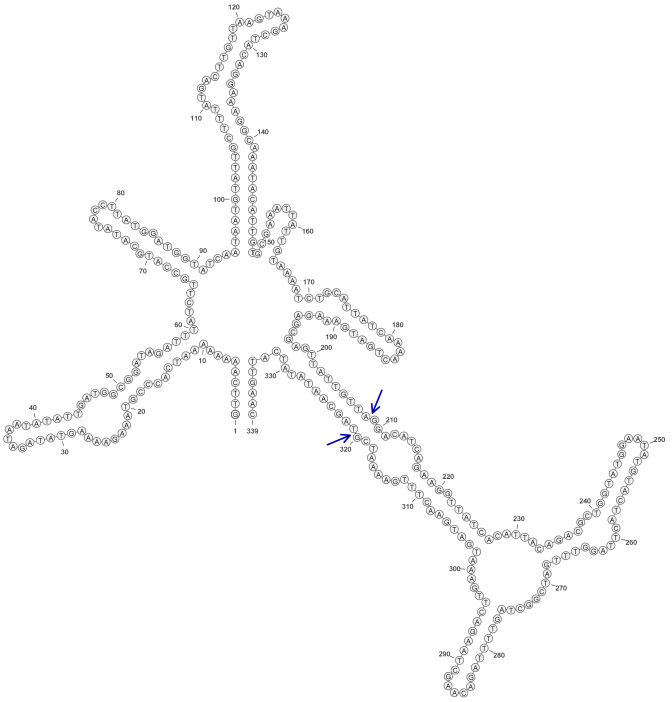

**1552**

### Retron Terminal 1578: Firmicutes

**Bacterial Vector for RTDNA: pSLS.687**

**Bacterial Vector for Editing: None**

**Phage Vector for Editing: None**

**Human Vector for Editing: pSLS.920**

**RTDNA Production (relative to Eco1): 0.014266788 by PAGE**

**Bacterial Editing: undetermined**

**Phage Editing: undetermined**

**Human Editing (demultiplexed): 0 percent precise (0x Eco1)**

**ncRNA:**

**AATAGTCAGTAGTCTTCTAGATATGGTAGAGAGTCTCTATCGTTATTAATGTTACTTAATTGATATG
TGAAAAGCAGGAAATCGCTACTTTTCACAACTACAACAAAAATCGTTGTAGTTGCAAATAATTTAT
GAAAACCACATCAATAATTTGCAGATTTTCAAAACAAGTTAAAGGTATTTTAAAAAACTAGATCAG
AGTAGAAGCTGCTGACTATT**

**RTDNA:**

**CTGATCTAGTTTTTTAAAATACCTTTAACTTGTTTTGAAAATCTGCAAATTATTGATGTGGTTTTCAT
AAATTATTTGCAACTACAACGATTTTTGTTGTAGTTGTGAAAAGTAGCGATTTCCTGCTTTTCACAT
ATCAATTAAGTAAC**

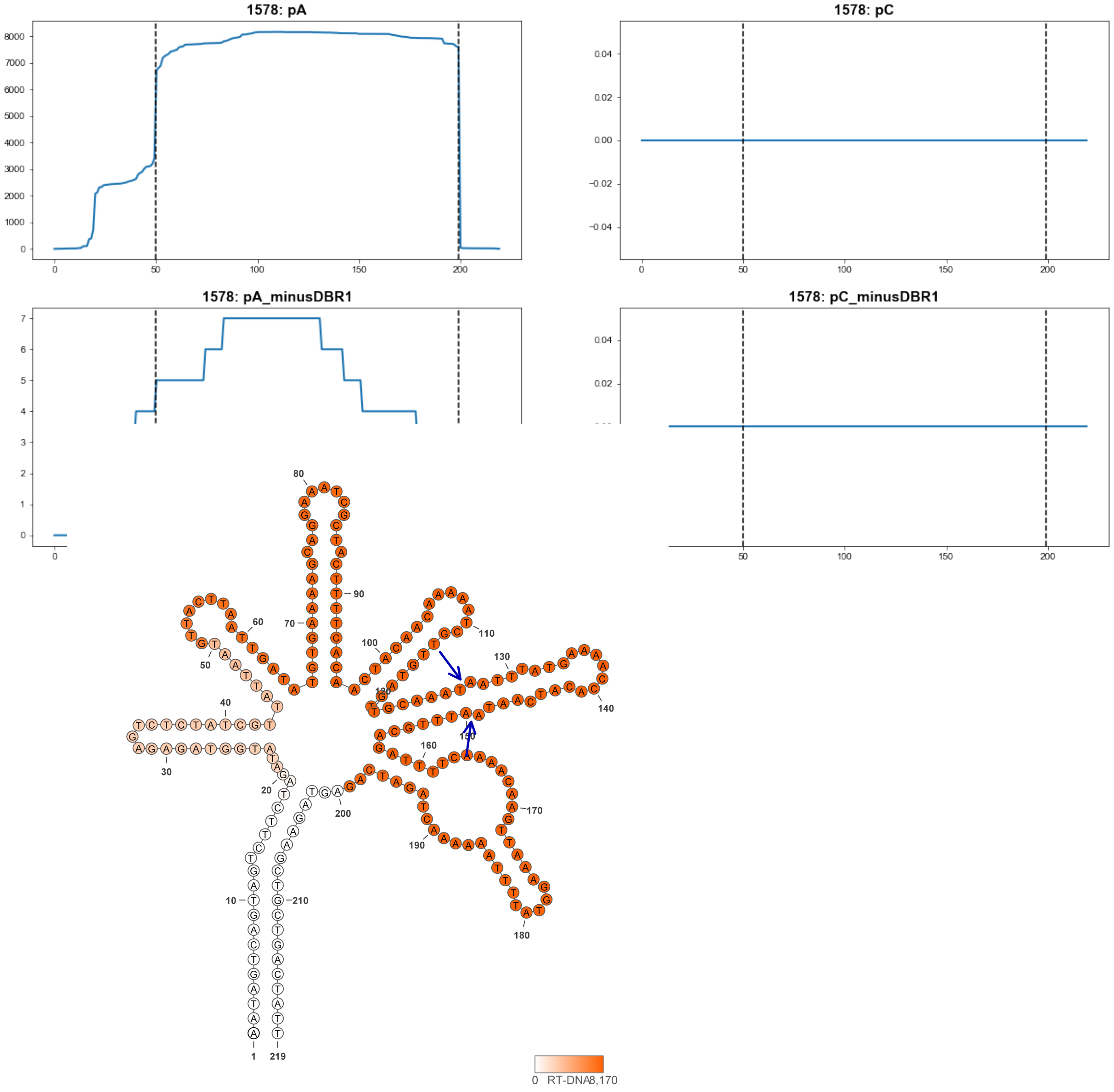


**1578**

### Retron Terminal 1595: Firmicutes

**Bacterial Vector for RTDNA: None**

**Bacterial Vector for Editing: None**

**Phage Vector for Editing: None**

**Human Vector for Editing: pSLS.940**

**RTDNA Production (relative to Eco1): undetermined**

**Bacterial Editing: undetermined**

**Phage Editing: undetermined**

**Human Editing (demultiplexed): 0.749506 percent precise (6.456193848x Eco1)**

**ncRNA:**

**GTTGTTTTCTAGAATTGAGCAGCGAGTCGCTGTTCGTGAAGTAAAGTTACTATGTATCTATGATACT
AATTTTCAAGAGCGTAAAATTAGTATCATACAACAAAGAAAATCACTTTGTTGTATACACCAGTAA
ACGTGGTGTATAGGTAAATTAAATAATCCAGTAGAAAATAAC**

**no RTDNA sequencing data**

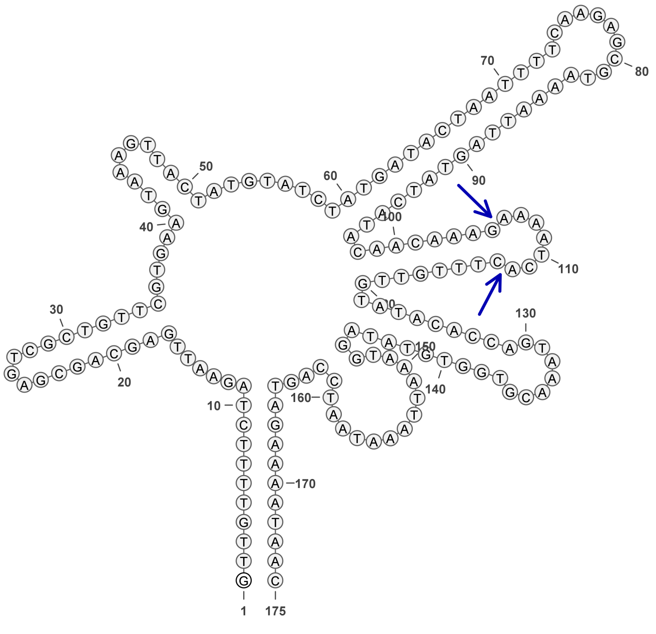

**1595**

### Retron Terminal 1609: Firmicutes

**Bacterial Vector for RTDNA: pSLS.688**

**Bacterial Vector for Editing: pAGD298**

**Phage Vector for Editing: None**

**Human Vector for Editing: pSLS.921**

**RTDNA Production (relative to Eco1): 2.015198951 by PAGE**

**Bacterial Editing: 1.62301306 percent precise (0.49942146x Eco1)**

**Phage Editing: undetermined**

**Human Editing (demultiplexed): 3.004598 percent precise (25.88140338x Eco1)**

**ncRNA:**

**GATAGGTTTCTTGGCCTTTATACTATGGTGGTGCGCCATGGTGGAGATTTGTTAGACACACATCAT**
**CAGGTTGCGCGCAATTCGCTACGCTACCAATACTGTGCGCTTTTAAAACCGATCCTTCGGTTAAGC**
**GACAGTATTGGTGCTACGCTGAAGTGTCACAACCAAATACAAGAATAGTTAGCAAGAAATCTATC**

**RTDNA:**

**TAACTATTCTTGTATTTGGTTGTGACACTTCAGCGTAGCACCAATACTGTCGCTTAACCGAAGGAT**
**CGGTTTTAAAAGCGCACAGTATTGGTAGCGTAGCGAATTGCGCGCAACCTGATGATGTGTGTCT**

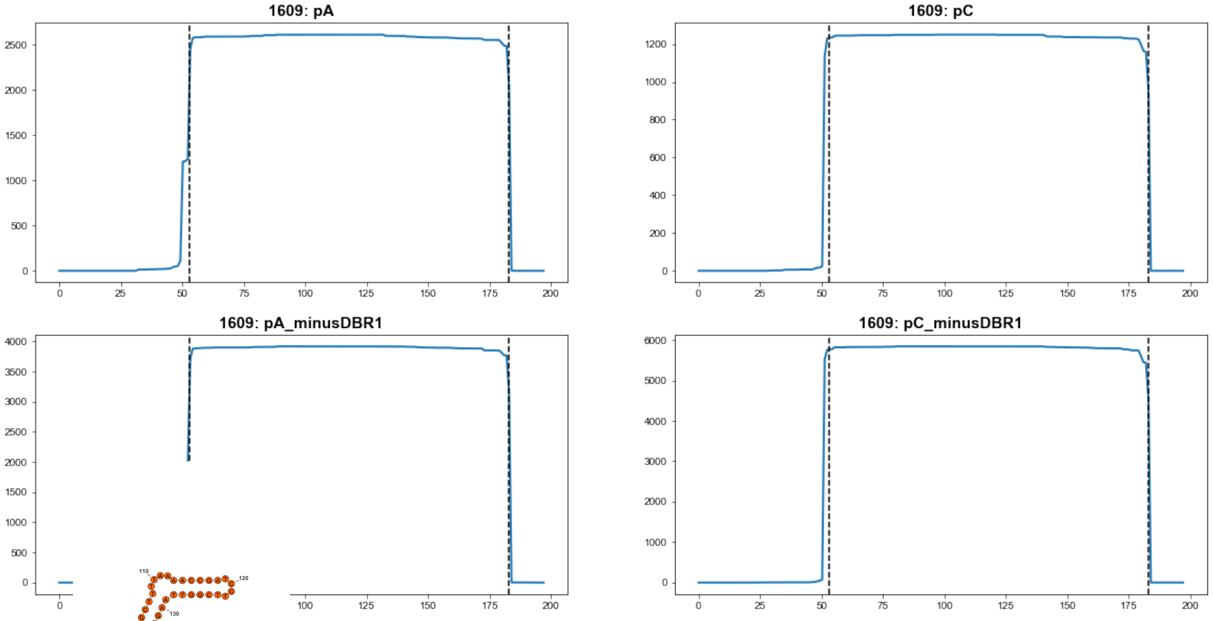


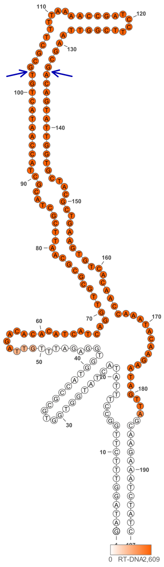

1609

### Retron Terminal 1627: Proteobacteria

**Bacterial Vector for RTDNA: None**

**Bacterial Vector for Editing: None**

**Phage Vector for Editing: None**

**Human Vector for Editing: pSLS.968**

**RTDNA Production (relative to Eco1): undetermined**

**Bacterial Editing: undetermined**

**Phage Editing: undetermined**

**Human Editing (demultiplexed): 0 percent precise (0x Eco1)**

**ncRNA:**

**GAGGCGCTCCGGACCCGCCGCAAGGGGCGCACGCGCTGAGTCAGTTGACCTGAACGCTGCCAC**
**CTTTGATGCGATTGACGCCGCTGGCGTGCGTCTCGACGTAGGTGCCCCGGAGGATGAGCTTGCCG**
**TTGCGGCGCAGGGTGATGCTCGCCTCTCCGCACTTCAGGACGATCTCGTCCGCGCCTT**

**no RTDNA sequencing data**

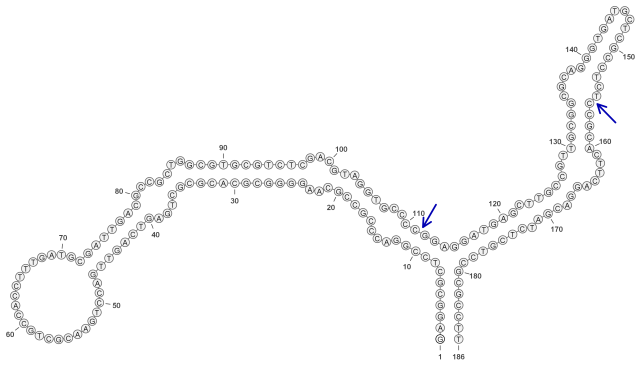

**1627**

### Retron Terminal 1630: Chloroflexi

**Bacterial Vector for RTDNA: None**

**Bacterial Vector for Editing: None**

**Phage Vector for Editing: None**

**Human Vector for Editing: pSLS.969**

**RTDNA Production (relative to Eco1): undetermined**

**Bacterial Editing: undetermined**

**Phage Editing: undetermined**

**Human Editing (demultiplexed): 0.172359 percent precise (1.484688736x Eco1)**

**ncRNA:**

**GGGTAGCTTGGTTAGCCGATTCTGCGATGGCACGTCGTGGAGGCGAATTGTTCACCACACTTCTT**
**AATTTAATCACCGAGTGATTAAATTACTTATGTCAAGGTTACGTGTAACCTTGACATAAGGTTCAGT**
**ATGTTAGTAACCAAGTTACCC**

**no RTDNA sequencing data**

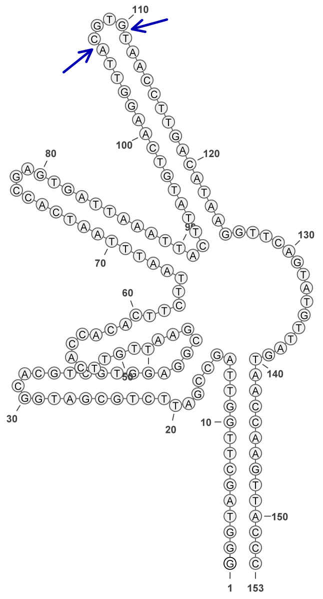


1630

### Retron Terminal 1644: Proteobacteria

**Bacterial Vector for RTDNA: None**

**Bacterial Vector for Editing: None**

**Phage Vector for Editing: None**

**Human Vector for Editing: pSLS.941**

**RTDNA Production (relative to Eco1): undetermined**

**Bacterial Editing: undetermined**

**Phage Editing: undetermined**

**Human Editing (demultiplexed): 1.460165 percent precise (12.57776227x Eco1)**

**ncRNA:**

**CTCGAGGAGGTCTGCGAGCACGAGCCCATCATGCTGCGGCGCTACGCCGTGGCGGAGCTTCGCT ACCCACCCGACCCCGCGCCCGATGCTCTGGGTTGGCGCCGCCGACGGCATGCGCCTGTCCCAAG GGCAACGCGGTAGGCTGCGCCTACCACGTCGACAGGCGCATGCCGTCGCGGCGCCGTGGGTGAT CGGACGCGGTCCCGAGCACGAAGTCGAGCTGGCAGACCTCCTCGAG**

**no RTDNA sequencing data**

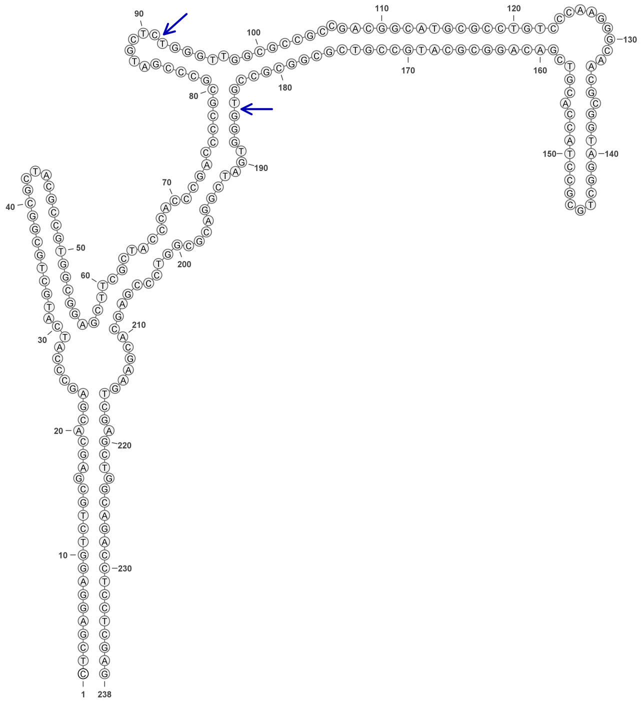

**1644**

### Retron Terminal 1648: Cyanobacteria

**Bacterial Vector for RTDNA: None**

**Bacterial Vector for Editing: None**

**Phage Vector for Editing: None**

**Human Vector for Editing: pSLS.970 (U6)**

**RTDNA Production (relative to Eco1): undetermined**

**Bacterial Editing: undetermined**

**Phage Editing: undetermined**

**Human Editing (demultiplexed): 0.537611 percent precise (4.63094469x Eco1)**

**ncRNA:**

**AATGGAAAATATCGTTGGCGGGCTCTTTTCCCATCTCTAAATCTTGAAAGTCAATCTGAAGCTGGA GTAGAGGAAAGAAAAGGTTTAGAGATATTCAAGTCAAATAAAATCAAGATAATTTCAGACCAGAT AACTTCCAAGGGTCGTGAACTTAAAGGTCTCGTAGAGTCACATGAAAAAATAAATCAACCTTCCAT T**

**no RTDNA sequencing data**

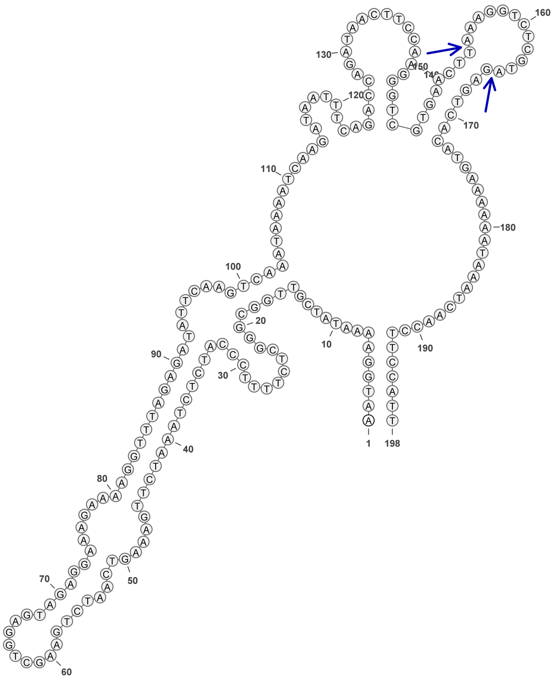

1648

### Retron Terminal 1673: Bacteroidetes

**Bacterial Vector for RTDNA: pSLS.624**

**Bacterial Vector for Editing: None**

**Phage Vector for Editing: None**

**Human Vector for Editing: pSLS.848**

**RTDNA Production (relative to Eco1): 1.969985491 by PAGE**

**Bacterial Editing: undetermined**

**Phage Editing: undetermined**

**Human Editing (demultiplexed): undetermined**

**ncRNA:**

**TTTGTAAAGAGGAATTGATTGCACTTGCGCCCGTAAAAGGGACGAATGATGTTTCGTCATTCAGGA
ATTATGACTATACATGCTTCACTAAATGCAATCTTTACTTTGGGAGGGGCGTACCTAGTGCGAATTT
ACGGTACGAGCCCCGGGGGCTGAAATTCACACTAGGTACGCCCCTCCCTGGAGTTGATTGATTTA
GTCAATGGCTTGCACGAAGGCCTTTTCCTCTTTACAAA**

**RTDNA:**

**GTGCAAGCCATTGACTAAATCAATCAACTCCAGGGAGGGGCGTACCTAGTGTGAATTTCAGCCCC
CGGGGCTCGTACCGTAAATTCGCACTAGGTACGCCCCTCCCAAAGTAAAGATTGCATTTAGTGAA
GCATGTATAGT**

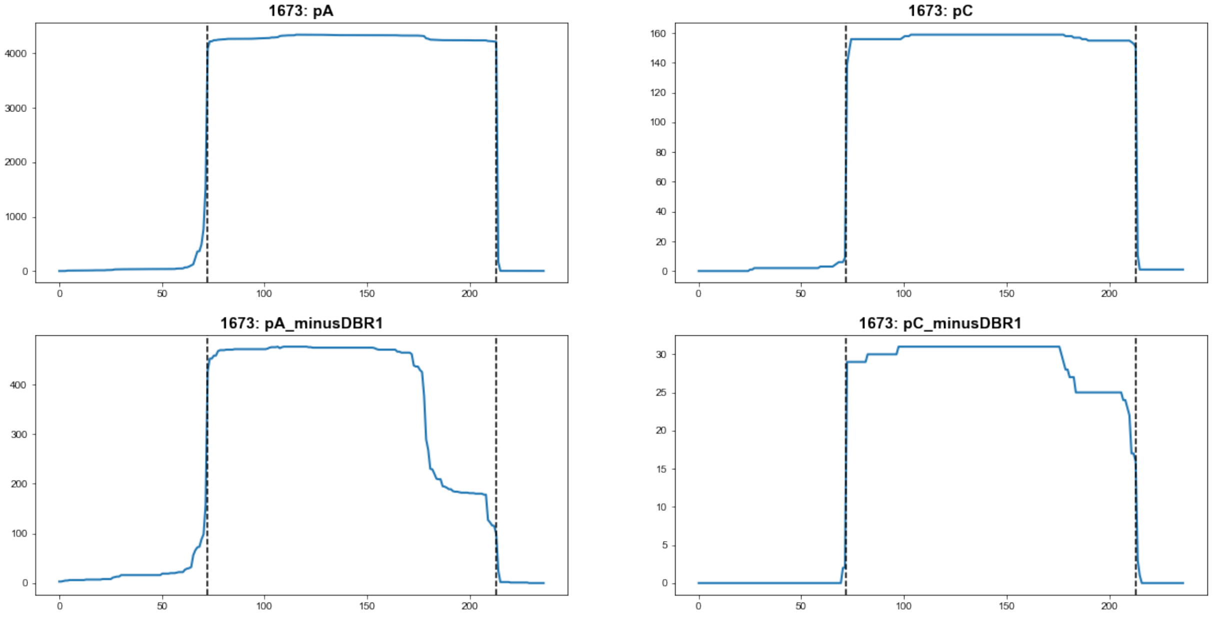


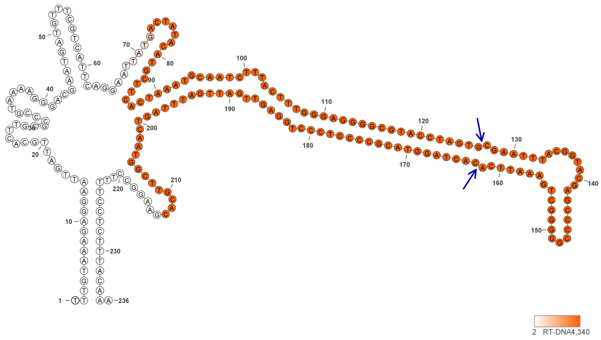

**1673**

### Retron Terminal 1687: Bacteroidetes

**Bacterial Vector for RTDNA: pSLS.689**

**Bacterial Vector for Editing: None**

**Phage Vector for Editing: None**

**Human Vector for Editing: pSLS.922**

**RTDNA Production (relative to Eco1): 0.347988202 by PAGE**

**Bacterial Editing: undetermined**

**Phage Editing: undetermined**

**Human Editing (demultiplexed): 12.26857 percent precise (105.6806299x Eco1)**

**ncRNA:**

**GGTAGTAATAAGATAACAACCTTGCAATTTGATTGGTGTATCGCCAATCTAGCTTACAGACACCCA ACGCGTCACTGACTCGAACTTAACTTGTGAAATAGGTACTATAATGCCATTTCCGGTTAAACTTCTT CGAAGTGAAATAGCACTATAGTACCTATTTCACTGGAGTTGTTCAATTCAGTCTTTAGAATTGTACG AAGGGAAAAAATATACATCTTATTGCTACC**

**RTDNA:**

**GTACAATTCTAAAGACTGAATTGAACAACTCCAGTGAAATAGGTACTATAGTGCTATTTCACTTCG AAGAAGTTTAACCGGAAATGGCATTATAGTACCTATTTCACAAGTTAAGTTCGAGTCAGTGACGCG TTGGGTGT**

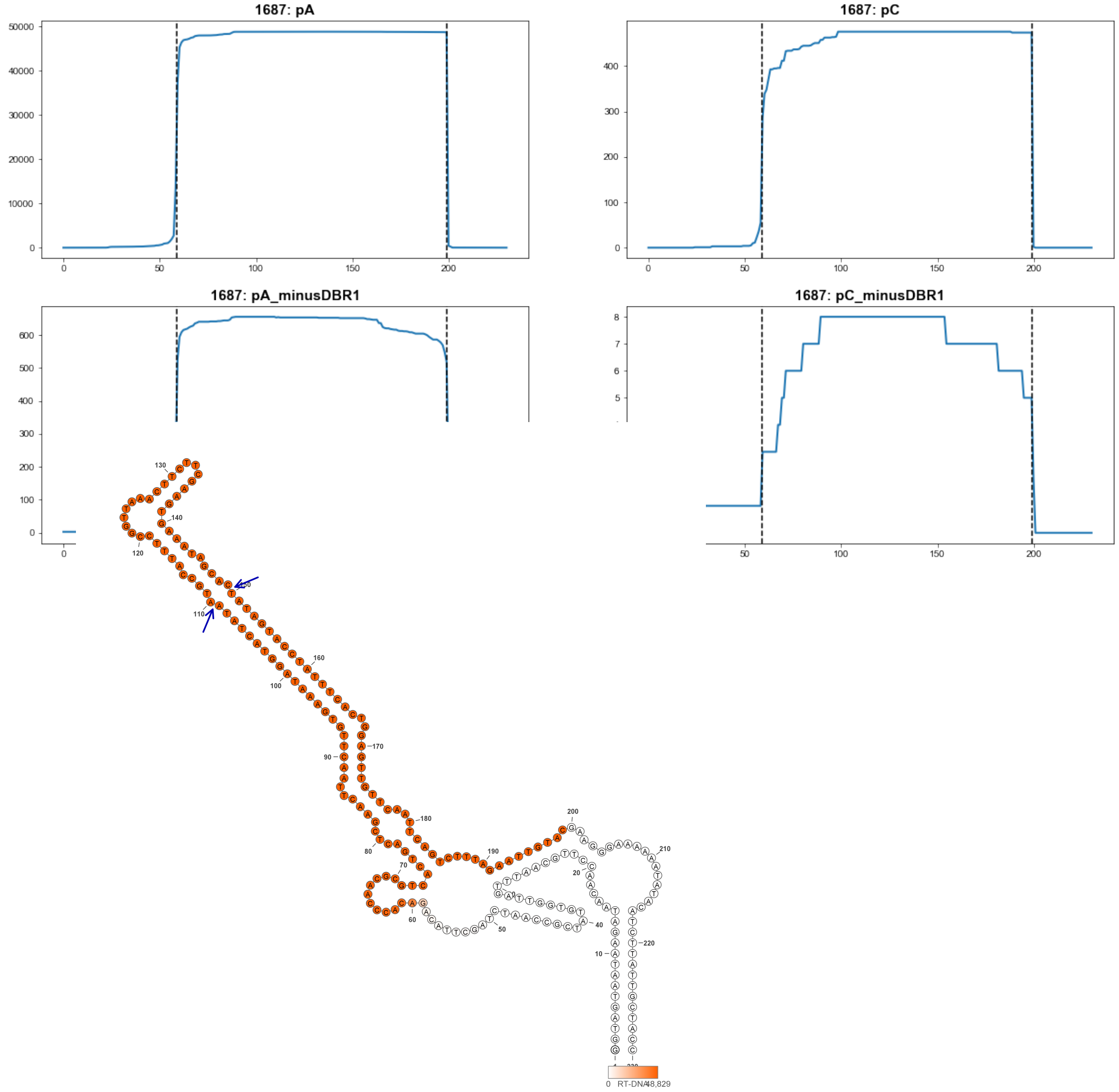


**1687**

### Retron Terminal 1703: Proteobacteria

**Bacterial Vector for RTDNA: pSLS.647**

**Bacterial Vector for Editing: pAGD300**

**Phage Vector for Editing: pAGD406**

**Human Vector for Editing: pSLS.887**

**RTDNA Production (relative to Eco1): 1.655638974 by PAGE**

**Bacterial Editing: 5.08805591 percent precise (1.565658574x Eco1)**

**Phage Editing: 26.3836458 percent precise (0.650743975x Eco1)**

**Human Editing (demultiplexed): 0.73231 percent precise (6.30806867x Eco1)**

**ncRNA:**

**TCGTGCAACGCGCAGGCTCTTGCAACCCGGACGGTGTTTCGCCGTCCATGACCTTTGCCGTTTCAT GCGCCACGGGGGACCGTTTTCACTGTGCCGAACATTGGCCTGCTGTGCACTTACGAAGAGGCTTG CGACAAGCCAATTGCACAGCAGGCCAAGTTCGGCGTCGAGTGAACAGCCCCCGTCCCGTGAGCC TTGCAGGCTCTTCGACGAGAAGCGCGTTGCACGA**

**RTDNA:**

**GAAGAGCCTGCAAGGCTCACGGGACGGGGGCTGTTCACTCGACGCCGAACTTGGCCTGCTGTGC AATTGGCTTGTCGCAAGCCTCTTCGTAAGTGCACAGCAGGCCAATGTTCGGCACAGTGAAAACGG TCCCCCGTGGCGCATGAAACG**

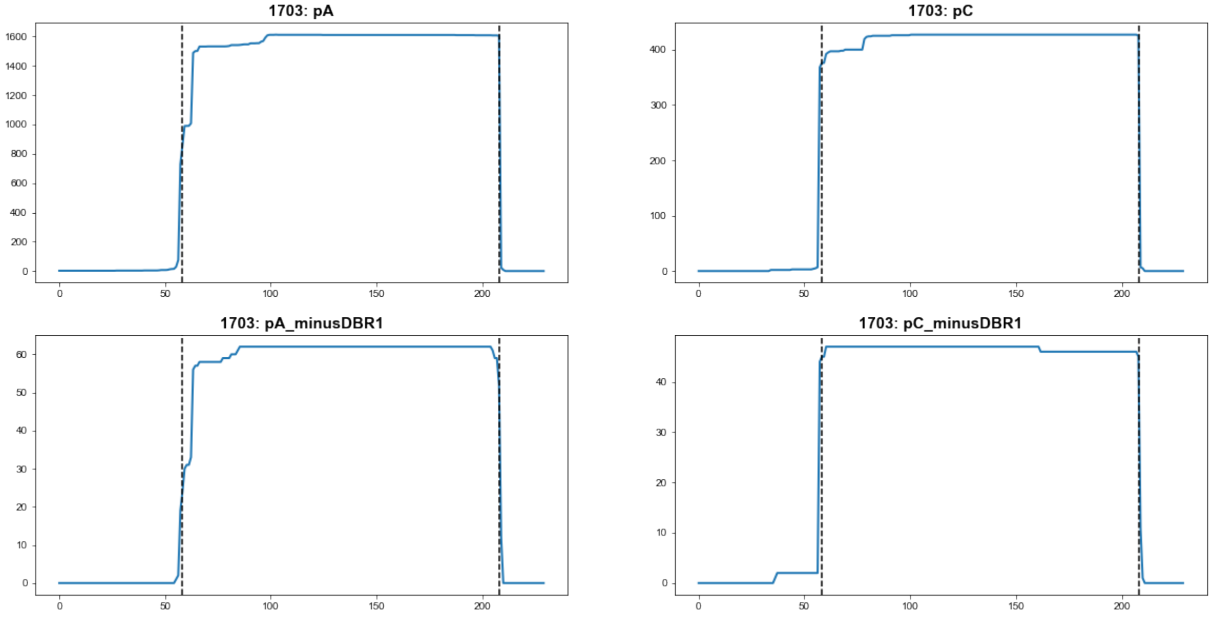


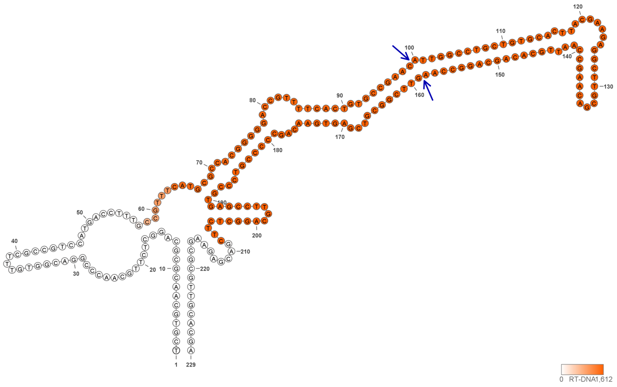

**1703**

### Retron Terminal 1710: Proteobacteria

**Bacterial Vector for RTDNA: pSLS.597**

**Bacterial Vector for Editing: pAGD294**

**Phage Vector for Editing: pAGD407**

**Human Vector for Editing: pSLS.832**

**RTDNA Production (relative to Eco1): 0.717899891 by PAGE**

**Bacterial Editing: 3.86430559 percent precise (1.189095263x Eco1)**

**Phage Editing: 4.932525205 percent precise (0.121659117x Eco1)**

**Human Editing (demultiplexed): 0 percent precise (0x Eco1)**

**ncRNA:**

**GGGCGTACGGGCAAGGCTCTTGCATCCTGGGTGGTGGATCGCCATCCGCGTTGTTTGCTCTGTCA TGCGTCACTGGCGCACTCTTCACTGTGCCGATGGCGCTACCCGCCGCTTTTGCCGTTTGACTTCCT TGAAGTCAAACAGCGGAGGGTAGCCCATCGGCGTCGAGTGGAGCGCGCAAGTCCGAAGGCTTCT TCGATGAGAACGCCCGTACGCCC**

**RTDNA:**

**GAAGAAGCCTTCGGACTTGCGCGCTCCACTCGACGCCGATGGGCTACCCTCCGCTGTTTGACTTC AAGGAAGTCAAACGGCAAAAGCGGCGGGTAGCGCCATCGGCACAGTGAAGAGTGCGCCAGTGA CGCATGACAGAG**

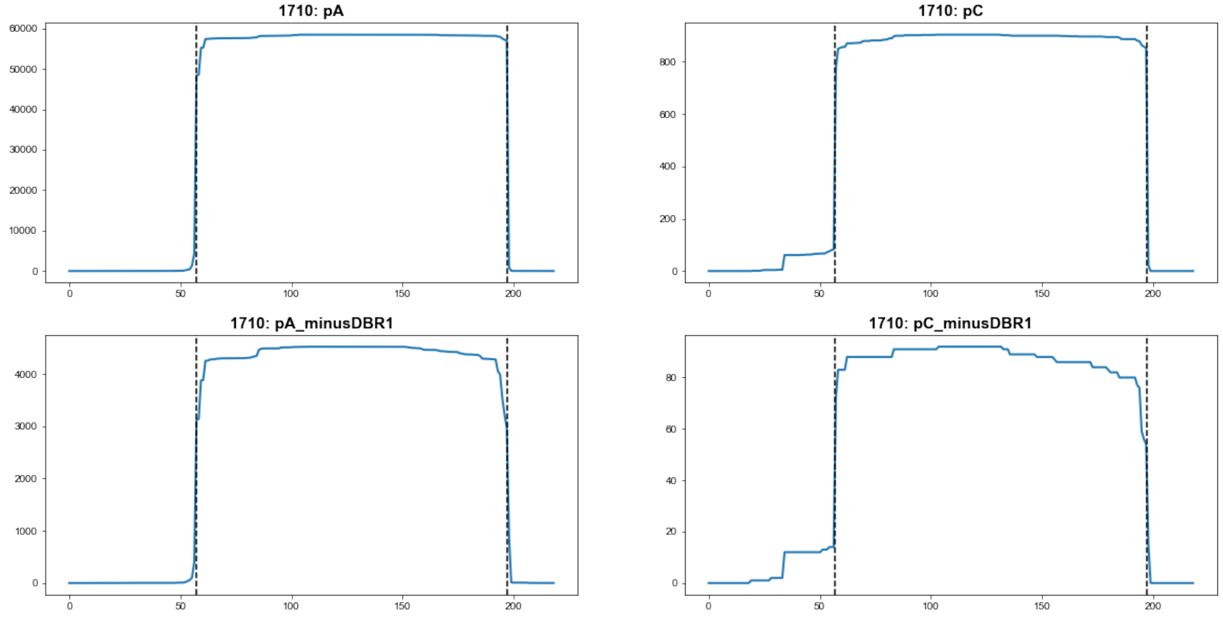


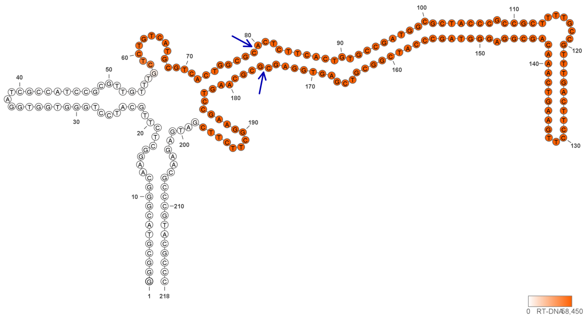


**1710**

### Retron Terminal 1718: Proteobacteria

**Bacterial Vector for RTDNA: None**

**Bacterial Vector for Editing: None**

**Phage Vector for Editing: None**

**Human Vector for Editing: pSLS.888**

**RTDNA Production (relative to Eco1): undetermined**

**Bacterial Editing: undetermined**

**Phage Editing: undetermined**

**Human Editing (demultiplexed): 0.140017 percent precise (1.206096941x Eco1)**

**ncRNA:**

**GCGGTAGCGCGACAAAACTACCTTGCAATCCGAGCGGTGGATCGCCGTTCTTGCTTTTAGACTTAT**
**CACGCTTCACTGGCCAGATCTTTACTGGGCCGTACTGCACCACCTATCGCTATTGCGGTTTGGTTT**
**CTACGAAACCAATGAGCGATAGGTGGTGCGTACGGCGTCGAGTTGATCAGTCCAGTTCTGAGAAT**
**TCTACGAAGGGAAACCATGTCGCGCTACCGC**

**no RTDNA sequencing data**

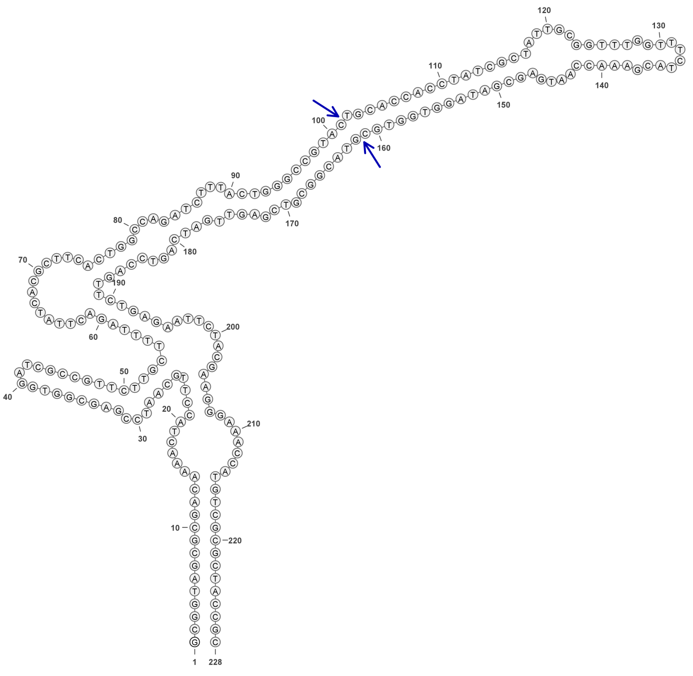

**1718**

### Retron Terminal 1731: Cyanobacteria

**Bacterial Vector for RTDNA: pSLS.690**

**Bacterial Vector for Editing: None**

**Phage Vector for Editing: None**

**Human Vector for Editing: pSLS.923**

**RTDNA Production (relative to Eco1): 0.03666868 by PAGE**

**Bacterial Editing: undetermined**

**Phage Editing: undetermined**

**Human Editing (demultiplexed): undetermined**

**ncRNA:**

**GGATGAAGCTAACGCGAATTATCAGCAAGTAACAACTTGCAACCCGGTCGGTGTATCGCCGTCCA TGCATTAAGCCCTAACATACTTCACTAGTCTGTTCTTTACTGTACGGTAGTAAGGTTCCTATCAGGG ATTTTCGGTGTGACTTCCATAAAGTAAATCCCTAATAGGAACCTTCTACCGCTTGAGTTGACCAGC CTAGTCCCCAGTAACTTACGAAGATAAACTTAGTGATAGTCGCGTTAGCTTTACTCT**

**RTDNA:**

**GTAAGTTACTGGGGACTAGGCTGGTCAACTCAAGCGGTAGAAGGTTCCTATTAGGGATTTACTTTA TGGAAGTCACACCGAAAATCCCTGATAGGAACCTTACTACCGTACAGTAAAGAACAGACTAGTGA AGTATGTTAGGGC**

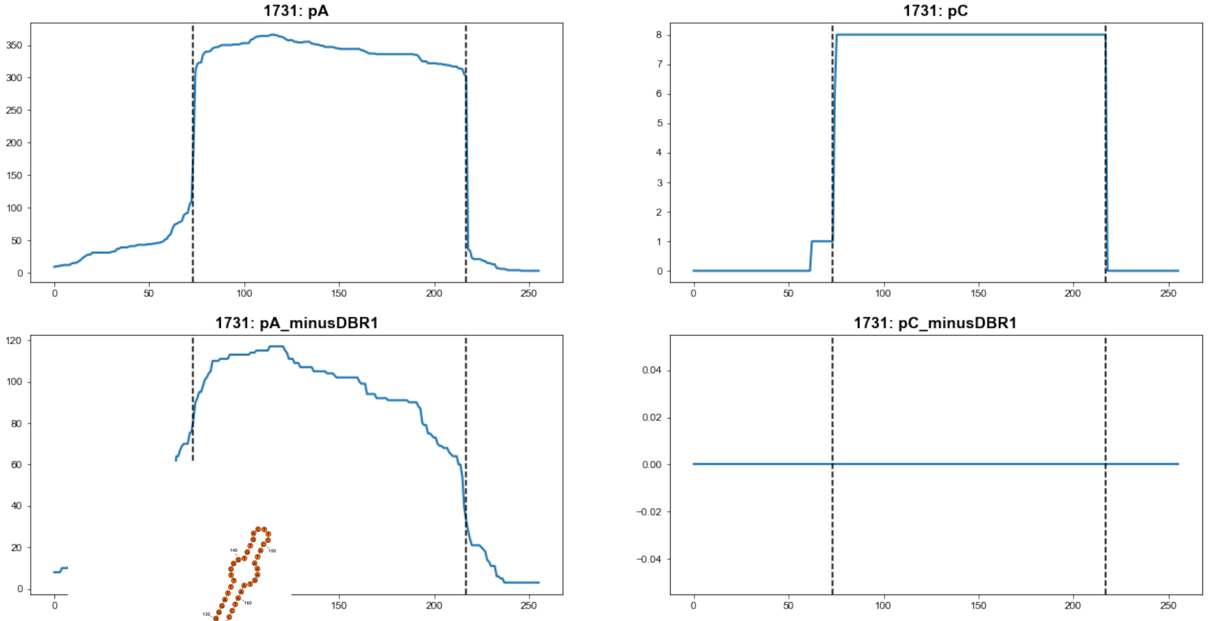


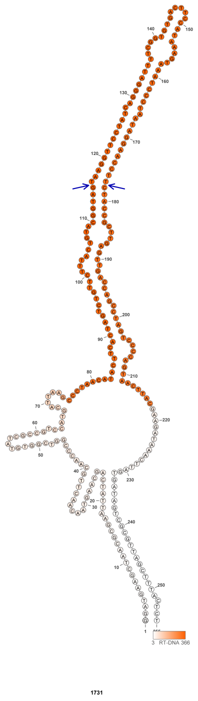

### Retron Terminal 1732: Cyanobacteria

**Bacterial Vector for RTDNA: pSLS.648**

**Bacterial Vector for Editing: pAGD297**

**Phage Vector for Editing: pAGD408**

**Human Vector for Editing: pSLS.889**

**RTDNA Production (relative to Eco1): 2.228198993 by PAGE**

**Bacterial Editing: 4.00776937 percent precise (1.233240866x Eco1)**

**Phage Editing: 42.08976525 percent precise (1.038130263x Eco1)**

**Human Editing (demultiplexed): 0.424507 percent precise (3.656674505x Eco1)**

**ncRNA:**

**TCACGCGCATCTGTAAGCTACGCGAAACAACCTTGCAAACCGGACGATGGTTCGTCGTCCTTGTG**
**CTTAGTCTAACCATGCATCACTAGCCTGCTCTCTACTTTGCGGAGCCACCCCCCGTACTACGGTTT**
**AGTTTCCGTGAAACTAATGACTATGGGGTGCTTGGTGGCTCCGCTGAGTTGAGCAGTCTAGTCCTG**
**AGACCTTTACGAAGGGGTAATTTGCACTTCGATGCGCGTGA**

**RTDNA:**

**GTAAAGGTCTCAGGACTAGACTGCTCAACTCAGCGGAGCCACCAAGCACCCCATAGTCATTAGTT**
**TCACGGAAACTAAACCGTAGTACGGGGGGTGGCTCCGCAAAGTAGAGAGCAGGCTAGTGATGCA**
**TGGTTAG**

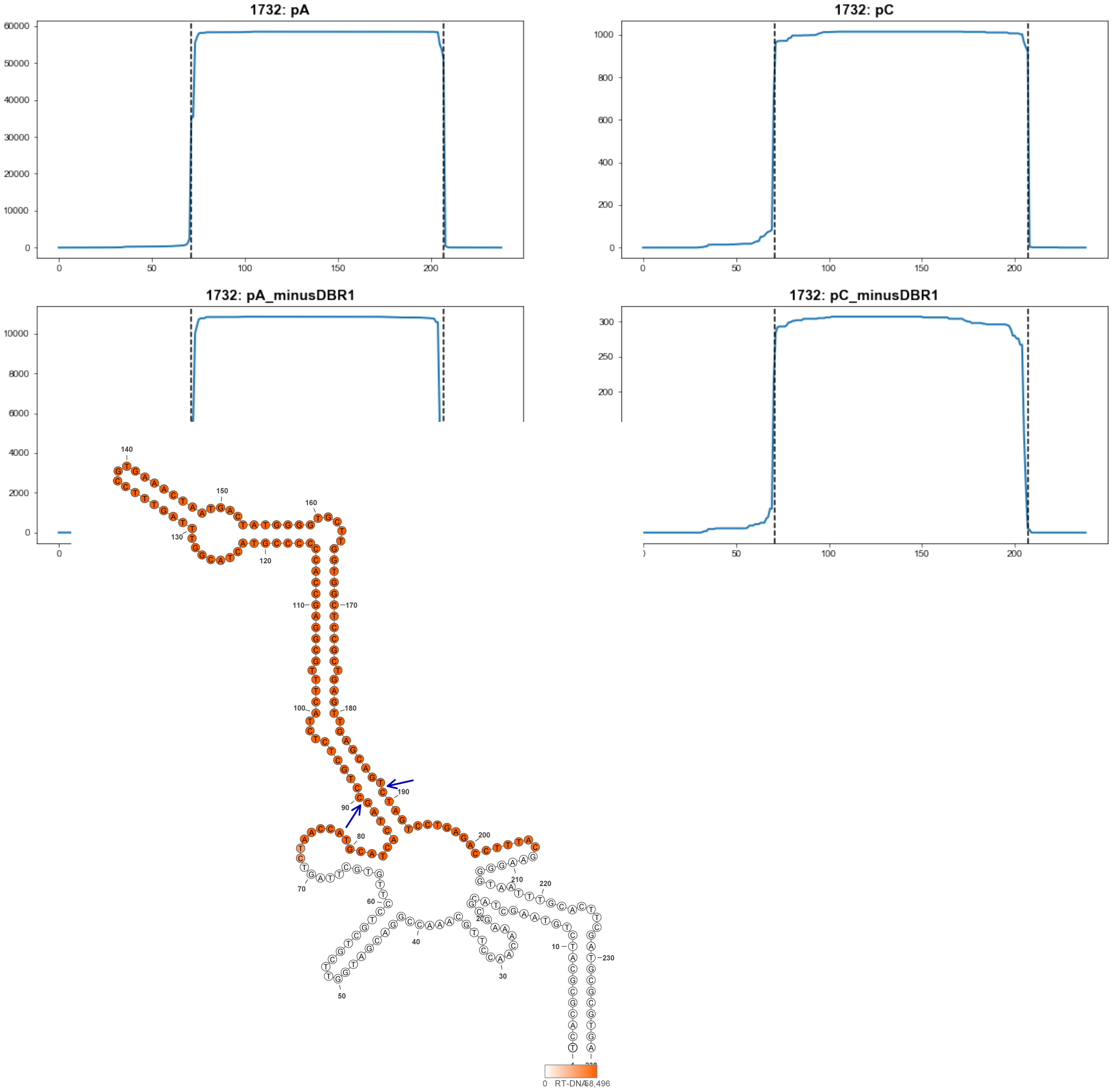


**1732**

### Retron Terminal 1739: Cyanobacteria

**Bacterial Vector for RTDNA: None**

**Bacterial Vector for Editing: None**

**Phage Vector for Editing: None**

**Human Vector for Editing: pSLS.1003**

**RTDNA Production (relative to Eco1): undetermined**

**Bacterial Editing: undetermined**

**Phage Editing: undetermined**

**Human Editing (demultiplexed): 0 percent precise (0x Eco1)**

**ncRNA:**

**CCTGTACCGGTGCCACCAAAGGGAATTTCCCCCATCAGCGACTTCATCAAGACAATGGTAAAAAA**
**TGTATAGCGGTTTCTAGCCTAATGAGGTACACAGGATTTTTATCTCTTGCCTCTTGCCTCTTGCCT**
**CTTTTTTGTTCCGGTGATCCGCTGTTCCCTATTCCCTGGGCATAGCGCTATAACAAACAACCTTTAA**
**CCAACCTTAAAAACTAAAAAATAGTTCCCATGTCAGACCAACCCCGTACCCGTCAGG**

**no RTDNA sequencing data**

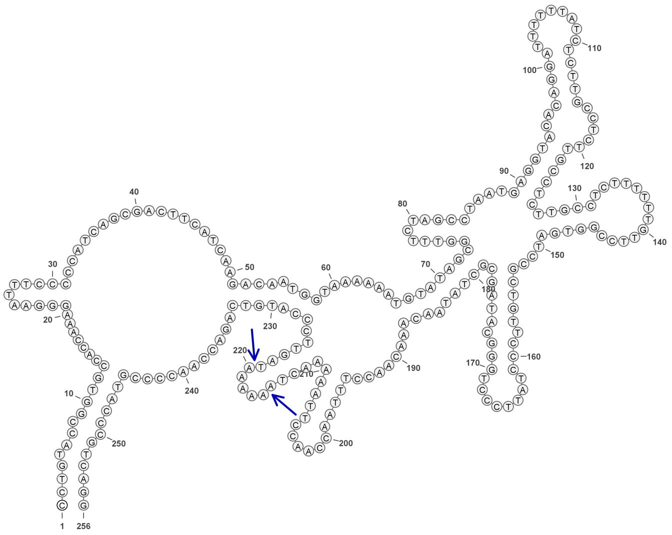

**1739**

### Retron Terminal 1746: Cyanobacteria

**Bacterial Vector for RTDNA: None**

**Bacterial Vector for Editing: None**

**Phage Vector for Editing: None**

**Human Vector for Editing: pSLS.1004**

**RTDNA Production (relative to Eco1): undetermined**

**Bacterial Editing: undetermined**

**Phage Editing: undetermined**

**Human Editing (demultiplexed): 0.106111 percent precise (0.914032957x Eco1)**

**ncRNA:**

**GCATCACATTGTAGGTAAAATCACCTTGCAATCTGGGTGGTGGATCGCCATCCATGTATAAAGTTA**

**AATCTCGATCTTTTATCGGTGAACTCGATCTTCACGAAGGTGTAACTTCATAAATGTGATGC**

**no RTDNA sequencing data**

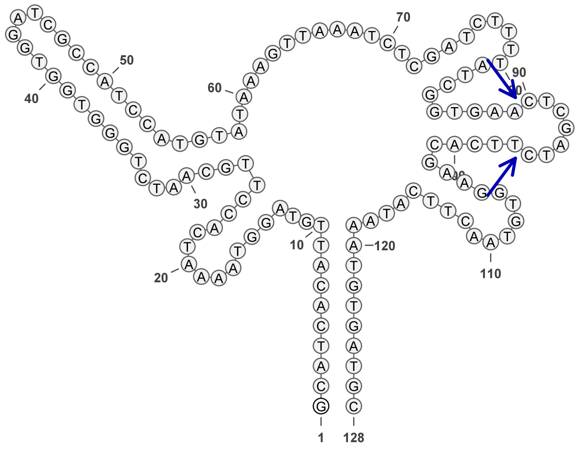


**1746**

### Retron Terminal 1765: Proteobacteria

**Bacterial Vector for RTDNA: pSLS.691**

**Bacterial Vector for Editing: None**

**Phage Vector for Editing: None**

**Human Vector for Editing: pSLS.924**

**RTDNA Production (relative to Eco1): undetermined**

**Bacterial Editing: undetermined**

**Phage Editing: undetermined**

**Human Editing (demultiplexed): 5.356156 percent precise (46.1375645x Eco1)**

**ncRNA:**

**CAGAGGGGCGCCGGCCGGCGAGGACCTTGCAACCCGGGCGGCGTTTCGCCGTCCTTGCACCCTG**
**CCCGGCAAAGCGTCACTGGCCCGCTCTTCACTGGGGCGGCACAGGCCACCGGTGCGGGTTTACG**
**GTTTGGGTTCCCTGAACCCAAACCCGCACCGGTGGCCGTGCCGCCTTCGAGTGGAGCGGGCCAG**
**TCCCAGGACGAGACGAAGGCGAGCCGTCCGGCGTCCCTCTG**

**no RTDNA sequencing data**

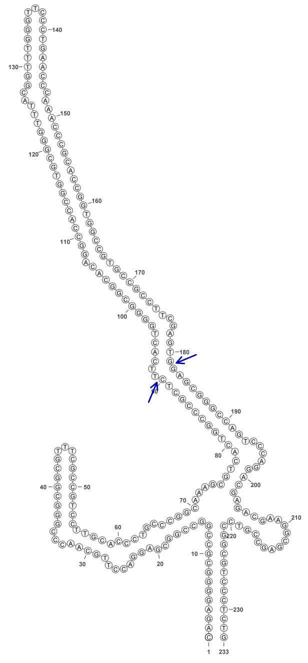

1765

### Retron Terminal 1795: Proteobacteria

**Bacterial Vector for RTDNA: None**

**Bacterial Vector for Editing: None**

**Phage Vector for Editing: None**

**Human Vector for Editing: pSLS.810 (H1)**

**RTDNA Production (relative to Eco1): undetermined**

**Bacterial Editing: undetermined**

**Phage Editing: undetermined**

**Human Editing (demultiplexed): 3.530519 percent precise (30.4116512x Eco1)**

**ncRNA:**

**GTGCGAGCGACCGAGAGAGGTCCCAAGCCATCAGCCTCAGCGCCTCGAGCGCGAGAGCGGCGT TGCGCCGCTCTGGTTGAATTGCAGGACACTCTCCGCAAGGTAGCCTGTTCTTGGCTCTCTTCCCTC CGGTGAGTACCTCTCCGGCCGGGGAGCTGAACCAACGACGCAACCGCCGTTTCCCCGGCCGGAG AGGTACTCACCGGAGGGGAGAGCCGGTGAGGCTACCGTGCCCCAGGTGAGAAGGTGGTGCCTTC GGGCCTCCCTCGACCGCTCGCGC**

**no RTDNA sequencing data**

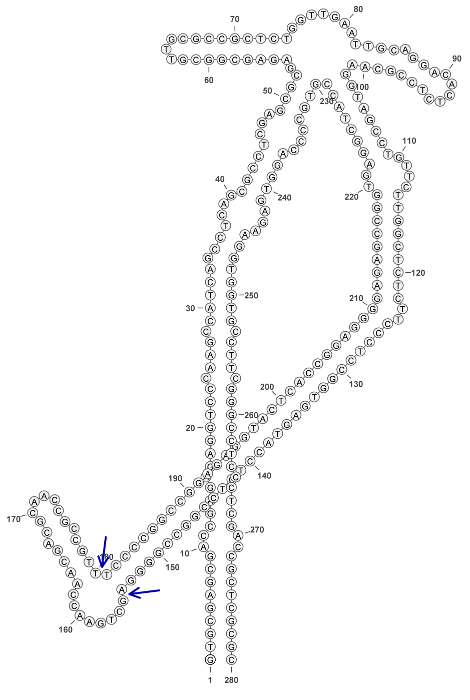

**1795**

### Retron Terminal 1807: Actinobacteria

**Bacterial Vector for RTDNA: pSLS.692**

**Bacterial Vector for Editing: None**

**Phage Vector for Editing: None**

**Human Vector for Editing: pSLS.925**

**RTDNA Production (relative to Eco1): undetermined**

**Bacterial Editing: undetermined**

**Phage Editing: undetermined**

**Human Editing (demultiplexed): 0.052495 percent precise (0.452188369x Eco1)**

**ncRNA:**

**CACGCGCACCGACGCTACTCCCACCGCCACGCGGGGTGACACGTCGTCGTCGCGGGCGGCTCTC**
**GAGGAGCGGCGCGCGGCCGTCGCGGCGACCGACCGACGGACTGGAGGCGTGGTGCGGGCCCG**
**CCCCGTGCCTCCCGTCCGTCAGGGGGTCGCCGTAGGGTGAGCGCGTG**

**no RTDNA sequencing data**

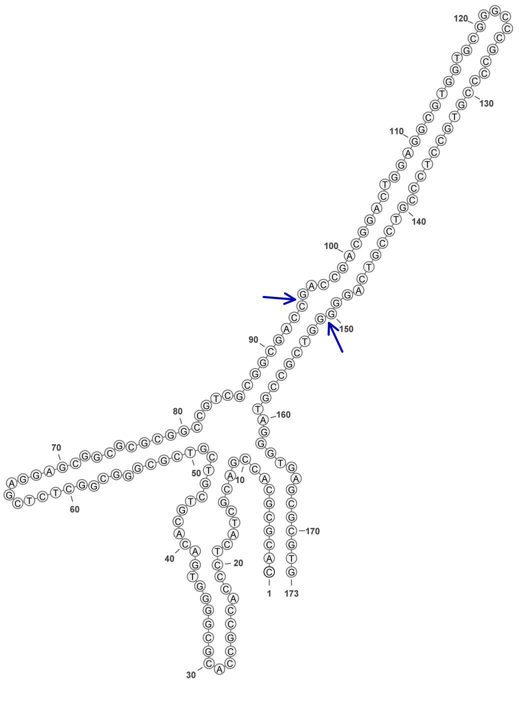


**1807**

### Retron Terminal 1823: Actinobacteria

**Bacterial Vector for RTDNA: None**

**Bacterial Vector for Editing: None**

**Phage Vector for Editing: None**

**Human Vector for Editing: pSLS.942**

**RTDNA Production (relative to Eco1): undetermined**

**Bacterial Editing: undetermined**

**Phage Editing: undetermined**

**Human Editing (demultiplexed): 0.509106 percent precise (4.385404553x Eco1)**

**ncRNA:**

**TTCCGGATGAGGTGGGTGGGCATTTGCGTGCCCTGCCCTCTCCGGGGATACCCCGGGGGACGAA**

**GAACGACCTCCTGCAAGCACCGCAGCCGCCTCGCAGCGCTCCGCGCAGCAGGGCGGCCGCACA**

**GACTGACGGGTCATCCGGAA**

**no RTDNA sequencing data**

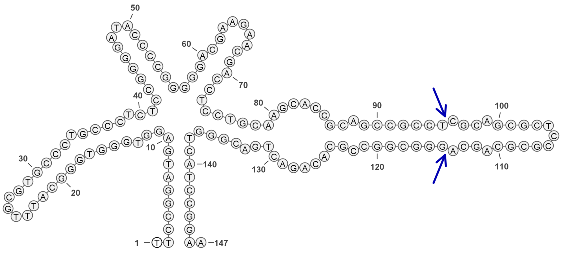


**1823**

### Retron Terminal 1840: Actinobacteria

**Bacterial Vector for RTDNA: pSLS.693**

**Bacterial Vector for Editing: None**

**Phage Vector for Editing: None**

**Human Vector for Editing: None**

**RTDNA Production (relative to Eco1): 0 by PAGE**

**Bacterial Editing: undetermined**

**Phage Editing: undetermined**

**Human Editing (demultiplexed): undetermined**

**ncRNA:**

**GGCGATGGTGGGCGAGAACTCCGGCCGGTGACGGTTCGGACGCTCCGGGGCACCCCCGGGGTG GGTTCACCGCAGTTGGCACCACCGAGCAGGCACATCGACGCGCTCCGCGCATGCAGGTGCCCGC CCAGCAGGGACAGGTCGCCCACCGTCGCC**

**RTDNA:**

**GGTGGGCGACCTGTCCCTGCTGGGCGGGCACCTGCATGCGCGGAGCGCGTCGATGTGCCTGCTC GGTGGTGCCAACTGCGGTGAACCCACCCCGG**

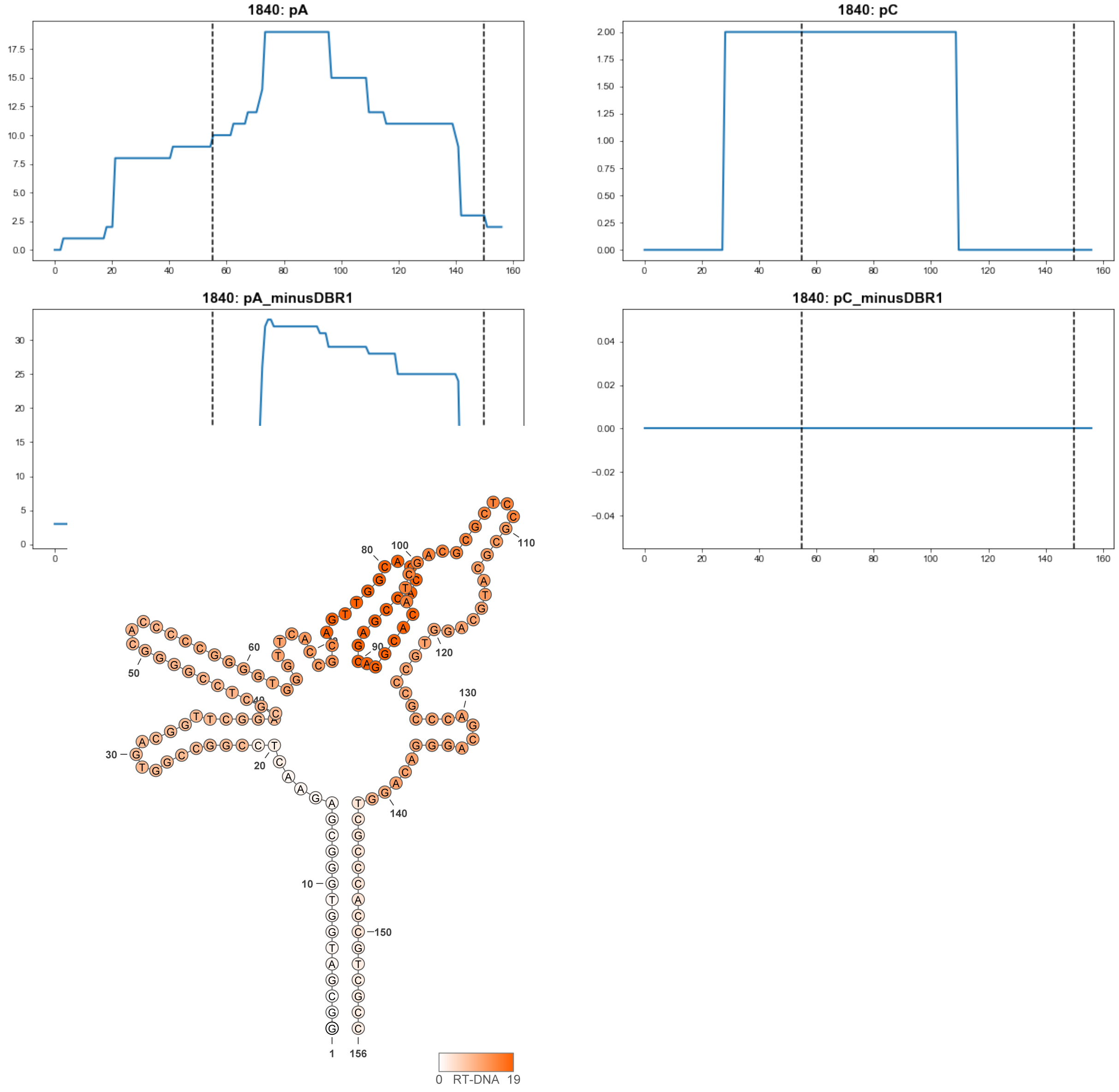


**1840**

### Retron Terminal 1841: Actinobacteria

**Bacterial Vector for RTDNA: None**

**Bacterial Vector for Editing: None**

**Phage Vector for Editing: None**

**Human Vector for Editing: pSLS.850**

**RTDNA Production (relative to Eco1): undetermined**

**Bacterial Editing: undetermined**

**Phage Editing: undetermined**

**Human Editing (demultiplexed): 0.012393 percent precise (0.106752461x Eco1)**

**ncRNA:**

**GTCGACGACGGCAGGCGAGACTGCTGGTTCGCCTCCGAGTGGTCAGCGGGGCCGATCAGCTGCT**
**CCGGGGATACCCCGGGGCCGGTTGTGGTGGTATCGCAGCACGTCACAGGCATGATGCCGCGCTC**
**CGCGCAGGGCAATGCCCGTCAGGCAGGACAGGTCGCCTGCCGTCGTCGAC**

**no RTDNA sequencing data**

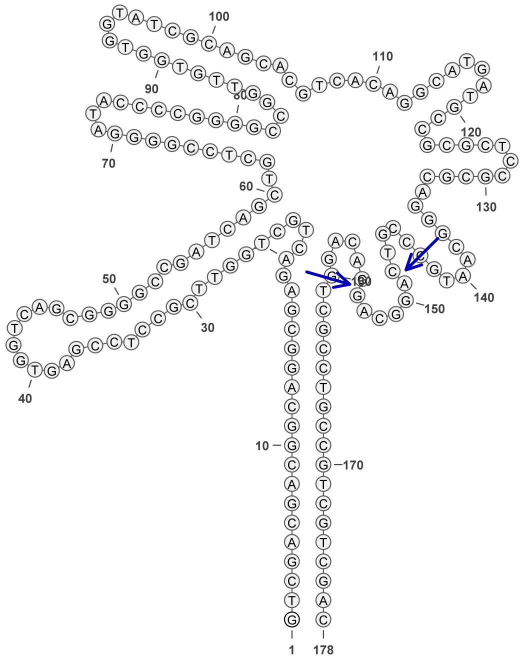

1841

### Retron Terminal 1856: Actinobacteria

**Bacterial Vector for RTDNA: pSLS.649**

**Bacterial Vector for Editing: None**

**Phage Vector for Editing: None**

**Human Vector for Editing: pSLS.890**

**RTDNA Production (relative to Eco1): 0 by PAGE**

**Bacterial Editing: undetermined**

**Phage Editing: undetermined**

**Human Editing (demultiplexed): 0 percent precise (0x Eco1)**

**ncRNA:**

**GGTGCGCGGCGGGGCGGGCAGGACGACCGGTGAGGACCGGTTCTCCCGGGGCAACCCCGGGG**

**ATCGCACTGTTCCCGAGCATCTCGTCAACCGCCAGGCCGTGCGCTCCGCGCACCTGATGGCGCCT**

**CGATCAGGGGCTGGCTGCCCGCCCCGCCGCGCACT**

**RTDNA: None**

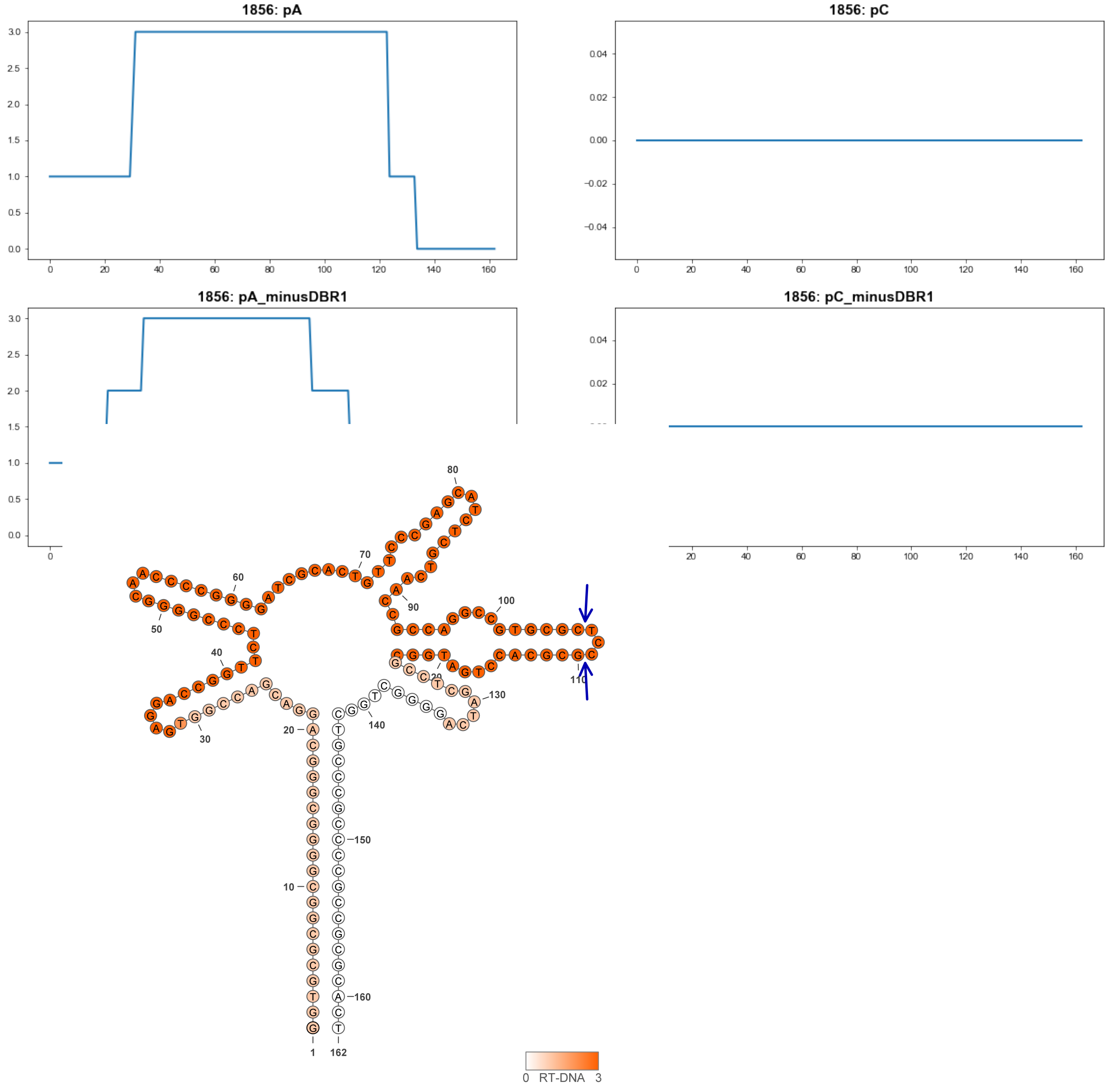


**1856**

### Retron Terminal 1873: Actinobacteria

**Bacterial Vector for RTDNA: None**

**Bacterial Vector for Editing: None**

**Phage Vector for Editing: None**

**Human Vector for Editing: pSLS.971**

**RTDNA Production (relative to Eco1): undetermined**

**Bacterial Editing: undetermined**

**Phage Editing: undetermined**

**Human Editing (demultiplexed): 0 percent precise (0x Eco1)**

**ncRNA:**

**ATGGCAGCGTGCCCCGAGCTGACCTGGCCGCGCCCCCACCGGGGACGCGGTCGGAGTCTCTCCG**
**GGGCAACCCCGGGGGCGGCACATCCACCTGGGCACCACACCACAGGTGCGTCGGCACGCTCCG**
**CGAGCCTGTGCACCCGTACGACCGGGATTCGTCGGGGCACGCTGTCAT**

**no RTDNA sequencing data**

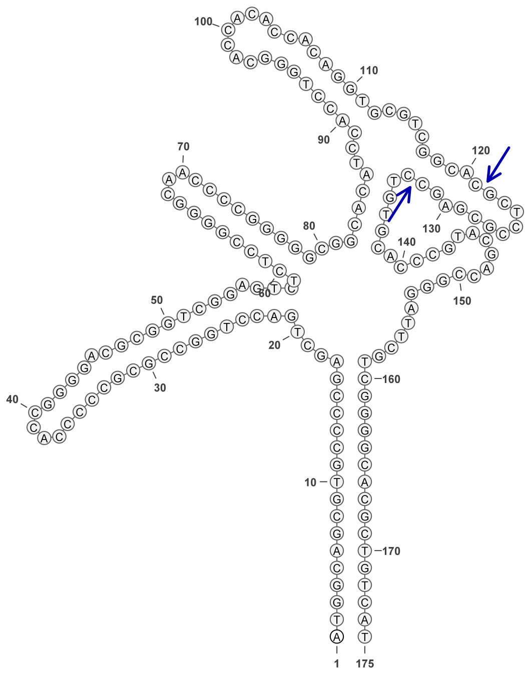

**1873**

### Retron Terminal 1875: Proteobacteria

**Bacterial Vector for RTDNA: pSLS.585**

**Bacterial Vector for Editing: pAGD287**

**Phage Vector for Editing: None**

**Human Vector for Editing: pSLS.818 (H1)**

**RTDNA Production (relative to Eco1): 0.02203482 by PAGE**

**Bacterial Editing: 0.47069324 percent precise (0.144838209x Eco1)**

**Phage Editing: undetermined**

**Human Editing (demultiplexed): 1.271094 percent precise (10.9491175x Eco1)**

**ncRNA:**

**GCCATGCGGCGGCTTGGACACCCAGCGGGTAGTCGGTCCGTGAAGCTGCGGTCCACTGCCTCTT**
**CGGGGTACCCCCGGGGACAACTCATCAGGAGCACACCGTCACGCGTGGCCGTCTCCGCGGCAAG**
**CCGCAGTTCGGGCCACGCAAGACACGACAGGTGGGGCCAGTCCGCCGCATGGC**

**RTDNA: None**

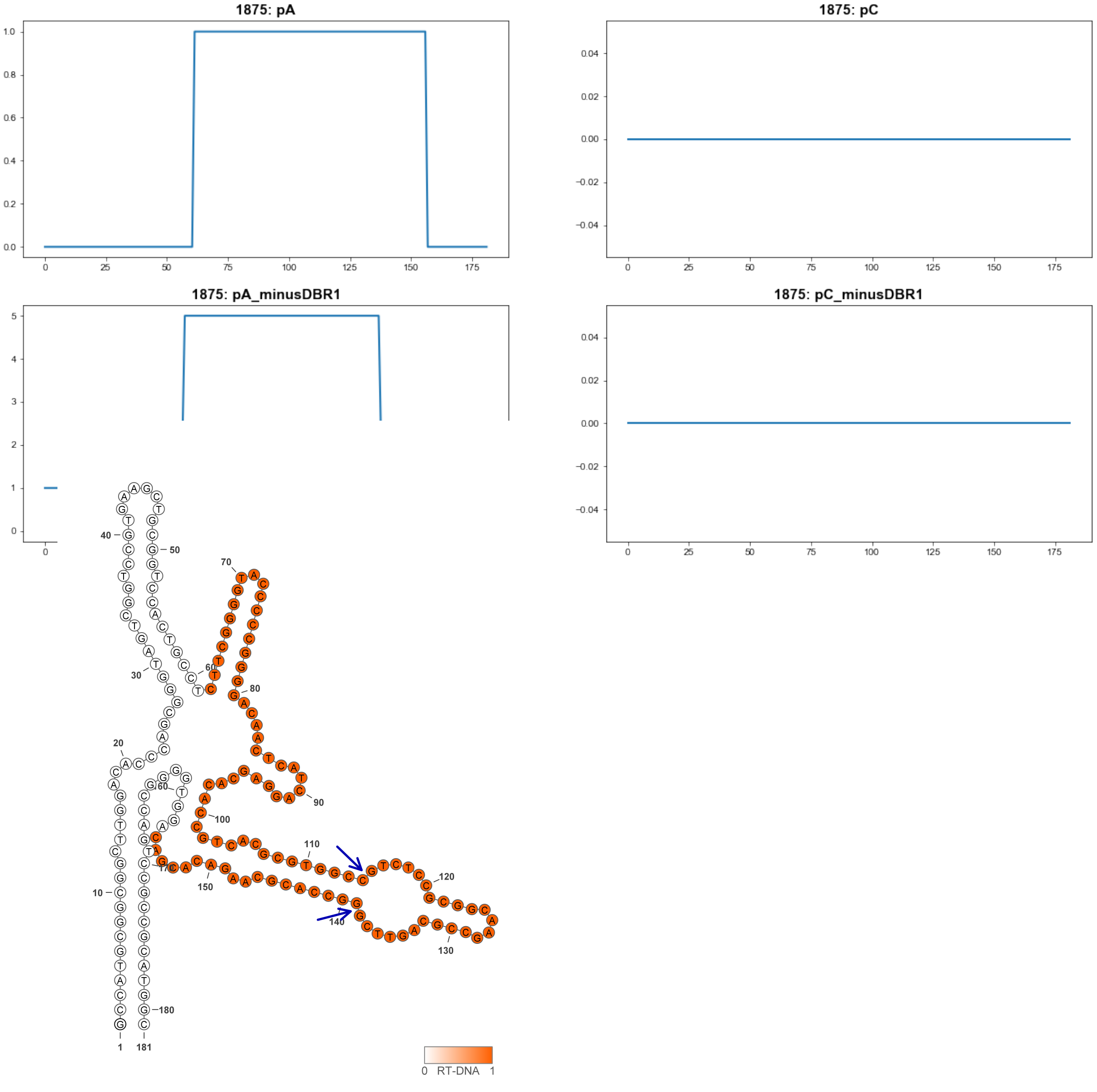


1875

### Retron Terminal 1878: Actinobacteria

**Bacterial Vector for RTDNA: None**

**Bacterial Vector for Editing: None**

**Phage Vector for Editing: None**

**Human Vector for Editing: pSLS.892**

**RTDNA Production (relative to Eco1): undetermined**

**Bacterial Editing: undetermined**

**Phage Editing: undetermined**

**Human Editing (demultiplexed): 0.001175 percent precise (0.01012137x Eco1)**

**ncRNA:**

**CGGCGACGCGGAGGTGAGCCGGTGGAGGACCACCCACTCCGGGGCAACCCCGGGGAGCAGCAC**

**GACACGTACCCACAGCGCCCCGCGCAGCGGCGGCACGCTCCGCGTACAAGACAGCGCGGATGA**

**ACGGAGAACGAGTCGCCTCCGCGTCGCTG**

**no RTDNA sequencing data**

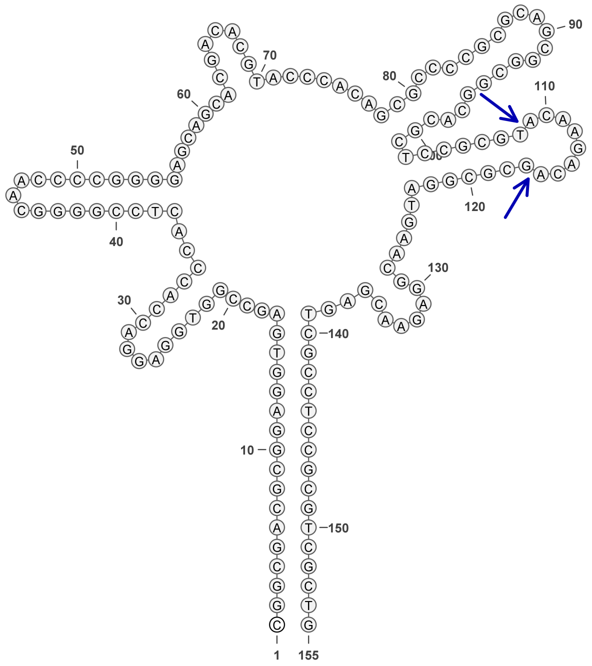

**1878**

### Retron Terminal 1884: Actinobacteria

**Bacterial Vector for RTDNA: None**

**Bacterial Vector for Editing: None**

**Phage Vector for Editing: None**

**Human Vector for Editing: pSLS.972**

**RTDNA Production (relative to Eco1): undetermined**

**Bacterial Editing: undetermined**

**Phage Editing: undetermined**

**Human Editing (demultiplexed): 0.421671 percent precise (3.632245394x Eco1)**

**ncRNA:**

**GCGGCGACACGGCAGTGAGCACGGTGGTGAACACCACCCCTCTGGGGCTACCCCAGGGAGCAC**

**GGATTTACCTGCGGATGGCGCCCCACGCGCGACAGCACACGCTGTGTACGCATGGGCGCGCGGA**

**TATCGGGAGAACAGGTCACTTCGTGTCGCCGC**

**no RTDNA sequencing data**

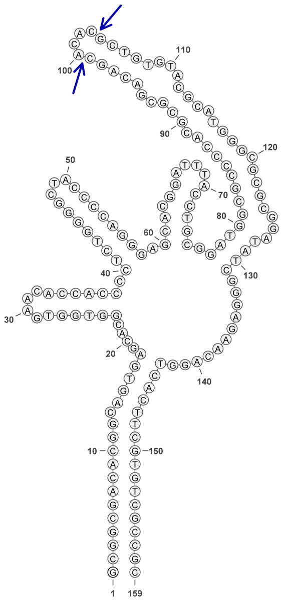

1884

### Retron Terminal 1894: Actinobacteria

**Bacterial Vector for RTDNA: pSLS.694**

**Bacterial Vector for Editing: None**

**Phage Vector for Editing: None**

**Human Vector for Editing: pSLS.894**

**RTDNA Production (relative to Eco1): 0.164705512 by PAGE**

**Bacterial Editing: undetermined**

**Phage Editing: undetermined**

**Human Editing (demultiplexed): 0.194151 percent precise (1.672403545x Eco1)**

**ncRNA:**

**TGGGCGGTCGGGGTGAGAGTGGATCGAGAGATCCGTCCCAGGGGCAACCCCTGGGTACATGTGA CAGTTGGTTCCTGCGTCCGACGCCGGTCCGCACGCTGCGCGTGCCGGGCGGCGACGAGCACTCG GCTCGTGGTCACCTCGACCGTCCA**

**RTDNA:**

**CCACGAGCCGAGTGCTCGTCGCCGCCCGGCACGCGCAGCGTGCGGACCGGCGTCGGACGCAGG AACCA**

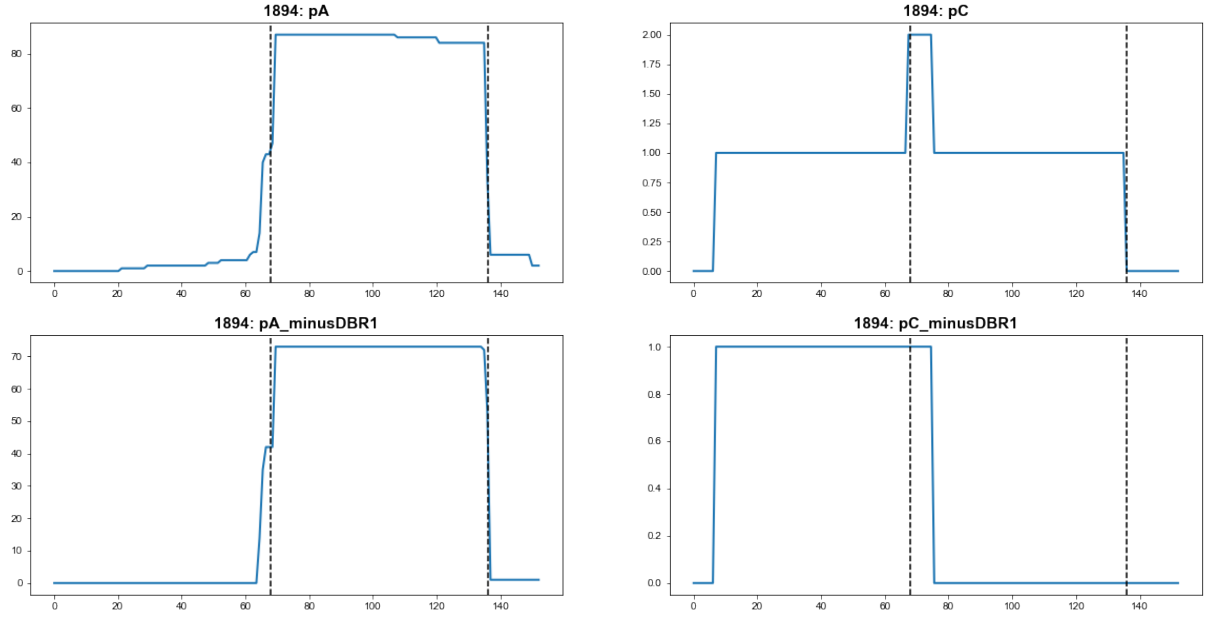


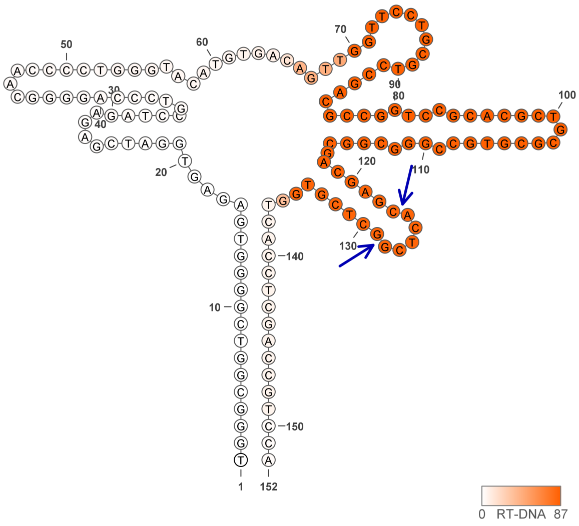


**1894**

### Retron Terminal 1895: Actinobacteria

**Bacterial Vector for RTDNA: None**

**Bacterial Vector for Editing: None**

**Phage Vector for Editing: None**

**Human Vector for Editing: pSLS.973**

**RTDNA Production (relative to Eco1): undetermined**

**Bacterial Editing: undetermined**

**Phage Editing: undetermined**

**Human Editing (demultiplexed): 0.868897 percent precise (7.484619824x Eco1)**

**ncRNA:**

**GGAGACCGGGTGCTGTGAGAGTGGATCGAGAGATCCGTCTCAGGGGTAACCCCTGGGTGCAGTT CACAGCAAGACCCCGCGGCCGGAGCCACGCCGCACGCTGTGCGTGCCGGATGGCACCGCGAAC ACCGGTGTGAGGTCACAGCACCCGGACTCC**

**no RTDNA sequencing data**

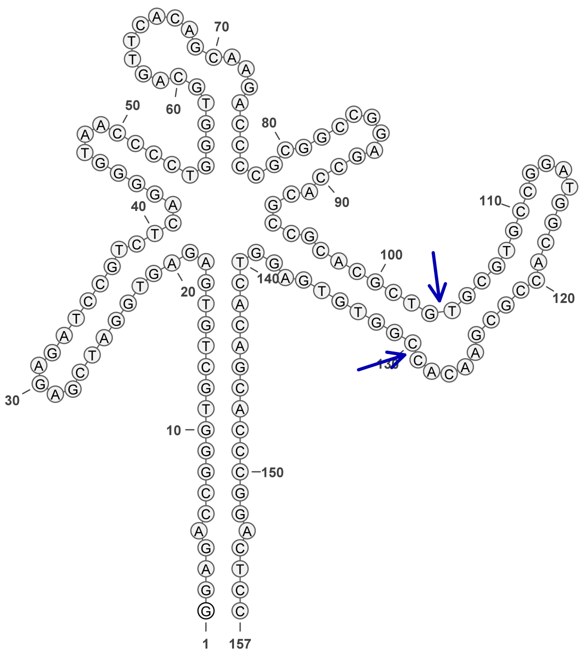

1895

### Retron Terminal 1898: Actinobacteria

**Bacterial Vector for RTDNA: None**

**Bacterial Vector for Editing: None**

**Phage Vector for Editing: None**

**Human Vector for Editing: pSLS.895**

**RTDNA Production (relative to Eco1): undetermined**

**Bacterial Editing: undetermined**

**Phage Editing: undetermined**

**Human Editing (demultiplexed): 0 percent precise (0x Eco1)**

**ncRNA:**

**GATGGGCGACGGGTGTGGTGAGAGGGGATCGGTTCGATCCGTCTCAGGGGTAACCCCTGGGTAC**
**AGTTCACAGTTGGGCCCTGCGTCCGACGCCGTGCAGCACGCTGCACGTGCAGGCCGGCGCCGCA**
**TCAAGGTTCTGGTCACCACACCCGTCGGCCATC**

**no RTDNA sequencing data**

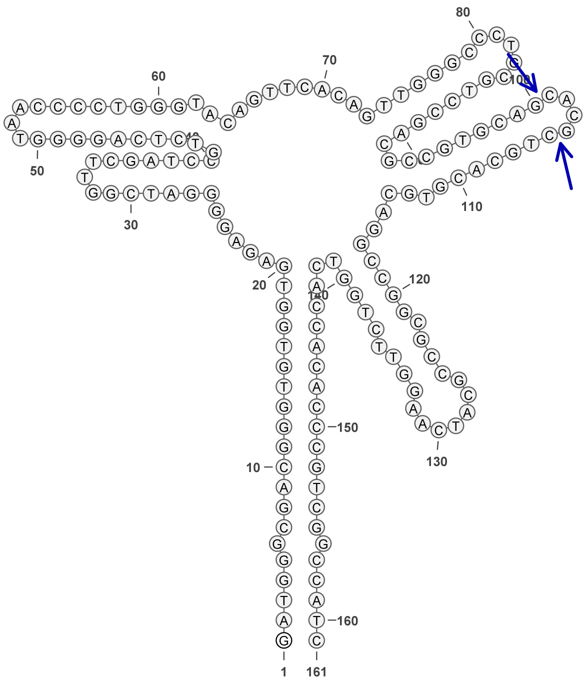

**1898**

### Retron Terminal 1912: Actinobacteria

**Bacterial Vector for RTDNA: pSLS.651**

**Bacterial Vector for Editing: None**

**Phage Vector for Editing: None**

**Human Vector for Editing: pSLS.896**

**RTDNA Production (relative to Eco1): 0 by PAGE**

**Bacterial Editing: undetermined**

**Phage Editing: undetermined**

**Human Editing (demultiplexed): 0 percent precise (0x Eco1)**

**ncRNA:**

**ATGCGGGGTGGGCGAGGCGAGAGTGGGTCGATGCGACCCGTCTCAGGGGTAACCCCTGGGTGC AGTTCACAGTTGGTATCAGCGTCCGACGCCGGACCACATGCTTCGCGTGTCCGGCGGCGCCGCG GACAGGTGCGGTTGGTCGCCCCGCCCGCCCCGCAT**

**RTDNA: None**

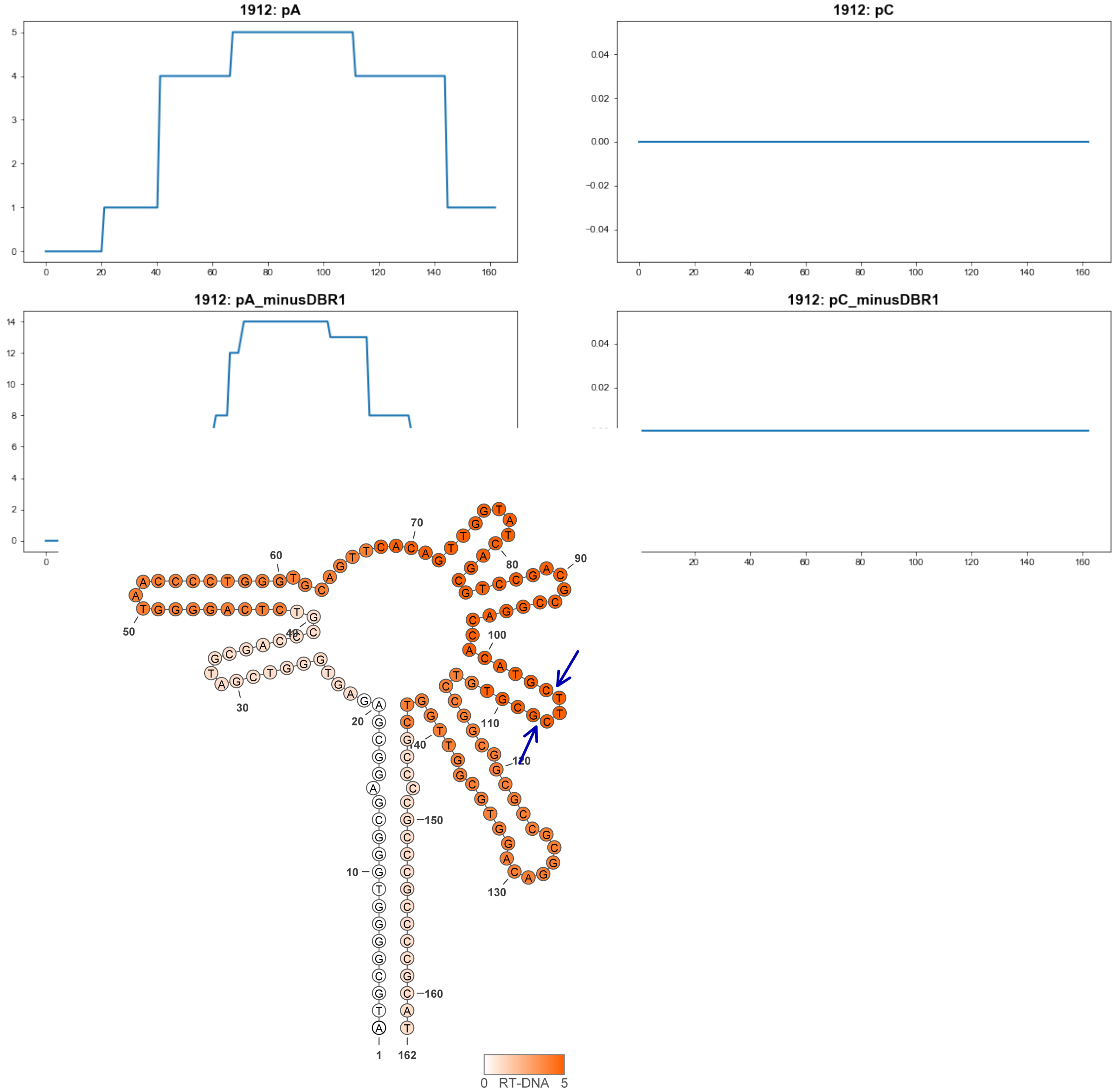


**1912**

### Retron Terminal 1913: Actinobacteria

**Bacterial Vector for RTDNA: pSLS.586**

**Bacterial Vector for Editing: pAGD288**

**Phage Vector for Editing: None**

**Human Vector for Editing: pSLS.819 (H1)**

**RTDNA Production (relative to Eco1): 0.020897778 by PAGE**

**Bacterial Editing: 0.40074262 percent precise (0.123313527x Eco1)**

**Phage Editing: undetermined**

**Human Editing (demultiplexed): 0.859534 percent precise (7.403967577x Eco1)**

**ncRNA:**

**GCGACGGTGGCGGGGCGAGAGCGGGTCGGGTCGACCCACCTCAGGGGTAACCCCTGGGAGCGT CTTGTCAGTTGGGCTCAGCGTCCGGAGCAAGGGCGCACGCTTCGCGTGCCTCGCAGCACCGCGG ACGAAGGATTCTGGTCGCCCCGCCACCGTCGC**

**RTDNA:**

**CCAGAATCCTTCGTCCGCGGTGCTGCGAGGCACGCGAAGCGTGCGCCCTTGCTCCGGACGCTGA GCCCAACT**

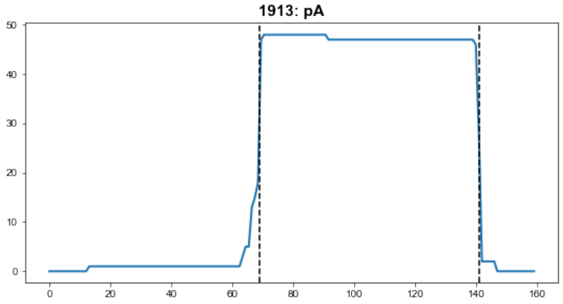


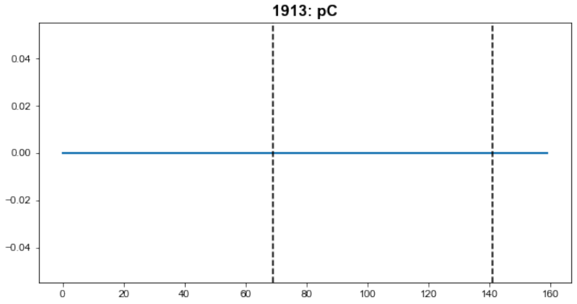


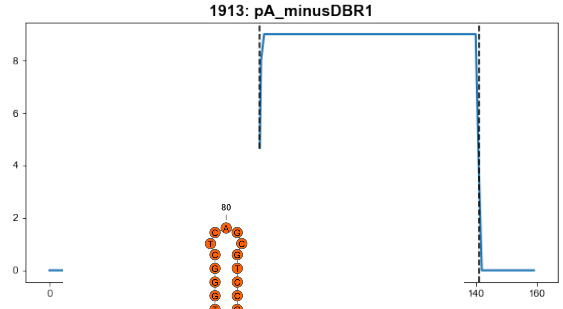


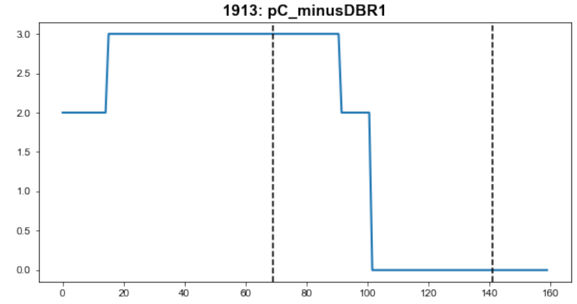


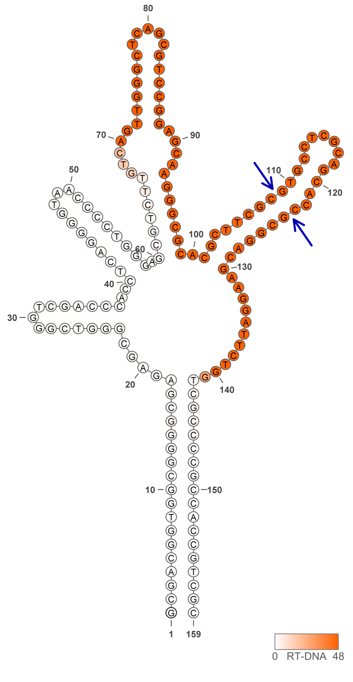


1913

### Retron Terminal 1914: Actinobacteria

**Bacterial Vector for RTDNA: pSLS.652**

**Bacterial Vector for Editing: None**

**Phage Vector for Editing: None**

**Human Vector for Editing: pSLS.897**

**RTDNA Production (relative to Eco1): 0 by PAGE**

**Bacterial Editing: undetermined**

**Phage Editing: undetermined**

**Human Editing (demultiplexed): undetermined**

**ncRNA:**

**GCGACGGTGGCGGGGTGAGAGCGGGTCGGTAGGACCCACCTCAGGGGCAACCCCTGGGAACGA**
**CACATCGATCGGGTACAGCGTCCGGAGCCGGCGCGCACGCACCGAGTGCCTCGCGGCACCGCG**
**GACGAGAGATTTCGGTCGCCACGCCACCGTCGC**

**RTDNA:**

**GGTGGCGTGGCGACCGAAATCTCTCGTCCGCGGTGCCGCGAGGCACTCGGTGCGTGCGCGCCGG**
**CTCCGGACGCTGTACCCGATCGATGTGTCGT**

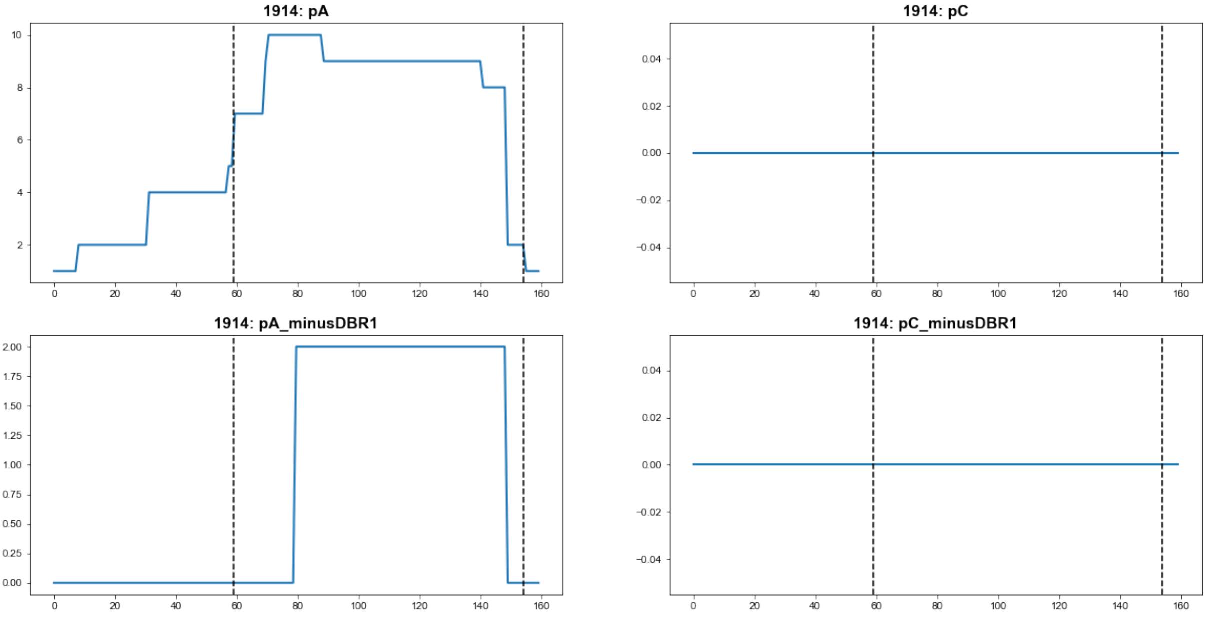


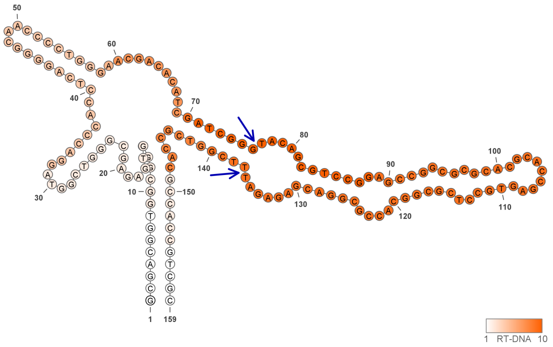


**1914**

### Retron Terminal 1922: Actinobacteria

**Bacterial Vector for RTDNA: None**

**Bacterial Vector for Editing: None**

**Phage Vector for Editing: None**

**Human Vector for Editing: pSLS.1005**

**RTDNA Production (relative to Eco1): undetermined**

**Bacterial Editing: undetermined**

**Phage Editing: undetermined**

**Human Editing (demultiplexed): 0.037297 percent precise (0.321273828x Eco1)**

**ncRNA:**

**GATGCGGCGGCGCGTGGGTGAGAGTTGGTCCACGCGACCAGTCTCAGGGGTAACCCCTGGGTCG**
**GCCAGAAGAATTGGATACTGCGTCCACCGGCGTTCGGCACGCTCCGCGTGCTTGCCGCCGCACC**
**GGACAAAGGATTTGGTCACCCCGCCCCGTCGCATC**

**no RTDNA sequencing data**

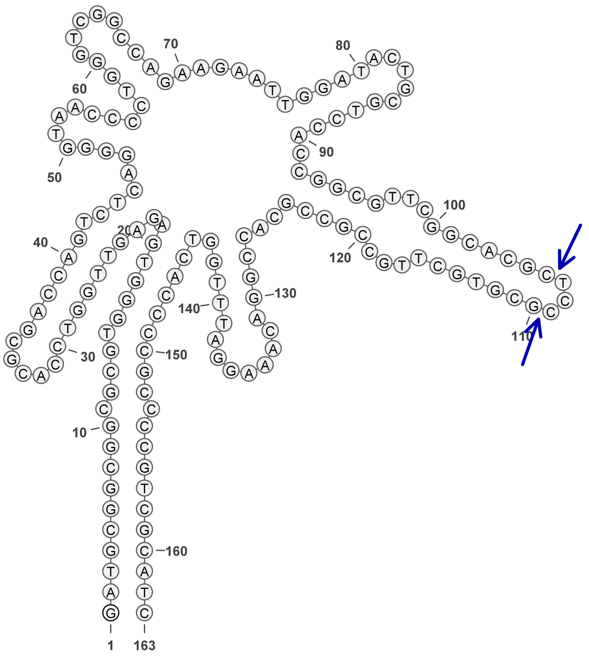


**1922**

### Retron Terminal 1925: Actinobacteria

**Bacterial Vector for RTDNA: pSLS.695**

**Bacterial Vector for Editing: None**

**Phage Vector for Editing: None**

**Human Vector for Editing: pSLS.927**

**RTDNA Production (relative to Eco1): 0.057008498 by PAGE**

**Bacterial Editing: undetermined**

**Phage Editing: undetermined**

**Human Editing (demultiplexed): 0.371421 percent precise (3.199395302x Eco1)**

**ncRNA:**

**GCGTGCAGCGGTGCCGGGGCGAGAGTGGGGTCGACCGGAGTTCGCGGCCGGTCACCCGTCTCG
GGGGCAACCCCCGGGAACGAGAACACAGCCGATCACTGCGTCCGACGCAGGAATGCACGCTCC
GCGTGCTGCGCTGCGCCGCCGACGGGAGAGCATGGTCGCCCCGGTCCCGCTGCCGC**

**RTDNA:**

**GGGGCGACCATGCTCTCCCGTCGGCGGCGCAGCGCAGCACGCGGAGCGTGCATTCCTGCGTCGG
ACGCAGTGATCGGCTGTGTTCTCGTTCCCGGGGGTTGCCCCCGAGACGGGTGACCGGCCGCGAA
CTCCGGTCGACCCC**

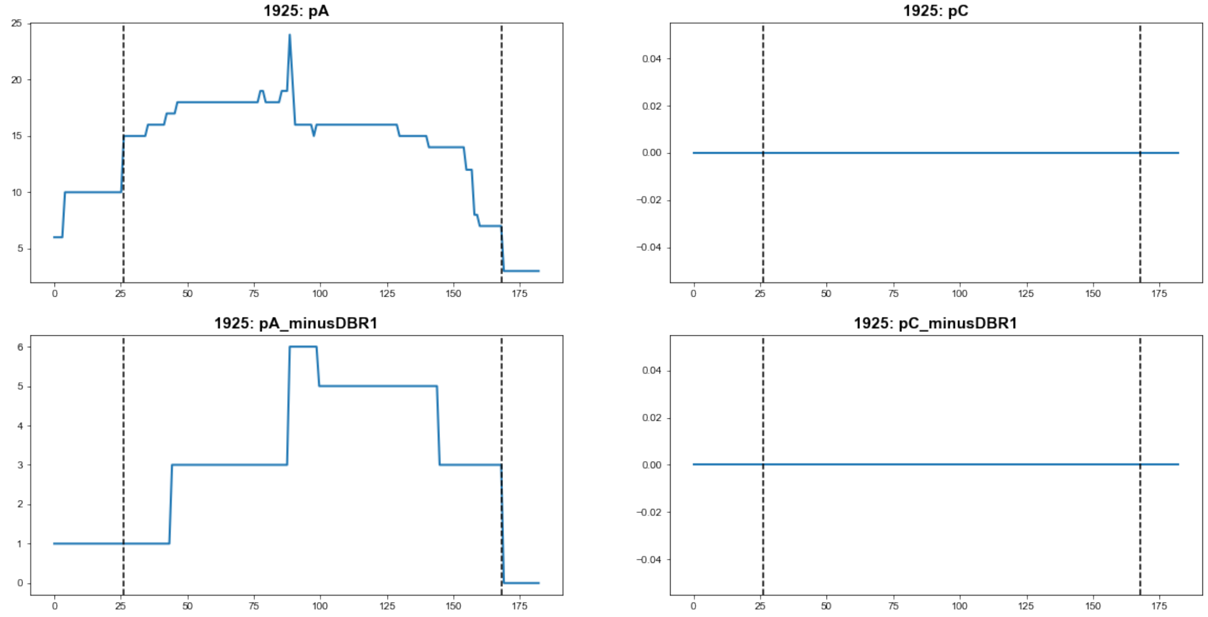


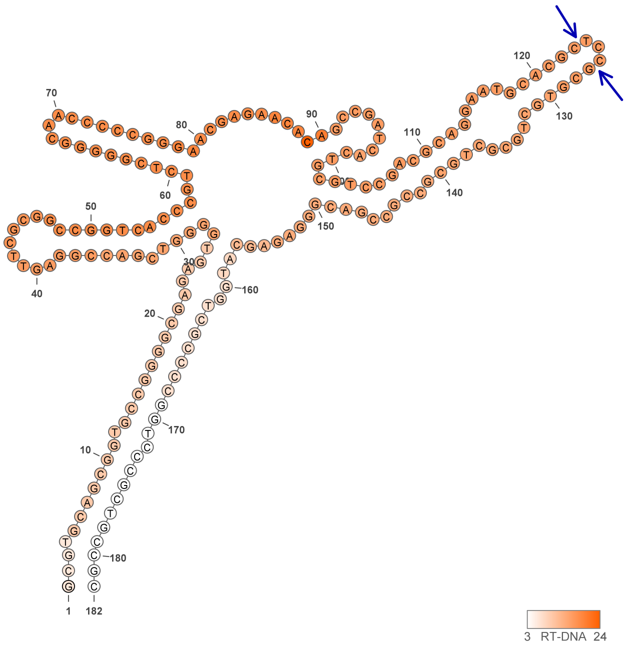


**1925**

### Retron Terminal 1928: Actinobacteria

**Bacterial Vector for RTDNA: pSLS.653**

**Bacterial Vector for Editing: None**

**Phage Vector for Editing: None**

**Human Vector for Editing: pSLS.898**

**RTDNA Production (relative to Eco1): 2.505321343 by PAGE**

**Bacterial Editing: undetermined**

**Phage Editing: undetermined**

**Human Editing (demultiplexed): undetermined**

**ncRNA:**

**GTGCGACGGTGCGGAGGTGAGAGTGGTGTGGCCGGTTCGCCGACCGCGACCACGTCAGGGGTA
ACCCCTGGCTCGACAGTGTTCACCGGAGACACCGTCCGACGCGCGACCGCATGCTCCGCGTGCT
CCGGCGCGCCGCAGACGACAGGTATCGGTCACCTCGCACCGTCGCAC**

**RTDNA:**

**CGATACCTGTCGTCTGCGGCGCGCCGGAGCACGCGGAGCATGCGGTCGCGCGTCGGACGGTGTC
TCCGGTGAACACTGTC**

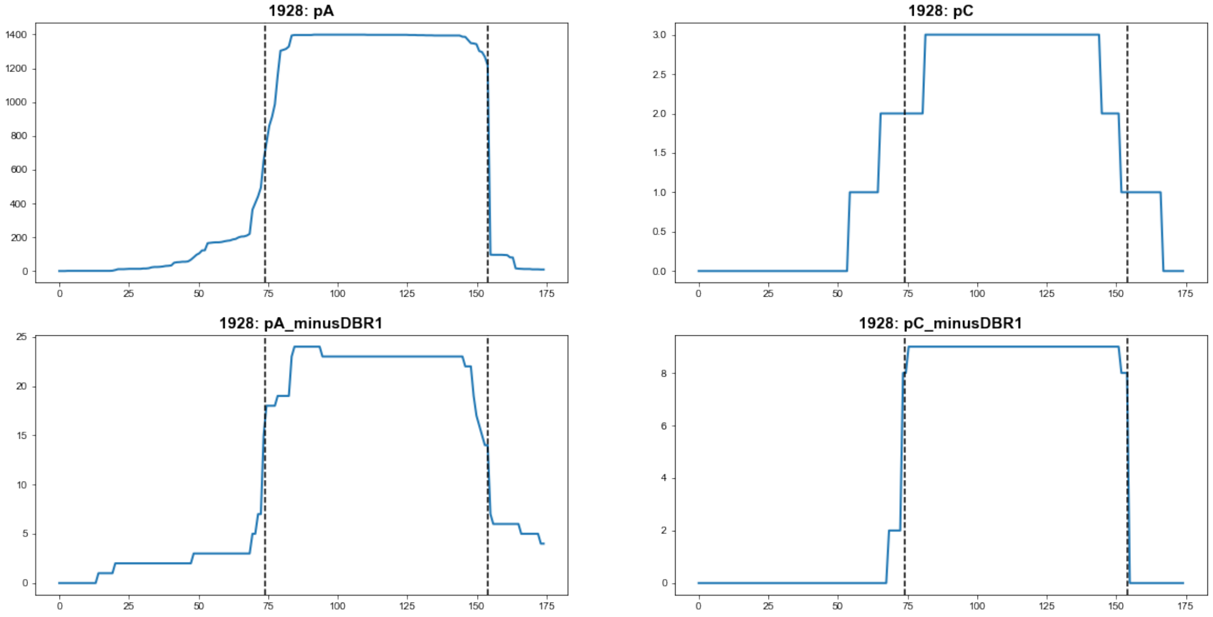


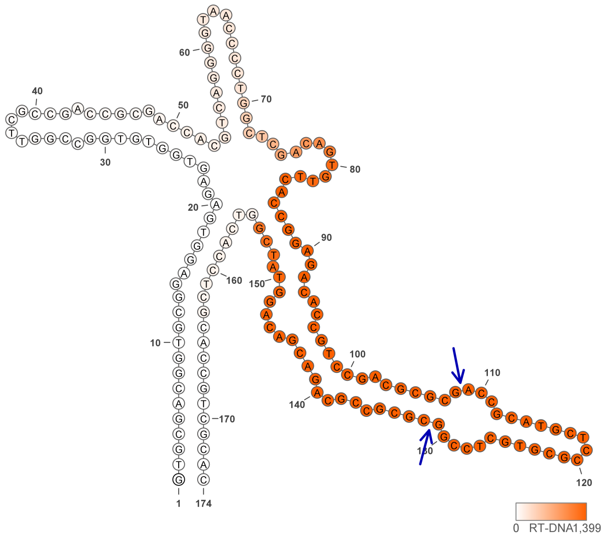


**1928**
